# Rule-Based Deconstruction and Reconstruction of Diterpene Libraries: Categorizing Patterns & Unravelling the Structural Landscape

**DOI:** 10.1101/2024.12.20.629783

**Authors:** Davis T. Mathieu, Nicholas Schlecht, Marvin van Aalst, Kevin M. Shebek, Lucas Busta, Nicole Babineau, Oliver Ebenhöh, Björn Hamberger

**Affiliations:** Michigan State University; Heinrich Heine Universitaet; Northwester University; University of Minnesota Duluth

**Keywords:** Carbocations, Diterpenes, Compound Libraries, Computational Modelling, Rule-Based Analysis

## Abstract

Terpenoids make up the largest class of specialized metabolites with over 180,000 reported compounds currently across all kingdoms of life. Their synthesis accentuates one of natures most choreographed enzymatic and non-reversible chemistries, leading to an extensive range of structural functionality and diversity. Current terpenoid repositories provide a seemingly endless landscape to systematically survey for information regarding structure, sourcing, and synthesis. Efforts here investigate entries for the 20-carbon diterpenoid variants and deconstruct the complex patterns into simple, categorical groups. This deconstruction approach reduces over 60,000 unique diterpenoid structures to less than 1,000 categorical structures. Furthermore, the majority of diterpene entries (over 75%) can be represented by less than 25 core skeletons. Natural diterpenoid abundance was mapped throughout the tree of life and structural diversity was correlated at an atom-and-bond resolution. Additionally, all identified core structures provide guidelines for predicting how diterpene diversity originates via the mechanisms catalyzed by diterpene synthases. Over 95% of diterpenoid structures rely on cyclization. Here a reconstructive approach is reapplied based on known biochemical rules to model the birth of compound diversity. Reconstruction enabled prediction of highly probable synthesis mechanisms for bioactive taxane-relatives, which were discovered over three decades ago. This computational synthesis validates previously identified reaction products and pathways, as well as enables predicting trajectories for synthesizing real and theoretical compounds. This deconstructive and reconstructive approach applied to the diterpene landscape provides modular, flexible, and an easy-to-use toolset for categorically simplifying otherwise complex or hidden patterns.

**Significance Statement:** We take a deconstructive and reconstructive approach to explore the origins of the diterpene landscape. Introduction of a navigational toolset enables users to survey compound libraries in ways formerly uncharted. Their utility demonstrated here, maps out diterpene cyclization routes, critical intermediate waypoints, and guidance for how to arrive at compounds previously off-the-map. Information acquired from these tools may imply the diterpene landscape is vastly unexplored, with the plateau for discovery potentially still out of sight.

## Introduction

Terpenoids are specialized metabolites known for their structural diversity, expansive utility, and vast distribution across the tree of life. They are structurally characterized by the combination of the universal 5-carbon building blocks, isopentenyl diphosphate (IDP) and dimethylallyl diphosphate (DMADP), to form the linear prenyl-diphosphates, which are further converted to form the mono- (C_10_), sesqui- (C_15_), di- (C_20_), sester- (C_25_), tri- (C_30_), tetra- (C_40_) and polyterpenoid classes. Currently, over 180,000 unique terpenoid structures have been reported in the Dictionary of Natural Products (DNP) and TeroKit databases (1–3). Most reported structural diversity originates from various cyclization mechanisms of the linear precursors via terpene synthase (TPS) activity (4–8) and through further modification via oxidative functionalization by NADPH-dependent cytochrome P450 mono-oxygenases, dehydrogenases, 2-oxoglutarate dependent oxygenases, or a range of transferase affording conjugates (7, 9–13). In nature, these compounds function in adaptation and interaction with the environment including defense, pollinator attraction, developmental signaling, and interspecies communication (14–36). The intrinsic role terpenoids serve in communication and defense likely has been a driving force for the observed diversity to date (29, 37–39). Additionally, there is a greater pressure for chemical responses in plants and other stationary organisms unable to run from threats,, leading to greater diversity overall seen in sessile systems (40–42). From a human perspective, terpenes offer broad applications as fuels, pharmaceuticals, nutraceuticals, fragrances, and pesticides (14, 43–57). Examining the diterpene libraries comprehensively instead of individual compounds provides high informational wealth with potential for synthesis elucidation, biochemical discovery, and the uncovering of otherwise hidden patterns.

Work presented here utilizes diterpenes as a case study to showcase development of software to extract patterns that exist within the complexities of a group of chemicals’ structural landscape.

Over 95% of diterpenes are cyclized either by a class II diTPS followed by a class I diTPS, or by a class I TPS alone. Schematics with examples of central terminology and visualized workflow can be found in the Supplemental Appendix. Class II/class I derived diterpenes represent structures with a characteristic decalin-core and are blanket referred to here as *“labdane-derived”*. These compounds are synthesized by an initial protonation at the precursor geranylgeranyl diphosphate (GGDP) tail (class II), which then leads to cationic cycloisomerization and subsequent quenching by deprotonation or hydroxylation. This is followed by the removal of the diphosphate (class I), forming another carbocation that can also cyclize (as in the case of kaurenes) and/or be quenched (as in the case of labdanes) (58, 59). The other most common synthesis, catalyzed by class I TPS, forms a carbocation by removing the diphosphate, which can lead to larger ring formations. This mode of synthesis is referred to here as *“macrocyclic-derived”*. Class II/class I and class I diTPS chemistries that orchestrate diterpene synthesis provide the foundation for most diterpene structural diversity. Here *“backbone”* and *“skeleton”* are used to describe two levels of detail regarding the GGDP-derived structure after cyclization. The term *“backbone*” refers to the portion of the molecule originating from GGDP but with retained information about stereochemistry, bond information, and identification of R-group modifications. “*Skeletons”* refer to the most simplified 20- carbon structures, only considering carbon-carbon linkage and ignoring all other details. Additionally, the simplicity of the skeletons in this work allows it to act as a common reference frame for comparison of all compounds with that shared skeleton. Demonstrated utility of how a common reference frame can be used to orchestrate multi-structural comparison in even greater detail (specific functional groups and cross-skeleton comparisons) and make additional comparative analyses can be viewed with additional context in our companion paper [Babineau et al. 2024].

Studying the immense size and scale of the current terpene landscape requires analysis to be performed in a computational space. This was previously approached with machine learning and rule-based approaches to mirror biochemical reactions (2, 8, 59–65). Machine learning can generate an abundance of compounds and parse out complex patterns within datasets, but also faces limitations where biases, and errors may not be readily detected, affecting interpretation and reproducibility (66, 67). Alternatively, a rule-based approach provides a higher degree of control to the user, grants ease to accommodate a continuously growing database, and emphasizes human guidance when modelling the metabolic landscape. Here, rule-based methods are utilized with Simplified Molecular Input Line Entry System (SMILEs) as inputs, which are the presentation of chemical structures as text in a computational space (68–72). The reaction rules applied are represented in a SMILES Arbitrary Target Specification (SMARTS) format for pattern recognition within compound structures and determining if a reaction is permissible (73–77).

The rule-based methodology here operates on the reported diterpenes in the DNP (>25K; v30.1) and TeroKit (>40K; v2.0) databases to uncover complex patterns regarding diterpene structural diversity, synthesis, and origin. Diterpene synthases catalyze complex, multi-step chemistries derived from diphosphate cleavage, protonation, carbocation rearrangements via nucleophilic attacks, methyl and hydride shifts, and eventual quenching and resolution of carbocations. A deconstructive and reconstructive approach are used here to model diterpene biochemistry targeting the synthesis of diterpene backbones and skeletons. The deconstructive approach isolates and identifies these diterpene backbones and skeletons. The reconstructive approach predicts diterpene backbone and skeleton formation using carbocation reactions mirroring the mechanisms of class II and class I diTPSs (63, 64) (workflow visualized in Supplemental Appendix). This reconstructive modelling provides a unique platform to demonstrate known carbocation rearrangements and stepwise mechanisms for both latent and known diterpene chemistries. Deconstructive and reconstructive paired methods provide a unique synergy for simplifying the abundance of reported compounds, examining the origin of diversity, and predicting mechanisms of synthesis. Positive outcomes illustrated by this test case can be expanded upon further for compound classes beyond diterpenes and has already shown similar capacity in triterpenes (see companion paper) [Babineau et al. 2024].

## Results

### Diterpene database deconstruction summary

Among all reported diterpenes (>60K) we identified a total of 924 topologically distinct C_20_ skeletons within the TeroKit (872 skeletons) and DNP datasets (671 skeletons) (Supplementary Data 2e, 2f; Supplemental Code 1). Compounds with greater or fewer than 20 carbons in their skeletons were not considered as candidates because nearly all of these structures were either misannotated (i.e. sesquiterpenes, triterpenes, alkaloids), further derivatized C_20_ structures that were already identified, or diterpene-diterpene dimers (C_40_). The majority of both databases could be deconstructed back to C_20_ (TeroKit 85.8%; DNP 88.0%), demonstrating that instances of methylation and demethylation are uncommon. The most common non-C_20_ skeletons (representing 10.5% of the TeroKit database) were structures containing C_19_ (2960), C_18_ (691), C_17_ (316), and C_21_ (312) (Supplemental Data 2f). The top 25 most common skeletons represent over 75% of all reported TeroKit diterpenes (∼26,000 compounds; Supplemental Data 2g). Echoed in the findings of previous work (6), monoterpenes (∼60), sesquiterpenes (∼320), triterpenes (∼70), and other compounds (such as alkaloids) were misannotated as diterpenes (∼1.10%; Supplemental Data 2h; Supplemental Code 1).

Skeleton structures from the DNP were compared to visualize compound distribution and similarities (Figure 1). Skeletons grouped largely based on whether or not cyclization occurred, representing principal component 1 (PC1) and 26% of identified skeleton variance, where the linear phytane backbone and other acyclic derivates, are separated from those that were cyclized (Figure 1, Group 1 vs Groups 2/3). PC2 represented 17% of variance and was determined by the mode of cyclization distinguishing labdane-derived structures (Figure 1, Group 2) from macrocyclic-derived structures (Figure 1, Group 3).

**Figure 1.**
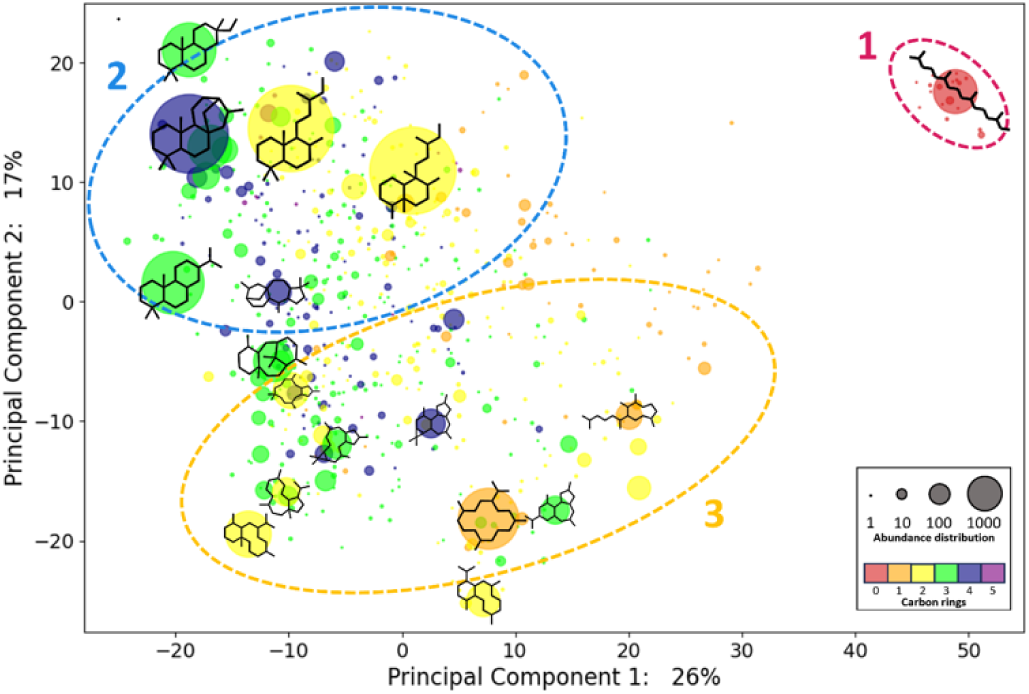
Principal component analysis of deconstructed DNP diterpene skeletons compared based on RDKit bit vector scores. Each point represents a diterpene skeleton in which point size is correlated to the number of reported compounds in the DNP database that deconstructed back to that specific skeleton structure. Color indicates the number of carbon rings present for each structure. PC1 defines 26% variance and is largely characterized by the presence or absence of rings. PC2 defines 17% variance and distinguishes itself based on mode of TPS cyclization. Cluster 1 (circled in **Red**) represents the phytanes, which are the dephosphorylated form of GGDP, the most common diterpene precursor. Cluster 2 (circled in **Blue**) contained the labdane-derived class II/class I compounds. Cluster 3 (circled in **Orange**) contained macrocyclic-derived compounds, synthesized via class I activity.

### Reconstructing diterpene backbone cyclization validates known biochemistries and charts hypothetical space

We modified Pickaxe parameters in order to evaluate whether known diterpene synthesis pathways can be mirrored computationally and to evaluate the total theoretical space possible solely through carbocation rearrangement and diTPS mechanisms seen in nature. Class II and class I carbocation rules were run separately to allow biochemistry to mirror the two main modes of diterpene synthesis (either class II/class I or class I) and to ensure compounds never had more than one carbocation at the same time. The minimum number of steps predicted to produce the complex clerodane (class II/class I product) and dolabadiene (class I product) each required 8 generations. As more generations were included, returns to diversity reached an upper threshold with minimal novel diversity arising. Quantification of class II and class I products generated >347,000 compounds, with 737 structures that matched 80 of the unique skeletons previously identified (Figure 2a). Among the resolved structures, a total of 48,979 hypothetical skeletons can originate solely from carbocation rules observed in nature (Supplemental Data 4) and represent putative undiscovered chemistries.

**Figure 2.**
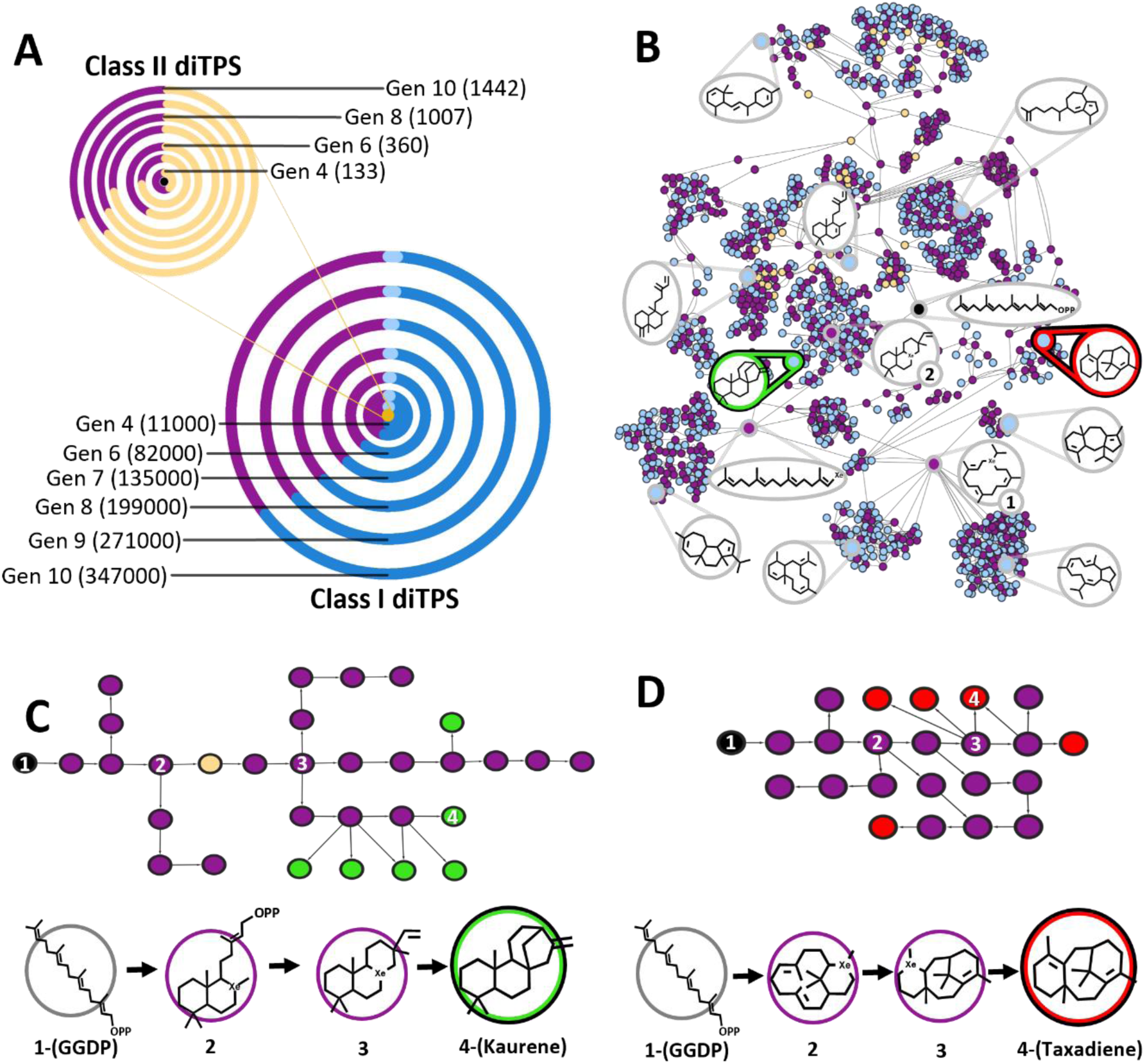
Reconstruction of known and construction of theoretical carbocation cyclization (TPS enzyme) reactions. Structures throughout are indicated as either **Black** (GGDP), **Purple** (carbocation containing), **Yellow** (diphosphate containing and class I precursors), **Blue** (resolved hypothetical backbone), **light Blue** (resolved backbone matching a known structure), **Green** (kaurene), or **Red** (taxadiene). **A** Products of diTPS class II reactions, which created 1442 total structures after 10 generations. The second iteration built upon the resolved class II products that retained the diphosphate (931 (yellow) and GGDP (black)) and were used as inputs, since the removal of the diphosphate is the first step in forming a carbocation in class I cyclization. The second iteration performed diTPS class I reactions, creating >347,000 compounds after 10 generations. **B** A network of filtered reactions exclusive to known structures. Two critical carbocation hubs driving cyclization are labeled for the cembrenyl-cation (1) and pimaranyl-cation (2). Local examples of synthesis are visualized for **C** the class II/class I synthesis of kaurene, and **D** the class I synthesis of taxadiene.

The acquisition and implementation of computational rules varied greatly. In 12,319 occurrences, multiple rules created the same product. This occurred as structures became more complex throughout each generation, and as such opportunities for redundancy in pattern matching increased. Additionally, there were multiple instances where rules intended for specific reactions were applied more commonly than expected, such as the reaction for the secondary beyeranyl carbocation, which is necessary to cyclize pimarane for beyerene, kaurene, atiserene, and trachylobane synthesis (90). This rule was designed that mirrors a precise secondary carbocation reaction (90) but in practice met the necessary conditions hundreds of times. As expected, rules designed to be generic such as those to quench carbocations, hydride shifts, or methyl shifts occurred most often, implemented >254,000, >115,000, and >24,000 times respectively. Rules for the 1,2-hydride and 1,2-methyl shifts, which only required an adjacent tertiary or quaternary carbon respectively, become more common as structures increasingly cyclized.

The 737 synthesized compounds that shared the same skeleton as those identified in the DNP were used to generate a reaction network and predict routes of synthesis of known structures (Figure 2a, 2c, 2d). Cembrene, pimarane, and taxane related carbocations had the highest centrality metric within the network, giving them extensive influence as central hubs in driving downstream synthesis and diversity (Figure 2B; Compounds 1 and 2) (78). Two cembrenyl ion variants had served as two of the most critical hubs, having the first and seventh highest centrality within the network (Figure 2B; Compound 1). The first cembrenyl cation, paralleled previous reaction reports leading to direct synthesis of cembrene A and cembrene C (91, 92). This cembrenyl cation also connects downstream with a wide variety of important macrocyclic diterpenes like taxanes and casbanes, which serve as additional relevant precursors for skeleton diversity, particularly among the taxadiene, cembrene, and casbene derived skeletons (12, 93, 94) (Compound 1; Figure 2b). Additionally, this ion variant provides an important first step in Pickaxe to create some of the most commonly reported marine-life derived skeletons, such as the briaranes and eunicellanes. It is of note that the eunicellanes are synthesized differently in coral and bacteria and variants of both mechanisms are represented here (95, 96). The second critical hub included the pimaranyl-cation, which is necessary to form trachylobane, abietane, atiserene, beyerene, kaurene, iso-pimarane, and cassane, with the second highest statistical significance (Compound 2; Figure 2b).

Cyclization by diterpene synthases contribute to the vast majority of all reported diterpene structures, with the 80 identified skeletons generated exclusively through diTPS action representing 72.6% of TeroKit database entries. Skeleton modifications that utilize additional enzymatic mechanisms after diTPS-cyclization are also known, for example the synthesis of lathryanes and lathryane-derivatives (alcohol dehydrogenase driven ring formation), seco-kauranes and enmeins (cytochrome P450 mediated ring breakage), and abeo-abietanes and tanshinones (cytochrome P450 driven methyl shift) (12, 80, 85, 97). Exploratory rules were implemented to account for additional structures among the diterpene skeleton landscape post-diTPS cyclization. These rules contributed an additional 528 skeletons, which account for a more complete representation of total diterpene landscape diversity (91.4%). These post-cyclization rules further derivatized skeletons through ring breakages (86 compound), alternative ring formations (59 compound), carbon side chains shifting (173 compound), ring and methyl groups collapsing/expanding (35 compound), or a combination of these conditions (255 compound).

### Fingerprinting diterpene backbone diversity from quenching patterns, additional (de)saturation of double bonds, and presence of aromaticity

Carbocation cycloisomerization from diTPS’s prevalently generates tertiary carbocations as they tend to be the lowest energy state intermediates (58–60). Carbocations are generally resolved either through deprotonation to form a double bond or through quenching with water. This leads to post-cyclized backbones having a mass of 272 m/z (C_20_H_32_), 290 m/z (C_20_H_34_O), or 308 m/z (C_20_H_36_O_2_; class II/class I synthesis only), depending on whether zero, one, or two carbocations were resolved with water. In the process of synthesis, labdane-derived compounds generate two carbocation rearrangements, macrocyclic-derived compounds and most phytanes generate one carbocation rearrangement, and some phytanes (like GGDP) generate zero carbocation rearrangement. Consequently, when backbone structures deviate from this formula, additional double bond (de)saturation events, and aromatization events can be predicted (Figure 3).

**Figure 3.**
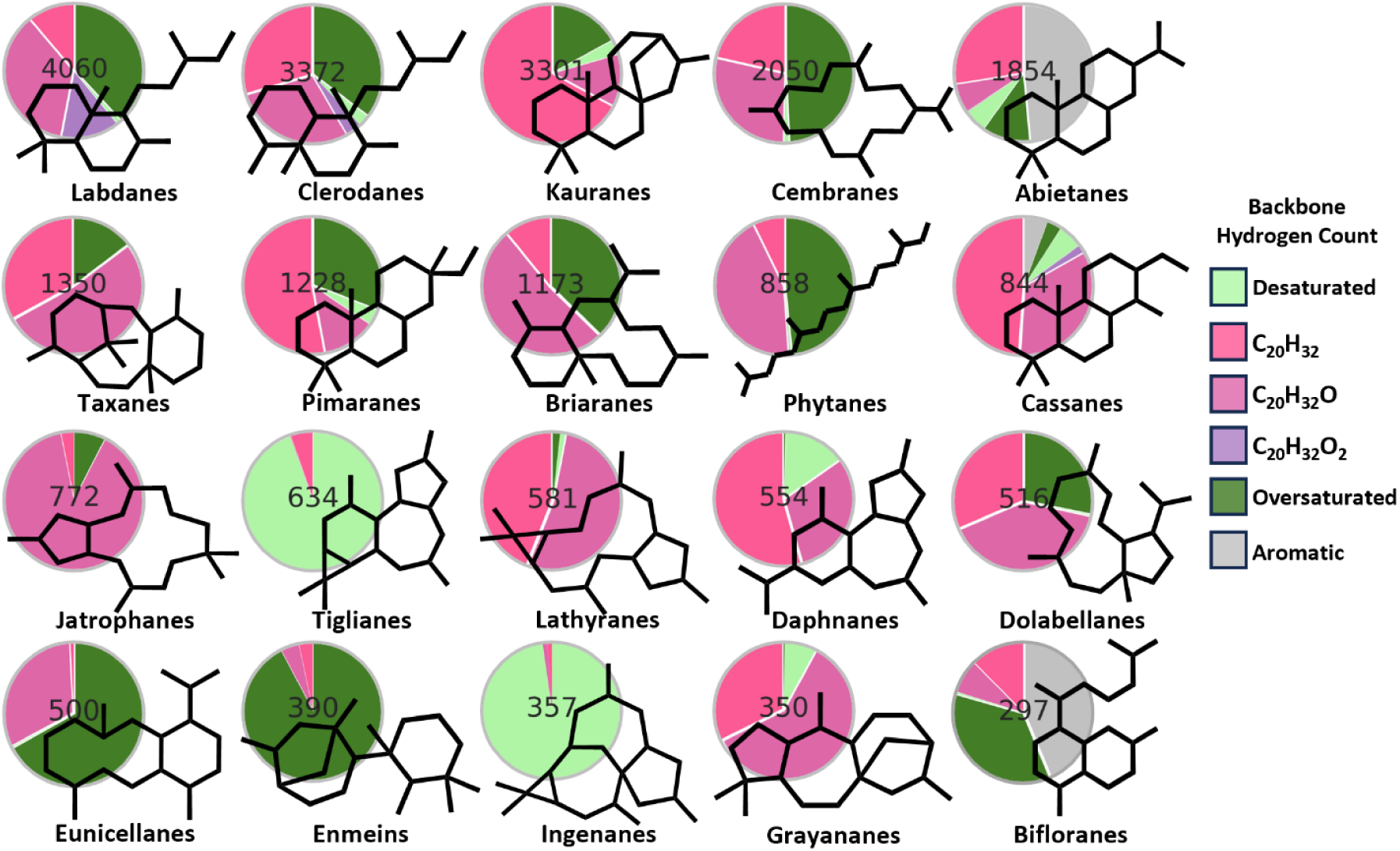
Summary of carbocation quenching patterns and post cyclization decoration for each of the top 20 most common diterpene skeleton classes in the TeroKit database Deconstructed diterpene backbones for each skeleton class were investigated based on carbocation resolution patterns. When a carbocation is generated by a diTPS, protonation or quenching with water will generate a final hydrogen count of 32, 34, or 36 (**Pink**, **Magenta**, or **Lavender** respectively). These in pink-scale represent typical diTPS activity. Occurrences of less than 32 hydrogens suggest additional desaturation to the diterpene backbone after cyclization, leading to the presence of more double bonds than were present prior to diTPS activity (**Mint Green**). When double bonds were absent from a compound and predicted to not be a consequence of diTPS quenching with water, these were estimated as oversaturated (**Forest Green**). Those in green-scale indicate additional modifications to the diterpene backbone after diTPS cyclization. Aromatic structures were identified with **Gray**.

Only three skeleton classes had reported aromaticity, with those being the cassanes, abietanes, and bifloranes, with the latter two being most common (Figure 3). All of these classes can have 1,4-cyclohexadiene, planar moieties that are known to spontaneously aromatize. For the abietanes, miltiradiene can spontaneously aromatize to form abietatriene (82, 83). Likewise, the biflorane derivative, dihydroserrulatene can spontaneously form the aromatic diterpene, serrulatane (7).

Abietanes, cassanes, daphnanes, and especially the tiglianes and ingenanes see desaturation events take place more commonly than in other groups (Figure 3; represented by lime green). Predicted oversaturation was common among most diterpene classes (Figure 3; represented by forest green). The commonality of double bond oversaturation after diterpene cyclization is not surprising however, as alkenes provide reactive centers for downstream decoration. Evaluating diterpene classes in this way allows fingerprinting for identifying distinctions between diterpene families and provides insight into additional decoration taking place post-cyclization.

### Multiple SMILEs alignment of diterpene backbones identifies atomic hotspots

Localized atom and bond diversity within specific diterpene classes was investigated using IQV values for each atom and bond among the top 20 TeroKit diterpene skeletons (Figure 4). Regardless of neighbor or position, labdane-derived compound carbons are decorated at a rate of 20.5% (Supplemental Code 5). The carbon, which neighbored the diphosphate in GGDP prior to cyclization, had reported decoration at a frequency of 56.2%, largely in consequence to diTPS and phosphatase activity (Figure 4). A second hotspot are the two methyl groups attached to the tertiary carbon on the labdane-derived decalin core that are more commonly decorated than other positions among this class (labdanes: 37%; clerodanes: 67%; kauranes: 51%; abietanes: 40%; pimaranes: 42%; cassanes: 42%). This localized decoration is a known product of the CYP701A subfamily and members of the related CYP71 clan, and the CYP720B subfamily of the CYP85 clan (98). The macrocyclic-derived compounds displayed a higher decoration rate of 28.7%. Among bond specific variations, the neighboring tertiary methyl groups saw the highest variation in bond type (Figure 4). Generally, these compounds saw the highest decoration among secondary carbons, with decoration occurring 52.9% of the time. Additionally, many of the macrocyclic-derived compounds had exceptionally high frequency for specific atoms (>95% decoration rate, Supplemental Data 7). Examples of this include the taxanes, jatrophanes, and jatrophane-derivatives such as ingenanes and lathyranes (Figure 4). Coincidentally, highly decorated atoms (IQV>0.79) in the taxane skeleton exactly overlap with those decorated in taxol, while low-decoration atoms (IQV<0.15) align with those not decorated in taxol. Six of these highly decorated taxane atomic sites have also previously been correlated to known locations acted upon by cytochrome P450 families (98).

**Figure 4.**
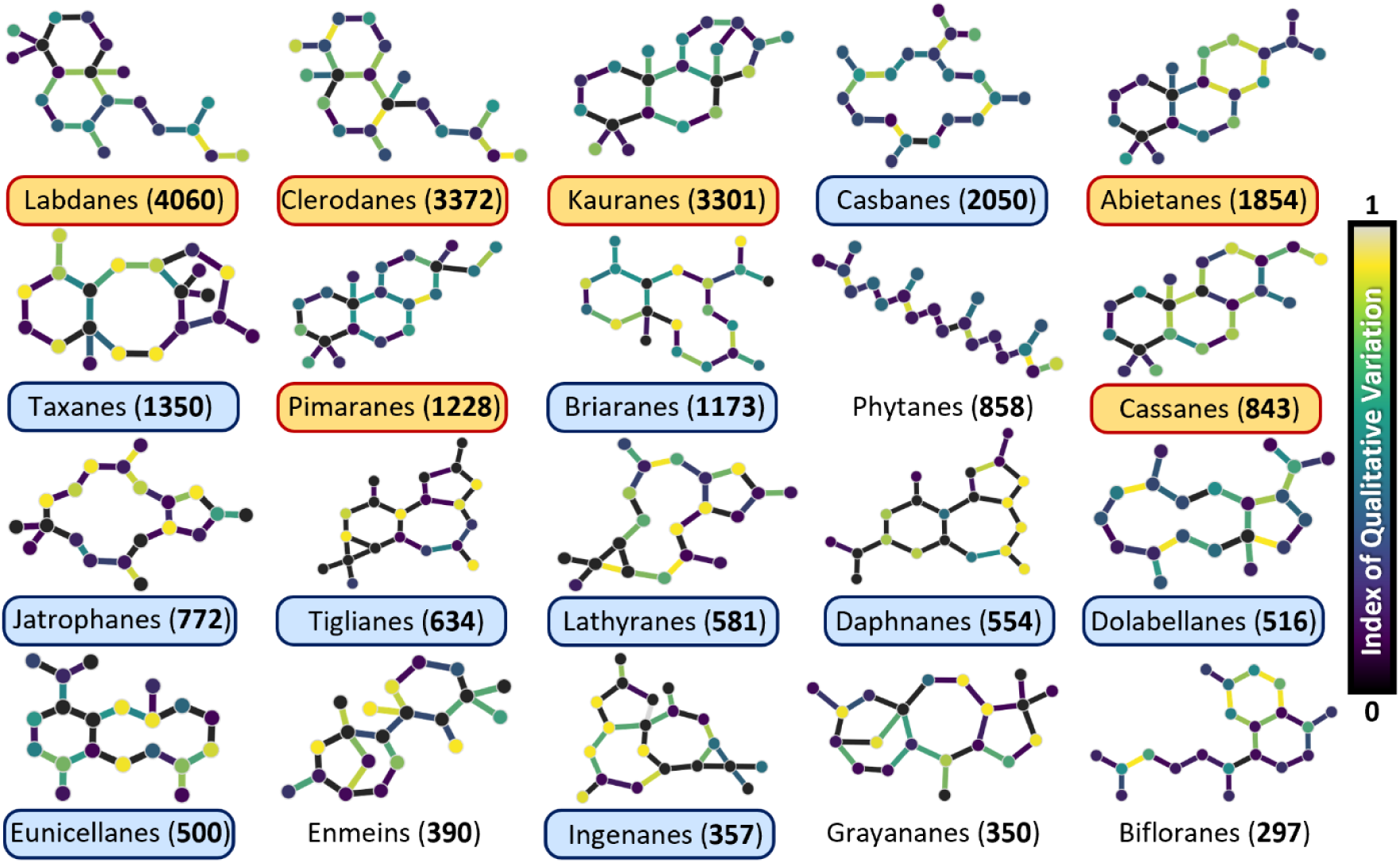
Visual of atom and bond variation among the top 20 most common diterpene skeletons and identified variation mapped to a common reference frame Skeleton names and the number of compounds with that representative skeleton were labeled for each. Compounds were distinguished as either labdane-derived (**Orange**), Macrocyclic (**Blue**), or ambiguous to these distinctions (**White**). Atom and bond variation was calculated using the index of qualitative variation (IQV).

Use of skeletons as a common reference frame, comparison of closely related structures, and alignment of multiple SMILEs for each of the reported diterpene classes allows fingerprinting of molecular families and localized comparison of decoration at an atom-and-bond resolution. When in conjunction with other evidence, this analysis may also aid in predicting synthesis. Here it demonstrates how specific patterns may indicate position-based sources of bioactivity, evolutionary driven effects, steric availability for decoration, and/or localized rigidity for common diterpene cores.

### Bringing to light family diterpene distribution among Viridiplantae, Rhodophyta, and Phaeophytes

The majority of reported diterpenes are sourced from plant life (2, 6). Within the DNP, 76.0% (149/196) of recorded taxonomic families and 76.9% (16,761/21804) of all reported compounds are sourced from Viridiplantae. Because specialized metabolism is often considered to be a potential driving force in speciation (99–103), it is investigated here whether the presence, absence, and/or distribution of compounds correlates with their phylogenetic divergence at a family level. The top 50 most common DNP skeletons (Supplemental Data 2a, 2e), representing 84.5% of all DNP diterpene entries, were used as the first metric, and the 67 land plant families, Charophytes (green algae), Rhodophytes (red algae), and Phaeophytes (brown algae) with at least 10 diterpenes reported were compared (Figure 5).

**Figure 5.**
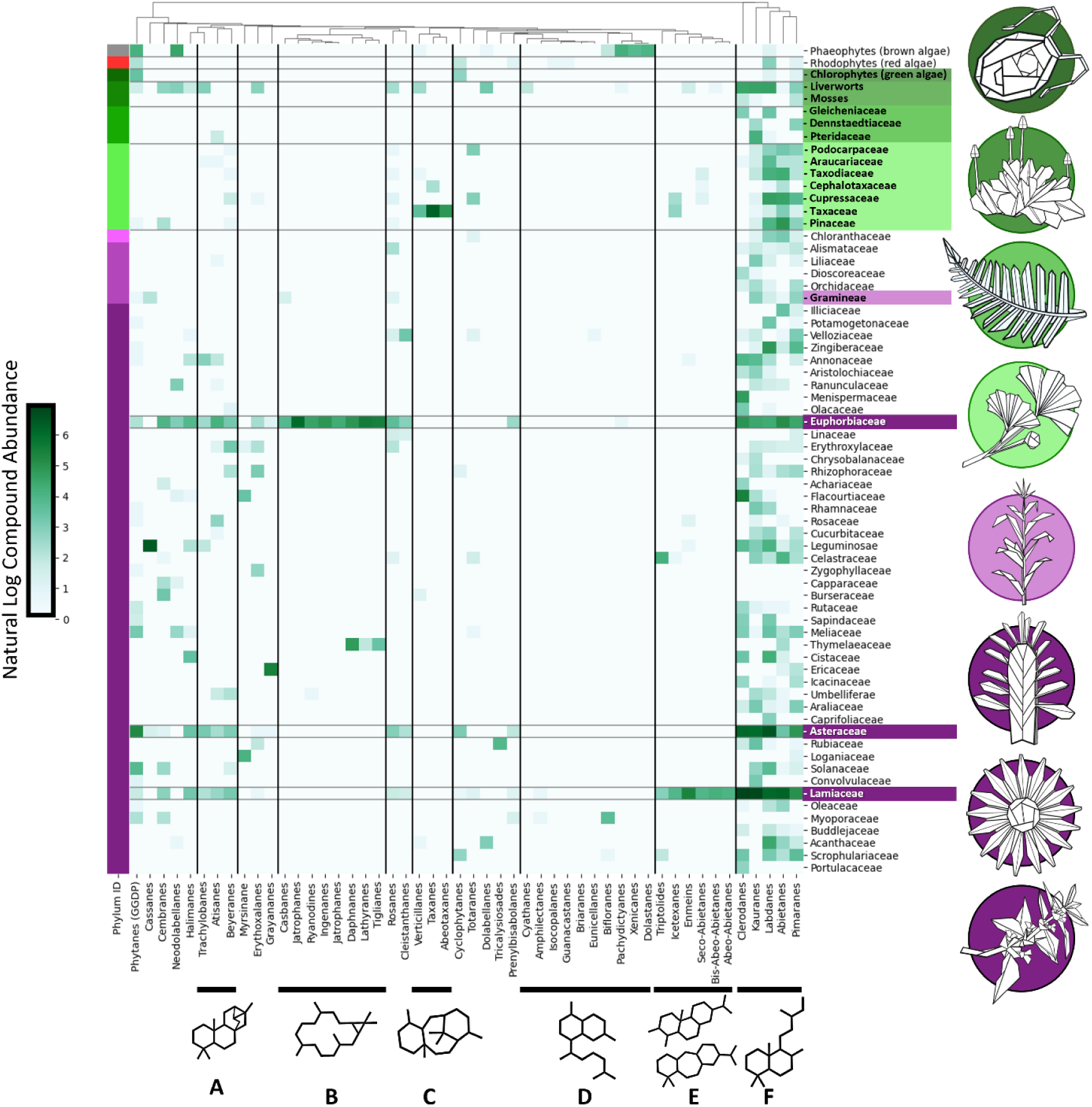
Heatmap of top 50 most common diterpene skeletons and their abundance in plants and algae. Families were manually organized based on phylum divergence in respect to each other (**Grey**: brown algae, **Red**: red algae, **Green (dark to light)**: Charophytes, Bryophytes, Lycophytes, and Gymnosperms, **Purple (light to dark)**: Magnoliids, Monocots, Eudicots). Compounds were hierarchically clustered, which generated groupings of **A** derived skeletons from pimaranes, **B)** cembrene and casbene derived diterpene skeletons in Euphorbeaceae, **C)** taxane and taxane derivates, **D)** diterpene skeletons most commonly found in fungi and marine life, **E)** rearranged abietane derivatives (i.e. abeo-, seco-abietane, and **F)** the most abundant labdane-derived compounds.

Among diterpenes, the Lamiaceae (3,595 diterpenes reported), Euphorbiaceae (2,285 diterpene reported), and Asteraceae (2,073 diterpenes reported) had the highest number of reported compound entries within the dataset (Figure 5). Despite these families being some of the richest in species abundance among plant clades, there appears to be no correlation between family size and the number of distinct diterpenes reported from that family, with Orchidaceae, Fabaceae, and Rubiaceae all being equivalently large with much fewer reported diterpenes. The Taxaceae, Ginkgoaceae, Taxodiaceae, and Cephalotaxaceae, despite being much smaller, also had a relatively high diterpene abundance (Supplemental Code 6, Supplemental Data 8). These results demonstrate that diterpene diversity is family dependent and the evolution of new pathways is different and arises differently in each clade.

Skeletons clustered based on the shared occurrence and/or absence within Viridiplantae families. The first major cluster (A; Figure 5), represents further derivatized versions of the pimarane skeletons, with them largely present in Liverworts and Angiosperms. Next, the Euphorbiaceae dominated reports of macrocyclic structures (B; Figure 5), and represented compounds further derived from the casbane and cembrane skeletons. The following cluster (C; Figure 5) contains compounds tied closely to the Taxaceae family, representing taxane and taxane-derived skeletons. Major compounds most commonly affiliated with fungi, coral, and other marine life (D; Figure 5) are prominently absent among most plant families with only one major exceptions being the Phaeophytes (brown algae), which diverged from Viridiplantae over 1 billion years ago (104). The next major cluster contained rearranged abietane derivates, such as seco-, abeo-, and bis-abeo- Abietanes (E; Figure 5), of which Lamiaceae represented the majority. The last cluster (F; Figure 5) contained compounds widely distributed among all Viridiplantae (8,725 compounds; 39.6%) and represent the major labdane derivatives, including the clerodanes, abietanes, pimaranes, and kauranes. Even though diterpene diversity and expansion are largely family specific, there are still instances of potential convergent evolution visualized here, for example the production of tiglianes in both Thymelaeaceae and Euphorbiaceae (12) and the bifloranes reported in the Australian Myoporaceae family, brown algae, and corals.

## Discussion

Work presented here provides a systematic and comprehensive analysis of the diterpene landscape, introduces a toolset for exploring that space in detail, and illustrates a high potential for remaining novelty among diterpene specialized metabolism yet to be identified. Deconstruction software was essential for extracting complex structures and curating them into manageable groups based on shared characteristics (Supplemental Code 1), a generalizable yet critical foundation for analyzing compound diversity at depth. This took a growing library of >60,000 structures and classified them into just 924 skeleton classes. Our adaptations to the reaction modeling software, Pickaxe, were able to simulate biochemistry leading to the origin of many diterpene skeletons solely through diterpene synthase mediated carbocation reactions, which is the first step in generating diterpene structural diversity (Supplemental Code 3) (63). Compound variance calculation software allowed the extraction and visualization of patterns among diterpene subclasses to further predict origin of synthesis, source of differentiation, and decoration localized to an atom and bond scale (Supplemental Code 4, 5). Lastly, tools were provided to show compound distribution by examining structural similarities and their phylogenetic abundance, which provides an accessible and easily obtainable dissection of the complete diterpene landscape (Supplemental Code 2, 6).

### Deconstruction driven curation of the diterpene landscape

While a huge feat, as metabolic databases continues to grow, information relevant to the user becomes more difficult to navigate. The deconstruction protocols and results presented here have successfully curated, categorized, and simplified the diterpene landscape. The sorting of compound skeletons has also demonstrated clear improvement from previous curation strategies in TeroKit (893 C_20_ scaffolds to 872 C_20_ scaffolds) and previous curation efforts of the DNP (1,092 C_20_ scaffolds to 671 C_20_ scaffolds) (6). While both the DNP and TeroKit databases have generated excellent compound libraries, library size makes manual curation tedious and labor-intensive. Computational deconstruction, however, identifies clear outliers among the libraries such as monoterpenoids, sesquiterpenoids, triterpenoids, alkaloids, and other compounds mislabeled as diterpenoid (approximately 500 entries). The presented indexable and class-separated lists allows identification of structural anomalies and analysis of compounds in unique and informative ways. As a resource, these analyses barely begin to unravel all patterns among diterpenes, with this approach showing strong potential for deciphering future complexities and sets a powerful precedent for navigating other compound libraries moving forward.

### Reconstruction facilitates hypothesis generation and potential for expansive diterpene discovery

Diterpenes are most prominently known for their diversity in cyclization and functional group decoration. Reconstructive efforts performed here modelled the class I and class II/class I diterpene synthase mechanisms observed in nature. The 80 skeletons produced solely through carbocation driven rearrangement represents 85% of all reported structures. Further analyses with this model also led to the identification of specific biosynthesis hubs, which serve as prerequisites for a large proportion of diterpene diversity. Especially notable identified intermediates here included cembrenyl-ions, taxanyl-ions, and pimiranyl-ions.

In order to provide a concrete example of how this model can be used, a highly probable map of synthesis mechanisms for common and uncommon structures related to taxanes were predicted (Figure 6, Supplemental Data 9). These mechanisms generated taxadiene, abeotaxane (105, 106), harziane (107, 108), and 3,11-cyclotaxane (109–111). While harzianes are only reported to be synthesized in coral-derived fungi, the mechanisms predicted here only slightly branches from core taxadiene synthesis (112, 113). Additionally, 3,11-cyclotaxane synthesis involves only one extra carbocation rearrangement than taxadiene synthesis and in nature is likely a consequence of a single additional enzyme. These findings demonstrate the capacity for our methods to not just categorize known diterpene synthesis but also the strength this approach has for hypothesis generation in predicting synthesis in unknown cases as well (Figure 6). When this model is not bound to reported structures, it illustrates that carbocation rearrangement reactions alone have high potential for creating diterpene skeleton diversity. Approximately, 49,000 unique skeletons were generated here, indicating an expansive pool of alternative structures to just the 924 skeletons identified in the database (Supplemental Data 2d, 2e, 2g). While likely an overestimation of products that occur naturally, this estimate may provide a theoretical plateau for diTPS driven skeleton diversity. This also suggests a high probability that additional structures exist and have yet to be found in nature or can be expanded upon through intervention via biosynthetic engineering, like enzyme evolution, and/or use of non-native promiscuous enzymes, which is a phenomenon not uncommon to terpene synthases (10, 63, 64, 114–117). The magnitude difference among reported structures and those predicted here suggests the terpenoid landscape still has novelty to offer and there still remains much to be discovered.

**Figure 6:**
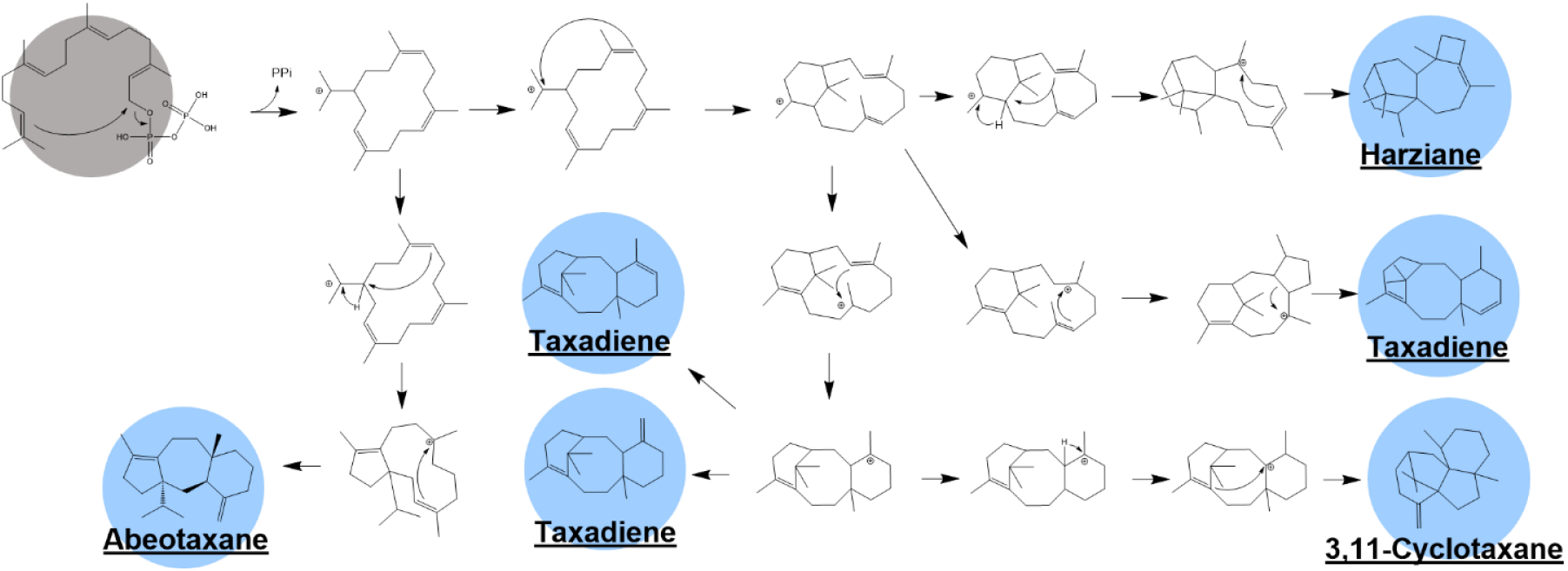
Abridged synthesis for taxane related structures, Taxadiene, Abeotaxane, Harziane, and 3,11-Cyclotaxane. Universal precursor GGDP (in **Grey**) provides the starting point followed by carbocation and cyclization cascades leading to the various final structures (in **Blue**). All intermediates and full predicted mechanisms can be viewed in Supplemental Data 9.

## Materials and Methods

For a full description of the methods, visual flowchart of software and data, and illustrated key terminology, see the Supplemental Appendix. All software and data presented in this manuscript can also be found at https://doi.org/10.5061/dryad.ksn02v7cb

### Deconstruction of the DNP and TeroKit libraries

Diterpene compounds from the DNP database^1^ (v30.2) (Supplemental Data 1a) and parsed datasets for diterpenes, sesquiterpenes, and triterpenes (Supplemental Data 1b-d) from the TeroKit molecule database^2^ (v2.0) were used as initial inputs for analysis (2, 65). Reported SMILEs from each dataset were used to deconstruct and isolate core terpene structures from each molecule (Supplemental Code 1) (74). Core structures were isolated by iteratively removing decoration of non-diTPS origin that were inherently non-terpene in origin. This methodology produced two derivatized models, diterpene backbones and diterpene skeletons, which served as common reference frames in downstream applications (see Supplemental Appendix). Backbones replaced where decoration was originally present with hydroxylation and retained stereochemistry and pi bonds when present. Diterpene skeletons were exclusively based on carbon-carbon sigma bonds, discounting all side-chains and stereochemistry. Diterpene backbones, skeletons, and the skeleton carbon numbers were appended to the original input files (Supplemental Data 1a, 1b) to create more comprehensive and categorical datasets for downstream analyses (Supplemental Data 2a- d). Skeleton abundance summaries for the DNP and TeroKit diterpenes were generated based on shared structures (Supplemental Data 2e, 2f). TeroKit skeleton abundance was visualized as a histogram to show the diversity distribution and compound abundance per skeleton (Supplemental Data 2g). The distribution of deconstructed carbon numbers was used to visualize deconstruction pathway effectiveness, detect database entry errors, and identify apo-terpenoids (Supplemental Data 2h, 2i).

The DNP diterpene skeletons were used for structural comparison (Supplemental Data 2e). All skeletons were converted to bit vectors, and an all-vs-all comparison matrix was generated with the Tanimoto coefficient of structures used as the metric for similarity (78). This metric measured the similarity between two molecules by comparing the ratio of bond intersection and cyclization to the retention of shared atomic structures. This data was visualized as a matrix, heatmap, and principal component analysis (PCA) (Figure 1; Supplemental Code 2; Supplemental Data 3).

### Reconstructing diterpene carbocation reactions in Pickaxe

The modelling software, Pickaxe (63), was modified for execution in bash and with custom rulesets (Supplemental Code 3a Supplemental Data 4). Pickaxe was executed three separate times with different rulesets and inputs to model diTPS activity, which were 1.) class II diTPS reactions 2.) class I diTPS reactions and 3.) combined class II/class I filtered to known TeroKit (Supplemental Data 5a-5d). The first Pickaxe execution used GGDP as the initial substrate, and only used class II diTPS reactions specifically excluding diphosphate cleavage and macrocyclic specific rules (Supplemental Data 5a). This Pickaxe run implemented 10 generations, which loaded in all reactants, applied all applicable reaction rules to compounds, produced new products (1 generation), then all products that were produced would be used as reactants for the next generation. The second execution also underwent 10 generations of reactions, used GGDP and class II products obtained from the first iteration as input compounds, and applied the class I diTPS reaction ruleset (Supplemental Data 5b). The unfiltered Pickaxe runs 1 and 2 represented the full reaction capacity of the model (Figure 2a). The third Pickaxe execution with filtered products identified from both Pickaxe and TeroKit were used as the framework for modeling known diterpene reactions (Figure 2b; Supplemental Code 3c; Supplemental Data 2f, 5c, 6). Results from the third run were visualized as a network in Cytoscape (v3.10.1) (Supplemental Data 6) (78). Kaurene and Taxadiene served as case studies to evaluate and visualize specific pathways (Figure 2c, 2d; Supplemental Data 6) (79). The Cytoscape network statistics for stress centrality (defined as the importance of a node based on how many shortest paths pass through it) and betweenness centrality (how often a node acts as a bridge along the shortest path between two other nodes) were used to identify highly important central hubs, driving diterpene biosynthesis (Figure 2; Supplemental Data 6).

While all known cyclic diterpenes require activity from diterpene synthases, with most reported structures only requiring diTPS activity, there remain known cyclizing modifications driven by alcohol dehydrogenases and cytochromes P450 (among others) (12, 80, 81) that lead to additional reported skeleton diversity. As a next step, an exploratory ruleset was implemented for 3 generations to identify what modifications could be involved leading to the synthesis of skeletons left unpredicted by the diTPS exclusive model. These rules broke carbon rings, created carbon rings, expanded/collapsed connecting rings, and shifted carbon side chains (Supplemental Data 5d).

### Fingerprinting diterpene backbone diversity from quenching patterns, additional (de)saturation of double bonds, and presence of aromaticity

The 20 most common TeroKit diterpene skeletons (Supplemental Data 2b, 2f) were used to fingerprint each compound class for specific quenching patterns (with or without water), post cyclization double bond (de)saturation modifications, and aromaticity (Supplemental Code 4). Deconstructed backbones were used as input for determining (de)saturation and quenching events by counting final hydrogens. If a diterpene is only acted upon by diTPS enzymes it will have a final molecular formula of either C_20_H_32_ (H_32_; 272 m/z), C_20_H_34_O (H_34_; 290 m/z), or C_20_H_36_O_2_ (H_36_; 308 m/z; class II/class I only) to reflect whether 0, 1, or 2 carbocations were quenched with water during cyclization. As an additional check that the hydrogen count was caused by the quenching of a carbocation with water, the expected number of quenched carbocations also required hydroxyl groups to be found at tertiary carbons. Otherwise, those structures were considered as oversaturated after diTPS cyclization. Also, because 2 carbocations are generated in class II/class I synthesis but only 1 carbocation is generated in class I only synthesis macrocyclic structures and phytane were also considered oversaturated if a backbone had 36 hydrogens. When there are fewer than 32 hydrogen atoms, it implies that additional desaturation events took place after cyclization. Likewise, when more than 36 hydrogen atoms are present, post cyclization saturation events are likely. Macrocyclic diterpenes and phytanes are acted upon by one class I diTPS; therefore, in these cases a hydrogen atom count above 34 would also imply post cyclization modification via saturation. If a compound was aromatic, it was labeled as “aromatic” and separated from quenching/saturation counts. Such aromatization is known to occur spontaneously, favoring a more stable conformation, in such cases as miltiradiene (82, 83)

### Determining atomic variability, decoration, and bonds among diterpene skeleton entries

Previously deconstructed TeroKit backbones and skeletons (Supplemental Data 2b, 2f) were used to evaluate atomic variability among the 20 most common diterpene classes (Figure 4; Supplemental Data 7). Canonical SMILEs were used as references to align all atoms and bonds within each class (Supplemental Code 5). To accomplish this, each compound within its respective skeleton class needed to be 20 atoms total (disregarding hydrogen). Previously identified TeroKit backbones (Supplemental Data 2b, 2f) were used to accomplish this by replacing any instances of hydroxyl groups and their connected carbon with xenon to track where decoration was present originally and convert structures to a final atom count of 20. Stereocenters between atoms were also used as a metric for measuring bond diversity. Final alignments were formatted paralleling that of a multiple sequence alignment used with proteins and DNA sequence (Supplemental Data 7).

Canonical references were converted to an edge (bond) and node (indexed atom) table to indicate which atoms were connected to each other and how often they differed (Supplemental Code 5). Bond and atom variability was calculated using the index of qualitative variation (IQV) scores (Figure 4) (84). Using IQV as the metric for variability was decided due to the categorical nature of atom decoration (presence/absence) and bonds (stereochemistry and pi bonds presence/absence). This showed how evenly or unevenly any point within a molecule exhibited variability throughout the dataset. Occurrences of atom decoration were assumed to always be a source of variation. Two separate IQV tables were generated for nodes/edges and were visualized in Cytoscape (v3.10.1) (Figure 4; Supplemental Data 7) (79).

Bias based on carbon position (primary, secondary, or tertiary) was investigated for the labdane- and macrocyclic-derived compounds. This was accomplished by calculating the percent diversity of each atom and determining their neighboring carbon positions for one degree of connection (Supplemental Code 5). Positions were averaged among all occurrences within that class and compared only when more than three occurrences of position/neighbor combinations were present (Supplemental Code 5; Figure 4).

### Phylogenetic distribution of diterpenes within Viridiplantae, Rhodophyta, and Chromista

The DNP skeleton abundance and sourcing (Supplemental Data 2a, 2e) examined compound distribution among land plants, green, red, and brown algae (Supplemental Code 6). Reported taxonomic families were extrapolated to include phylum and kingdom classification. Skeleton occurrences per family were quantified, disregarding families with fewer than 10 entries. The remaining families were manually organized based on their phylogenetic divergence and the 50 most common diterpenes were hierarchically clustered and annotated based on external and DNP reports^3^ (85–89). A scatterplot illustrating diterpenes per family and number of species within each family was also generated (Supplemental Code 6; Supplemental Data 8).

## Supporting information

Zip of all Code used

Zip of all data and Metadata

## Acknowledgments

DM, and BH acknowledge support for this project from the NSF Dimensions of Biodiversity Grant (DEB 1737898) and the NSF-funded doctoral student training grant Integrated training Model in Plant And Compu-Tational Sciences (IMPACTS; DGE-1828149). DM acknowledges financial support from the Jeff Schell Fellowship provided by the Bayer Foundation for in-person travel and collaboration with Heinrich Heine Universität (JS-2022-029). BH acknowledges funding from the Great Lakes Bioenergy Research Center, U.S. Department of Energy, Office of Science, Office of Biological and Environmental Research under Award Number DE-SC0018409 and support from the Department of Biochemistry and Molecular Biology startup funding and support from AgBioResearch (BH=MICL02454). BH gratefully acknowledges support from the MSU James K. Billman, Jr. MD endowment. OE is funded by the Deutsche Forschungsgemeinschaft (DFG) under Germany’s Excellence Strategy EXC 2048/1, Project ID: 390686111. MvA is funded by EU’s Horizon 2020 research and innovation program under the Grant Agreement 862087. We collectively acknowledge that Michigan State University occupies the ancestral, traditional, and contemporary Lands of the Anishinaabeg – Three Fires Confederacy of Ojibwe, Odawa, and Potawatomi peoples. In particular, the University resides on Land ceded in the 1819 Treaty of Saginaw. We recognize, support, and advocate for the sovereignty of Michigan’s twelve federally- recognized Indian nations, for historic Indigenous communities in Michigan, for Indigenous individuals and communities who live here now, and for those who were forcibly removed from their Homelands. By offering this Land Acknowledgement, we affirm Indigenous sovereignty and will work to hold Michigan State University more accountable to the needs of American Indian and Indigenous peoples.

## Author Contributions

DM, MvA, OE, and BH designed the research; NS and BH provided oversight and guidance on all biochemistry; MvA and OE provided oversight for computational modeling; KS provided tutorial, suggestion, and troubleshooting for Pickaxe; DM, NS, and MvA wrote code for visualization and adaptation of Pickaxe for diterpene synthesis; NS and DM designed SMARTs rules for diterpene synthesis; DM wrote and implemented exploratory cyclization SMARTs rules; MvA wrote script for conversion of Pickaxe result output to network data; NS implemented and oversaw all diterpene synthase Pickaxe runs and conversion to network in Cytoscape; NS provided visualization of all SMARTs rules in supplemental; NS and DM generated taxanes related mechanisms supplemental from Pickaxe results; NS provided the dictionary of Natural compounds database file; NS and DM extracted and prepared TeroKit database files; LB and NB provided initial idea and suggestions for atomic hotspot analysis; LB suggested additional analyses for QC of atomic hotspot based on carbon position and bias check related to plant family size. NS and BH edited the manuscript; DM wrote all scripts for deconstructing terpene libraries, cross-structure comparison with PCA, analyzing diterpene saturation and quenching patterns, diterpene class atomic hotspot analysis, and phylogenetic analysis. DM generated all the figures. DM wrote the paper.

## Access and Organization of Supplemental Data

Dryad link for Supplemental Data & Supplemental Code: https://doi.org/10.5061/dryad.ksn02v7cb

## Supplemental Data

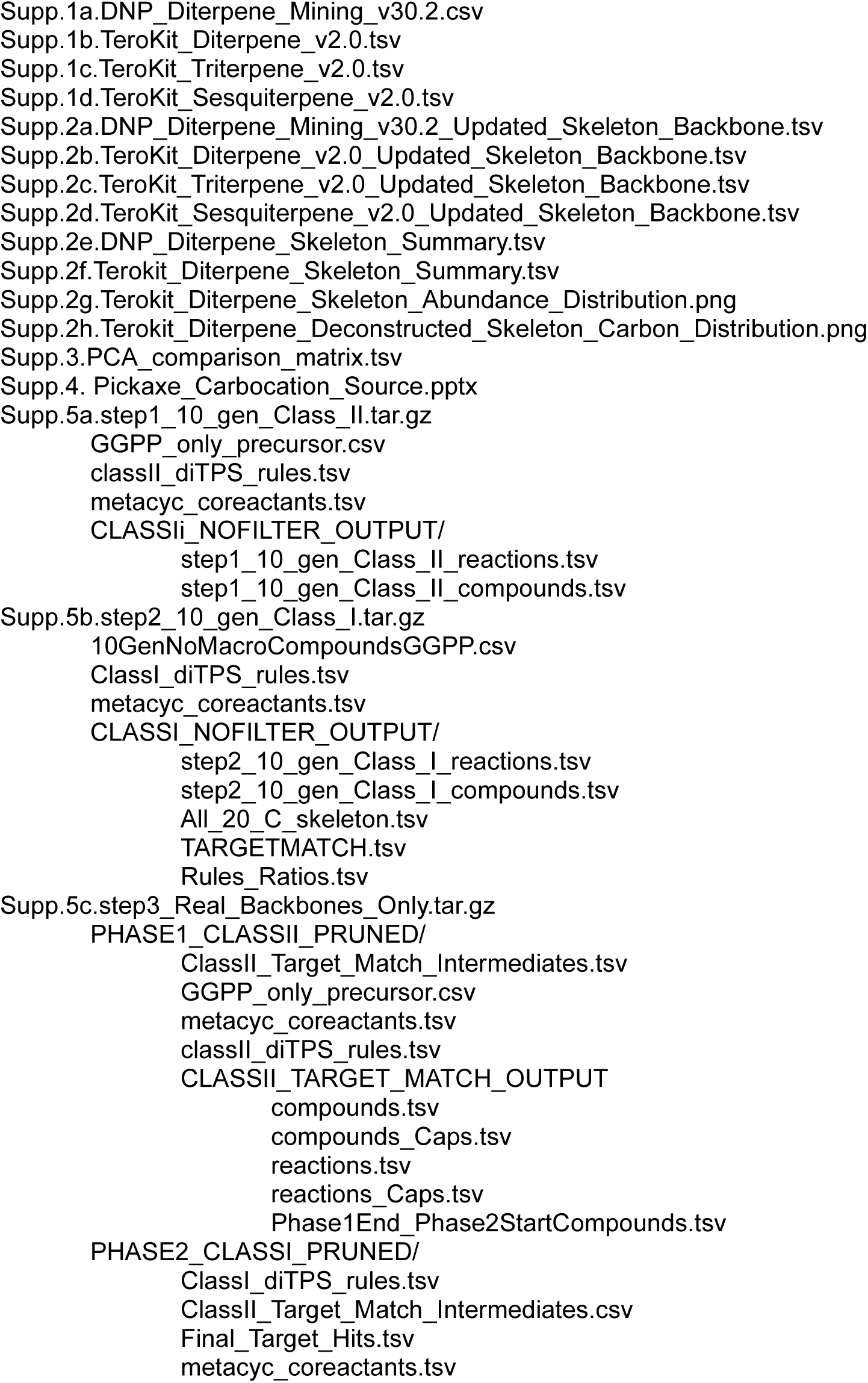
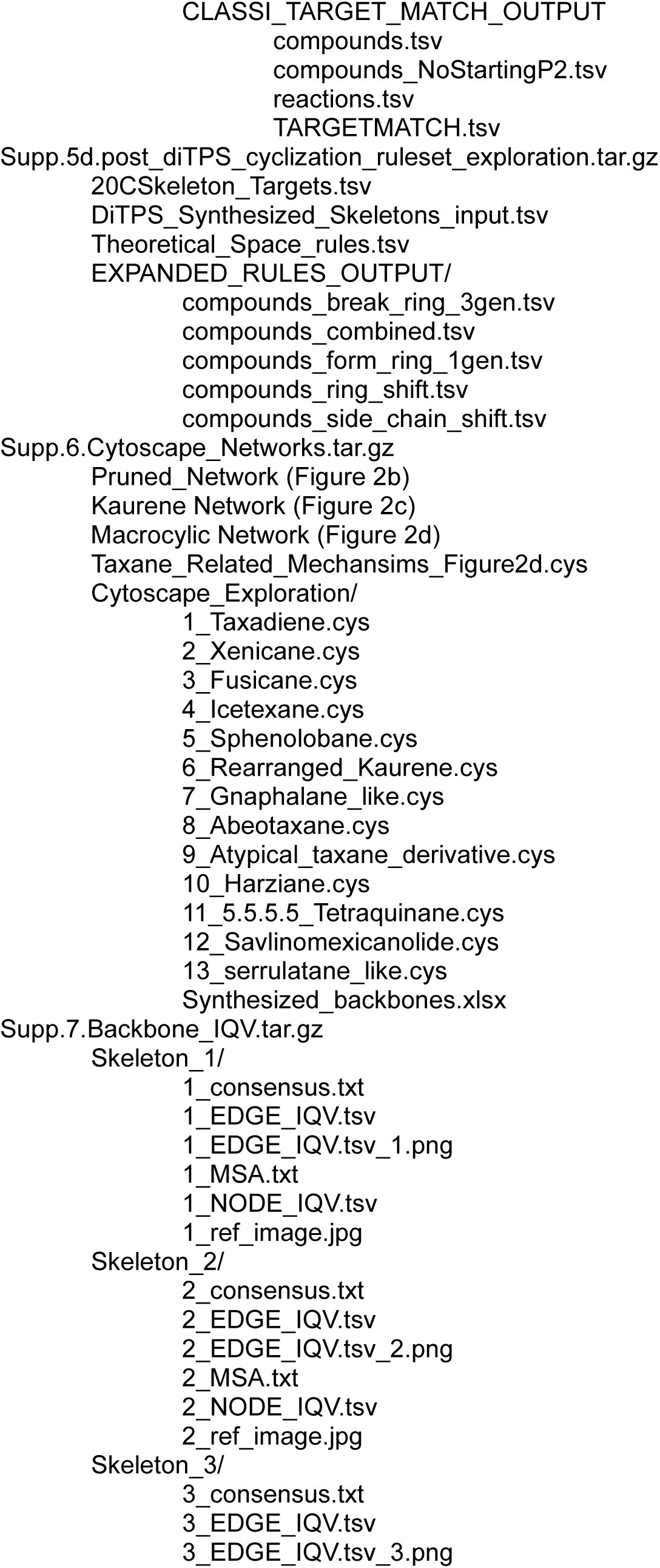
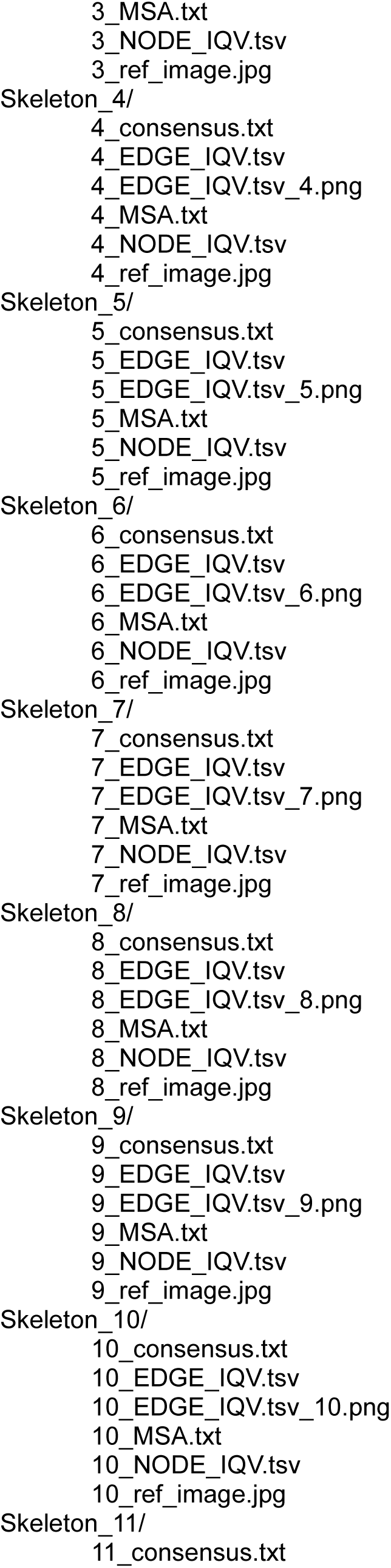
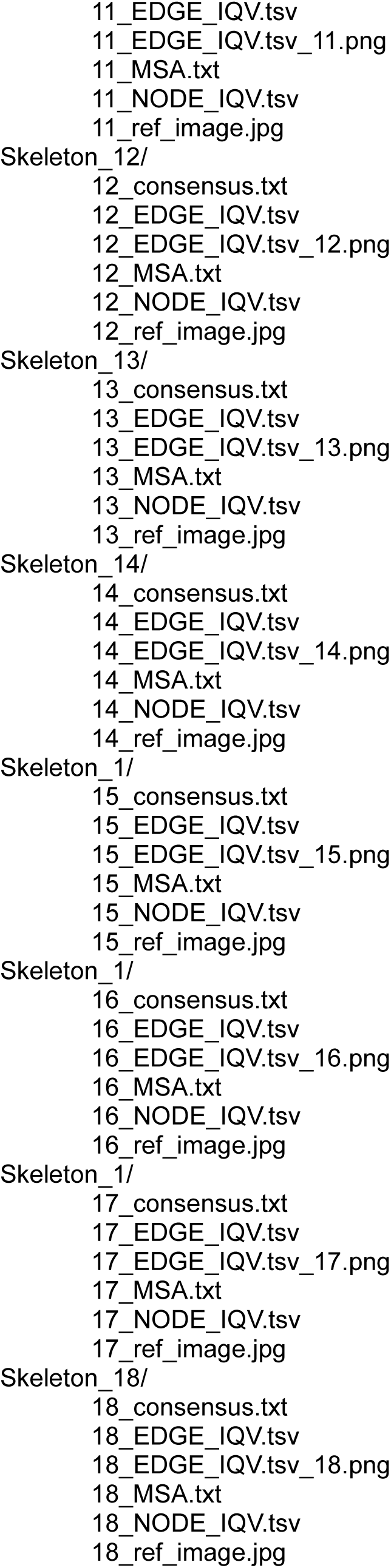
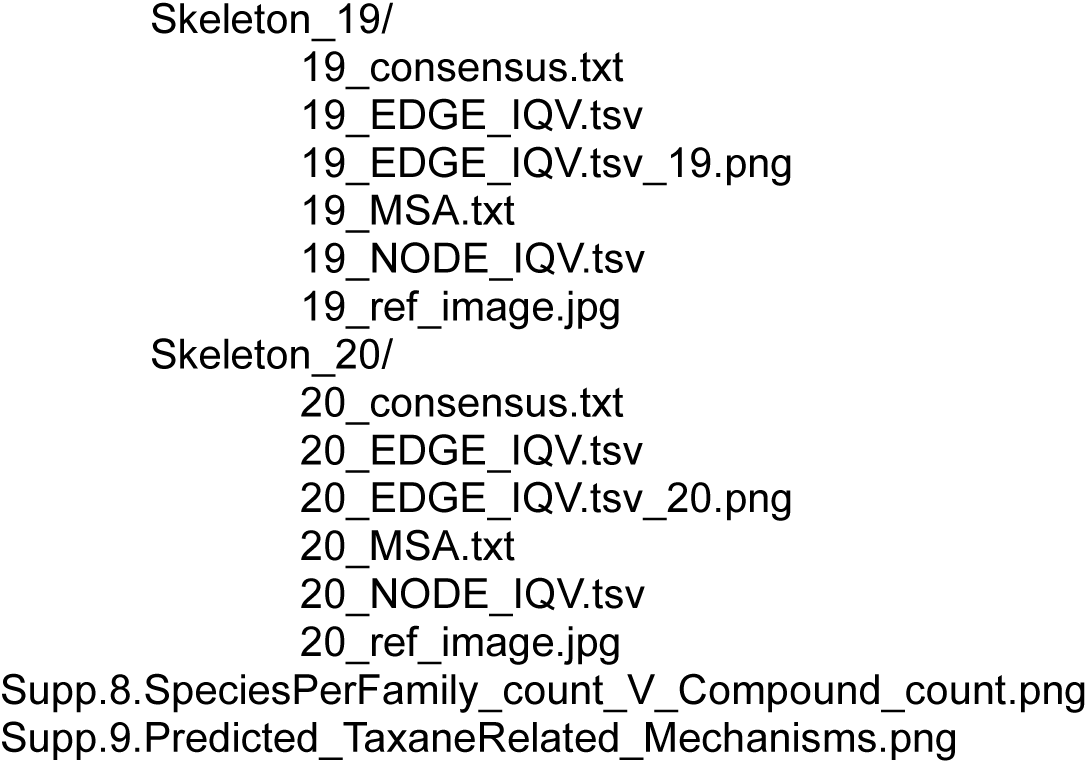

## Supplemental Code

Supp_Code.1.Terpenoid_Deconstruction.ipynb

Supp_Code.2.Skeleton_PCA.ipynb

Supp_Code.3a.Pickaxe_DM_NS.py

Supp_Code.3b.Match.py

Supp_Code.3c.Network_maker.py

Supp_Code.4.Carbocation_Quench_Predictor.ipynb

Supp_Code.5.Backbone_MSA.ipynb

Supp_Code.6.Phylogenetic_Skeleton_Abundance_Heatmap.ipynb

## Supplemental Appendix

### Deconstruction of the DNP and TeroKit diterpene libraries

The DNP database^4^ (v30.2) was queried within the rubric ‘Type of Compound Words’ for “*\*diterpen\**”. Database hits were downloaded, extracted, and concatenated, collecting information on Chemical name, formula, weight, SMILE, InChi, compound type, and biological source for each compound (Supplemental Data 1a). The TeroKit molecule database^5^ (v2.0) was downloaded and subsequently parsed into “sesquiterpene”, “diterpene”, and “triterpene” datasets using the grep command (Supplemental Data 1b-1d) (1, 2). These datasets included TeroKit compound ID, molecular formula, InChi, and SMILEs. SMILEs from each datasets were used as input for deconstruction. Python (v3.9) was used with libraries, pandas (v1.5.1), numpy (v1.25.0), re (v2.2.1), matplotlib (v3.7.1), and RDKit (v2022.09.1) (Supplemental Code 1) (3). Formatted SMILEs from each database were subsequently and iteratively deconstructed to backbones and skeletons using SMART reaction rules. The first function removed any portion of the molecule containing: boron, halogens, silicon, phosphate, sulfur, selenium, tin, fatty acids, saccharides, coumarin, nitrogenous bases, other nitrogen containing R groups, ester linked R groups, ether linked R groups, and additional non-skeleton derived carbon side chains. Instances of isotopic carbon or charged carbon atoms were also converted to a neutral ^12^C. The output returned a backbone with any R-group substituted with an alcohol moiety. Bond and stereochemical variation was retained, along with a list of each step taken to deconstruct the original compound. These backbones were then flattened to a carbon skeleton by converting all covalent bonds to single bonds and removing hydroxyl groups, resulting in exclusively carbon-carbon connections. The original input SMILEs from DNP and TeroKit (Supplemental Data 1a-1d) were processed to include Backbone, Skeleton, and Carbon Number (Supplemental Data 2a-2d). The reported skeletons and their abundances were used to generate DNP (Supplemental Data 2e) and TeroKit summaries (Supplemental Data 2f). Deconstruction of sesquiterpene and triterpene databases were performed as proof-of-functionality and to support robustness of programs (Supplemental Data 2c, 2d). Representation of TeroKit skeletons was visualized based on abundance to identify percentile coverages for the total diterpenoid database (Supplemental Data 2g). Carbon number of deconstructed diterpenoids identified database outliers, database entry errors, and non-C_20_ diterpenoids, such as cleavage products and apo-terpenoids (Supplemental Data 2h, 2i).

### Structural comparison of the DNP diterpene skeletons

Due to a higher degree of curation, skeletons identified from the DNP were used for structural comparison (Supplemental Data 2e). A comparison matrix was generated by converting the 671 skeleton SMILEs to bit vectors (binary vectors) using RDKit (v2022.09.1) (3). All skeleton comparisons were made based on Tanimoto coefficients to determine molecular similarity, ranging from 0 to 1 and to create a 671x671 matrix (Supplemental Data 3) (4). This metric measured the similarity between two molecules by comparing the ratio of bond intersections and cyclization compared to the retention of shared atomic structures. This matrix was visualized with seaborn (v0.12.2) as a heatmap (Figure 1; Supplemental Code 2). A principal component analysis (PCA) was conducted on this matrix using sklearn (v1.1.3) functions for transformation, PCA generation, and variance calculations (Supplemental Code 2). This PCA scaled each point based on the number of compounds within the database containing that structure and was colored based on the total number of rings present.

### Modelling diterpene carbocation reactions in Pickaxe

The software, Pickaxe (5), was modified for job executed in a bash terminal (Supplemental Code 3a). The following Pickaxe settings were changed from their default: “input_cpds”, “product_cpds”, “output_dir”, “generations”, “sample_size=1000000”, “coreactant_list”, “rules_list”, “processes=24”, “kekulize=False”, “processes=24”, “quiet=False”, “neutralise=False”. Quality of life additions were made to print an abundance summary of rule usage and to include date and time in output naming. Because Pickaxe has multiple checks for validating compound structures, the inclusion of charged carbocation atoms required adaptation of the rules. The solution used substituted the noble gas xenon, to represent carbocations in all reactions instead. The creation of diTPS SMARTs rules had three guiding principles. First, the majority of carbocation rules prioritized the formation of reactive centers on tertiary carbons, because of the higher commonality and stability of carbocations at these positions (6, 7). Generally, these rules represented a nucleophilic attack of a double bond, ring formation, and subsequent generation of a new tertiary carbocation. Second, when possible, reaction rules focused on the substructure of the molecule involved in the modeled biochemistry as opposed to the entire molecule. For example, typical Class II mechanisms rely on the cascading carbocation attacks from adjacent isoprene unit’s double bonds to form the labdanoid decalin core. One rule models a single isoprene unit attacking a tertiary carbocation 6 carbons away, leading to cyclization. This one rule is used by the model for both the first and second ring formations in labdanoid biosynthesis, despite using different precursors. Focusing on these substructures allowed rules to maintain specificity to known reactions while still allowing the same rule to be robust to other biochemistries as well, thus reducing redundancy within the ruleset. Third, specificity for intermediates were used in cases where a rule could not be generalized to a portion of the molecule, such as in the case of concerted reactions (i.e. cyathane biosynthesis) (8), carbocation stabilization via delocalization or hyperconjugation (i.e. taxadiene biosynthesis) (6), or the generation of secondary carbocations (i.e. cyathane, kaurane, and taxadiene biosynthesis) (Supplemental Data 9) (6, 8, 9). These respective rules were generated to resemble the identified mechanism as closely as possible within the context of Pickaxe. Illustrations of specific reaction rules and their sourcing can be found in Supplemental Data 4.

Pickaxe was executed with four different rulesets where 1.) represented unfiltered class II diTPS action, 2.) represented unfiltered class I diTPS action, 3.) represented the class II and class I rules from steps 1 and 2 but now filtered to known diterpene backbones and 4.) explored where remaining skeleton diversity may be originating. Updated coreactant lists and custom biochemical rulesets were generated for each of the four iterations (Supplemental Data 5a-5d). The first execution applied 10 generations of reactions and used GGDP as the initial substrate. This iteration used the class II diTPS reaction ruleset, specifically excluding diphosphate cleavage and any rules generating macrocyclic compounds (Supplemental Data 5a). The second execution also applied 10 generations of reactions, used GGDP and all compounds produced from execution 1 as inputs. The second iteration used the class I diTPS reaction ruleset (Supplemental Data 5b). All compounds generated from the first and second iteration not containing a diphosphate or carbocation were converted to carbon skeletons and compared to all previously identified TeroKit and DNP skeletons (Supplemental Data 2e, 2f). If a compound matched a previously identified skeleton, the original compound was marked as a successful target match (Supplemental Code 3c). The first and second Pickaxe executions demonstrated the full potential of the unfiltered model, while the filtered, third execution specifically focused on known diterpenes and the validation of the model (Figure 2a, 2b; Supplemental Data 5c, 6).

All filtered reactions from execution 3 were concatenated into a single file and converted into a network containing information on edge (reactions) and node (compound) data (Supplemental Code 3c) along with key identifiers like, SMILES, and compound classifications (GGDP, class II intermediates, class II products, class I intermediates, final targets). These were made into a network in Cytoscape (v3.10.1) to represent known diTPS carbocation reactions with taxadiene and kaurene acting as case studies (Figure 2b-2d; Supplemental Data 6) (10). Within Cytoscape the “analyze network function” was performed to evaluate stress centrality (defined as the importance of a node based on how many shortest paths pass through it) and betweenness centrality (how often a node acts as a bridge along the shortest path between two other nodes) were used to identify central hubs driving diterpene biosynthesis.

While all known cyclic diterpenes require activity from diterpene synthases, with most reported structures only requiring diTPS activity, there remain known cyclizing modifications driven by alcohol dehydrogenases and cytochromes P450 (among others) (11–13) that lead to additional reported skeleton diversity. As a next step, an exploratory ruleset was implemented for 3 generations to identify what modifications could be involved leading to the synthesis of skeletons left unpredicted by the diTPS exclusive model. These rules broke carbon rings, created carbon rings, expanded/collapsed connecting rings, and shifted carbon side chains (Supplemental Data 5d).

### Fingerprinting diterpene backbone diversity from quenching patterns, additional (de)saturation of double bonds, and presence of aromaticity

The 20 most common TeroKit diterpene skeletons (Supplemental Data 2b, 2f) were used to fingerprint each compound class for specific quenching patterns: post cyclization double bonds (de)saturation modifications, and aromaticity (Supplemental Code 4). Deconstructed backbones were used as input for determining (de)saturation and quenching events by counting final hydrogens. Using the diterpene class specific index from the skeleton count (Supplemental Data 2f) and corelated index among deconstructed backbones (Supplemental Data 2b), all hydroxyl groups were temporarily stripped. Reduced backbones then were converted to molecular formula and total hydrogen per molecule were counted. Outlier cases where a molecule had an odd number of hydrogens (an ion) were ignored. If a diterpene is only acted upon by diTPS enzymes it will have a final molecular formula of either C_20_H_32_ (272 m/z), C_20_H_34_O (290 m/z), or C_20_H_36_O_2_ (308 m/z) to reflect whether 0, 1, or 2 carbocations were quenched with water during cyclization. As an additional check that the hydrogen count was caused by the quenching of a carbocation with water, the expected number of quenched carbocations also required hydroxyl groups to be found at tertiary carbons. Otherwise, those structures were considered as oversaturated after diTPS cyclization. Also, because 2 carbocations are generated in class II/class I synthesis but only 1 carbocation is generated in class I only synthesis macrocyclic structures and phytane were also considered oversaturated if a backbone had 36 hydrogens. When there are fewer than 32 hydrogen atoms, it implies that additional desaturation events took place after cyclization. Likewise, when more than 36 hydrogen atoms are present, post cyclization desaturation events are implied. Macrocyclic diterpenes and phytanes are acted upon by one class I diTPS; therefore, in these cases a hydrogen atom count above 34 would also imply post cyclization modification via saturation. If a compound was aromatic, it was labeled as “aromatic” and separated from quenching/saturation counts. Events that are considered exclusive products from diTPS activity (C_20_H_32_, C_20_H_34_O, and C_20_H_36_O_2_) were graphed in pink-scale, with additional (de)saturation and aromaticity in green-scale and grey, respectively (Figure 3).

### Determining diterpene molecular activity from variability of atomic decoration and bonds

Because of the higher compound count and inclusion of stereochemistry, the output skeletons and backbones from the deconstructed TeroKit database (Supplemental Data 2b, 2f) were used to evaluate atomic variability among the top 20 most common diterpene classes (Figure 4; Supplemental Data 7). This analysis compared atom-to-atom and bond-to-bond of all letter and symbol characters from the SMILEs backbones among each individual diterpene skeleton group (Supplemental Code 5). A canonical SMILE for each of the flattened skeletons was generated and served as the reference character string for each diterpene group and to determine where each atom in each model was ordered. To compare within each skeleton class, all backbones had to first be harmonized to have the same number of non-hydrogen atoms. Meeting these configuration requirements was accomplished again by replacing any hydroxyl groups left from deconstruction and their connecting carbons with xenon. If the canonical SMILE for any individual compound completely aligned with the reference SMILEs, the carbon stereocenters, “[C@]”, “[C@@]”, “[C@H]”, “[C@@H]”, were replaced with L, R, D, or U respectively, otherwise stereocenter information was ignored. When aligning strings, all occurrences of “[Xe]” within a given SMILE were replaced with “X”. These conversions of stereocenters and “[Xe]” made it, so all SMILE names were an identical number of atoms and characters L, R, U, and D were later converted back into “C” after the bonds with indicated stereochemistry were replaced with the symbols “^” and “*” to represent the chirality of their referenced bonds. At completion this method generates an output that mirrors a multiple sequence alignment (Supplemental Data 7) that would be produced when aligning, for example protein or DNA sequence. A visual example of the biflorane skeleton reference (underlined) and select entries aligning to that reference are shown below:

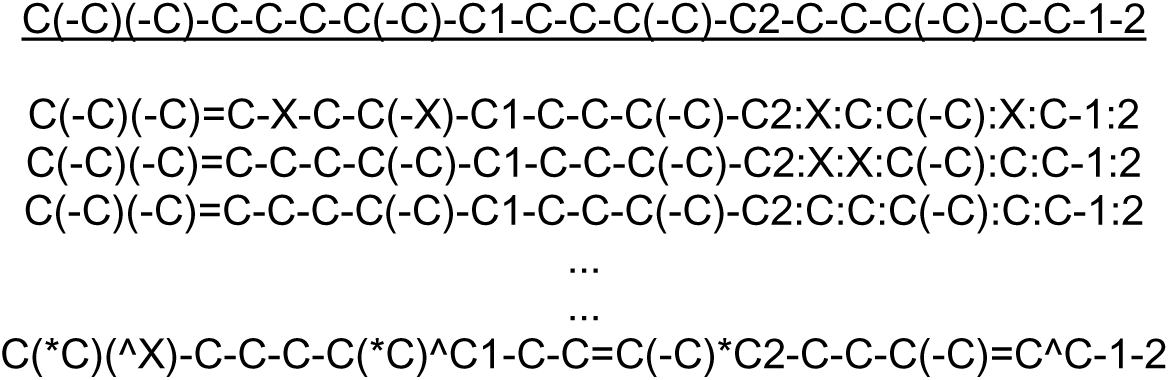

Occasionally a backbone SMILE did not conform to the canonical reference. To correct for this, whichever entry was not cooperating was iteratively changed to match the nonconforming SMILE by replacing bonds with the matching number of double bonds, triple bonds, or aromatic bonds equal to those present in the nonconforming SMILE. When checking for bonds, occurrences of xenon were substituted with carbon. When all unique bond patterns matched, carbon atoms were iteratively replaced with xenon until the updated string aligned to the canonical reference SMILE and was identified as identical to the original SMILE that did not align. In order to make sure all SMILEs had conforming atom order to the canonical SMILE, two approaches were used. The first approach checked compound similarity as each bond or Xenon was replaced, then used the top 5 most similar compounds to the product at each step until number of bonds and Xenon matched the non-conforming SMILE (comparing <60 compounds per iteration). When this method did not work an alternative second approach was implemented. This second approach created all possible combinations of bonds and/or xenon based on the number of occurrences identified from the nonconforming SMILE, formatted in the reference SMILE. This second approach has the potential to generate many compounds for comparison and directly parallels the binomial coefficient equation to estimate compound abundance. For example, if a compound contained ten Xenon atoms, there would be ∼184,000 different possible position combinations that could be generated and only one that will match our nonconforming reference in canonical order. This method, while highly accurate, was only used as a last resort and faces even greater computational demand if tried with molecules larger than diterpenes, like triterpenes (possible upper limit of ∼155M compounds) or carotenoids (possible upper limit of ∼138B compounds). When backbones aligned, the reference SMILE was converted to an edge/node table (Supplementary Code 5). The level of diversity for each bond (edge) and atom (node) was calculated using the index of qualitative variation (IQV). Using IQV as the metric for variability was decided due to the categorical nature of atom decoration (presence/absence) and bonds (stereochemistry and pi bonds presence/absence). This showed how evenly or unevenly any point within a molecule exhibited variability throughout the dataset. All occurrences where xenon was present were assumed to be unique as the decoration that existed prior to deconstruction was converted to a presence/absence value instead. By making this assumption, this analysis weighs the occurrences of decoration more highly than it may have otherwise. IQV values for atoms were used and saved as the intensity for each node and the IQV values for edges (bonds) were used to describe the intensity connecting node-to-node (Supplemental Data 7). Edge/node tables were used as input in Cytoscape (v3.10.1) to visualize each diterpene class (Figure 4; Supplemental Data 7) (10).

Decoration bias based on carbon position (primary, secondary, or tertiary) for the two major classes, labdane-derived and macrocyclic-derived, were investigated. This was accomplished by calculating the percentage of decoration at each carbon location with consideration to the number of connecting carbons and the position of all neighboring carbons (Supplemental Code 5). All conditions were then compared to determine effects based on diterpene class and position-based variation. Only carbon positions that had at least three instances of atomic location and neighbors among labdane- or macrocyclic-derived diterpenes were used.

### Phylogenetic distribution of diterpenes within Viridiplantae, Rhodophyta, and Chromista

The previously generated DNP skeleton summary datasets (Supplemental Data 2a, 2e) were used to 1.) quantify compound occurrences and 2.) identify their sourcing among land plants, green algae, red algae, and brown algae (out groups) (Supplemental Code 6). The reported plant taxonomic family data (Supplemental Data 1a, 2a) was used to extrapolate phylum and kingdom taxonomic classification as well. Comparing and indexing the overlap where particular families and skeletons were both recorded helped to quantify where and how many compounds were reported throughout the Viridiplantae, Rhodophyta, and Chromista clades. Only the top 50 most common diterpene skeletons were investigated. Families with fewer than 10 reported compounds were disregarded in downstream phylogenetic analysis. The remaining families in the heatmap (Figure 5) were manually sorted based on phylum divergence, not hierarchically clustered. Diterpene skeletons, which were hierarchically clustered, were manually annotated based on external (14–18) and DNP reports^6^. A scatterplot illustrating diterpenes per plant family and number of species within each family was also generated (Supplemental Code 6; Supplemental Data 8).

## Supplemental Workflow

Inputs are represented in **Green**. Software is represented in **Purple**. Skeletons and backbones acquired from deconstruction are represented in **Orange**. Examples of the two major diterpene families, the macrocyclic- (**Blue**) and labdane-derived (**Yellow**), which both originate from a geranylgeranyl diphosphate precursor (GGDP; **White**). Terpenoids from the Dictionary of Natural Product (DNP) and TeroKit databases were the input for deconstruction. **1:** Deconstruction isolated diterpene structures to identify backbones and skeletons. **2:** Skeletons were compared to determine structural similarity and distribution of core diterpene classes. **3:** Carbocation biochemistry was modelled for known and hypothetical synthesis. **4:** Structures were compared to determine common diterpene class fingerprints. **5:** Atom-and-Bond variation of major diterpene classes, using each skeleton as a common reference frame. **6:** Distribution of reported diterpene was mapped phylogenetically among plant life.

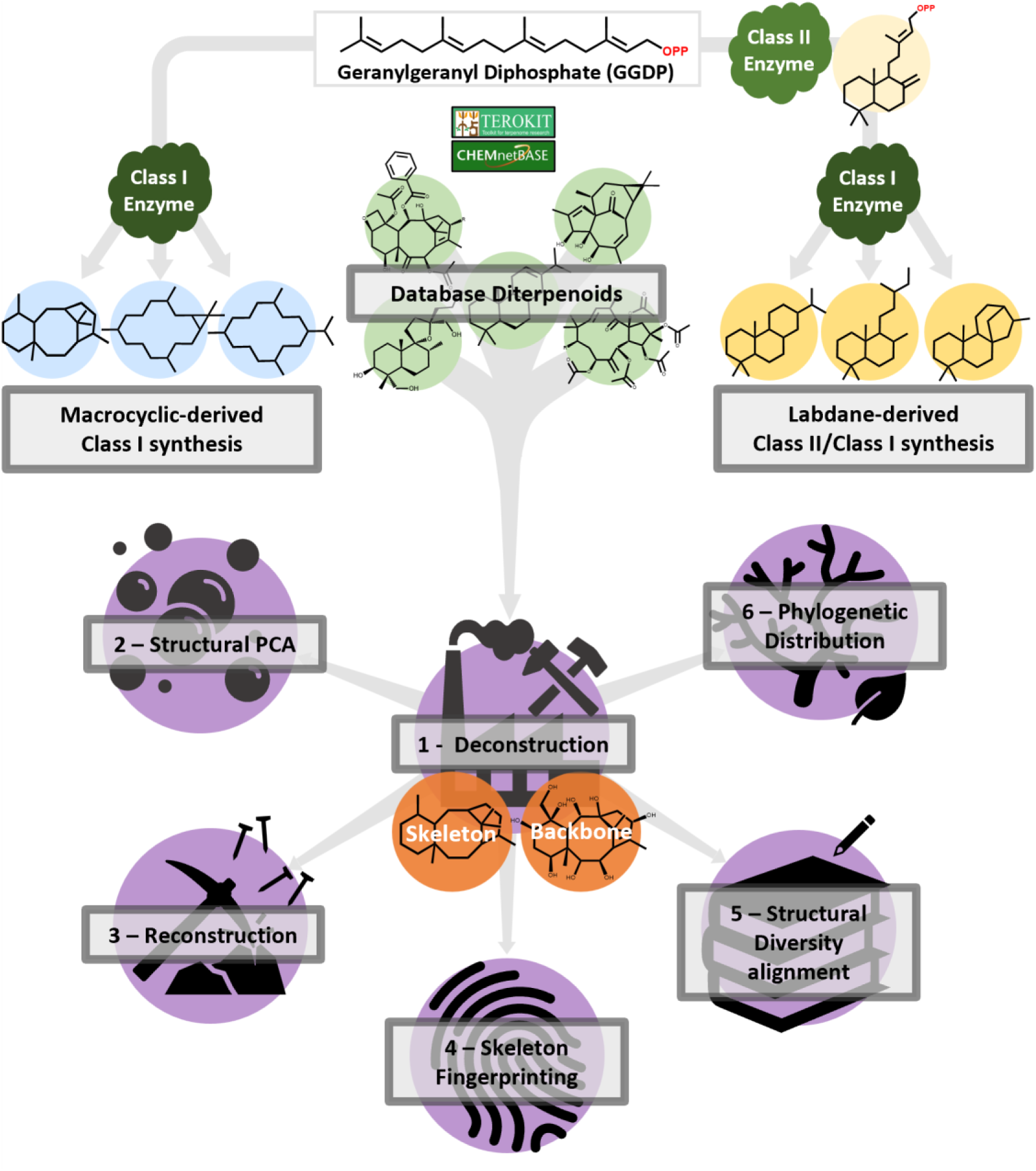

## Footnotes

1 http://dnp.chemnetbase.com

2 http://terokit.qmclab.com/data.html

3 https://dnp.chemnetbase.com/HelpFiles/DNP_Introduction.pdf

4 http://dnp.chemnetbase.com

5 http://terokit.qmclab.com/data.html

6 https://dnp.chemnetbase.com/HelpFiles/DNP_Introduction.pdf

