## Supplementary material for "Rule-Based Deconstruction and Reconstruction of Diterpene Libraries: Categorizing Patterns & Unravelling the Structural Landscape": Zip of all Code used: Supp_Code.1.Terpenoid_Deconstruction.ipynb

```
# %% [code]
# IMPORT ALL RELEVENT MODULES
import pandas as pd
import numpy as np
import os
import re
import matplotlib.pyplot as plt

import rdkit
from rdkit import Chem
from rdkit.Chem import AllChem
from rdkit import DataStructs
from rdkit.Chem import Descriptors
from rdkit.Chem.rdMolDescriptors import CalcMolFormula
from rdkit.Chem.Draw import IPythonConsole
from rdkit.Chem import Draw

Water_coreactant = AllChem.MolFromSmiles('O')

# %% [code]
print("Pandas:",pd.__version__)
print("Numpy:",np.__version__)
print("RE:", re.__version__)
print("Matplotlib:",plt.matplotlib.__version__)
print("RDKit:", rdkit.__version__)
print("OS:",os.__file__)

# %% [markdown]
# Overview of program
## Supplemental data used and generated by this program

# %% [code]
#Supplemental Code 1 Terpenoid Deconstruction Program

### DATASET(S) INTENDED AS INPUT FOR THIS CODE
# Dataset_S1.xlsx (Johnson et al.2019 doi: https://doi.org/10.1074/jbc.RA118.006025)
# Supplemental_Data.1a.DNP_Diterpene_Mining_v30.2.csv
# Supplemental_Data.1b.TeroKit_Diterpene_v2.0.tsv
# Supplemental_Data.1c.TeroKit_Triterpene_v2.0.tsv
# Supplemental_Data.1d.TeroKit_Sesquiterpene_v2.0.tsv)

### SUPPLEMENTAL DATASETS GENERATED BY THIS FUNCTION (GENERATION OF BACKBONE AND SKELETON INFORMATION)
# Supp.2a.DNP_Diterpene_Mining_v30.2_Updated_Skeleton_Backbone.tsv
# Supp.2b.TeroKit_Diterpene_v2.0_Updated_Skeleton_Backbone.tsv
# Supp.2c.TeroKit_Triterpene_v2.0_Updated_Skeleton_Backbone.tsv
# Supp.2d.TeroKit_Sesquiterpene_v2.0_Updated_Skeleton_Backbone.tsv
# Supp.2e.DNP_Diterpene_Skeleton_Summary.tsv
# Supp.2f.Terokit_Diterpene_Skeleton_Summary.tsv
# Supp.2g.Terokit_Diterpene_Skeleton_Abundance_Distribution.png
# Supp.2h.Terokit_Diterpene_Deconstructed_Skeleton_Carbon_Distribution.png
# Supp.2i.10X_Zoomed_Terokit_Diterpene_Deconstructed_Skeleton_Carbon_Distribution.png

'''
### OVERVIEW OF FUNCTIONALITIY ###
### Davis Mathieu
### Jan 1, 2024

This program functions to take a large dataset containing terpenoids
and revert them to their terpene backbones and terpene skeletons. We
define a terpene backbone as the stripped terpenoid structure of all 
functional groups, breaking any oxygen linkages with the aim to 
return just the (cyclized) isoprene derived ring structure (based on
retained carbon-carbon linkages), but retaining double bond variation,
stereochemical variation, and hydroxyl groups present where functional
groups once were. The terpene skeletons take this one step further and 
strip all stereochemistry, bond variation, and hydroxyl groups, leaving
exclusively the "flattened" carbon-carbon linkages, exclusively. This
program is intended to take mined DNP data (found from Johnson et al.
2019 doi: https://doi.org/10.1074/jbc.RA118.006025) Dataset_S1.xlsx with
InChis converted to SMILEs using Obabel and updated to a newer version 
(Supplemental_Data.1a.DNP_Diterpene_Mining_v30.2.csv) as well as the 
table formatted acquired from bulk download on TeroKit 
(http://terokit.qmclab.com/data.html).

Functions:
- Deconstruct2backbone: Takes a terpene in SMILE format and strips
                        functional groups to return the terpene 
                        backbone and the list of all steps taken to
                        deconstruct the molecule.

- backbone_flattener:   Takes the deconstructed backbone and strips
                        all stereochemistry, double bonds, triple 
                        bonds, aromatics, and hydroxyl groups.
                        
These produce updated information on terpene backbones, skeletons,
database deconstruction information (final carbon numbers), and 
skeleton distribution.                    
'''
x=0

# %% [markdown]
## Deconstruction Rules Function

# %% [code]
def deconstruct2backbone(sub):
    The_Process = [sub]
    #sub = Chem.MolFromSmiles(Chem.MolToSmiles(sub), sanitize=False)
    sub = Chem.MolFromSmiles(Chem.MolToSmiles(sub))
    

    """
    Remove Fatty Acids
    """        
    fatty_acid_present = sub.GetSubstructMatches(Chem.MolFromSmarts("[#6:21]!@[#8:0]!@[#6:1]!@[#6:2]!@[#6:3]!@[#6:4]!@[#6:5]!@[#6:6]!@[#6:7]!@[#6:8]!@[#6:9]!@[#6:10]!@[#6:11]!@[#6:12]!@[#6:13]!@[#6:14]!@[#6:15]!@[#6:16]!@[#6:17]!@[#6:18]"))
    while(fatty_acid_present):
        remove_fatty_acid = AllChem.ReactionFromSmarts("[#6:21]!@[#8:0]!@[#6:1]!@[#6:2]!@[#6:3]!@[#6:4]!@[#6:5]!@[#6:6]!@[#6:7]!@[#6:8]!@[#6:9]!@[#6:10]!@[#6:11]!@[#6:12]!@[#6:13]!@[#6:14]!@[#6:15]!@[#6:16]!@[#6:17]!@[#6:18]>>[#6:21][#8:0].[#6:1][#6:2][#6:3][#6:4][#6:5][#6:6][#6:7][#6:8][#6:9][#6:10][#6:11][#6:12][#6:13][#6:14][#6:15][#6:16][#6:17][#6:18]")
        sub1=remove_fatty_acid.RunReactants((sub,))[0][0]
        if(re.search("CCCCCCCCCC", Chem.MolToSmiles(remove_fatty_acid.RunReactants((sub,))[0][1]))):
            sub=Chem.MolFromSmiles(Chem.MolToSmiles(sub1))
            fatty_acid_present = sub.GetSubstructMatches(Chem.MolFromSmarts("[#6:21]!@[#8:0]!@[#6:1]!@[#6:2]!@[#6:3]!@[#6:4]!@[#6:5]!@[#6:6]!@[#6:7]!@[#6:8]!@[#6:9]!@[#6:10]!@[#6:11]!@[#6:12]!@[#6:13]!@[#6:14]!@[#6:15]!@[#6:16]!@[#6:17]!@[#6:18]"))
        else:
            fatty_acid_present = []
        The_Process.append(sub)

    glucose_present = sub.GetSubstructMatches(Chem.MolFromSmarts("[#6:1]-1(-[#8:8][#6:15])[#8:3][#6:4](-[#6:9][#8:10])[#6:5](-[#8:11])[#6:6](-[#8:12])[#6:7](-[#8:13])-1"))
    while(len(glucose_present) > 0):
        condition =""
        rxn4 = AllChem.ReactionFromSmarts("[#6:1]-1(-[#8:8][#6:15])[#8:3][#6:4](-[#6:9][#8:10])[#6:5](-[#8:11])[#6:6](-[#8:12])[#6:7](-[#8:13])-1.[#8:14]>>[#6:15][#8:14].[#6:1]-1(-[#8:8])[#8:3][#6:4](-[#6:9][#8:10])[#6:5](-[#8:11])[#6:6](-[#8:12])[#6:7](-[#8:13])-1")
        sub1 = rxn4.RunReactants((sub,Water_coreactant))
        for i in sub1:
            #print(CalcMolFormula(Chem.MolFromSmiles(Chem.MolToSmiles(i[1]))))
            if(Chem.MolFromSmiles(Chem.MolToSmiles(i[1]))==None):
                condition=""
            elif(CalcMolFormula(Chem.MolFromSmiles(Chem.MolToSmiles(i[1])))=="C6H12O6"):
                condition="skip"
                sub= i[0]; break
            elif(CalcMolFormula(Chem.MolFromSmiles(Chem.MolToSmiles(i[1])))=="C18H32O15"):
                condition="skip"
                sub= i[0]; break
            elif(CalcMolFormula(Chem.MolFromSmiles(Chem.MolToSmiles(i[1])))=="C18H32O14"):
                condition="skip"
                sub= i[0]; break
            elif(int(re.findall("\d+",CalcMolFormula(Chem.MolFromSmiles(Chem.MolToSmiles(i[0]))))[0])%5==0):
                condition="skip"
                sub=i[0];break
            elif(int(re.findall("\d+",CalcMolFormula(Chem.MolFromSmiles(Chem.MolToSmiles(i[0]))))[0])==19):
                condition="skip"
                sub=i[0];break
            elif(len(re.findall("S",CalcMolFormula(Chem.MolFromSmiles(Chem.MolToSmiles(i[1])))))>0):
                condition="skip"
                sub=i[0];break
        if(condition=="skip"):
            condition==""
        elif(len(re.findall("C",Chem.MolToSmiles(sub1[0][0]),re.IGNORECASE))<len(re.findall("C",Chem.MolToSmiles(sub1[0][1]),re.IGNORECASE))):
            if(Chem.MolFromSmiles(Chem.MolToSmiles(sub1[0][1]))!=None and CalcMolFormula(Chem.MolFromSmiles(Chem.MolToSmiles(sub1[0][1])))!="C6H12O6" and len(re.findall("C",Chem.MolToSmiles(sub1[0][1]),re.IGNORECASE))>len(re.findall("C",Chem.MolToSmiles(sub1[0][0]),re.IGNORECASE))):
                sub=sub1[0][1]
            elif(Chem.MolFromSmiles(Chem.MolToSmiles(sub1[0][0]))!=None and CalcMolFormula(Chem.MolFromSmiles(Chem.MolToSmiles(sub1[0][0])))!="C6H12O6"):
                sub=sub1[0][0]
            else:
                break
        else:
            sub = sub1[0][0]
        sub = Chem.MolFromSmiles(Chem.MolToSmiles(sub))
        glucose_present = sub.GetSubstructMatches(Chem.MolFromSmarts("[#6:1]-1(-[#8:8][#6:15])[#8:3][#6:4](-[#6:9][#8:10])[#6:5](-[#8:11])[#6:6](-[#8:12])[#6:7](-[#8:13])-1"))
        The_Process.append(sub) 

    sub = Chem.MolFromSmiles(Chem.MolToSmiles(sub))
    C13_present = sub.GetSubstructMatches(Chem.MolFromSmarts("[#6:1]!@[13#6H]"))
    if(len(C13_present)>0):
        sub =str(Chem.MolToSmiles(sub))
        sub = re.sub("\[13CH3\]","C",sub)
        sub = re.sub("\[13CH2\]","C",sub)
        sub = re.sub("\[13CH\]","C",sub)
        sub = re.sub("\[13C\]","C",sub)
        sub = Chem.MolFromSmiles(sub)

    sub = Chem.MolFromSmiles(Chem.MolToSmiles(sub))
    C14_present = sub.GetSubstructMatches(Chem.MolFromSmarts("[#6:1]!@[14#6H]"))
    if(len(C14_present)>0):
        sub =str(Chem.MolToSmiles(sub))
        sub = re.sub("\[14CH3\]","C",sub)
        sub = re.sub("\[14CH2\]","C",sub)
        sub = re.sub("\[14CH\]","C",sub)
        sub = re.sub("\[14C\]","C",sub)
        sub = Chem.MolFromSmiles(sub)
        
    carbocation_present = sub.GetSubstructMatches(Chem.MolFromSmarts("[#6+:1]"))
    if(len(carbocation_present)>0):
        sub= str(Chem.MolToSmiles(sub))
        sub = re.sub("\[CH2\+\]", "C",sub)
        sub = re.sub("\[CH\+\]", "C",sub)
        sub = re.sub("\[C\+\]", "C",sub)
        sub = Chem.MolFromSmiles(sub)

    carbanion_present = sub.GetSubstructMatches(Chem.MolFromSmarts("[#6-:1]"))
    if(len(carbanion_present)>0):
        sub = str(Chem.MolToSmiles(sub))
        sub = re.sub("\[CH2\-\]", "C",sub) 
        sub = re.sub("\[CH\-\]", "C",sub)
        sub = re.sub("\[C\-\]", "C",sub)
        sub = Chem.MolFromSmiles(sub)

 
    glucose_present = sub.GetSubstructMatches(Chem.MolFromSmarts("[#6:1]-1[#8:2][#6:3](-[#6:4]-[#6:5](=[#8:6])-[#8:7])[#6:8](-[#8:9])[#6:10](-[#8:11])[#6:12](-[#8:13])-1"))
    while(len(glucose_present)>0):
        remove_glucose = AllChem.ReactionFromSmarts("[#6:1]-1[#8:2][#6:3](-[#6:4]-[#6:5](=[#8:6])-[#8:7])[#6:8](-[#8:9])[#6:10](-[#8:11])[#6:12](-[#8:13])-1>>[#8:7].[#6:1]-1[#8:2][#6:3](-[#6:4]-[#6:5](=[#8:6]))[#6:8](-[#8:9])[#6:10](-[#8:11])[#6:12](-[#8:13])-1")
        sub = remove_glucose.RunReactants((sub,))[0][0]
        glucose_present = sub.GetSubstructMatches(Chem.MolFromSmarts("[#6:1]-1[#8:2][#6:3](-[#6:4]-[#6:5](=[#8:6])-[#8:7])[#6:8](-[#8:9])[#6:10](-[#8:11])[#6:12](-[#8:13])-1"))
        The_Process.append(sub)
           
    pentopyranose_sugar_present = sub.GetSubstructMatches(Chem.MolFromSmarts("[#6:1](-[#8:2][#6:11])(-[#8:3]-1)[#6:4](-[#8:5])[#6:6](-[#8:7])[#6:8](-[#8:9])[#6:10]-1"))
    while(len(pentopyranose_sugar_present) > 0):
        rxn7 = AllChem.ReactionFromSmarts("[#6:1](-[#8:2][#6:11])(-[#8:3]-1)[#6:4](-[#8:5])[#6:6](-[#8:7])[#6:8](-[#8:9])[#6:10]-1>>[#8:2][#6:11].[#6:1](-[#8:3]-1)[#6:4](-[#8:5])[#6:6](-[#8:7])[#6:8](-[#8:9])[#6:10]-1")
        sub1 = rxn7.RunReactants((sub,))
        if(len(re.findall("C",Chem.MolToSmiles(sub1[0][0]), re.IGNORECASE))<len(re.findall("C",Chem.MolToSmiles(sub1[0][1]), re.IGNORECASE))):
            sub = sub1[0][1]
        else:
            sub = sub1[0][0]
        pentopyranose_sugar_present = sub.GetSubstructMatches(Chem.MolFromSmarts("[#6:1](-[#8:2][#6:11])(-[#8:3]-1)[#6:4](-[#8:5])[#6:6](-[#8:7])[#6:8](-[#8:9])[#6:10]-1"))
        #print("pentopyranose present")
        The_Process.append(sub)

    furan_present = sub.GetSubstructMatches(Chem.MolFromSmarts("[#6:1][#6:2]-1=[#6:3](-[#8:4])[#6:5](=[#8:6])-[#8:7][#6:8](-[#8:9])-1"))
    while(len(furan_present)>0):
        rxn_8 = AllChem.ReactionFromSmarts("[#6:1][#6:2]-1=[#6:3](-[#8:4])[#6:5](=[#8:6])-[#8:7][#6:8](-[#8:9])-1>>[#6:1].[#6:2]-1=[#6:3](-[#8:4])[#6:5](=[#8:6])-[#8:7][#6:8](-[#8:9])-1")
        sub = rxn_8.RunReactants((sub,))[0][0]
        furan_present = sub.GetSubstructMatches(Chem.MolFromSmarts("[#6:1][#6:2]-1=[#6:3](-[#8:4])[#6:5](=[#8:6])-[#8:7][#6:8](-[#8:9])-1"))
        The_Process.append(sub)
      
    furanose_sugar_present = sub.GetSubstructMatches(Chem.MolFromSmarts("[#8:1]-1-[#6:2](-[#8:3][#6:11])[#6:4](-[#8:5])[#6:6](-[#8:7])-[#6:8]-1(-[#6:9][#8:10])"))
    while(len(furanose_sugar_present) > 0):
        rxn8 = AllChem.ReactionFromSmarts("[#8:1]-1-[#6:2](-[#8:3][#6:11])[#6:4](-[#8:5])[#6:6](-[#8:7])-[#6:8]-1(-[#6:9][#8:10]).[#8:12]>>[#6:11][#8:12].[#8:1]-1-[#6:2](-[#8:3])[#6:4](-[#8:5])[#6:6](-[#8:7])-[#6:8]-1(-[#6:9][#8:10])")
        sub = rxn8.RunReactants((sub,Water_coreactant))[0][0]
        furanose_sugar_present = sub.GetSubstructMatches(Chem.MolFromSmarts("[#8:1]-1-[#6:2](-[#8:3][#6:11])[#6:4](-[#8:5])[#6:6](-[#8:7])-[#6:8]-1(-[#6:9][#8:10])"))
        #print("furanose present")
        The_Process.append(sub)
   

    coumarin_present = sub.GetSubstructMatches(Chem.MolFromSmarts("[#6:1]:1(-[#8:7]):[#6:2]:[#6:3]:[#6:4]:[#6:5]:2:[#6:6]:1:[#8:8]:[#6:9](=[#8:10]):[#6:11]:[#6:12]:2"))
    while(len(coumarin_present) > 0):
        rxn6 = AllChem.ReactionFromSmarts("[#6:1]:1(-[#8:7]):[#6:2]:[#6:3]:[#6:4]:[#6:5]:2:[#6:6]:1:[#8:8]:[#6:9](=[#8:10]):[#6:11]:[#6:12]:2>>[#6:1]:1:[#6:2]:[#6:3]:[#6:4]:[#6:5]:2:[#6:6]:1:[#8:8]:[#6:9](=[#8:10]):[#6:11]:[#6:12]:2.[#8:7]")
        sub = rxn6.RunReactants((sub,))[0][1]
        coumarin_present = sub.GetSubstructMatches(Chem.MolFromSmarts("[#6:1]:1(-[#8:7]):[#6:2]:[#6:3]:[#6:4]:[#6:5]:2:[#6:6]:1:[#8:8]:[#6:9](=[#8:10]):[#6:11]:[#6:12]:2"))    
        #print("Coumarin_present")
        The_Process.append(sub)

    phosphate_present = sub.GetSubstructMatches(Chem.MolFromSmarts("[#6:1]-[#8:2]-[#15:3](-[#8:4])(-[#8:5])=[#8:6]"))
    while(len(phosphate_present)>0):
        remove_phosphate = AllChem.ReactionFromSmarts("[#6:1]-[#8:2]-[#15:3](-[#8:4])(-[#8:5])=[#8:6]>>[#6:1]-[#8:2].[#15:3](-[#8:4])(-[#8:5])=[#8:6]")
        sub1 = remove_phosphate.RunReactants((sub,))
        if(len(re.findall("C",Chem.MolToSmiles(sub1[0][0]),re.IGNORECASE))<len(re.findall("C",Chem.MolToSmiles(sub1[0][1]),re.IGNORECASE))):
            sub=sub1[0][1]
        else:
            sub=sub1[0][0]
        phosphate_present = sub.GetSubstructMatches(Chem.MolFromSmarts("[#6:1]-[#8:2]-[#15:3](-[#8:4])(-[#8:5])=[#8:6]"))
        The_Process.append(sub)

    phosphate_present = sub.GetSubstructMatches(Chem.MolFromSmarts("[#6:1]-[#15:3](-[#8:4])(-[#8:5])=[#8:6]"))
    while(len(phosphate_present)>0):
        remove_phosphate = AllChem.ReactionFromSmarts("[#6:1]-[#15:3](-[#8:4])(-[#8:5])=[#8:6].[#8:2]>>[#6:1]-[#8:2].[#15:3](-[#8:4])(-[#8:5])=[#8:6]")
        sub1 = remove_phosphate.RunReactants((sub,Water_coreactant))#[0][0]
        if(Chem.MolToSmiles(sub1[0][0])=="CO"):
            sub=sub1[0][1]
        else:
            sub=sub1[0][0]
        phosphate_present = sub.GetSubstructMatches(Chem.MolFromSmarts("[#6:1]-[#15:3](-[#8:4])(-[#8:5])=[#8:6]"))
        The_Process.append(sub)        

    diphosphate_present = sub.GetSubstructMatches(Chem.MolFromSmarts("[#8:1]-[#15:2]-[#8:3]-[#15:4]"))
    while(len(diphosphate_present)>0):
        remove_DP = AllChem.ReactionFromSmarts("[#8:1]-[#15:2]-[#8:3]-[#15:4]>>[#8:1].[#15:2]-[#8:3]-[#15:4]")
        sub = remove_DP.RunReactants((sub,))[0][0]
        diphosphate_present = sub.GetSubstructMatches(Chem.MolFromSmarts("[#8:1]-[#15:2]-[#8:3]-[#15:4]"))
        The_Process.append(sub)

    phosphate_present = sub.GetSubstructMatches(Chem.MolFromSmarts("[#6:0]-[#8:1]-[#15:2]"))    
    while(len(phosphate_present)>0):
        remove_phosphate = AllChem.ReactionFromSmarts("[#6:0]-[#8:1]-[#15:2]>>[#6:0]-[#8:1].[#15:2]")
        sub = remove_phosphate.RunReactants((sub,))[0][0]
        phosphate_present = sub.GetSubstructMatches(Chem.MolFromSmarts("[#6:0]-[#8:1]-[#15:2]"))
        The_Process.append(sub) 

    silicon_present = sub.GetSubstructMatches(Chem.MolFromSmarts("[#6:0][#14:1]"))
    while(len(silicon_present)>0):
        remove_silicon=AllChem.ReactionFromSmarts("[#6:0][#14:1].[#8:2]>>[#6:0][#8:2].[#14:1]")
        sub1 = remove_silicon.RunReactants((sub,Water_coreactant))
        if(len(re.findall("C",Chem.MolToSmiles(sub1[0][0]),re.IGNORECASE))>len(re.findall("C",Chem.MolToSmiles(sub1[0][1]),re.IGNORECASE))):
            sub = sub1[0][0]
        else:
            sub = sub1[0][1]
        silicon_present = sub.GetSubstructMatches(Chem.MolFromSmarts("[#6:0][#14:1]"))
        The_Process.append(sub)
    
    selenium_present = sub.GetSubstructMatches(Chem.MolFromSmarts("[#6:0]-[#34:1]"))
    while(len(selenium_present)>0):
        remove_selenium=AllChem.ReactionFromSmarts("[#6:0]-[#34:1].[#8:2]>>[#6:0][#8:2].[#34:1]")
        sub1 = remove_selenium.RunReactants((sub,Water_coreactant))
        if(len(re.findall("C",Chem.MolToSmiles(sub1[0][0]),re.IGNORECASE))>len(re.findall("C",Chem.MolToSmiles(sub1[0][1]),re.IGNORECASE))):
            sub = sub1[0][0]
        else:
            sub = sub1[0][1]
        selenium_present = sub.GetSubstructMatches(Chem.MolFromSmarts("[#6:0]-[#34:1]"))
        The_Process.append(sub)
    
    
    tin_present = sub.GetSubstructMatches(Chem.MolFromSmarts("[#6:1][#8:2][#50:3]"))
    while(len(tin_present)>0):
        remove_tin=AllChem.ReactionFromSmarts("[#6:1][#8:2][#50:3]>>[#6:1][#8:2].[#50:3]")
        sub = remove_tin.RunReactants((sub,))[0][0]
        tin_present = sub.GetSubstructMatches(Chem.MolFromSmarts("[#6:1][#8:2][#50:3]"))
        The_Process.append(sub)
    
    boron_present = sub.GetSubstructMatches(Chem.MolFromSmarts("[#6:1]-[#5:2]"))
    while(len(boron_present)>0):
        remove_boron=AllChem.ReactionFromSmarts("[#6:1]-[#5:2].[#8:3]>>[#6:1]-[#8:3].[#5:2]")
        sub = remove_boron.RunReactants((sub,Water_coreactant))[0][0]
        boron_present = sub.GetSubstructMatches(Chem.MolFromSmarts("[#6:1]-[#5:2]"))
        The_Process.append(sub)
    
    silicon_present = sub.GetSubstructMatches(Chem.MolFromSmarts("[#6:0][#8:1][#14:2]"))
    while(len(silicon_present)>0):
        remove_silicon=AllChem.ReactionFromSmarts("[#6:0][#8:1][#14:2]>>[#6:0][#8:1].[#14:2]")        
        sub = remove_silicon.RunReactants((sub,))[0][0]
        silicon_present = sub.GetSubstructMatches(Chem.MolFromSmarts("[#6:0][#8:1][#14:2]"))
        The_Process.append(sub)
        
    sulfur_compound_present = sub.GetSubstructMatches(Chem.MolFromSmarts("[#16:1](=[#8:2])(=[#8:3])(-[#8:4])-[#6:5]"))
    while(len(sulfur_compound_present)>0):
        remove_sulfur=AllChem.ReactionFromSmarts("[#16:1](=[#8:2])(=[#8:3])(-[#8:4])-[#6:5]>>[#16:1](=[#8:2])(=[#8:3]).[#8:4][#6:5]")
        sub = remove_sulfur.RunReactants((sub, ))[0][1]
        sulfur_compound_present = sub.GetSubstructMatches(Chem.MolFromSmarts("[#16:1](=[#8:2])(=[#8:3])(-[#8:4])-[#6:5]"))
        #print("Sulfur present")
        The_Process.append(sub)        
      
    sulfur_compound_present = sub.GetSubstructMatches(Chem.MolFromSmarts("[#16:1]-[#6:2]"))
    while(len(sulfur_compound_present) > 0):
        remove_sulfur=AllChem.ReactionFromSmarts("[#6:1]-[#16:2].[#8:4]>>[#6:1][#8:4].[#16:2]")
        sub1 =remove_sulfur.RunReactants((sub,Water_coreactant))
        if(len(re.findall("C",Chem.MolToSmiles(sub1[0][0])))<len(re.findall("C",Chem.MolToSmiles(sub1[0][1])))):
            sub = sub1[0][1]
        else:
            sub = sub1[0][0]
        sulfur_compound_present = sub.GetSubstructMatches(Chem.MolFromSmarts("[#16:1]-[#6:2]"))
        #print("Sulfur present")
        The_Process.append(sub)
        
    sulfur_compound_present = sub.GetSubstructMatches(Chem.MolFromSmarts("[#16:1][#8:2][#6:3]"))
    while(len(sulfur_compound_present)>0):
        remove_sulfur=AllChem.ReactionFromSmarts("[#16:1][#8:2][#6:3]>>[#16:1].[#8:2][#6:3]")
        sub = remove_sulfur.RunReactants((sub, ))[0][1]
        sulfur_compound_present = sub.GetSubstructMatches(Chem.MolFromSmarts("[#16:1][#8:2][#6:3]"))
        #print("Sulfur present")
        The_Process.append(sub)


### RESOLVING NITROGEN BASED SIDE CHAINS ################################
    pyrimadine_present = sub.GetSubstructMatches(Chem.MolFromSmarts("[#7:1]2[#6:2][#7:3][#6:4]1[#7:5][#6:6][#7:7](-[#6:8])[#6:9]1[#6:10]2"))
    while(len(pyrimadine_present) > 0):
        rxn5 = AllChem.ReactionFromSmarts("[#7:1]2[#6:2][#7:3][#6:4]1[#7:5][#6:6][#7:7](-[#6:8])[#6:9]1[#6:10]2.[#8:11]>>[#6:8][#8:11].[#7:1]2[#6:2][#7:3][#6:4]1[#7:5][#6:6][#7:7][#6:9]1[#6:10]2")
        sub1 = rxn5.RunReactants((sub,Water_coreactant))
        if(Chem.MolToSmiles(sub1[0][0])=="CO"):
            sub = sub1[0][1]
        else:
            sub = sub1[0][0]
        pyrimadine_present = sub.GetSubstructMatches(Chem.MolFromSmarts("[#7:1]2[#6:2][#7:3][#6:4]1[#7:5][#6:6][#7:7](-[#6:8])[#6:9]1[#6:10]2"))
        The_Process.append(sub)
        #print("pyrimadine present")
 
    pyrimadine_present = sub.GetSubstructMatches(Chem.MolFromSmarts("[#7:1]2[#6:2][#7:3][#6:4]1[#7:5](-[#6:8])[#6:6][#7:7][#6:9]1[#6:10]2"))
    while(len(pyrimadine_present) > 0):
        rxn5 = AllChem.ReactionFromSmarts("[#7:1]2[#6:2][#7:3][#6:4]1[#7:5](-[#6:8])[#6:6][#7:7][#6:9]1[#6:10]2.[#8:11]>>[#6:8][#8:11].[#7:1]2[#6:2][#7:3][#6:4]1[#7:5][#6:6][#7:7][#6:9]1[#6:10]2")
        sub1 = rxn5.RunReactants((sub,Water_coreactant))
        if(Chem.MolToSmiles(sub1[0][0])=="CO"):
            sub = sub1[0][1]
        elif(len(re.findall("C",Chem.MolToSmiles(sub1[0][1]), re.IGNORECASE))>len(re.findall("C",Chem.MolToSmiles(sub1[0][0]), re.IGNORECASE))):
            sub = sub1[0][1]
        else:
            sub = sub1[0][0]
        pyrimadine_present = sub.GetSubstructMatches(Chem.MolFromSmarts("[#7:1]2[#6:2][#7:3][#6:4]1[#7:5](-[#6:8])[#6:6][#7:7][#6:9]1[#6:10]2"))
        The_Process.append(sub)
        #print("pyrimadine present")        
        
    #remove pyrole
    pyrole_present = sub.GetSubstructMatches(Chem.MolFromSmarts("[#8:1]-[#6:2]=[#6:3]-1-[#6:4]=[#6:5]-[#6:6]=[#7:7]-1"))
    while(len(pyrole_present)>0):
        remove_pyrole = AllChem.ReactionFromSmarts("[#8:1]-[#6:2]=[#6:3]-1-[#6:4]=[#6:5]-[#6:6]=[#7:7]-1>>[#8:1].[#6:2]=[#6:3]-1-[#6:4]=[#6:5]-[#6:6]=[#7:7]-1")
        sub = remove_pyrole.RunReactants((sub,))[0][0]
        pyrole_present = sub.GetSubstructMatches(Chem.MolFromSmarts("[#8:1]-[#6:2]=[#6:3]-1-[#6:4]=[#6:5]-[#6:6]=[#7:7]-1"))
        The_Process.append(sub)
        #print("Pyrole present")

    sub = Chem.MolFromSmiles(Chem.MolToSmiles(sub))
    indole_present = sub.GetSubstructMatch(Chem.MolFromSmarts("[#6:0]-[#6:1]:1:[#6:2]:[#7:3]:[#6:4]:2:[#6:5]:1:[#6:6]:[#6:7]:[#6:8]:[#6:9]:2"))
    while(len(indole_present)>0):
        remove_indole = AllChem.ReactionFromSmarts("[#6:0]-[#6:1]:1:[#6:2]:[#7:3]:[#6:4]:2:[#6:5]:1:[#6:6]:[#6:7]:[#6:8]:[#6:9]:2>>[#6:0].[#6:1]:1:[#6:2]:[#7:3]:[#6:4]:2:[#6:5]:1:[#6:6]:[#6:7]:[#6:8]:[#6:9]:2")
        sub = remove_indole.RunReactants((sub,))[0][0]
        sub = Chem.MolFromSmiles(Chem.MolToSmiles(sub))
        indole_present = sub.GetSubstructMatch(Chem.MolFromSmarts("[#6:0]-[#6:1]:1:[#6:2]:[#7:3]:[#6:4]:2:[#6:5]:1:[#6:6]:[#6:7]:[#6:8]:[#6:9]:2"))
        The_Process.append(sub)
        

    pyrimidine_present = sub.GetSubstructMatches(Chem.MolFromSmarts("[#7:0]-[#6:1]:1:[#6:2]:[#7:3]:[#6:4]:[#7:5]:[#6:6]:1"))
    while(len(pyrimidine_present)>0):
        remove_pyrimidine = AllChem.ReactionFromSmarts("[#7:0]-[#6:1]:1:[#6:2]:[#7:3]:[#6:4]:[#7:5]:[#6:6]:1>>[#7:0].[#6:1]:1:[#6:2]:[#7:3]:[#6:4]:[#7:5]:[#6:6]:1")
        sub = remove_pyrimidine.RunReactants((sub,))[0][0]        
        pyrimidine_present = sub.GetSubstructMatches(Chem.MolFromSmarts("[#7:0]-[#6:1]:1:[#6:2]:[#7:3]:[#6:4]:[#7:5]:[#6:6]:1"))
        The_Process.append(sub)
        #print("Pyrimidine present")
 
    pyrolidine_present = sub.GetSubstructMatches(Chem.MolFromSmarts("[#6:1]-1-[#6:2]-[#6:3]=[#7:4]-[#6:5]-1"))
    while(len(pyrolidine_present)>0):
        resolve_pyrolidine = AllChem.ReactionFromSmarts("[#6:1]-1-[#6:2]-[#6:3]=[#7:4]-[#6:5]-1>>[#6:1](-[#6:5])-[#6:2]-[#6:3]=[#7:4]")
        sub = resolve_pyrolidine.RunReactants((sub,))[0][0]
        pyrolidine_present = sub.GetSubstructMatches(Chem.MolFromSmarts("[#6:1]-1-[#6:2]-[#6:3]=[#7:4]-[#6:5]-1"))
        The_Process.append(sub)
        #print("pyrolidine present")
         
    pyrolidine_present = sub.GetSubstructMatches(Chem.MolFromSmarts("[#6:1]-1=[#6:2]-[#6:3][#7:4]=[#6:5]-1"))
    while(len(pyrolidine_present)>0):
        resolve_pyrolidine = AllChem.ReactionFromSmarts("[#6:1]-1=[#6:2]-[#6:3][#7:4]=[#6:5]-1>>[#6:1](-[#6:5]=[#7:4])-[#6:2]-[#6:3]")
        sub = resolve_pyrolidine.RunReactants((sub,))[0][0]
        pyrolidine_present = sub.GetSubstructMatches(Chem.MolFromSmarts("[#6:1]-1=[#6:2]-[#6:3][#7:4]=[#6:5]-1"))
        The_Process.append(sub)
        #print("pyrolidine present")

    nitrogenous_base = sub.GetSubstructMatches(Chem.MolFromSmarts("[#7:1]:1:[#6:2]:[#7:3](-[#6:10]):[#6:4]-2:[#6:5]:1-[#7:6][#6:7]=[#7:8]-[#6:9]-2"))
    while(len(nitrogenous_base)>0):
        remove_nitrogenous_base = AllChem.ReactionFromSmarts("[#7:1]:1:[#6:2]:[#7:3](-[#6:10]):[#6:4]-2:[#6:5]:1-[#7:6][#6:7]=[#7:8]-[#6:9]-2>>[#6:10].[#7:1]:1:[#6:2]:[#7:3]:[#6:4]-2:[#6:5]:1-[#7:6][#6:7]=[#7:8]-[#6:9]-2")
        sub = remove_nitrogenous_base.RunReactants((sub,))[0][0]
        nitrogenous_base = sub.GetSubstructMatches(Chem.MolFromSmarts("[#7:1]:1:[#6:2]:[#7:3](-[#6:10]):[#6:4]-2:[#6:5]:1-[#7:6][#6:7]=[#7:8]-[#6:9]-2"))
        The_Process.append(sub)

    nitrogenous_base = sub.GetSubstructMatches(Chem.MolFromSmarts("[#7:1]:1:[#7:2]:[#7:3]:[#6:4](-[#6:6]):[#6:5]:1"))
    while(len(nitrogenous_base)>0):
        remove_nitrogenous_base = AllChem.ReactionFromSmarts("[#7:1]:1:[#7:2]:[#7:3]:[#6:4](-[#6:6]):[#6:5]:1>>[#6:6].[#7:1]:1:[#7:2]:[#7:3]:[#6:4]:[#6:5]:1")
        sub = remove_nitrogenous_base.RunReactants((sub,))[0][0]
        nitrogenous_base = sub.GetSubstructMatches(Chem.MolFromSmarts("[#7:1]:1:[#7:2]:[#7:3]:[#6:4](-[#6:6]):[#6:5]:1"))
        The_Process.append(sub)


    nitrogenous_base = sub.GetSubstructMatches(Chem.MolFromSmarts("[#7:1]:1:[#6:2](=[#8:7]):[#7:3]:[#6:4](=[#8:8]):[#6:5](-[#6:9]):[#6:6]:1"))
    while(len(nitrogenous_base)>0):
        remove_nitrogenous_base = AllChem.ReactionFromSmarts("[#7:1]:1:[#6:2](=[#8:7]):[#7:3]:[#6:4](=[#8:8]):[#6:5](-[#6:9]):[#6:6]:1>>[#6:9].[#7:1]:1:[#6:2](=[#8:7]):[#7:3]:[#6:4](=[#8:8]):[#6:5]:[#6:6]:1")
        sub = remove_nitrogenous_base.RunReactants((sub,))[0][0]
        nitrogenous_base = sub.GetSubstructMatches(Chem.MolFromSmarts("[#7:1]:1:[#6:2](=[#8:7]):[#7:3]:[#6:4](=[#8:8]):[#6:5](-[#6:9]):[#6:6]:1"))
        The_Process.append(sub)
    
    nitrogenous_base = sub.GetSubstructMatches(Chem.MolFromSmarts("[#6:1]:1:[#6:2]([#7:7]):[#6:3](-[#6:8](=[#8:9])-[#8:10][#6:11]):[#6:4]:[#6:5]:[#6:6]:1"))
    while(len(nitrogenous_base)>0):
        remove_nitrogenous_base = AllChem.ReactionFromSmarts("[#6:1]:1:[#6:2]([#7:7]):[#6:3](-[#6:8](=[#8:9])-[#8:10][#6:11]):[#6:4]:[#6:5]:[#6:6]:1>>[#8:10][#6:11].[#6:1]:1:[#6:2]([#7:7]):[#6:3](-[#6:8](=[#8:9])):[#6:4]:[#6:5]:[#6:6]:1")
        sub = remove_nitrogenous_base.RunReactants((sub,))[0][0]
        nitrogenous_base = sub.GetSubstructMatches(Chem.MolFromSmarts("[#6:1]:1:[#6:2]([#7:7]):[#6:3](-[#6:8](=[#8:9])-[#8:10][#6:11]):[#6:4]:[#6:5]:[#6:6]:1"))
        The_Process.append(sub)
  
    nitrogenous_base = sub.GetSubstructMatches(Chem.MolFromSmarts("[#6:1]:1([#7:7]-[#6:8]):[#6:2]([#8:9][#6:10][#7:11][#6:12]2):[#6:3]2:[#6:4]:[#6:5]:[#6:6]:1"))
    while(len(nitrogenous_base)>0):
        remove_nitrogenous_base = AllChem.ReactionFromSmarts("[#6:1]:1([#7:7]-[#6:8]):[#6:2]([#8:9][#6:10][#7:11][#6:12]2):[#6:3]2:[#6:4]:[#6:5]:[#6:6]:1.[#8:13]>>[#6:8][#8:13].[#6:1]:1([#7:7]):[#6:2]([#8:9][#6:10][#7:11][#6:12]2):[#6:3]2:[#6:4]:[#6:5]:[#6:6]:1")
        sub = remove_nitrogenous_base.RunReactants((sub, Water_coreactant))[0][0]
        nitrogenous_base = sub.GetSubstructMatches(Chem.MolFromSmarts("[#6:1]:1([#7:7]-[#6:8]):[#6:2]:[#6:3]:[#6:4]:[#6:5]:[#6:6]:1"))
        The_Process.append(sub)
 
    nitrogenous_base = sub.GetSubstructMatches(Chem.MolFromSmarts("[#6:0]-[#6:1]:2:[#7:2]:[#6:3]:1:[#6:4]:[#6:5]:[#6:6]:[#6:7]:[#6:8]:1:[#7:9]:2"))
    while(len(nitrogenous_base)>0):
        remove_nitrogenous_base= AllChem.ReactionFromSmarts("[#6:0]-[#6:1]:2:[#7:2]:[#6:3]:1:[#6:4]:[#6:5]:[#6:6]:[#6:7]:[#6:8]:1:[#7:9]:2.[#8:13]>>[#6:0][#8:13].[#6:1]:2:[#7:2]:[#6:3]:1:[#6:4]:[#6:5]:[#6:6]:[#6:7]:[#6:8]:1:[#7:9]:2")
        sub = remove_nitrogenous_base.RunReactants((sub,Water_coreactant))[0][0]
        nitrogenous_base = sub.GetSubstructMatches(Chem.MolFromSmarts("[#6:0]-[#6:1]:2:[#7:2]:[#6:3]:1:[#6:4]:[#6:5]:[#6:6]:[#6:7]:[#6:8]:1:[#7:9]:2"))
        The_Process.append(sub)

    amine_based_sidechain_present = sub.GetSubstructMatches(Chem.MolFromSmarts("[#6:0][#8:1][#6:2](=[#8:3])[#7:4]"))
    while(len(amine_based_sidechain_present)>0):
        resolve_pyrolidine = AllChem.ReactionFromSmarts("[#6:0][#8:1][#6:2](=[#8:3])[#7:4].[#8:5]>>[#6:0][#8:5].[#8:1][#6:2](=[#8:3])[#7:4]")
        sub1 = resolve_pyrolidine.RunReactants((sub,Water_coreactant))
        if(len(re.findall("C",Chem.MolToSmiles(sub1[0][0]),re.IGNORECASE))>len(re.findall("C",Chem.MolToSmiles(sub1[0][1]),re.IGNORECASE))):
            sub=sub1[0][0]
        else:
            sub=sub1[0][1]
        amine_based_sidechain_present = sub.GetSubstructMatches(Chem.MolFromSmarts("[#6:1]-1-[#6:2]-[#6:3]=[#7:4]-[#6:5]-1"))
        The_Process.append(sub)  
        
#     Spongidine_present = sub.GetSubstructMatches(Chem.MolFromSmarts("[+#7:1]:1:[#6:2]:[#6:3](-[#6:7]-[#6:8]-[#6:9]-[#6:10]-2):[#6:4]2:[#6:5]:[#6:6]:1"))
#     while(len(Spongidine_present)>0):
#         resolve_spongidine = AllChem.ReactionFromSmarts("[+#7:1]:1:[#6:2]:[#6:3](-[#6:7]-[#6:8]-[#6:9]-[#6:10]-2):[#6:4]2:[#6:5]:[#6:6]:1.[#8:11]>>[#6:7]=[#6:8]-[#6:9]-[#6:10]-[#8:11].[+#7:1]:1:[#6:2]:[#6:3]:[#6:4]:[#6:5]:[#6:6]:1")
#         sub = resolve_spongidine.RunReactants((sub,Water_coreactant))[0][0]
#         sub = Chem.MolFromSmiles(Chem.MolToSmiles(sub))
#         Spongidine_present = sub.GetSubstructMatches(Chem.MolFromSmarts("[+#7:1]:1:[#6:2]:[#6:3](-[#6:7]-[#6:8]-[#6:9]-[#6:10]-2):[#6:4]2:[#6:5]:[#6:6]:1"))
#         The_Process.append(sub)

    aromatic_mess = sub.GetSubstructMatches(Chem.MolFromSmarts("[#8:0]:[#6:1]:[#7:2]:[#6:3]:[#8:4]:[#6:5]:1:[#6:6]:[#6:7]:[#6:8]:[#6:9]:[#6:10]:1"))
    if(len(aromatic_mess)>0):
        #This solution loses information but RDKit cannot deal with this particular condition:
        # Cc1cc2c(C(C)C)c(O)c(O)c(C=N)c2c(O)c1-c1c(C)cc2c(C(C)C)c(O)c(=O)c3c4[nH]c(oc1c2-3)c1c(=O)c(O)c(C(C)C)c2cc(C)c(-c3c(C)cc5c(C(C)C)c(O)c(O)c(C=O)c5c3O)c(o4)c2-1
        sub = Chem.MolFromSmiles(str.upper(Chem.MolToSmiles(sub)))
        
    aromatic_nitrogen = sub.GetSubstructMatches(Chem.MolFromSmarts("[#6:1]:1([#7:7]-[#6:8]):[#6:2]:[#6:3]:[#6:4]:[#6:5]:[#6:6]:1"))
    while(len(aromatic_nitrogen)>0):
        resolve_aromatic_nitrogen= AllChem.ReactionFromSmarts("[#6:1]:1([#7:7]-[#6:8]):[#6:2]:[#6:3]:[#6:4]:[#6:5]:[#6:6]:1.[#8:13]>>[#6:8][#8:13].[#6:1]:1([#7:7]):[#6:2]:[#6:3]:[#6:4]:[#6:5]:[#6:6]:1")
        sub1 = resolve_aromatic_nitrogen.RunReactants((sub, Water_coreactant))#[0][0]
        if(len(re.findall("C",Chem.MolToSmiles(sub1[0][0]),re.IGNORECASE))<len(re.findall("C",Chem.MolToSmiles(sub1[0][1]),re.IGNORECASE)) and Chem.MolFromSmiles(Chem.MolToSmiles(sub1[0][1]))!=None):
            sub = sub1[0][1]
        else:
            Carbon_number = len(re.findall("C",Chem.MolToSmiles(sub1[0][0]), re.IGNORECASE)); reference=0
            for i in range(0,np.shape(sub1)[0]):
                test = len(re.findall("C", Chem.MolToSmiles(sub1[i][0]), re.IGNORECASE))

                if(test>Carbon_number):
                    Carbon_number=test
                    reference = i
            if(Carbon_number<15):
                break
            sub = sub1[reference][0]
        sub = Chem.MolFromSmiles(Chem.MolToSmiles(sub))
        aromatic_nitrogen = sub.GetSubstructMatches(Chem.MolFromSmarts("[#6:1]:1([#7:7]-[#6:8]):[#6:2]:[#6:3]:[#6:4]:[#6:5]:[#6:6]:1"))
        The_Process.append(sub)
       
    aromatic_nitrogen =sub.GetSubstructMatches(Chem.MolFromSmarts("[#7:1]:1:[#6:2]:[#8:3]:[#6:4]:[#6:5]:1"))
    while(len(aromatic_nitrogen)>0):
        resolve_aromatic_nitrogen = AllChem.ReactionFromSmarts("[#7:1]:1:[#6:2]:[#8:3]:[#6:4]:[#6:5]:1>>[#8:3]-[#6:4][#6:5].[#7:1]-[#6:2]")
        sub =resolve_aromatic_nitrogen.RunReactants((sub,))[0][0]
        sub = Chem.MolFromSmiles(Chem.MolToSmiles(sub))
        aromatic_nitrogen =sub.GetSubstructMatches(Chem.MolFromSmarts("[#7:1]:1:[#6:2]:[#8:3]:[#6:4]:[#6:5]:1"))
        The_Process.append(sub)
        #print("aromatic nitrogen present")

    aromatic_nitrogen =sub.GetSubstructMatches(Chem.MolFromSmarts("[#7:1]:1:[#6:2]:[#6:3](-[#6:7](=[#8:8])-[#8:9]):[#6:4]:[#6:5]:[#6:6]:1"))
    while(len(aromatic_nitrogen)>0):
        resolve_aromatic_nitrogen = AllChem.ReactionFromSmarts("[#7:1]:1:[#6:2]:[#6:3](-[#6:7](=[#8:8])-[#8:9]):[#6:4]:[#6:5]:[#6:6]:1>>[#8:9].[#7:1]:1:[#6:2]:[#6:3](-[#6:7](=[#8:8])):[#6:4]:[#6:5]:[#6:6]:1")
        sub1 =resolve_aromatic_nitrogen.RunReactants((sub,))#[0][0]
        if(Chem.MolToSmiles(sub1[0][0])=="O"):
            sub = sub1[0][1]; aromatic_nitrogen=[]
            break
        else:
            sub = sub1[0][0]
            aromatic_nitrogen =sub.GetSubstructMatches(Chem.MolFromSmarts("[#7:1]:1:[#6:2]:[#6:3](-[#6:7](=[#8:8])-[#8:9]):[#6:4]:[#6:5]:[#6:6]:1"))
        The_Process.append(sub)
        #print("aromatic nitrogen present")        

    aromatic_nitrogen =sub.GetSubstructMatches(Chem.MolFromSmarts("[#7:1]:1:[#6:3](-[#6:7](=[#8:8])-[#8:9]):[#6:4]:[#6:5]:[#6:6]:1"))
    while(len(aromatic_nitrogen)>0):
        resolve_aromatic_nitrogen = AllChem.ReactionFromSmarts("[#7:1]:1:[#6:3](-[#6:7](=[#8:8])-[#8:9]):[#6:4]:[#6:5]:[#6:6]:1>>[#8:9].[#7:1]:1:[#6:3](-[#6:7](=[#8:8])):[#6:4]:[#6:5]:[#6:6]:1")
        sub =resolve_aromatic_nitrogen.RunReactants((sub,))[0][0]
        aromatic_nitrogen =sub.GetSubstructMatches(Chem.MolFromSmarts("[#7:1]:1:[#6:3](-[#6:7](=[#8:8])-[#8:9]):[#6:4]:[#6:5]:[#6:6]:1"))
        The_Process.append(sub)
        #print("aromatic nitrogen present")

    #remove fulminate
    sub = Chem.MolFromSmiles(Chem.MolToSmiles(sub))
    fulminate_present = sub.GetSubstructMatches(Chem.MolFromSmarts("[#6:1]!@[#7:2]=[#6:3]"))
    while(len(fulminate_present)>0):
        resolve_fulminate = AllChem.ReactionFromSmarts("[#6:1]!@[#7:2]=[#6:3].[#8:5]>>[#7:2]=[#6:3].[#6:1][#8:5]")
        sub1 = resolve_fulminate.RunReactants((sub, Water_coreactant))
        if(str(Chem.MolToSmiles(sub1[0][0]))=="N=CO"):
            sub=sub1[0][1];
        elif(len(re.findall("C", Chem.MolToSmiles(sub1[0][1]), re.IGNORECASE))>len(re.findall("C", Chem.MolToSmiles(sub1[0][0]), re.IGNORECASE))):
            sub=sub1[0][1]
        else:
            sub=sub1[0][0]
        sub = Chem.MolFromSmiles(Chem.MolToSmiles(sub))
        fulminate_present = sub.GetSubstructMatches(Chem.MolFromSmarts("[#6:1]!@[#7:2]=[#6:3]"))
        The_Process.append(sub)
        #print("Fulminate present")

    Nitrogenous_sidechain =sub.GetSubstructMatch(Chem.MolFromSmarts("[#6:0]-[#7:1]@2@[#6:2]@[#6:3](@[#6:4]@[#7:5]@[#6:6]1)@[#6:7]@[#6:8]1@[#6:9]@2"))
    while(len(Nitrogenous_sidechain)>0):
        Nitrogenous_sidechain =AllChem.ReactionFromSmarts("[#6:0]-[#7:1]@2@[#6:2]@[#6:3](@[#6:4]@[#7:5]@[#6:6]1)@[#6:7]@[#6:8]1@[#6:9]@2.[#8:10]>>[#6:0]-[#8:10].[#7:1]@2@[#6:2]@[#6:3](@[#6:4]@[#7:5]@[#6:6]1)@[#6:7]@[#6:8]1@[#6:9]@2")
        sub = Nitrogenous_sidechain.RunReactants((sub,Water_coreactant))[0][0]
        sub = Chem.MolFromSmiles(Chem.MolToSmiles(sub))
        Nitrogenous_sidechain =sub.GetSubstructMatch(Chem.MolFromSmarts("[#6:0]-[#7:1]@2@[#6:2]@[#6:3](@[#6:4]@[#7:5]@[#6:6]1)@[#6:7]@[#6:8]1@[#6:9]@2"))
        The_Process.append(sub)

    sub = Chem.MolFromSmiles(Chem.MolToSmiles(sub))
    Nitrogenous_sidechain =sub.GetSubstructMatches(Chem.MolFromSmarts("[#6:1]!@[#7:2]=[#6:3]"))
    while(len(Nitrogenous_sidechain)>0):
        Nitrogenous_sidechain =AllChem.ReactionFromSmarts("[#6:1]!@[#7:2]=[#6:3]>>[#6:1]-[#7:2].[#6:3]")
        sub1 = Nitrogenous_sidechain.RunReactants((sub,))   
        if(len(re.findall("C",Chem.MolToSmiles(sub1[0][0])))<10):
            sub = sub1[0][1]
        else:
            sub = sub1[0][0]
        sub = Chem.MolFromSmiles(Chem.MolToSmiles(sub))
        Nitrogenous_sidechain =sub.GetSubstructMatches(Chem.MolFromSmarts("[#6:1]!:[#7:2]=[#6:3]"))
        The_Process.append(sub)
        #print("Nitrogenous sidechain")

    Nitrogen_bridge = sub.GetSubstructMatches(Chem.MolFromSmarts("[#7:1](!:[#6:2])(!:[#6:3])!:[#6:4]"))
    if(len(Nitrogen_bridge)>0):
        counter =1
        resolve_nitrogen = AllChem.ReactionFromSmarts("[#7:1]!:[#6:2]>>[#7:1].[#6:2]")
        while(counter==1):
            sub1 = resolve_nitrogen.RunReactants((sub,))
            counter=0
            side_chain =0
            for i in sub1:
                if(len(re.findall("C",Chem.MolToSmiles(i[1])))%5==0 and len(re.findall("C",Chem.MolToSmiles(i[1])))>=15):
                    sub = i[1]
                    break;

                elif(len(re.findall("C", Chem.MolToSmiles(i[1])))>side_chain):
                    side_chain = len(re.findall("C", Chem.MolToSmiles(i[1])))
                    sub = i[1]
            sub = Chem.MolFromSmiles(Chem.MolToSmiles(sub))
            Nitrogen_bridge = sub.GetSubstructMatches(Chem.MolFromSmarts("[#7:1](!:[#6:2])(!:[#6:3])!:[#6:4]"))
            The_Process.append(sub)
            if(len(Nitrogen_bridge)>0):
                counter=1
            The_Process.append(sub)
            #print("Nitrogen bridge present1")
    sub = Chem.MolFromSmiles(Chem.MolToSmiles(sub))
    Nitrogen_bridge = sub.GetSubstructMatches(Chem.MolFromSmarts("[#6:1]!@[#7:2]!@[#6:3]"))
    while(len(Nitrogen_bridge)>0):
        resolve_nitrogen = AllChem.ReactionFromSmarts("[#6:1]!@[#7:2]!@[#6:3].[#8:4]>>[#6:1][#8:4].[#7:2][#6:3]")
        sub1 = resolve_nitrogen.RunReactants((sub,Water_coreactant))
        for i in sub1:
            if(len(re.findall("C", Chem.MolToSmiles(i[0])))>len(re.findall("C", Chem.MolToSmiles(i[1])))):
                sub = i[0]
            else:
                sub = i[1]
        sub = Chem.MolFromSmiles(Chem.MolToSmiles(sub))
        Nitrogen_bridge = sub.GetSubstructMatches(Chem.MolFromSmarts("[#6:1]!@[#7:2]!@[#6:3]"))
        #print("Nitrogen Bridge Present2")
        The_Process.append(sub)

    sub = Chem.MolFromSmiles(Chem.MolToSmiles(sub))
    nitrogen_ring = sub.GetSubstructMatches(Chem.MolFromSmarts("[#7:1]!:1!:[#6:2]!:[#6:3]!:[#6:4]!:[#6:5]!:[#6:6]!:1"))
    while(len(nitrogen_ring)>0): 
        resolve_nitrogen_ring = AllChem.ReactionFromSmarts("[#7:1]!:1!:[#6:2]!:[#6:3]!:[#6:4]!:[#6:5]!:[#6:6]!:1>>[#7:1]-[#6:2]-[#6:3]-[#6:4]-[#6:5]-[#6:6]")
        sub = resolve_nitrogen_ring.RunReactants((sub,))[0][0]
        sub = Chem.MolFromSmiles(Chem.MolToSmiles(sub))
        nitrogen_ring = sub.GetSubstructMatches(Chem.MolFromSmarts("[#7:1]!:1!:[#6:2]!:[#6:3]!:[#6:4]!:[#6:5]!:[#6:6]!:1"))
        The_Process.append(sub)
   
    #resolve Nitrogen
    double_bond_nitrogen = sub.GetSubstructMatches(Chem.MolFromSmarts("[#6:1]=[#7:2]"))
    if(len(double_bond_nitrogen)>0):
        counter=1
        while(counter==1):
            resolve_nitrogen = AllChem.ReactionFromSmarts("[#6:1]=[#7:2].[#8:3]>>[#6:1]=[#8:3].[#7:2]")
            sub1 = resolve_nitrogen.RunReactants((sub,Water_coreactant))
            counter =0
            for i in sub1:
                if(str(Chem.MolToSmiles(i[1]))=="N"):
                    sub=i[0]
                    counter =1; break
                elif(len(re.findall("C", Chem.MolToSmiles(i[1])))<5):
                    sub=i[0]
                else:
                    counter=0
            The_Process.append(sub)

   
    nitrogen_present = sub.GetSubstructMatches(Chem.MolFromSmarts("[#7:0]-[#8:1]-[#6:2]"))
    while(len(nitrogen_present)>0):
        remove_nitrogen =AllChem.ReactionFromSmarts("[#7:0]-[#8:1]-[#6:2].[#8:3]>>[#7:0]-[#8:1].[#8:3][#6:2]")
        sub1 = remove_nitrogen.RunReactants((sub,Water_coreactant))
        if(Chem.MolToSmiles(sub1[0][1])=="CO"):
            sub = sub1[0][0]
        elif(len(re.findall("C",Chem.MolToSmiles(sub1[0][0]),re.IGNORECASE))>len(re.findall("C",Chem.MolToSmiles(sub1[0][1]),re.IGNORECASE))):
            sub = sub1[0][0]
        elif(len(re.findall("C",Chem.MolToSmiles(sub1[0][0]),re.IGNORECASE))<len(re.findall("C",Chem.MolToSmiles(sub1[0][1]),re.IGNORECASE))):
            sub = sub1[0][1]
        else:
            sub = sub1[0][1]
        nitrogen_present = sub.GetSubstructMatches(Chem.MolFromSmarts("[#7:0]-[#8:1]-[#6:2]"))
        The_Process.append
        
    sub = Chem.MolFromSmiles(Chem.MolToSmiles(sub))
    terminal_nitrogen = sub.GetSubstructMatches(Chem.MolFromSmarts("[#6:1]!@[#7:2]"))
    while(len(terminal_nitrogen)>0):
        remove_terminal_nitrogen = AllChem.ReactionFromSmarts("[#6:1]!@[#7:2].[#8:3]>>[#6:1]-[#8:3].[#7:2]")
        sub1 = remove_terminal_nitrogen.RunReactants((sub,Water_coreactant))
        if(len(re.findall("C", Chem.MolToSmiles(sub1[0][1])))<5):
            sub=sub1[0][0]
            sub = Chem.MolFromSmiles(Chem.MolToSmiles(sub))
            terminal_nitrogen = sub.GetSubstructMatches(Chem.MolFromSmarts("[#6:1]!@[#7:2]"))
        else:
            terminal_nitrogen =[]
        The_Process.append(sub)

    # remove aromatic adjacent nitrogens
    sub = Chem.MolFromSmiles(Chem.MolToSmiles(sub))
    aromatic_nitrogen = sub.GetSubstructMatches(Chem.MolFromSmarts("[#6:0]:[#6:1]-[#7:2]@[#6:4]"))
    while(len(aromatic_nitrogen)>0):
        sub = Chem.MolFromSmiles(Chem.MolToSmiles(sub))
        remove_aromatic_nitrogen = AllChem.ReactionFromSmarts("[#6:0]:[#6:1]-[#7:2].[#8:4]>>[#6:0]:[#6:1]-[#8:4].[#7:2]")
        sub = remove_aromatic_nitrogen.RunReactants((sub,Water_coreactant))[0][0]
        sub = Chem.MolFromSmiles(Chem.MolToSmiles(sub))
        aromatic_nitrogen = sub.GetSubstructMatches(Chem.MolFromSmarts("[#6:0]:[#6:1]-[#7:2]@[#6:4]"))
        The_Process.append(sub)     

    # resolve non-aromatic ring nitrogens
    sub = Chem.MolFromSmiles(Chem.MolToSmiles(sub))
    ring_nitrogen = sub.GetSubstructMatches(Chem.MolFromSmarts("[#6:1]@[#7:2]@[#6:3]"))
    aromatic_nitrogen1 = sub.GetSubstructMatches(Chem.MolFromSmarts("[#6:0]:[#6:1]-[#7:2]=[#6:3]"))
    aromatic_nitrogen2 = sub.GetSubstructMatches(Chem.MolFromSmarts("[#6:0]:[#7:2]"))
    while(len(aromatic_nitrogen1)==0 and len(aromatic_nitrogen2)==0 and len(ring_nitrogen)>0):
        break_ring = AllChem.ReactionFromSmarts("[#6:1]@[#7:2]@[#6:3].[#8:4].[#8:5]>>[#6:1][#8:4][#7:2][#8:5][#6:3]")
        resolve_nitrogen = AllChem.ReactionFromSmarts("[#6:0]@[#8:3]@[#7:2]>>[#6:0][#8:3].[#7:2]") 
        sub1 = break_ring.RunReactants((sub, Water_coreactant,Water_coreactant))
        if(np.shape(sub1)[0]>2):
            if(Chem.MolFromSmiles(Chem.MolToSmiles(sub1[0][0]))==None and Chem.MolFromSmiles(Chem.MolToSmiles(sub1[1][0]))!=None):
                sub1 =Chem.MolFromSmiles(Chem.MolToSmiles(sub1[1][0]))
            else:
                sub1 =Chem.MolFromSmiles(Chem.MolToSmiles(sub1[0][0]))
        else:
            sub1= Chem.MolFromSmiles(Chem.MolToSmiles(sub1[0][0]))
        sub = resolve_nitrogen.RunReactants((sub1,))[0][0]
        sub = Chem.MolFromSmiles(Chem.MolToSmiles(sub))
        ring_nitrogen = sub.GetSubstructMatches(Chem.MolFromSmarts("[#6:1]@[#7:2]@[#6:3]"))
        The_Process.append(sub)

           
    # remove aromatic adjacent nitrogens
    sub = Chem.MolFromSmiles(Chem.MolToSmiles(sub))
    aromatic_nitrogen = sub.GetSubstructMatches(Chem.MolFromSmarts("[#6:0]:[#6:1]-[#7:2]=[#6:4]"))
    while(len(aromatic_nitrogen)>0):
        remove_aromatic_nitrogen = AllChem.ReactionFromSmarts("[#6:0]:[#6:1]-[#7:2].[#8:4]>>[#6:0]:[#6:1]-[#8:4].[#7:2]")
        sub = remove_aromatic_nitrogen.RunReactants((sub,Water_coreactant))[0][0]
        aromatic_nitrogen = sub.GetSubstructMatches(Chem.MolFromSmarts("[#6:0]:[#6:1]-[#7:2]=[#6:4]"))
        The_Process.append(sub)
        
    # remove cyanide
    sub = Chem.MolFromSmiles(Chem.MolToSmiles(sub))
    Cyanide_present = sub.GetSubstructMatches(Chem.MolFromSmarts("[#6:1]#[#7:2]!@[#6:3]"))
    while(len(Cyanide_present)>0):
        remove_cyanide = AllChem.ReactionFromSmarts("[#6:1]#[#7:2]!@[#6:3].[#8:4]>>[#6:3][#8:4].[#6:1]#[#7:2]")
        sub = remove_cyanide.RunReactants((sub, Water_coreactant))[0][0]
        sub = Chem.MolFromSmiles(Chem.MolToSmiles(sub))
        Cyanide_present = sub.GetSubstructMatches(Chem.MolFromSmarts("[#6:1]#[#7:2]!@[#6:3]"))
        The_Process.append(sub)
        
    ###################################################################################################

    tanshinlactone_present = sub.GetSubstructMatches(Chem.MolFromSmarts("[#6:1]:1:[#6:2]:[#6:3]:[#6:4]:[#6:5]:2:[#6:6]:1:[#6:7]:[#6:8]:[#6:9]:4:[#6:10]:2:[#6:11](=[#8:12]):[#8:13]:[#6:14]:3:[#6:15]:[#6:16]:[#8:17]:[#6:18]:3:4"))
    #https://doi.org/10.1039/C0NP00035C
    while(len(tanshinlactone_present)>0):
        tanshinlactone_condition = AllChem.ReactionFromSmarts('[#6:1]:1:[#6:2]:[#6:3]:[#6:4]:[#6:5]:2:[#6:6]:1:[#6:7]:[#6:8]:[#6:9]:4:[#6:10]:2:[#6:11](=[#8:12]):[#8:13]:[#6:14]:3:[#6:15]([#6:20]):[#6:16]:[#8:17]:[#6:18]:3:4.[#8:19]>>[#6:1]-1=[#6:2]-[#6:3]=[#6:4]-[#6:5]-2=[#6:6]-1-[#6:7]=[#6:8]-[#6:9](-[#6:14](=[#8:17])-[#6:15]-[#6:16](-[#6:20])-[#6:18](=[#8:19]))=[#6:10]-2-[#6:11](=[#8:12])-[#8:13]')
        sub = tanshinlactone_condition.RunReactants((sub,Water_coreactant))[0][0]
        tanshinlactone_present = sub.GetSubstructMatches(Chem.MolFromSmarts("[#6:1]:1:[#6:2]:[#6:3]:[#6:4]:[#6:5]:2:[#6:6]:1:[#6:7]:[#6:8]:[#6:9]:4:[#6:10]:2:[#6:11](=[#8:12]):[#8:13]:[#6:14]:3:[#6:15]:[#6:16]:[#8:17]:[#6:18]:3:4"))
        #print("tanshinlactone present")
        The_Process.append(sub)  
          
    neotanshinlactone_present = sub.GetSubstructMatches(Chem.MolFromSmarts("[#6:1]:1:[#6:2]:[#6:3]:[#6:4]:[#6:5]:2:[#6:6]:1:[#6:7]:[#6:8]:[#6:9]:4:[#6:10]:2:[#8:13]:[#6:11](=[#8:12]):[#6:14]:3:[#6:15]:[#6:16]:[#8:17]:[#6:18]:3:4"))
    #https://doi.org/10.1039/C0NP00035C
    while(len(neotanshinlactone_present)>0):
        neotanshinlactone_condition = AllChem.ReactionFromSmarts('[#6:1]:1:[#6:2]:[#6:3]:[#6:4]:[#6:5]:2:[#6:6]:1:[#6:7]:[#6:8]:[#6:9]:4:[#6:10]:2:[#8:13]:[#6:11](=[#8:12]):[#6:14]:3:[#6:15]:[#6:16]:[#8:17]:[#6:18]:3:4>>[#6:1]-1=[#6:2]-[#6:3]=[#6:4]-[#6:5]-2=[#6:6]-1-[#6:7]=[#6:8]-[#6:9](-[#6:18]=3-[#8:17]-[#6:16]=[#6:15]-[#6:14]=3-[#6:11](-[#8:13])(=[#8:12]))=[#6:10]-2')
        sub = neotanshinlactone_condition.RunReactants((sub,))[0][0]
        neotanshinlactone_present = sub.GetSubstructMatches(Chem.MolFromSmarts("[#6:1]:1:[#6:2]:[#6:3]:[#6:4]:[#6:5]:2:[#6:6]:1:[#6:7]:[#6:8]:[#6:9]:4:[#6:10]:2:[#8:13]:[#6:11](=[#8:12]):[#6:14]:3:[#6:15]:[#6:16]:[#8:17]:[#6:18]:3:4"))
        #print("neotanshinlactone present")
        The_Process.append(sub) 
        
    swartziarboreal_present =sub.GetSubstructMatches(Chem.MolFromSmarts("[#6:1]@1(=[#8:2])@[#8:3]@[#6:4]@[#6:5]@[#6:6]@[#6:7]@1"))
    # https://doi.org/10.26850/1678-4618eqj.v35.3.2010.p169-177
    while(len(swartziarboreal_present) > 0):
        swartziarboreol_condition = AllChem.ReactionFromSmarts("[#6:1]@1(=[#8:2])@[#8:3]@[#6:4]@[#6:5]@[#6:6]@[#6:7]@1.[#8:8]>>[#8:8]-[#6:4][#6:5][#6:6][#6:7][#6:1](=[#8:2])-[#8:3]")
        sub1 = swartziarboreol_condition.RunReactants((sub, Water_coreactant))[0][0]
        if(Chem.MolFromSmiles(Chem.MolToSmiles(sub1))!=None):
            sub=Chem.MolFromSmiles(Chem.MolToSmiles(sub1))
        else:break
        swartziarboreal_present =sub.GetSubstructMatches(Chem.MolFromSmarts("[#6:1]@1(=[#8:2])@[#8:3]@[#6:4]@[#6:5]@[#6:6]@[#6:7]@1"))
        #print("Swartziarboreal exception")
        The_Process.append(sub)
        
    hepta_abietane_present = sub.GetSubstructMatches(Chem.MolFromSmarts("[#6:1](=[#8:8]):1:[#6:2]:[#6:3]:[#6:4]:[#6:5]:[#6:6]:[#6:7]:1"))
    #https://doi.org/10.1021/np040219h
    while(len(hepta_abietane_present)>0):
        hepta_abietane_condition = AllChem.ReactionFromSmarts('[#6:1](=[#8:8]):1:[#6:2]:[#6:3]:[#6:4]:[#6:5]:[#6:6]:[#6:7]:1>>[#6:1](-[#8:8])-1-[#6:2]=[#6:3]-[#6:4]=[#6:5]-[#6:6]=[#6:7]-1')
        sub = hepta_abietane_condition.RunReactants((sub,))[0][0]
        hepta_abietane_present = sub.GetSubstructMatches(Chem.MolFromSmarts("[#6:1](=[#8:8]):1:[#6:2]:[#6:3]:[#6:4]:[#6:5]:[#6:6]:[#6:7]:1"))
        #print("hepta_abietane_present")
        The_Process.append(sub)  

    avinosol_present = sub.GetSubstructMatches(Chem.MolFromSmarts("[#6:1]-[#7:2]:2:[#6:3]:[#7:4]:[#6:5]:1:[#7:6]:[#6:7]:[#7:8]:[#6:9]:1:[#6:10]:2"))
    #https://doi.org/10.1021/ol061333p
    while(len(avinosol_present)>0):
        avinosol_condition = AllChem.ReactionFromSmarts('[#6:1]-[#7:2]:2:[#6:3]:[#7:4]:[#6:5]:1:[#7:6]:[#6:7]:[#7:8]:[#6:9]:1:[#6:10]:2>>[#6:1].[#7:2]:2:[#6:3]:[#7:4]:[#6:5]:1:[#7:6]:[#6:7]:[#7:8]:[#6:9]:1:[#6:10]:2')
        sub = avinosol_condition.RunReactants((sub,))[0][0]
        avinosol_present = sub.GetSubstructMatches(Chem.MolFromSmarts("[#6:1]-[#7:2]:2:[#6:3]:[#7:4]:[#6:5]:1:[#7:6]:[#6:7]:[#7:8]:[#6:9]:1:[#6:10]:2"))
        #print("avinosol present")
        The_Process.append(sub)

    #replace halogen(s) with water

    halogen_present = sub.GetSubstructMatches(Chem.MolFromSmarts("[#6:1][F,Cl,Br,I:3]"))  
    while(len(halogen_present)>0):
        halogen_condition = AllChem.ReactionFromSmarts("[#6:1][F,Cl,Br,I:3].[#8:2]>>[#6:1][#8:2].[F,Cl,Br,I:3]")
        sub = halogen_condition.RunReactants((sub, Water_coreactant))[0][0]
        sub = Chem.MolFromSmiles(Chem.MolToSmiles(sub))
        halogen_present = sub.GetSubstructMatches(Chem.MolFromSmarts("[#6:1][F,Cl,Br,I:3]")) 
        The_Process.append(sub)      

    #replace halogen(s) with water
    halogen_present = sub.GetSubstructMatches(Chem.MolFromSmarts("[#6:1]-[#8:2][F,Cl,Br,I:3]"))  
    while(len(halogen_present)>0):
        halogen_condition = AllChem.ReactionFromSmarts("[#6:1]-[#8:2][F,Cl,Br,I:3]>>[#6:1]-[#8:2].[F,Cl,Br,I:3]")
        sub = halogen_condition.RunReactants((sub,))[0][0]
        halogen_present = sub.GetSubstructMatches(Chem.MolFromSmarts("[#6:1]-[#8:2][F,Cl,Br,I:3]")) 
        The_Process.append(sub)  
  
    #break Ester bonds
    sub = Chem.MolFromSmiles(Chem.MolToSmiles(sub))
    Ester_bonds = sub.GetSubstructMatches(Chem.MolFromSmarts('[#6:1](=[#8:2])-[#8:3]-[#6:4]'))
    while(len(Ester_bonds) > 0):
        sub = Chem.MolFromSmiles(Chem.MolToSmiles(sub))
        rxn = AllChem.ReactionFromSmarts('[#6:1](=[#8:2])-[#8:3]-[#6:4].[#8:5]>>[#6:1](-[#8:2])-[#8:5]-[#8:3]-[#6:4]')
        rxn2 = AllChem.ReactionFromSmarts('[#8:1]-[#8:2]>>[#8:1].[#8:2]')
        sub1 = rxn.RunReactants((sub, Water_coreactant))[0][0]
        sub1 = Chem.MolFromSmiles(Chem.MolToSmiles(sub1))
        sub2 = rxn2.RunReactants((sub1,))
        #carbon_number = len(re.findall("C",Chem.MolToSmiles(sub2[0][1]), re.IGNORECASE))
        if(len(re.findall("C",Chem.MolToSmiles(sub2[0][1]), re.IGNORECASE))==20):
            sub = sub2[0][1]
        elif(len(re.findall("C",Chem.MolToSmiles(sub2[0][0]), re.IGNORECASE))==20):
            sub = sub2[0][0]
        elif(len(re.findall("C",Chem.MolToSmiles(sub2[0][1]), re.IGNORECASE))>len(re.findall("C",Chem.MolToSmiles(sub2[0][0]), re.IGNORECASE))):
            sub = sub2[0][1]
        else:
            sub = sub2[0][0]
        The_Process.append(sub)
        sub = Chem.MolFromSmiles(Chem.MolToSmiles(sub))
        Ester_bonds = sub.GetSubstructMatches(Chem.MolFromSmarts('[#6:1](=[#8:2])[#8:3]-[#6:4]'))
        #print("succesfully broke ester bonds")
 
    covalent_O = sub.GetSubstructMatches(Chem.MolFromSmarts("[#8:1][#8:2]"))
    while(len(covalent_O)>0):
        rxn2 = AllChem.ReactionFromSmarts('[#8:1][#8:2]>>[#8:1].[#8:2]')
        sub1 = rxn2.RunReactants((sub,))
        if(len(re.findall("C",Chem.MolToSmiles(sub1[0][0])))>len(re.findall("C",Chem.MolToSmiles(sub1[0][1])))):
            sub = sub1[0][0]
        else:
            sub = sub1[0][1]
        covalent_O = sub.GetSubstructMatches(Chem.MolFromSmarts("[#8:1][#8:2]"))
    
    
    #reduce ketones/aldehydes
    sub = Chem.MolFromSmiles(Chem.MolToSmiles(sub))
    double_bonds = sub.GetSubstructMatches(Chem.MolFromSmarts("[#6:0]!:[#6:1]=[#8:2]"))
    while(len(double_bonds) > 0):
        rxn3= AllChem.ReactionFromSmarts("[#6:0]!:[#6:1]=[#8:2]>>[#6:0]-[#6:1]-[#8:2]")
        sub = rxn3.RunReactants((sub,))[0][0]
        sub = Chem.MolFromSmiles(Chem.MolToSmiles(sub))
        double_bonds = sub.GetSubstructMatches(Chem.MolFromSmarts("[#6:0]!:[#6:1]=[#8:2]"))
        The_Process.append(sub)
        #print("succesfully reduced all ketones and aldehydes")        
        
    #remove oxygen when more than one on a carbon
    sub = Chem.MolFromSmiles(Chem.MolToSmiles(sub))
    hydroxy2many = sub.GetSubstructMatches(Chem.MolFromSmarts("[#6:1](!@[#8:2])!@[#8:3]"))
    while(len(hydroxy2many) > 0):
        rxn3= AllChem.ReactionFromSmarts("[#6:1](!@[#8:2])!@[#8:3]>>[#6:1](-[#8:2]).[#8:3]")
        sub1 = rxn3.RunReactants((sub,))
        if(Chem.MolToSmiles(sub1[0][1])=='O'):
            sub = sub1[0][0]
        elif(Chem.MolToSmiles(sub1[0][1])=='CO'):
            sub = sub1[0][0]
        elif(len(re.findall("C",Chem.MolToSmiles(sub1[0][0]))) < 10):
            sub = sub1[0][1]
        else:
            sub = sub1[0][0]
        sub = Chem.MolFromSmiles(Chem.MolToSmiles(sub))
        hydroxy2many = sub.GetSubstructMatches(Chem.MolFromSmarts("[#6:1](!@[#8:2])!@[#8:3]"))
        The_Process.append(sub)
        #print("Succesfully removed assumed P450 hydroxyl groups")
    
    # resolve epoxy groups
    epoxy_present = sub.GetSubstructMatches(Chem.MolFromSmarts("[#6:1]-1-[#8:2]-[#6:3]-1"))
    resolve_epoxy = AllChem.ReactionFromSmarts("[#6:1]-1-[#8:2]-[#6:3]-1.[#8:4]>>[#6:1]-[#6:3][#8:4].[#8:2]")
    while(len(epoxy_present)>0):
        sub = resolve_epoxy.RunReactants((sub, Water_coreactant))[0][0]
        epoxy_present = sub.GetSubstructMatches(Chem.MolFromSmarts("[#6:1]-1-[#8:2]-[#6:3]-1"))
        The_Process.append(sub)
  
    # remove methanol groups
    sub = Chem.MolFromSmiles(Chem.MolToSmiles(sub))
    methanol_present = sub.GetSubstructMatches(Chem.MolFromSmarts("[#6:1]!@[#8:2]!@[#6:3]"))
    remove_methanol= AllChem.ReactionFromSmarts("[#6:1]!@[#8:2]!@[#6:3].[#8:4]>>[#6:1]-[#8:2].[#6:3][#8:4]")
    if(len(methanol_present)>0):
        counter = 1
        while(counter==1):
            sub = Chem.MolFromSmiles(Chem.MolToSmiles(sub))
            sub1 = remove_methanol.RunReactants((sub, Water_coreactant))
            counter=0
            for i in sub1:
                if(Chem.MolToSmiles(i[0])=="CO"):
                    sub = i[1]
                    counter = 1; break
                elif(Chem.MolToSmiles(i[1])=="CO"):
                    sub = i[0]
                    counter = 1; break
                elif(Chem.MolToSmiles(i[0])=="CCO"):
                    sub = i[1]
                    counter = 1; break
                elif(Chem.MolToSmiles(i[0])=="CC(C)O"):
                    sub = i[1]
                    counter = 1; break
                elif(len(re.findall("C", Chem.MolToSmiles(i[0]), re.IGNORECASE))<len(re.findall("C", Chem.MolToSmiles(i[1]), re.IGNORECASE))):
                    sub = i[1]
                    counter = 1; break
                else:
                    counter =0

            The_Process.append(sub)
        #print("removed methanol")

    # remove methanol groups
    methanol_present = sub.GetSubstructMatches(Chem.MolFromSmarts("[#6:1][#7:2]-[#8:3]-[#6:4]"))
    remove_methanol= AllChem.ReactionFromSmarts("[#6:1][#7:2]-[#8:3]-[#6:4].[#8:5]>>[#6:1]-[#8:5].[#7:2]-[#8:3]-[#6:4]")
    if(len(methanol_present)>0):
        counter = 1
        while(counter==1):
            sub1 = remove_methanol.RunReactants((sub, Water_coreactant))
            counter=0
            for i in sub1:
                if(Chem.MolToSmiles(i[1])=="CON"):
                    sub = i[0]
                    counter = 1; break
                else:
                    counter =0
            The_Process.append(sub)
        #print("removed methanol")
 
    sub = Chem.MolFromSmiles(Chem.MolToSmiles(sub))
    cyclic_ether_present=sub.GetSubstructMatches(Chem.MolFromSmarts("[#8:1]!:1!:[#6:2]!:[#6:3]!:[#6:4]!:[#6:5]!:1"))
    while(len(cyclic_ether_present)>0):
        break_ether = AllChem.ReactionFromSmarts("[#8:1]!:1!:[#6:2]!:[#6:3]!:[#6:4]!:[#6:5]!:1.[#8:6]>>[#8:1]-[#6:2]-[#6:3]-[#6:4]-[#6:5][#8:6]")
        sub = break_ether.RunReactants((sub,Water_coreactant))[0][0]
        sub = Chem.MolFromSmiles(Chem.MolToSmiles(sub))
        cyclic_ether_present=sub.GetSubstructMatches(Chem.MolFromSmarts("[#8:1]!:1!:[#6:2]!:[#6:3]!:[#6:4]!:[#6:5]!:1"))
        The_Process.append(sub)
        
    cyclic_ether_present=sub.GetSubstructMatches(Chem.MolFromSmarts("[#8:1]-1-[#6:2]-[#6:3]-[#6:4]-1"))
    while(len(cyclic_ether_present)>0):
        break_ether = AllChem.ReactionFromSmarts("[#8:1]-1-[#6:2]-[#6:3]-[#6:4]-1.[#8:5]>>[#8:1]-[#6:2]-[#6:3]-[#6:4][#8:5]")
        sub = break_ether.RunReactants((sub, Water_coreactant))[0][0]
        cyclic_ether_present=sub.GetSubstructMatches(Chem.MolFromSmarts("[#8:1]-1-[#6:2]-[#6:3]-[#6:4]-[#6:5]-1"))
        The_Process.append(sub) 

    sub = Chem.MolFromSmiles(Chem.MolToSmiles(sub))
    oxygen_ring = sub.GetSubstructMatches(Chem.MolFromSmarts("[#8:2]!:1!:[#6:1]!:[#8:5]!:[#6:3]!:[#6:4]!:[#6:6]!:[#6:7]!:1"))
    while(len(oxygen_ring)>0): #and len(aromatic_oxygen_ring)==0):
        oxygen_ring_resolve = AllChem.ReactionFromSmarts("[#8:2]!:1!:[#6:1]!:[#8:5]!:[#6:3]!:[#6:4]!:[#6:6]!:[#6:7]!:1>>[#8:2]-[#6:1]-[#8:5]-[#6:3]-[#6:4]-[#6:6]-[#6:7]")
        sub = oxygen_ring_resolve.RunReactants((sub,))[0][0]
        sub = Chem.MolFromSmiles(Chem.MolToSmiles(sub))
        oxygen_ring = sub.GetSubstructMatches(Chem.MolFromSmarts("[#8:2]!:1!:[#6:1]!:[#8:5]!:[#6:3]!:[#6:4]!:[#6:6]!:[#6:7]!:1"))
        The_Process.append(sub)     
        
    sub = Chem.MolFromSmiles(Chem.MolToSmiles(sub))
    oxygen_ring = sub.GetSubstructMatches(Chem.MolFromSmarts("[#8:2]!:1!:[#6:1]!:[#8:5]!:[#6:3]!:[#6:4]!:[#6:6]!:1"))
    while(len(oxygen_ring)>0): #and len(aromatic_oxygen_ring)==0):
        oxygen_ring_resolve = AllChem.ReactionFromSmarts("[#8:2]!:1!:[#6:1]!:[#8:5]!:[#6:3]!:[#6:4]!:[#6:6]!:1>>[#8:2]-[#6:1]-[#8:5]-[#6:3]-[#6:4]-[#6:6]")
        sub = oxygen_ring_resolve.RunReactants((sub,))[0][0]
        sub = Chem.MolFromSmiles(Chem.MolToSmiles(sub))
        oxygen_ring = sub.GetSubstructMatches(Chem.MolFromSmarts("[#8:2]!:1!:[#6:1]!:[#8:5]!:[#6:3]!:[#6:4]!:[#6:6]!:1"))
        The_Process.append(sub)

    sub = Chem.MolFromSmiles(Chem.MolToSmiles(sub))
    oxygen_ring = sub.GetSubstructMatches(Chem.MolFromSmarts("[#8:2]@1@[#6:1]@[#6:3]@[#8:5]@[#6:4]@[#6:6]@1"))
    while(len(oxygen_ring)>0):
        oxygen_ring_resolve = AllChem.ReactionFromSmarts("[#8:2]@1@[#6:1]@[#6:3]@[#8:5]@[#6:4]@[#6:6]@1>>[#8:2][#6:1][#6:3][#8:5][#6:4][#6:6]")
        sub = oxygen_ring_resolve.RunReactants((sub,))[0][0]
        sub = Chem.MolFromSmiles(Chem.MolToSmiles(sub))
        oxygen_ring = sub.GetSubstructMatches(Chem.MolFromSmarts("[#8:2]@1@[#6:1]@[#6:3]@[#8:5]@[#6:4]@[#6:6]@1"))
        The_Process.append(sub)

    sub = Chem.MolFromSmiles(Chem.MolToSmiles(sub))
    oxygen_ring = sub.GetSubstructMatches(Chem.MolFromSmarts("[#8:2]!:1!:[#6:1]!:[#8:5]!:[#6:3]!:[#6:4]!:1"))
    while(len(oxygen_ring)>0): #and len(aromatic_oxygen_ring)==0):
        oxygen_ring_resolve = AllChem.ReactionFromSmarts("[#8:2]!:1!:[#6:1]!:[#8:5]!:[#6:3]!:[#6:4]!:1>>[#8:2]-[#6:1]-[#8:5]-[#6:3]-[#6:4]")
        sub = oxygen_ring_resolve.RunReactants((sub,))[0][0]
        sub = Chem.MolFromSmiles(Chem.MolToSmiles(sub))
        oxygen_ring = sub.GetSubstructMatches(Chem.MolFromSmarts("[#8:2]!:1!:[#6:1]!:[#8:5]!:[#6:3]!:[#6:4]!:1"))
        The_Process.append(sub)


    ether_present=sub.GetSubstructMatches(Chem.MolFromSmarts("[#6:1]!@[#8:2]!@[#6:3]"))
    while(len(ether_present)>0):
        break_ether=AllChem.ReactionFromSmarts("[#6:1]!@[#8:2]!@[#6:3].[#8:4]>>[#6:1][#8:4].[#8:2]-[#6:3]")
        sub1 = break_ether.RunReactants((sub,Water_coreactant))
        if(len(re.findall("C",Chem.MolToSmiles(sub1[0][0])))>len(re.findall("C",Chem.MolToSmiles(sub1[0][1])))):
            sub = sub1[0][0]
        else: sub = sub1[0][1]
        sub = Chem.MolFromSmiles(Chem.MolToSmiles(sub))
        ether_present=sub.GetSubstructMatches(Chem.MolFromSmarts("[#6:1]!@[#8:2]!@[#6:3]"))
        The_Process.append(sub)    
    
    ## Edge Case Cleanup
    sub = Chem.MolFromSmiles(Chem.MolToSmiles(sub))
    nitrogen_present = sub.GetSubstructMatches(Chem.MolFromSmarts("[#7:1]:1:[#6:2](!@[#6:7]):[#6:3]:[#6:4]:[#6:5]:[#6:6]:1"))
    while(len(nitrogen_present)>0):
        remove_nitrogen = AllChem.ReactionFromSmarts("[#7:1]:1:[#6:2](!@[#6:7]):[#6:3]:[#6:4]:[#6:5]:[#6:6]:1>>[#7:1]-[#6:2](-[#6:7]).[#6:3]-[#6:4]-[#6:5]-[#6:6]")
        if(Chem.MolFromSmiles(Chem.MolToSmiles(remove_nitrogen.RunReactants((sub,))[0][1]))!=None):
            sub = remove_nitrogen.RunReactants((sub,))[0][1]
        elif(Chem.MolFromSmiles(Chem.MolToSmiles(remove_nitrogen.RunReactants((sub,))[0][0]))!=None):
            sub = remove_nitrogen.RunReactants((sub,))[0][0]

        else:break
        sub = Chem.MolFromSmiles(Chem.MolToSmiles(sub))

        nitrogen_present = sub.GetSubstructMatches(Chem.MolFromSmarts("[#7:1]:1:[#6:2]:[#6:3]:[#6:4]:[#6:5]:[#6:6]:1"))
        The_Process.append(sub)
     
    
    sub = Chem.MolFromSmiles(Chem.MolToSmiles(sub))
    nitrogen_present = sub.GetSubstructMatches(Chem.MolFromSmarts("[#7:1]:1:[#6:2]:[#6:3]:[#6:4]:[#6:5]:[#6:6]:1"))
    while(len(nitrogen_present)>0):
        remove_nitrogen = AllChem.ReactionFromSmarts("[#7:1]:1:[#6:2]:[#6:3]:[#6:4]:[#6:5]:[#6:6]:1>>[#6:2][#6:3][#6:4][#6:5]=[#6:6].[#7:1]")
        sub1 = remove_nitrogen.RunReactants((sub,))
        if(Chem.MolFromSmiles(Chem.MolToSmiles(sub1[0][0]))!=None):
            sub = sub1[0][0]
        elif(Chem.MolFromSmiles(Chem.MolToSmiles(sub1[1][0]))!=None):
            sub = sub1[1][0]
            sub = Chem.MolFromSmiles(re.sub("N","O",Chem.MolToSmiles(sub)))
        else:
            break
        sub = Chem.MolFromSmiles(Chem.MolToSmiles(sub))
        nitrogen_present = sub.GetSubstructMatches(Chem.MolFromSmarts("[#7:1]:1:[#6:2]:[#6:3]:[#6:4]:[#6:5]:[#6:6]:1"))
        The_Process.append(sub)
    
    nitrogen_present = sub.GetSubstructMatches(Chem.MolFromSmarts("[#6:1]:[#7:2]:[#6:3]"))
    nitrogen_present2 = sub.GetSubstructMatches(Chem.MolFromSmarts("[#7:1]@1@[#6:2]@[#6:3]@[#6:4]@[#6:5]@1"))
    while(len(nitrogen_present)>0 and len(nitrogen_present2)>0):
        remove_nitrogen = AllChem.ReactionFromSmarts("[#7:1]@1@[#6:2]@[#6:3]@[#6:4]@[#6:5]@1.[#8:6].[#8:7]>>[#7:1].[#8:7]-[#6:2][#6:3][#6:4][#6:5]-[#8:6]")
        sub1 = remove_nitrogen.RunReactants((sub,Water_coreactant,Water_coreactant))[0][1]
        sub1 = Chem.MolFromSmiles(Chem.MolToSmiles(sub1))
        if(sub1==None):
            remove_nitrogen = AllChem.ReactionFromSmarts("[#7:1]@1@[#6:2]@[#6:3]@[#6:4]@[#6:5]@1.[#8:6].[#8:7]>>[#7:1].[#8:7]=[#6:2]-[#6:3]=[#6:4]-[#6:5]-[#8:6]")
            sub1 = remove_nitrogen.RunReactants((sub,Water_coreactant,Water_coreactant))[0][1]
            sub1 = Chem.MolFromSmiles(Chem.MolToSmiles(sub1))
            if(sub1==None):
                break
            else:
                sub=sub1
        else:
            sub=sub1
        nitrogen_present = sub.GetSubstructMatches(Chem.MolFromSmarts("[#6:1]:[#7:2]:[#6:3]"))
        The_Process.append(sub)

    nitrogen_present = sub.GetSubstructMatches(Chem.MolFromSmarts("[#6:1]:[#7:2]"))
    nitrogen_present2 = sub.GetSubstructMatches(Chem.MolFromSmarts("[#6:1]!@[#7:2]"))
    while(len(nitrogen_present)>0 and len(nitrogen_present2)>0):
        remove_nitrogen = AllChem.ReactionFromSmarts("[#6:1]:[#7:2].[#8:7]>>[#6:1]-[#8:7].[#7:2]")
        sub = remove_nitrogen.RunReactants((sub,Water_coreactant))[0][0]
        sub = Chem.MolFromSmiles(Chem.MolToSmiles(sub))
        nitrogen_present = sub.GetSubstructMatches(Chem.MolFromSmarts("[#6:1]:[#7:2]"))
        The_Process.append(sub)

    nitrogen_present = sub.GetSubstructMatches(Chem.MolFromSmarts("[#6:1]!@[#7:2]!@[#6:3]"))
    while(len(nitrogen_present)):
        remove_nitrogen = AllChem.ReactionFromSmarts("[#6:1]!@[#7:2]!@[#6:3].[#8:7]>>[#6:1]-[#8:7].[#7:2][#6:3]")
        sub1 = remove_nitrogen.RunReactants((sub,Water_coreactant))
        if(len(re.findall("C",Chem.MolToSmiles(sub1[0][0])))>len(re.findall("C",Chem.MolToSmiles(sub1[0][1])))):
            sub = sub1[0][0]
        else: sub = sub1[0][1]
        sub = Chem.MolFromSmiles(Chem.MolToSmiles(sub))
        nitrogen_present = sub.GetSubstructMatches(Chem.MolFromSmarts("[#6:1]!@[#7:2]"))
        The_Process.append(sub)    

########################## Oxygen Testing #######################################

    oxygen_present = sub.GetSubstructMatches(Chem.MolFromSmarts("[#6:0]:[#6:1](=[#8:2])[#8:3][#6:4](=[#8:5]):[#6:8]"))
    while(len(oxygen_present)>0):
        reduce_oxygen = AllChem.ReactionFromSmarts("[#6:0]:[#6:1](=[#8:2])[#8:3][#6:4](=[#8:5]):[#6:8].[#8:6].[#8:7]>>[#6:0]-[#6:1](-[#8:2])[#8:3][#8:6][#8:7][#6:4](-[#8:5])-[#6:8]")
        reduce_oxygen2 = AllChem.ReactionFromSmarts("[#6:1][#8:2][#8:3]>>[#6:1][#8:2].[#8:3]")
        sub = reduce_oxygen.RunReactants((sub,Water_coreactant,Water_coreactant))[0][0]; sub = Chem.MolFromSmiles(Chem.MolToSmiles(sub))
        sub = reduce_oxygen2.RunReactants((sub,))[0][0]; sub = Chem.MolFromSmiles(Chem.MolToSmiles(sub))
        oxygen_present = sub.GetSubstructMatches(Chem.MolFromSmarts("[#6:0]:[#6:1](=[#8:2])[#8:3][#6:4](=[#8:5]):[#6:8]"))
        
    oxygen_present = sub.GetSubstructMatches(Chem.MolFromSmarts("[#8:0]=[#6:1][#6:2]=[#8:3]"))
    while(len(oxygen_present)>0):
        reduce_oxygen = AllChem.ReactionFromSmarts("[#8:0]=[#6:1][#6:2]=[#8:3]>>[#8:0]-[#6:1]=[#6:2]-[#8:3]")
        sub = reduce_oxygen.RunReactants((sub,))[0][0]
        sub = Chem.MolFromSmiles(Chem.MolToSmiles(sub))        
        oxygen_present = sub.GetSubstructMatches(Chem.MolFromSmarts("[#8:0]=[#6:1][#6:2]=[#8:3]"))
        The_Process.append(sub)

    oxygen_present = sub.GetSubstructMatches(Chem.MolFromSmarts("[#6:0]!=!:[#6:1]!:&=[#8:2]"))
    while(len(oxygen_present)>0):
        reduce_oxygen = AllChem.ReactionFromSmarts("[#6:0]!:[#6:1]!:&=[#8:2]>>[#6:0]!:[#6:1]-[#8:2]")
        sub1 = reduce_oxygen.RunReactants((sub,))[0][0]
        if(Chem.MolFromSmiles(Chem.MolToSmiles(sub1))!=None):
            sub = Chem.MolFromSmiles(Chem.MolToSmiles(sub1))
            oxygen_present = sub.GetSubstructMatches(Chem.MolFromSmarts("[#6:0]!:[#6:1]!:&=[#8:2]"))
        else:break
        The_Process.append(sub)
      
    oxygen_present = sub.GetSubstructMatches(Chem.MolFromSmarts("[#6:0]!:&=[#6:1]=[#8:2]"))
    while(len(oxygen_present)>0):
        reduce_oxygen = AllChem.ReactionFromSmarts("[#6:0]!:&=[#6:1]=[#8:2]>>[#6:0]=[#6:1][#8:2]")
        sub = reduce_oxygen.RunReactants((sub,))[0][0]
        sub = Chem.MolFromSmiles(Chem.MolToSmiles(sub))        
        oxygen_present = sub.GetSubstructMatches(Chem.MolFromSmarts("[#6:0]!:&=[#6:1]=[#8:2]"))
        The_Process.append(sub)
        
    furan_ring = sub.GetSubstructMatches(Chem.MolFromSmarts("[#6D2:1]:1:[#6:2]:[#6:3](-[#6:6][#6:7]):[#6D2:4]:[#8:5]:1"))
    while(len(furan_ring)>0):
        resolve_furan = AllChem.ReactionFromSmarts("[#6D2:1]:1:[#6:2]:[#6:3](-[#6:6][#6:7]):[#6D2:4]:[#8:5]:1>>[#6:1]=[#6:2][#6:3](-[#6:6][#6:7])=[#6:4].[#8:5]")
        sub = resolve_furan.RunReactants((sub,))[0][0]
        sub = Chem.MolFromSmiles(Chem.MolToSmiles(sub))
        furan_ring = sub.GetSubstructMatches(Chem.MolFromSmarts("[#6D2:1]:1:[#6:2]:[#6:3](-[#6:6][#6:7]):[#6D2:4]:[#8:5]:1"))        
        The_Process.append(sub)
        
    furan_ring = sub.GetSubstructMatches(Chem.MolFromSmarts("[#6D2:1]:1:[#6:2]:[#6:3](-[#6:6]):[#6D2:4]:[#8:5]:1"))
    while(len(furan_ring)>0):
        resolve_furan = AllChem.ReactionFromSmarts("[#6D2:1]:1:[#6:2]:[#6:3](-[#6:6]):[#6D2:4]:[#8:5]:1>>[#6D2:1]=[#6:2][#6:3](-[#6:6])=[#6D2:4].[#8:5]")
        sub = resolve_furan.RunReactants((sub,))[0][0]
        sub = Chem.MolFromSmiles(Chem.MolToSmiles(sub))
        furan_ring = sub.GetSubstructMatches(Chem.MolFromSmarts("[#6D2:1]:1:[#6:2]:[#6:3](-[#6:6]):[#6D2:4]:[#8:5]:1"))          
        The_Process.append(sub)

    furan_ring = sub.GetSubstructMatches(Chem.MolFromSmarts("[#6D3:1]:1:[#6:2]:[#6:3](-[#6:6]):[#6D2:4]:[#8:5]:1"))
    while(len(furan_ring)>0):
        resolve_furan = AllChem.ReactionFromSmarts("[#6D3:1]:1:[#6:2]:[#6:3](-[#6:6]):[#6D2:4]:[#8:5]:1.[#8:7]>>[#6:1](-[#8:7])[#6:2]-[#6:3](-[#6:6])=[#6:4][#8:5]")
        sub = resolve_furan.RunReactants((sub, Water_coreactant))[0][0]
        sub = Chem.MolFromSmiles(Chem.MolToSmiles(sub))
        furan_ring = sub.GetSubstructMatches(Chem.MolFromSmarts("[#6D3:1]:1:[#6:2]:[#6:3](-[#6:6]):[#6D2:4]:[#8:5]:1"))          
        The_Process.append(sub)

    furan_ring = sub.GetSubstructMatches(Chem.MolFromSmarts("[#6D2H1:1]:1:[#6:2]:[#6:3](-[#6:6]):[#6D3H0:4]:[#8:5]:1"))
    while(len(furan_ring)>0):
        resolve_furan_ring = AllChem.ReactionFromSmarts("[#6D2H1:1]:1:[#6:2]:[#6:3](-[#6:6]):[#6D3H0:4]:[#8:5]:1.[#8:7]>>[#8:7]-[#6D2H1:1]=[#6:2]-[#6:3](-[#6:6])=[#6D3H0:4]-[#8:5]")
        sub = resolve_furan_ring.RunReactants((sub, Water_coreactant))[0][0]
        sub = Chem.MolFromSmiles(Chem.MolToSmiles(sub))
        furan_ring = sub.GetSubstructMatches(Chem.MolFromSmarts("[#6D2H1:1]:1:[#6:2]:[#6:3](-[#6:6]):[#6D3H0:4]:[#8:5]:1"))          
        The_Process.append(sub)
    
    furan_ring = sub.GetSubstructMatches(Chem.MolFromSmarts("[#6D3H0:0](-[#6:1]):1:[#6:2]:[#6:3]:[#6D3H0:4](-[#6:5]):[#8:6]:1"))
    while(len(furan_ring)>0):
        resolve_furan_ring = AllChem.ReactionFromSmarts("[#6D3H0:0](-[#6:1]):1:[#6:2]:[#6:3]:[#6D3H0:4](-[#6:5]):[#8:6]:1.[#8:7]>>[#8:7][#6D3H0:0](-[#6:1])=[#6:2]-[#6:3]=[#6D3H0:4](-[#6:5])-[#8:6]")
        sub = resolve_furan_ring.RunReactants((sub, Water_coreactant))[0][0]
        sub = Chem.MolFromSmiles(Chem.MolToSmiles(sub))
        furan_ring = sub.GetSubstructMatches(Chem.MolFromSmarts("[#6D2H1:1]:1:[#6:2]:[#6:3](-[#6:6]):[#6D3H0:4]:[#8:5]:1"))          
        The_Process.append(sub)
    
    furan_ring = sub.GetSubstructMatches(Chem.MolFromSmarts("[#6D2H1:0]:1:[#6:2]:[#6:3]:[#6D3H0:4]:[#8:6]:1"))
    while(len(furan_ring)>0):
        resolve_furan_ring = AllChem.ReactionFromSmarts("[#6D2H1:0]:1:[#6:2]:[#6:3]:[#6D3H0:4]:[#8:6]:1.[#8:7]>>[#8:7][#6D2H1:0]=[#6:2]-[#6:3]=[#6D23H0:4]-[#8:6]")
        sub = resolve_furan_ring.RunReactants((sub, Water_coreactant))[0][0]
        sub = Chem.MolFromSmiles(Chem.MolToSmiles(sub))
        furan_ring = sub.GetSubstructMatches(Chem.MolFromSmarts("[#6D2H1:1]:1:[#6:2]:[#6:3](-[#6:6]):[#6D3H0:4]:[#8:5]:1"))          
        The_Process.append(sub)    
  
    cyclic_oxygen = sub.GetSubstructMatches(Chem.MolFromSmarts("[#6:1]@[#8:2]!:[#6:3]"))
    while(len(cyclic_oxygen)>0):
        resolve_cyclic_O  = AllChem.ReactionFromSmarts("[#6:1]@[#8:2]!:[#6:3].[#8:4].[#8:5]>>[#6:1][#8:2][#8:4][#8:5][#6:3]")
        resolve_cyclic_O_2 = AllChem.ReactionFromSmarts("[#6:1]@[#8:2]@[#8:3]>>[#6:1][#8:2].[#8:3]")
        sub = resolve_cyclic_O.RunReactants((sub,Water_coreactant,Water_coreactant))[0][0]; sub = Chem.MolFromSmiles(Chem.MolToSmiles(sub))
        sub = resolve_cyclic_O_2.RunReactants((sub,))[0][0]; sub = Chem.MolFromSmiles(Chem.MolToSmiles(sub))
        cyclic_oxygen = sub.GetSubstructMatches(Chem.MolFromSmarts("[#6:1]@[#8:2]!:[#6:3]"))
        The_Process.append(sub)
  
    sub = Chem.MolFromSmiles(Chem.MolToSmiles(sub))
    lone_electron = sub.GetSubstructMatches(Chem.MolFromSmarts("[#6D1H1:1]-[#6D2:2]!=[#6:3]"))
    while(len(lone_electron)>0):
        resolve_electron_pair = AllChem.ReactionFromSmarts("[#6D1H1:1]-[#6D2:2]!=[#6:3]>>[#6:1]=[#6:2]-[#6:3]")
        sub = resolve_electron_pair.RunReactants((sub,))[0][0]
        sub = Chem.MolFromSmiles(Chem.MolToSmiles(sub))
        lone_electron = sub.GetSubstructMatches(Chem.MolFromSmarts("[#6D1H1:1]-[#6D2:2]!=[#6:3]"))
        The_Process.append(sub)

    lone_electron = sub.GetSubstructMatches(Chem.MolFromSmarts("[#6D1H1:1]-[#6D4:2]"))
    while(len(lone_electron)>0):    
        resolve_electron_pair = AllChem.ReactionFromSmarts("[#6D1H1:1]-[#6D4:2]>>[#6H3:1]-[#6D4:2]")
        sub = resolve_electron_pair.RunReactants((sub,))[0][0]
        sub = Chem.MolFromSmiles(Chem.MolToSmiles(sub))
        lone_electron = sub.GetSubstructMatches(Chem.MolFromSmarts("[#6D1H1:1]-[#6D4:2]"))
        The_Process.append(sub)

    lone_electron = sub.GetSubstructMatches(Chem.MolFromSmarts("[#6:0]-[#6D2H1:1]-[#8:2]"))
    while(len(lone_electron)>0):    
        resolve_electron_pair = AllChem.ReactionFromSmarts("[#6:0]-[#6D2H1:1]-[#8:2]>>[#6:0]-[#6D2H2:1]-[#8:2]")
        sub = resolve_electron_pair.RunReactants((sub,))[0][0]
        sub = Chem.MolFromSmiles(Chem.MolToSmiles(sub))
        lone_electron = sub.GetSubstructMatches(Chem.MolFromSmarts("[#6:0]-[#6D2H1:1]-[#8:2]"))
        The_Process.append(sub)
        
    lone_electron = sub.GetSubstructMatches(Chem.MolFromSmarts("[#6:0]-[#6D2H1:1]-[#6:2]"))
    while(len(lone_electron)>0):    
        resolve_electron_pair = AllChem.ReactionFromSmarts("[#6:0]-[#6D2H1:1]-[#6:2]>>[#6:0]-[#6D2H2:1]-[#6:2]")
        sub = resolve_electron_pair.RunReactants((sub,))[0][0]
        sub = Chem.MolFromSmiles(Chem.MolToSmiles(sub))
        lone_electron = sub.GetSubstructMatches(Chem.MolFromSmarts("[#6:0]-[#6D2H1:1]-[#6:2]"))
        The_Process.append(sub)
    
    hydroxy2many = sub.GetSubstructMatches(Chem.MolFromSmarts("[#6:1](!@[#8:2])!@[#8:3]"))
    while(len(hydroxy2many) > 0):
        rxn3= AllChem.ReactionFromSmarts("[#6:1](!@[#8:2])!@[#8:3]>>[#6:1](-[#8:2]).[#8:3]")
        sub1 = rxn3.RunReactants((sub,))
        if(len(re.findall("C",Chem.MolToSmiles(sub1[0][0]),re.IGNORECASE))<len(re.findall("C",Chem.MolToSmiles(sub1[1][0]),re.IGNORECASE))):
            sub = sub1[1][0]
        else:
            sub = sub1[0][0]
        sub = Chem.MolFromSmiles(Chem.MolToSmiles(sub))
        hydroxy2many = sub.GetSubstructMatches(Chem.MolFromSmarts("[#6:1](!@[#8:2])!@[#8:3]"))
        The_Process.append(sub)
   
    ether_present = sub.GetSubstructMatches(Chem.MolFromSmarts("[#6:0]-[#8:1]-[#6:2]"))
    while(len(ether_present)>0):
        remove_ether=AllChem.ReactionFromSmarts("[#6:0]-[#8:1]-[#6:2]>>[#6:0]-[#8:1].[#6:2]")
        sub1 = remove_ether.RunReactants((sub,))
        if(len(re.findall("C",Chem.MolToSmiles(sub1[0][0]),re.IGNORECASE))<len(re.findall("C",Chem.MolToSmiles(sub1[1][0]),re.IGNORECASE))):
            sub = sub1[1][0]
        else:
            sub = sub1[0][0]
        ether_present = sub.GetSubstructMatches(Chem.MolFromSmarts("[#6:0]-[#8:1]-[#6:2]"))
    
    sub = Chem.MolFromSmiles(Chem.MolToSmiles(sub))
    return sub, The_Process#(CalcMolFormula(Chem.MolFromSmiles(Chem.MolToSmiles(sub))))


# %% [markdown]
## Function used to strip backbones exclusively to carbon-carbon linkages (Skeletons)

# %% [code]
break_O = AllChem.ReactionFromSmarts("[#6:1]@[#8:2]>>[#6:1].[#8:2]")
#for i in flattened_backbone:
def backbone_flattener(i):
    i = re.sub("c","C",i)
    i = re.sub("o","O",i)
    i = re.sub("H\d+", "",i)
    i = re.sub("H", "",i )
    i = re.sub("\(\)","", i)
    i = re.sub("\[13C","[C",i)
    i = Chem.MolFromSmiles(str(i))
    oxygen_ring = i.GetSubstructMatches(Chem.MolFromSmarts("[#6:1]@[#8:2]@[#6:3]"))
    while(len(oxygen_ring)>0):
        i = break_O.RunReactants((i,))[0][0]
        i = Chem.MolFromSmiles(Chem.MolToSmiles(i))
        oxygen_ring = i.GetSubstructMatches(Chem.MolFromSmarts("[#6:1]@[#8:2]@[#6:3]"))
    i = Chem.MolToSmiles(i)
    i = re.sub("\+","",i)
    i = re.sub("\=","", i)
    i = re.sub("\#","",i)
    i = re.sub("\(O\)","",i)
    i = re.sub("O","",i)
    i = re.sub("\\\\","", i)
    i = re.sub("/", "",i)
    i = re.sub("CO\)","C)", i)
    i = re.sub("\@","",i)
    i = re.sub("H", "",i )
    i = re.sub("\[", "",i )
    i = re.sub("\]", "", i)
    i = re.sub("P","",i)
    i = re.sub("\(\)","", i)
    i = re.sub("\-","",i)
    i = Chem.MolToSmiles(Chem.MolFromSmiles(i))
    return(i)


# %% [markdown]
## Example of deconstruction

# %% [code]
taxol = Chem.MolFromSmiles("CC1=C2[C@H](C(=O)[C@@]3([C@H](C[C@@H]4[C@](C3[C@@H]([C@@](C2(C)C)(C[C@@H]1OC(=O)[C@@H]([C@H](C5=CC=CC=C5)NC(=O)C6=CC=CC=C6)O)O)OC(=O)C7=CC=CC=C7)(CO4)OC(=O)C)O)C)OC(=O)C")
formula, deconstruction = deconstruct2backbone(taxol)

#formula = final backbone with decorated oxygens
#deconstruction = the appended list of each action along the way for troubleshooting

print(CalcMolFormula(formula))
formula = Chem.MolFromSmiles(Chem.MolToSmiles(formula))
# produce image of deconstructed taxol
print(Chem.MolToSmiles(formula, allBondsExplicit=True))
formula

# %% [markdown]
# Dictionary of Natural Products

# %% [markdown]
## Import DNP dataset(s) and deconstruct

# %% [code]
# Updated DNP (V30.2) library
deconstruction_mol = pd.read_csv("Supp.1a.DNP_Diterpene_Mining_v30.2.csv", sep=",", encoding='utf-8')
deconstruction_mol = deconstruction_mol.dropna(subset= ["SMILES"],axis=0).reset_index(drop=True)

print("Shape of DNP Database before formatting:", np.shape(deconstruction_mol))
counter = 0
empty = []
#format some DNP Smiles for interpretation by RDKit
for i in deconstruction_mol["SMILES"]:
    counter = counter+1
    i = re.findall("\S+", i)[0]
    i = re.split("\.",i)[0]
    i = re.sub("C=\[N\]\(C\)c","C=[N+](C)c",i)
    i = re.sub("\(\[#6\]\)\(\[N\]#\[C-\]\)","([#6])([N+]#[C-])",i)
    empty.append(i)
deconstruction_mol["SMILES"] = empty

counter = 0

#convert SMILES to canonical form when possible
for i in deconstruction_mol["SMILES"]:
    if(str(Chem.CanonSmiles(i))!=str(i)):
        deconstruction_mol["SMILES"].iloc[counter]=str(Chem.CanonSmiles(i))        
    counter+=1
    
#some entries are missing "Molecular Formula" and "Molecular Weight", this calculates that
all_mol_formula = []
all_mol_weight = []
for i in deconstruction_mol["SMILES"]:
    all_mol_formula.append(CalcMolFormula(Chem.MolFromSmiles(i)))
    all_mol_weight.append(Descriptors.MolWt(Chem.MolFromSmiles(i)))
deconstruction_mol["Molecular_Formula"]=all_mol_formula
deconstruction_mol["Molecular_Weight"]=all_mol_weight

print("Shape of DNP Database after formatting:", np.shape(deconstruction_mol))
print("^^^ These should be the same... ^^^")
deconstruction_mol.head()

# %% [code]
#IF YOU WOULD LIKE TO SKIP TO A PROBLEM AREA
start=0
deconstruction_mol = deconstruction_mol.iloc[start:,]
index = start

#MAKE LIST OF COMPOUNDS THAT DO NOT FIT INTO A 20C BACKBONE CRITERIA
work2do = 1
work_list1 = []
length_comp = 0
carbon_number = []

#EMPTY LIST OF BACKBONES, SKELETONS, AND CARBON_NUMBER 
flattened_backbone = []
backbone = []

#LOOP THROUGH ALL SMILES IN DNP DATABASE AND CONVERT TO BACKBONE AND SKELETON
for i in deconstruction_mol["SMILES"]:
    diterpene = Chem.MolFromSmiles(i)
    formula, deconstruction = deconstruct2backbone(diterpene)
    Chemical_formula = CalcMolFormula(Chem.MolFromSmiles(Chem.MolToSmiles(formula)))
    match = re.search('C20', Chemical_formula)
    # Check if not C20, could be real, coulbe be an error
    if (match):
        if(len(deconstruction)>length_comp):
            length_comp = len(deconstruction)
            length_index = index
    else:
        #print(work2do,"\t",index, "\t", formula, "\t", Chemical_formula)
        work_list1.append(index)
        work2do = work2do+1
    # create list of Backbones (backbone), Skeletons (flattened_backbone), and the final number of carbons (carbon_number)
    backbone.append(Chem.MolToSmiles(formula))
    flattened_backbone.append(backbone_flattener(Chem.MolToSmiles(formula)))
    carbon_number.append(re.search("\d+", CalcMolFormula(formula))[0])

    index=index+1

# %% [code]
# GENERATE SUPPLEMENTAL DATA 2A
# Add columns to Original DNP Dataset to include Backbone, Skeleton, and Carbon number
deconstruction_mol["Backbone"]=backbone
deconstruction_mol["Skeleton"]=flattened_backbone
deconstruction_mol["Carbon Number"]=carbon_number
deconstruction_mol.to_csv("Supp.2a.DNP_Diterpene_Mining_v30.2_Updated_Skeleton_Backbone.tsv", sep='\t', encoding='utf-8',index=False)
deconstruction_mol.head()

# %% [markdown]
## Summarize diterpene skeleton data for later characterization

# %% [code]
# GENERATE DNP DERIVED SKELETON DATA FOR SUPPLEMENTAL DATA 2E
DNP_Terpene_dataset_v2 = pd.read_csv("Supp.2a.DNP_Diterpene_Mining_v30.2_Updated_Skeleton_Backbone.tsv",sep="\t")

sample = pd.DataFrame()
sample["frequency"] = np.unique(DNP_Terpene_dataset_v2["Skeleton"], return_counts=True)[1]
sample["SMILES"] = np.unique(DNP_Terpene_dataset_v2["Skeleton"], return_counts=True)[0]

test = []
for i in sample["SMILES"]:
    x = len(re.findall("C", i, re.IGNORECASE))
    test.append(x)

sample["Carbon Number"] = test

test = []
for i in np.unique(DNP_Terpene_dataset_v2["Skeleton"]):
    x = 0
    index = []
    for j in DNP_Terpene_dataset_v2["Skeleton"]:
        if i == j:
            index.append(x)  
        x = x+1
    test.append(index)
sample["index"] = test
sample = sample.sort_values(by = ['frequency'], ascending=False )
sample = sample.reset_index(drop=True)
print(sample.index)
print("Total Skeleton:", np.shape(sample))

# FILTERING STEPS
#These values used for pruning cases skeleton has more or less than 20C
#sample = sample.sort_values(by=['Carbon Number'])
# sample = sample.loc[sample['Carbon Number']>15]
# sample = sample.loc[sample['Carbon Number']<25]

# Keep skeletons with 20C structure, those that do not are likely mislabeled or not generated by terpene synthase activity
sample = sample.loc[sample['Carbon Number']==20]
sample = sample.reset_index(drop=True)
print("20C Skeleton:", np.shape(sample))
sample.head(10)
#sample = sample.loc[sample["frequency"]>1]

# %% [code]
sample.to_csv("Supp.2e.DNP_Diterpene_Skeleton_Summary.tsv", sep='\t', index=False)

# %% [markdown]
# TeroKit

# %% [markdown]
## Import TeroKit dataset(s) and deconstruct
### Diterpene, Triterpene, Sesquiterpene

# %% [code]
# CHOOSE WHICH DATASET INTENDED TO ASSESS
TeroKit_mol = pd.read_csv("Supp.1b.TeroKit_Diterpene_v2.0.tsv", sep="\t")
#TeroKit_mol = pd.read_csv("Supp.1c.TeroKit_Triterpene_v2.0.tsv", sep="\t")
#TeroKit_mol = pd.read_csv("Supp.1d.TeroKit_Sesquiterpene_v2.0.tsv", sep="\t")

TeroKit_mol = TeroKit_mol.rename(columns={'smiles': 'SMILES'})
TeroKit_mol.head()

# %% [code]
# ensure correct formatting for deconstruction (i.e. formatting matches RDKit) 
index = 0
print("Shape of TeroKit Database before formatting:", np.shape(TeroKit_mol))
for i in TeroKit_mol['mol_id']:
    if(isinstance(i, float)):
        TeroKit_mol['mol_id'].iloc[index]="Unknown"
        TeroKit_mol['category'].iloc[index]=TeroKit_mol['SMILES'].iloc[index]
        TeroKit_mol['SMILES'].iloc[index]=TeroKit_mol['inchi'].iloc[index]
        TeroKit_mol['inchi'].iloc[index]=str(TeroKit_mol['formula'].iloc[index])
        TeroKit_mol['formula'].iloc[index]=CalcMolFormula(Chem.MolFromSmiles(TeroKit_mol['SMILES'].iloc[index]))
        #print(TeroKit_mol.iloc[index])
    index+=1
counter = 0
for i in TeroKit_mol["SMILES"]:
    if(str(Chem.CanonSmiles(i))!=str(i)):
        TeroKit_mol["SMILES"].iloc[counter]=str(Chem.CanonSmiles(i))
    counter+=1
print("Shape of TeroKit Database after formatting:", np.shape(TeroKit_mol))
print("^^^ These should be the same... ^^^")

# %% [code]
#IF YOU WOULD LIKE TO SKIP TO A PROBLEM AREA
start=0
TeroKit_mol = TeroKit_mol.iloc[start:,]
index = start

#MAKE LIST OF COMPOUNDS THAT DO NOT FIT INTO A 20C BACKBONE CRITERIA
work2do = 1
work_list1 = []

#EMPTY LIST OF BACKBONES, SKELETONS, AND CARBON_NUMBER
flattened_backbone = []
backbone = []
carbon_number = []

#LOOP THROUGH ALL SMILES IN DNP DATABASE AND CONVERT TO BACKBONE AND SKELETON
for i in TeroKit_mol["SMILES"]:
    diterpene = Chem.MolFromSmiles(i)
    formula, deconstruction = deconstruct2backbone(diterpene)
    Chemical_formula = CalcMolFormula(Chem.MolFromSmiles(Chem.MolToSmiles(formula)))
    match = re.search('C20', Chemical_formula)
    # Check if not C20, could be real, coulbe be an error
    if(match):
        if(len(deconstruction)>length_comp):
            length_comp = len(deconstruction)
            length_index = index
    else:
        #print(work2do,"\t",index, "\t", formula, "\t", Chemical_formula)
        work_list1.append(index)
        work2do = work2do+1
    flattened_backbone.append(backbone_flattener(Chem.MolToSmiles(formula)))
    backbone.append(Chem.MolToSmiles(formula))
    carbon_number.append(re.search("\d+", CalcMolFormula(formula))[0])
    
    # Print Progress
    if(index%1000==0):
        print(index, "finished thus far; \t", format(100*(index/len(list(TeroKit_mol["SMILES"]))), ".2f"), "% complete")

    index=index+1

# %% [code]
# GENERATE TEROKIT DERIVED SUPPLEMENTAL DATA 2B, 2C, or 2D

# Add columns to Original DNP Dataset to include Backbone, Skeleton, and Carbon number
TeroKit_mol["Backbone"]=backbone
TeroKit_mol["Skeleton"]=flattened_backbone
TeroKit_mol["Carbon Number"]=carbon_number

'''NOTE: MAKE SURE TO SPECIFIY WHICH DATASET OUTPUT YOU WOULD LIKE TO REFERENCE HERE'''
TeroKit_mol.to_csv("Supp.2b.TeroKit_Diterpene_v2.0_Updated_Skeleton_Backbone.tsv", sep='\t', index=False)
#TeroKit_mol.to_csv("Supp.2c.TeroKit_Triterpene_v2.0_Updated_Skeleton_Backbone.tsv", sep='\t', index=False)
#TeroKit_mol.to_csv("Supp.2d.TeroKit_Sesquiterpene_v2.0_Updated_Skeleton_Backbone.tsv", sep='\t', index=False)

TeroKit_mol.head()

# %% [markdown]
## Summarize diterpene skeleton data for later characterization

# %% [code]
# GENERATE TEROKIT DERIVED SUPPLEMENTAL DATA 2F
Tero_Terpene_dataset_v2 = pd.read_csv("Supp.2b.TeroKit_Diterpene_v2.0_Updated_Skeleton_Backbone.tsv",sep="\t")

sample = pd.DataFrame()
sample["frequency"] = np.unique(Tero_Terpene_dataset_v2["Skeleton"], return_counts=True)[1]
sample["SMILES"] = np.unique(Tero_Terpene_dataset_v2["Skeleton"], return_counts=True)[0]

test = []
for i in sample["SMILES"]:
    x = len(re.findall("C", i, re.IGNORECASE))
    test.append(x)

sample["Carbon Number"] = test

test = []
for i in np.unique(Tero_Terpene_dataset_v2["Skeleton"]):
    x = 0
    index = []
    for j in Tero_Terpene_dataset_v2["Skeleton"]:
        if i == j:
            index.append(x)  
        x = x+1
    test.append(index)
sample["index"] = test
sample = sample.sort_values(by = ['frequency'], ascending=False )
sample = sample.reset_index(drop=True)
print(sample.index)
print("Total Skeleton:", np.shape(sample))

# FILTERING STEPS
#These values used for pruning cases skeleton has more or less than 20C
#sample = sample.sort_values(by=['Carbon Number'])
# sample = sample.loc[sample['Carbon Number']>15]
# sample = sample.loc[sample['Carbon Number']<25]

# Keep skeletons with 20C structure, those that do not are likely mislabeled or not generated by terpene synthase activity
sample = sample.loc[sample['Carbon Number']==20]
sample = sample.reset_index(drop=True)
print("20C Skeleton:", np.shape(sample))
sample.head(10)
#sample = sample.loc[sample["frequency"]>1]

# %% [code]
sample.to_csv("Supp.2f.Terokit_Diterpene_Skeleton_Summary.tsv", sep='\t', index=False)

# %% [markdown]
## Graphical representation of diterpene skeleton abundance distribution

# %% [code]
# Graphic Representation of Backbone abundance vs distribution
TeroKit_diterpene_summary = pd.read_csv("Supp.2f.Terokit_Diterpene_Skeleton_Summary.tsv", sep='\t')
plt.figure(figsize=(13,4))
plt.bar(TeroKit_diterpene_summary.index, TeroKit_diterpene_summary["frequency"], align="center", color="black", width=1)

plt.vlines(7,0,4000, linewidth=1.5, color="blue"); plt.text(12,3800,"50% (top 7 skeletons)", color = "blue")
plt.vlines(34,0,3500, linewidth=1.5, color="red"); plt.text(39,3300,"75% (top 25 skeletons)", color = "red")
plt.vlines(83,0,3000, linewidth=1.5, color="green"); plt.text(69,2800,"<  90% (top 83 skeletons)", color = "green")
plt.vlines(169,0,2500, linewidth=1.5, color="indigo"); plt.text(155,2300,"<  95% (top 169 skeletons)", color = "indigo")
plt.vlines(529,0,2000, linewidth=1.5, color="deeppink"); plt.text(516,1800,"<  99% (top 529 skeletons)", color = "deeppink")

plt.xlim(-2,872)
plt.xlabel("Unique 20C Diterpene Skeletons ordered by abundance")
plt.ylabel("Diterpene Skeleton Abundance")
plt.title("Diterpene Skeleton Dataset Abundance and Distribution")
plt.savefig("Supp.2g.Terokit_Diterpene_Skeleton_Abundance_Distribution.png",dpi=600,bbox_inches='tight')
plt.show()

# %% [markdown]
## Graphical representation of diterpene deconstruction based on Carbon Number

# %% [code]
# Graphic Representation of Compound abundance vs carbon number
TeroKit_diterpene_updated = pd.read_csv("Supp.2b.TeroKit_Diterpene_v2.0_Updated_Skeleton_Backbone.tsv", sep="\t")
Carbon_deconstruction = list(TeroKit_diterpene_updated["Carbon Number"])
plt.hist(Carbon_deconstruction, bins=np.unique(Carbon_deconstruction), color="grey", align="left")
plt.gcf().set_size_inches(10, 3)
plt.ylim(0,40000)
plt.text(19.4,35500,"20C", color = "black", fontsize=8)
plt.text(18.3,3200,"19C", color = "black", fontsize=8)
plt.text(17.3,950,"18C", color = "black", fontsize=8)
plt.text(20.5,550,"21C", color = "black", fontsize=8)
plt.text(14.3,500,"15C", color = "black", fontsize=8)
plt.text(39.4,500,"40C", color = "black", fontsize=8)

plt.xlabel("Carbon Number")
plt.ylabel("Compound Abundance")
plt.title("Deconstruction Effectiveness Based on Final Carbon Count")
plt.tight_layout()
plt.savefig("Supp.2h.Terokit_Diterpene_Deconstructed_Skeleton_Carbon_Distribution.png",dpi=600,bbox_inches='tight')
plt.show()

# %% [code]
for i in range(len(np.unique(TeroKit_diterpene_updated["Carbon Number"]))):
    print(np.unique(TeroKit_diterpene_updated["Carbon Number"], return_counts=True)[0][i],np.unique(TeroKit_diterpene_updated["Carbon Number"], return_counts=True)[1][i])

# %% [code]
#Zoom into outliers and non-20-Carbon compounds
plt.hist(Carbon_deconstruction, bins=np.unique(Carbon_deconstruction), color="grey", align="left")
plt.gcf().set_size_inches(10, 3)
plt.ylim(0,4000)
plt.text(18.1,3200,"19C", color = "black", fontsize=8)
plt.text(17.1,950,"18C", color = "black", fontsize=8)
plt.text(20.5,500,"21C", color = "black", fontsize=8)


plt.text(14.3,500,"15C", color = "black", fontsize=8)
plt.text(14.5,700,"↓", color = "black", fontsize=8)
plt.text(10,900,"sesquiterpenes", color = "black", fontsize=8)

plt.text(39.4,500,"40C", color = "black", fontsize=8)
plt.text(39.7,700,"↓", color = "black", fontsize=8)
plt.text(35,900,"diterpene-diterpene dimers", color = "black", fontsize=8)

plt.text(9.7,200,"↓", color = "black", fontsize=8)
plt.text(5,400,"monoterpenes", color = "black", fontsize=8)

plt.text(20.5,3500,"20C (~35,000)", color = "black", fontsize=8)
plt.text(20.6,3750,"↑", color = "black", fontsize=8)


plt.xlabel("Carbon Number")
plt.ylabel("Compound Abundance")
plt.title("Zoomed Deconstruction Effectiveness Based on Final Carbon Count")
plt.tight_layout()
plt.savefig("Supp.2i.10X_Zoomed_Terokit_Diterpene_Deconstructed_Skeleton_Carbon_Distribution.png",dpi=600,bbox_inches='tight')


plt.show()

# %% [code]
skeleton1 = pd.read_csv("Supp.2e.DNP_Diterpene_Skeleton_Summary.tsv", sep="\t")
skeleton2 = pd.read_csv("Supp.2f.Terokit_Diterpene_Skeleton_Summary.tsv", sep="\t")

# %% [code]
print("DNP",len(np.unique(list(skeleton1["SMILES"]))))
print("Tero",len(np.unique(list(skeleton2["SMILES"]))))
print("Both",len(np.unique(list(skeleton1["SMILES"])+list(skeleton2["SMILES"]))))

# %% [code]
print("DNP conversion", sum(list(skeleton1['frequency']))/len(DNP_Terpene_dataset_v2["Skeleton"]))
print("Tero conversion", sum(list(skeleton2["frequency"]))/len(Tero_Terpene_dataset_v2["Skeleton"]))

# %% [code]
DNP_Terpene_dataset_v2 = pd.read_csv("Supp.2a.DNP_Diterpene_Mining_v30.2_Updated_Skeleton_Backbone.tsv",sep="\t")
Tero_Terpene_dataset_v2 = pd.read_csv("Supp.2b.TeroKit_Diterpene_v2.0_Updated_Skeleton_Backbone.tsv",sep="\t")

# %% [code]
print("Tero:\t",len(np.unique(Tero_Terpene_dataset_v2["Skeleton"])))

# %% [code]
print("DNP:\t",len(np.unique(DNP_Terpene_dataset_v2["Skeleton"])))
```
