## Supplementary material for "Rule-Based Deconstruction and Reconstruction of Diterpene Libraries: Categorizing Patterns & Unravelling the Structural Landscape": Zip of all Code used: Supp_Code.2.Skeleton_PCA.ipynb

```
# %% [code]
import pandas as pd
import seaborn as sns
import numpy as np
import re
import matplotlib.pyplot as plt

import rdkit
from rdkit.Chem import AllChem as Chem
from rdkit import DataStructs

import sklearn
from sklearn.decomposition import PCA
from sklearn.preprocessing import StandardScaler

# %% [code]
print("Pandas:",pd.__version__)
print("Seaborn:",sns.__version__)
print("Numpy:",np.__version__)
print("Sklearn:", sklearn.__version__)
print("RE:", re.__version__)
print("Matplotlib:",plt.matplotlib.__version__)
print("RDKit:", rdkit.__version__)

# %% [code]
temp = pd.read_csv("Supp.3.PCA_comparison_matrix.tsv", sep="\t")
temp.columns[temp.isna().any()]

# %% [code]
#Supplemental Code 2 Skeleton Structure PCA generation

### DATASET(S) INTENDED AS INPUT FOR THIS CODE
# Supp.2e.DNP_Diterpene_Skeleton_Summary.tsv
# Supp.2f.Terokit_Diterpene_Skeleton_Summary.tsv

### SUPPLEMENTAL DATASETS GENERATED BY THIS FUNCTION (GENERATION OF BACKBONE AND SKELETON INFORMATION)
# Supp.3.PCA_comparison_matrix.tsv

'''
### OVERVIEW OF FUNCTIONALITIY ###
### Davis Mathieu
### Jan 15, 2024

This program takes the skeleton summary data generated in Supplemental
Code 1 and compares all structures to one another by converting to bit
vector and making all possible comparisons. This generates a matrix 
that can then be used to map compounds based on similarity. We assume
that compounds that are similar will rank closely to one another and 
also compare more similar to other compounds as well.

This generates figure 1 and the Supplemental structure PCA
'''
X=()

# %% [code]
# IMPORT RELEVANT PACKAGES FOR COMPARISON TO BE MADE
skeleton =pd.read_csv("Supp.2e.DNP_Diterpene_Skeleton_Summary.tsv", sep="\t")
# skeleton = pd.read_csv("Supp.2f.Terokit_Diterpene_Skeleton_Summary.tsv", sep="\t")
skeleton = skeleton[::-1]
skeleton.tail(20)

# %% [code]
#empty list for appending each comparison
fplist = []
visual = []
#investigate Skeleton SMILEs
for mol1 in skeleton["SMILES"]:
    fp = Chem.GetMorganFingerprintAsBitVect(Chem.MolFromSmiles(mol1), 2)
    # and compare to every other skeleton SMILEs
    for mol2 in skeleton["SMILES"]:
        fp2= Chem.GetMorganFingerprintAsBitVect(Chem.MolFromSmiles(mol2), 2)
        fplist.append(DataStructs.DiceSimilarity(fp,fp2))
    
    visual.append(fplist);fplist = []

# %% [code]
# GENERATE COMPARISON MATRIX AND CONVERT TO HEATMAP; SAVE MATRIX AS SUPPLEMENTAL 3
skeleton_PCA_matrix = pd.DataFrame(visual)

color_spec = sns.light_palette("red", as_cmap=True)
skeleton_heatmap = skeleton_PCA_matrix.style.background_gradient(cmap=color_spec)
#skeleton_heatmap

# %% [code]
# Save comparison matrix of PCA for Supplemental 
skeleton_PCA_matrix.to_csv("Supp.3.PCA_comparison_matrix.tsv", sep='\t', index=False)

# %% [code]
# PROVIDE COLOR AND SIZE SETTINGS FOR PCA
#determine point size based on number of compounds represented
point_size =[]
for i in skeleton["frequency"]:
    point_size.append(i*1.5)

# Hexcode for various shades of the rainbow to distinguish #of rings
rainbow_gradient = [
    '#D71F1F',   # Dark Red ------ 0 rings
    '#FFA500',   # Orange -------- 1 ring
    '#FFFF00',   # Yellow -------- 2 rings
    '#00FF00',   # Green --------- 3 rings
    '#000080',   # Navy Blue ----- 4 rings
    '#800080',   # Purple -------- 5 rings
    '#9400D3'    # Dark Purple --- 6 rings
]

color_map = []
max_num = 0
for i in skeleton['SMILES']:
    x = re.findall("\d", i, re.IGNORECASE)
    target = int(len(x)/2)
    color_map.append(rainbow_gradient[target])

# %% [code]
# PERFORM PCA ANALYSIS
scaler = StandardScaler()
scaler.fit(skeleton_PCA_matrix)
scaled_data = scaler.transform(skeleton_PCA_matrix)
pca = PCA(n_components=2)
pca.fit(scaled_data)
pca_variance = pca.explained_variance_ratio_

coord_pca = pca.transform(scaled_data)
x_coord_pca = coord_pca[:,0]
y_coord_pca = coord_pca[:,1]
# DETERMINE PC1 AND PC2 THEN GRAPH PCA COORDINATES
plt.figure(figsize=(12,8))
plt.scatter(x_coord_pca,-y_coord_pca, c =color_map,cmap=color_map,alpha=0.6, s=point_size)

x_axis_title = 'Principal Component 1:   '+str(int(round(pca_variance[0]*100)))+"%"
y_axis_title = 'Principal Component 2:   '+str(int(round(pca_variance[1]*100)))+"%"
plt.xlabel(x_axis_title, fontsize=16)
plt.ylabel(y_axis_title, fontsize=16)
plt.xticks(fontsize=14)
plt.yticks(fontsize=14)

plt.savefig("Structure_similarity.png", dpi='figure', bbox_inches="tight")
plt.show

# %% [code]
count = 0
total_linear = 0
for i in skeleton["SMILES"]:
    if(re.search("\d+",i)==None):
        print(i, skeleton["frequency"].iloc[count])
        total_linear= total_linear+skeleton["frequency"].iloc[count]
    count+=1
print(total_linear/sum(skeleton['frequency']))

# %% [code]
count=0
print("SMILES FOR TOP 20 MOST ABUNDANT SKELETONS W/ COORDINATES FOR MANUAL ANNOTATION")
print("\nSMILES\t\t\t\t\t x_coord\t y_coord",)
for i in skeleton["SMILES"].tail(20):
    if(len(str(i))<=30):
        print(i,"\t\t",round(x_coord_pca[count-20],5),"\t",round(-y_coord_pca[count-20],3))
    else:
        print(i,"\t",round(x_coord_pca[count-20],5),"\t",round(-y_coord_pca[count-20],3))
    count+=1

# %% [code]
```
