## Supplementary material for "Rule-Based Deconstruction and Reconstruction of Diterpene Libraries: Categorizing Patterns & Unravelling the Structural Landscape": Zip of all Code used: Supp_Code.3a.Pickaxe_DM_NS.py

```
# import relevant packages
from __future__ import annotations
import datetime
import multiprocessing
import pickle
import time
import numpy as np
import os
import matplotlib.pyplot as plt
import pandas as pd  # type: ignore
import rdkit  # type: ignore
import mordred  # type: ignore
import sklearn  # type: ignore
import libsbml  # type: ignore
import lxml  # type: ignore
import sys

####################
 # WHAT TO CHANGE FOR EACH RUN #
#rule_list = "./minedatabase/data/metacyc_rules/test.tsv"
#rule_list = "./minedatabase/data/metacyc_rules/Custom_Rules_Only.tsv"
#rule_list = "./inputs/WorkingXenonRules/Xenon_protonation.tsv"
#rule_list = "./inputs/WorkingXenonRules/20231207_AllRules.tsv"
#rule_list = "./inputs/WorkingXenonRules/20240101_AllRules.tsv"
rule_list = "./inputs/WorkingXenonRules/20240101_All_NoProtonation.tsv"
#rule_list = "./inputs/WorkingXenonRules/20240101_NoMacrosOrCleavage.tsv"
#rule_list = "./inputs/WorkingXenonRules/20231207_AllRules.tsv"
#rule_list = "./inputs/WorkingXenonRules/20231207_AllRules_NoDiphosphateCleave.tsv"

#input_cpds = "./inputs/GGPP_only_precursor.csv"
#input_cpds = "./inputs/GGPP_only_precursor.csv"
input_cpds = "./inputs/10GenNoMacroCompoundsGGPP.csv"
#input_cpds = "./inputs/8GenNoMacroCompoundsGGPP.csv"
#products_cpds = "./inputs/All_20_C_backbone.tsv"
products_cpds = "./inputs/identified_target_hits.tsv"

generations = 10
similarity_threshold = [0,0,0,0,0,0,0,0,0,0,0,0,0,0,0]

increasing_similarity = True
load_targets_without_filter = True
react_targets = True
prune_to_targets = True		#want for final True; False for testing
filter_after_final_gen = True 	#want True; False for testing

sample_size = 100000000

cutoff_fingerprint_args = {"radius": 4}
sample_fingerprint_args = {"radius": 4}
rule_abundance_print= True

write_to_csv = True
output_dir = "outputs/20240101_10GenNoMacro_10GenAll_PruneToIdentifiedTargets"
####################

# minedatabase specific imports
from minedatabase.filters import (
    AtomicCompositionFilter,
    MCSFilter,
    MetabolomicsFilter,
    MWFilter,
    SimilarityFilter,
    SimilaritySamplingFilter,
)
from minedatabase.pickaxe import Pickaxe
from minedatabase.rules import metacyc_generalized, metacyc_intermediate

####################

start = time.time()

####################

### DATABASE INFORMATION OPTIONS ###

write_db = False
# Database name and message to print in metadata
database = "example_db"
message = "Example run to show how pickaxe is run."
# Whether to write compounds to core compound database with extra info
write_core = True
# Force overwrite existing database
database_overwrite = True
# Use local DB, i.e. localhost:27017
use_local = True

####################
# RULES TO USE FOR PICKAXE

#rule_list = "./minedatabase/data/metacyc_rules/metacyc_intermediate_rules.tsv"
#rule_list = "./minedatabase/data/metacyc_rules/metacyc_generalized_rules.tsv"

coreactant_list = "./minedatabase/data/metacyc_rules/metacyc_coreactants.tsv"

####################

### HOW MANY ROUNDS OF TRANSFORMATION TO OCCUR ###

processes = 24                # Number of processes for parallelization
verbose = True               # Display RDKit warnings and errors

####################

### ADVANCED OPTIONS ###
inchikey_blocks_for_cid = 1  # Number of inchi key blocks to gen cid
explicit_h = False          # use explicit hydrogens in rules
kekulize = False              # Assign Double bonds?
neutralise = False            # Neutralise all molecules when loading
quiet = False                # Suppress the output??
indexing = False             

####################

### IF YOU WANT TRANSFORMATIONS TO APPROACH A CERTAIN MOLECULAR TYPE###
''' Will want to create this'''
#target_example = "./example_data/target_list_many.csv"

# Target molecular weight end compound (probably don't want)
MW_filter = False
min_MW = 100
max_MW = 150

# Target final molecule based on composition (probably don't want)
atomic_composition_filter = False
atomic_composition_constraints = {
    "C": [4, 7],
    "O": [5, 5]
}

########################

thermo_filter = None

#########################

# Filter based on approaching a final compound based on similarity to structure

feasibility_filter = False

similarity_filter = True               # Should be true?
cutoff_fingerprint_method = "Morgan"

# Arguments to pass to fingerprint_method
cutoff_fingerprint_args = {"radius": 4}
cutoff_similarity_method = "Tanimoto"

similarity_sample = True
# Number of compounds per generation to sample

#########################

sample_fingerprint_method = "RDKit"
# arguments to pass to fingerprint_method
sample_fingerprint_args = {"radius": 4}
sample_similarity_method = "Tanimoto"

#########################

def weight(score):
    """weight is a function that accepts a similarity score as the sole argument
    and returns a scaled value. 
    """
    return score**4

weight_representation = "score^4"

#########################

def rule_abundance():
        df = pd.read_csv(output_dir + "/reactions.tsv", sep="\t", index_col=0).drop(columns=["Name"])
        Rule_freq = np.unique(df['Reaction rules'], return_counts=True)
        for i in range(len(Rule_freq[0])):
            print(Rule_freq[0][i],"---", Rule_freq[1][i])

#########################

mcs_filter = False
metabolomics_filter = False

#########################

print_parameters = True

#########################

def print_run_parameters():
    """Write relevant parameters."""
    def print_parameter_list(plist):
        for i in plist:
            print(f"--{i}: {eval(i)}")
    print("\n-------------Run Parameters-------------")
    print("\nRun Info")
    print("rules = " + rule_list)
    print("Output folder is output_dir " + output_dir)
    print_parameter_list(["coreactant_list", "input_cpds"])
    #print_parameter_list(["coreactant_list", "rule_name", "input_cpds"])
    print("\nExpansion Options")
    print_parameter_list(["generations", "processes"])
    print("\nGeneral Filter Options")
    print_parameter_list(
        [
            "filter_after_final_gen",
            "react_targets",
            "prune_to_targets",
        ]
    )
    if similarity_sample:
        print("\nTanimoto Sampling Filter Options")
        print_parameter_list(
            [
                "sample_size",
                "weight_representation",
                "sample_fingerprint_args",
                "sample_fingerprint_method",
                "sample_similarity_method"
            ]
        )
    if similarity_filter:
        print("\nTanimoto Threshold Filter Options")
        print_parameter_list(
            [
                "similarity_threshold",
                "increasing_similarity",
                "cutoff_fingerprint_args",
                "cutoff_fingerprint_method",
                "cutoff_similarity_method"
            ]
        )
    if mcs_filter:
        print("\nMaximum Common Substructure Filter Options")
        print_parameter_list(["crit_mcs"])
    if metabolomics_filter:
        print("\nMetabolomics Filter Options")
        print_parameter_list(["met_data_path", "met_data_name",
                              "possible_adducts", "mass_tolerance"])
    if MW_filter:
        print("\nMolecular Weight Filter Options")
        print_parameter_list(["min_MW", "max_MW"])
    if atomic_composition_filter:
        print("\nAtomic Composition Filter")
        print_parameter_list(["atomic_composition_constraints"])
    print("\nPickaxe Options")
    print_parameter_list(
        [
            "verbose",
            "explicit_h",
            "kekulize",
            "neutralise",
            "quiet",
            "indexing"
        ]
    )
    print("----------------------------------------\n")

    ###############################################################################

#####################

# Use "spawn" for multiprocessing
# multiprocessing.set_start_method("spawn")

mongo_uri = False
# Change database to none if not writing
if write_db is False:
    database = None

### Initialize the Pickaxe class
# print parameters
if print_parameters:
    print_run_parameters()
pk = Pickaxe(
    coreactant_list=coreactant_list,
    rule_list=rule_list,
    errors=verbose,
    explicit_h=explicit_h,
    kekulize=kekulize,
    neutralise=neutralise,
    image_dir=None,
    inchikey_blocks_for_cid=inchikey_blocks_for_cid,
    database=database,
    database_overwrite=database_overwrite,
    mongo_uri=mongo_uri,
    quiet=quiet,
    react_targets=react_targets,
    filter_after_final_gen=filter_after_final_gen
)

# Load compounds
pk.load_compound_set(compound_file=input_cpds)

# Load target compounds for filters
if (
    similarity_filter or mcs_filter or similarity_sample
    or load_targets_without_filter or MW_filter
    or atomic_composition_filter or thermo_filter
    or feasibility_filter):
    #pass
    pk.load_targets(products_cpds)

# Apply filters in this order
if MW_filter:
    pk.filters.append(MWFilter(min_MW, max_MW))
if atomic_composition_filter:
    pk.filters.append(AtomicCompositionFilter(atomic_composition_constraints))
if similarity_filter:
    taniFilter = SimilarityFilter(
        crit_similarity=similarity_threshold,
        increasing_similarity=increasing_similarity,
        fingerprint_method=cutoff_fingerprint_args,
        fingerprint_args=cutoff_fingerprint_args,
        similarity_method=cutoff_similarity_method
    )
    pk.filters.append(taniFilter)
if similarity_sample:
    taniSampleFilter = SimilaritySamplingFilter(
        sample_size=sample_size,
        weight=weight,
        fingerprint_method=sample_fingerprint_method,
        fingerprint_args=sample_fingerprint_args,
        similarity_method=sample_similarity_method)
    pk.filters.append(taniSampleFilter)
if mcs_filter:
    mcsFilter = MCSFilter(crit_mcs=crit_mcs)
    pk.filters.append(mcsFilter)
if metabolomics_filter:
    if rt_predictor_pickle_path:
        with open(rt_predictor_pickle_path, "rb") as infile:
            rt_predictor = pickle.load(infile)
    else:
        rt_predictor = None
    metFilter = MetabolomicsFilter(
        filter_name="ADP1_Metabolomics_Data",
        met_data_name=met_data_name,
        met_data_path=met_data_path,
        possible_adducts=possible_adducts,
        mass_tolerance=mass_tolerance,
        rt_predictor=rt_predictor,
        rt_threshold=rt_threshold,
        rt_important_features=rt_important_features
    )
    pk.filters.append(metFilter)
if feasibility_filter:
    pk.filers.append(feasibility_filter)
if thermo_filter:
    pk.filters.append(thermo_filter)

# Transform compounds (the main step)
pk.transform_all(processes, generations)
if pk.targets and prune_to_targets:
    pk.prune_network_to_targets()

if write_to_csv:
    pk.assign_ids()
    pk.write_compound_output_file(output_dir + "/compounds.tsv")
    pk.write_reaction_output_file(output_dir + "/reactions.tsv")
    print("Succesfully wrote output to\n")
    print(os.getcwd())
print("----------------------------------------")
print(f"Overall run took {round(time.time() - start, 2)} seconds.")
print("----------------------------------------")

if rule_abundance_print:
	rule_abundance()

#####################
```
