## Supplementary material for "Rule-Based Deconstruction and Reconstruction of Diterpene Libraries: Categorizing Patterns & Unravelling the Structural Landscape": Zip of all Code used: Supp_Code.3b.Match.py

```
from rdkit import Chem
from rdkit.Chem.Draw import IPythonConsole
from rdkit.Chem import Draw
from rdkit.Chem import AllChem
from rdkit import DataStructs
import re
import os
import numpy as np
import pandas as pd
import sys

from rdkit import RDLogger                                                                                                                                                               
RDLogger.DisableLog('rdApp.*')

##################################### PARAMETERS ####################################

# Load Files 

DIR = sys.argv[1]

#path = os.getcwd() + "/outputs/20231207_4GenNoMacro_4All/"
path = os.getcwd() + "/outputs/" + DIR 
output_path =  path + "/compounds.tsv"
print(output_path)
#output_path = os.getcwd() + "/20231002_15GensAllRules/compounds_abreviated.tsv"
target_path = os.getcwd() + "/All_20_C_backbone.tsv"

print("file matching: " + output_path)
# Do you want to strip "=" and "#" from compounds?
flatten=True

#####################################################################################
#include "./"
MatchOutputFile = path + "/MatchedCompounds.tsv"


# loop through Pickaxe compound list
output_path = pd.read_csv(output_path, sep='\t')
target_path = pd.read_csv(target_path, sep='\t')
temp = []
for i in target_path["SMILES"]:
	temp.append(Chem.MolFromSmiles(i))
target_path["Mol"]= temp

hits = {}
print("Beginning Target Search")
for i in range(np.shape(output_path)[0]):
	if(i%500==0):
		print(i,"/",np.shape(output_path)[0])
	if(flatten==True):
		x = re.sub("=","",output_path["SMILES"].iloc[i])
		potential_match = re.sub("#", "", x)
		x = Chem.MolFromSmiles(re.sub("#","",x))
	else:
		x = Chem.MolFromSmiles(output_path["SMILES"].iloc[i])

	# compare compound to target list
	if(re.search("Xe", output_path["SMILES"].iloc[i],re.IGNORECASE)):
		null=False
	elif(re.search("P", output_path["SMILES"].iloc[i],re.IGNORECASE)):
		null=False

	elif(re.search("N", output_path["SMILES"].iloc[i],re.IGNORECASE)):
		null=False
	else: null=True	

	if(null==True):
		if not (target_path[target_path["SMILES"]==potential_match].empty):
			j = target_path[target_path["SMILES"]==potential_match]["id"].iloc[0]
			cont = False
			if (j in hits):
				hits[j].append(output_path["ID"].iloc[i])
			else:
				hits[j] = [output_path["ID"].iloc[i]]
			#print(i, target_path[target_path["SMILES"]==potential_match]["id"].iloc[0])
		else: cont=True

	if(null==True and cont==True):
		for j in range(np.shape(target_path)[0]):
			Comparison_Score = DataStructs.FingerprintSimilarity(Chem.RDKFingerprint(x),Chem.RDKFingerprint(target_path["Mol"].iloc[j]), metric=DataStructs.TanimotoSimilarity)
			if(Comparison_Score==1):
				if(j in hits):
					hits[j].append(output_path["ID"].iloc[i])
				else:
					hits[j] = [output_path["ID"].iloc[i]]
				#print(Comparison_Score,output_path["SMILES"].iloc[i], j)
				break
print("Completed list of tentative hits, pending QC check:")
for key in sorted(hits, key=lambda x: int(x)):
    print(key)
#Checking match success
print("\n\nBeginning Quality Check")
for key in hits:
	for j in hits[key]:
		j = str(output_path[output_path["ID"]==j]["SMILES"].iloc[0])
		if(flatten==True):
			j = re.sub("=","",j)
			j = re.sub("#","",j)
		for k in hits[key]:
			k = str(output_path[output_path["ID"]==k]["SMILES"].iloc[0])
			if(flatten==True):
				k = re.sub("=","",k)
				k = re.sub("#","",k)
			Comparison_Score = DataStructs.FingerprintSimilarity(Chem.RDKFingerprint(Chem.MolFromSmiles(j)),Chem.RDKFingerprint(Chem.MolFromSmiles(k)), metric=DataStructs.TanimotoSimilarity)
			if(Comparison_Score!=1):
				print("ERROR FOR THIS TARGET:", key)

Final = pd.DataFrame(columns=['Target','Frequency','Compounds'])
#print("\nTarget\tFrequency\tCompounds")
for key in sorted(hits, key=lambda x: int(x)):
	#print(key,"\t", np.shape(hits[key])[0], "\t",hits[key])
	all_compounds = [hits[key]]
	Final = pd.concat([Final,pd.DataFrame({'Target':key, 'Frequency':np.shape(hits[key])[0], 'Compounds':all_compounds})], ignore_index=True)
Final.to_csv(MatchOutputFile, sep='\t', index=False)
```
