## Supplementary material for "Rule-Based Deconstruction and Reconstruction of Diterpene Libraries: Categorizing Patterns & Unravelling the Structural Landscape": Zip of all Code used: Supp_Code.3c.Network_Maker.py

```
# %%
from __future__ import annotations

import itertools as it
import matplotlib.pyplot as plt
import networkx as nx
import pandas as pd
from matplotlib.lines import Line2D
from pathlib import Path
from shared.parser import lex, parse
from shared.utils import split_stoichiometries
import csv

def get_colors_from_cmap(
    cmap_name: str, reaction_rules: list[str]
) -> dict[str, tuple[float, float, float, float]]:
    from matplotlib import colors

    cmap = plt.get_cmap(cmap_name, len(reaction_rules))  # type: ignore
    if isinstance(cmap, colors.ListedColormap):
        vals = cmap.colors  # type: ignore
    elif isinstance(cmap, colors.LinearSegmentedColormap):
        vals = [cmap(i) for i in range(len(reaction_rules))]
    else:
        raise ValueError("Unknown type of colormap")

    return dict(zip(reaction_rules, vals))


# %%
df = pd.read_csv(
    Path(__file__).parent / "." / "PrunedPickaxes/Phase1_Phase2_AllReactions.tsv" ,  # type: ignore
    sep="\t",
    index_col=0,
)
reactions = df["ID equation"].apply(lambda x: parse(lex(x))).to_dict()
compounds = set(j for i in reactions.values() for j in i)
OutFileName = "PrunedPhase1Phase2_Edges.tsv"

#print(df)

to_ignore = {
    "PYROPHOSPHATE_ACCEPTOR_CoF",
    "PYROPHOSPHATE_DONOR_CoF",
    "Test",
    "WATER",
    "XENON_1",
    "CARBON",
    "XENON_ProtonateAcceptor",
    "SecondaryXenonDonor",
    "SecondaryXenonAcceptor",
    "TertiaryXenonDonor",
    "TertiaryXenonAcceptor",
    "BadXENON_CleavageDonor",
    "BadXENON_CleavageDonor",
    "BadXENON_CleavageAcceptor2",
    "BadXENON_CleavageAcceptor2",
    "XENON_CleavageDonor3",
    "XENON_CleavageAcceptor3",
    "XenonCarbon",
    "Carbon",
    "Ethane",
    "Ethene",
    "PrimaryToTertiaryDonor",
    "PrimaryToTertiaryAcceptor",
    "PrimaryToTertiaryDonor2",
    "PrimaryToTertiaryAcceptor2",
    "PrimaryToTertiaryDubAcceptor2",
    "PrimaryToTertiaryDubAcceptor",
    "PrimaryToTertiaryDubDonator",
    "CCO",
    "CleavageResonanceDonor",
    "CleavageResonanceAcceptor",
    "H2",
    "CARBON",
    "XENON_1",
    "CH3",
    "XeH3",
    "XENON_2",
    "XENON",
    "CARBOCATION_SUBSTITUTE_XE",
    "XENONProtonator"
}

reactions = {k1: {k2: v2 for k2, v2 in v1.items() if k2 not in to_ignore} for k1, v1 in reactions.items()}
# df.loc["pkr0000001", "Reaction rules"]

reaction_rules = sorted(df.loc[:, "Reaction rules"].unique())
color_per_rule = get_colors_from_cmap("tab20", reaction_rules)
g = nx.Graph()


with open(OutFileName, 'w', newline='') as csvfile:
     output = csv.writer(csvfile, delimiter='\t',quotechar='|')
     for rxn, stoich in reactions.items():
          subs, prods = split_stoichiometries(stoich)
          for u, v in it.product(subs, prods):
              RULE = df.loc[rxn, "Reaction rules"]	
              #print(prods)
              output.writerow([u,RULE,v])	
		
              #print(outdf)	
        #print(u, '\t', RULE, v)	
        #print(df.loc[rxn, "Reaction rules"])	
        #print(v)	
        #print(subs)	
        #print(prods)	
        #print(df.loc[rxn, "Reaction rules"])	
        #g.add_edge(u, v, color=color_per_rule[df.loc[rxn, "Reaction rules"]])  # type: ignore

#print("Graph is {}a tree".format("" if nx.is_tree(g) else "not "))
#print(g)
#fig, ax = plt.subplots(figsize=(12, 12))
#
#nx.draw(
#    g,
#    with_labels=True,
#    edge_color=nx.get_edge_attributes(g, "color").values(),
#    width=3,
#    ax=ax,
#    pos=nx.spring_layout(g, seed=3),
#)
#ax.legend(
#    [Line2D([0], [0], color=v) for v in color_per_rule.values()],  # type: ignore
#    color_per_rule.keys(),
#    title="Reaction rule",
#)
#plt.show()
#plt.savefig('reaction_fig.jpg', dpi='figure')

# %%
```
