## Supplementary material for "Rule-Based Deconstruction and Reconstruction of Diterpene Libraries: Categorizing Patterns & Unravelling the Structural Landscape": Zip of all Code used: Supp_Code.4.Quench_Predictor.ipynb

```
# %% [code]
# IMPORT ALL RELEVENT MODULES
import pandas as pd
import numpy as np
import os
import re
import matplotlib.pyplot as plt
import colorsys

import rdkit
from rdkit import Chem
from rdkit.Chem import AllChem
from rdkit import DataStructs
from rdkit.Chem.rdMolDescriptors import CalcMolFormula
from rdkit.Chem.Draw import IPythonConsole
from rdkit.Chem import Draw
Water_coreactant = AllChem.MolFromSmiles('O')

# %% [code]
print("Pandas:",pd.__version__)
print("Numpy:",np.__version__)
print("RE:", re.__version__)
print("Matplotlib:",plt.matplotlib.__version__)
print("RDKit:", rdkit.__version__)
print("OS:",os.__file__)

# %% [markdown]
## IMPORT RELEVANT DATASETS

# %% [code]
#Supplemental Code 4: CARBOCATION PREDICTION SOFTWARE AND BACKBONE DIVERSITY PROFILING

### DATASET(S) INTENDED AS INPUT FOR THIS CODE
# PAIRED FOR DNP
# Supp.2a.DNP_Diterpene_Mining_v30.2_Updated_Skeleton_Backbone.tsv
# Supp.2e.DNP_Diterpene_Skeleton_Summary.tsv

# PAIRED FOR TEROKIT
# Supp.2b.TeroKit_Diterpene_v2.0_Updated_Skeleton_Backbone.tsv
# Supp.2f.Terokit_Diterpene_Skeleton_Summary.tsv 

### DATASET OUTPUT(S) (20 IN TOTAL)
# Figure 3

'''
### OVERVIEW OF FUNCTIONALITIY ###
### Davis Mathieu
### Jan 17, 2024

This program takes all the backbones for a specific skeleton class
(performed on the top 20 most common diterpene skeletons in TeroKit 
here) and evaluates the resolution of the carbocation position based
on tertiary carbons with double bonds. Normal quenching should render
a molecule with 32H, 34H, or 36H depending on if anything was quenched
with water, or two waters. Any compound with Hydrogen content greater 
than 36H indicates post cyclization saturation of double bonds. Those
with less than 32H indicate additional desaturation to form additional 
double bonds. Aromatics were also identified within the groups. 

Functions
- color_range_list              Returns a list of hexcodes equal to the number
                                of conditions wished and equally spaced between
                                two prechosen hexcodes (generally chosen to be
                                color blind friendly)

- estimate_carbocation          Predict a carbocation at all tertiary carbon
                                locations containing a double bond. If there is
                                a double bond connection two tertiary carbons
                                consider both possible locations for position
                                and check later

- reduce_oxygen                 Based on Hydrogen count, we can predict how many 
                                Hydroxyl groups were used to quench carbocations
                                or saturate double bonds. This produces a list of 
                                all possible combinations of hydroxyl side chains
                                that would be predicted as necessary for normal 
                                carbocation quenching (for example a compound with
                                5 hydroxyl side chains and 34H, suggests one quenching
                                event occured to resolve a carbocation; 5 structures
                                would be returned for further analysis)

- estimate_carbocations_from_O  Works the same as "estimat_carbocation" but instead 
                                guages where carbocations may have been due to the 
                                placement of water where if the water was removed 
                                would produce a double bond on a tertiary carbon. The
                                acception to this being also investigating the secondary
                                carbon neighboring the position of diphosphate in GGDP; 
                                This condition is only relevant for labdanes. 

#-----------------------------------------------------------------#
This program is intended to produce two pie charts. The first pie chart 
shows quenching patterns of carbocations, post cyclization (de)saturation
events, and presence of aromatic compounds. The second pie chart shows the 
number of estimated positions of unique and semi-unambigious predictions of 
carbocation rearrangement/resolution via the formation of a double bond or 
quenching with water'''
x=0

# %% [code]
# IMPORT PAIRED DATASETS FOR RUNNING MSA ALIGNMENT
Backbones = pd.read_csv("Supp.2b.TeroKit_Diterpene_v2.0_Updated_Skeleton_Backbone.tsv", sep="\t")
Skeletons = pd.read_csv("Supp.2f.Terokit_Diterpene_Skeleton_Summary.tsv", sep="\t")

#Choose skeleton index and create an empty list to iterate thorugh
backbone = []
atom_value = 19

#Determine in the backbone is a product of Class II/Class I (Class_I_only==False) synthesis or Class I (Class_I_only==True)
#synthesis. This is important to predict how many quenching events are predicted to take place.
Class_I_only=True


for i in eval(Skeletons["index"].iloc[atom_value]):
    backbone.append(Backbones["Backbone"].iloc[i])

# %% [code]
# to visualize structure for creating Figure 3
print(Skeletons["SMILES"].iloc[atom_value])
Chem.MolFromSmiles(Skeletons["SMILES"].iloc[atom_value])

# %% [code]
# Create a SMARTs rule for removing all hydroxyl groups to check for <32H, 32H, 34H, 36H, 36H<, and aromatics
#x=0
remove_O = AllChem.ReactionFromSmarts("[#6:1]-[#8:2]>>[#6:1].[#8:2]")
Quenching_pattern = []
Quenching_pattern.append(999999)
Compounds2quench = []
for j in backbone:
    original_compound = j
    i =Chem.MolFromSmiles(j)
    while(i.GetSubstructMatches(Chem.MolFromSmarts("[#6:1]-[#8:2]"))):
        i = remove_O.RunReactants((i,))[0][0]
        i = Chem.MolFromSmiles(Chem.MolToSmiles(i))
    if(i.GetSubstructMatches(Chem.MolFromSmarts("[#6:0]:[#6:1]"))):
        Quenching_pattern.append("aromatic")
    else:
        Quenching_pattern.append(re.findall("\d+",CalcMolFormula(i))[1])
        if((int(re.findall("\d+",CalcMolFormula(i))[1])%2)==0):
            Compounds2quench.append(original_compound)
            #break
#        if((int(re.findall("\d+",CalcMolFormula(i))[1]))==32):
#            print(original_compound)
#             if(x<3):
#                 x+=1
#             else: break

# %% [code]
# correct for instances where a compound class has multiple orders of saturation (like phytanes)
count= -1
for i in Quenching_pattern:
    count+=1
    try:
        int_value = int(i)
    except ValueError: continue
    if(int(i)>36):
        Quenching_pattern[count] = 999999
    

# %% [code]
### FUNCTION FOR CREATING EVENLY SPACED COLORS
def color_range_list (start_hex, stop_hex, list_len):
    #You can create a Python list of colors ranging from "#3bd2f7" to "#1b5766" using a loop to generate the intermediate colors. Here's a code snippet to do that:
    start_color = start_hex
    end_color = stop_hex
    num_colors = list_len

    # Convert the hex colors to RGB tuples
    start_rgb = tuple(int(start_color[i:i+2], 16) for i in (1, 3, 5))
    end_rgb = tuple(int(end_color[i:i+2], 16) for i in (1, 3, 5))

    # Generate a list of colors
    colors = []
    if(int(num_colors)==1): 
        colors.append(start_hex)
    elif(int(num_colors)==0):
        colors =[]
    else:
        for i in range(num_colors):
            # Interpolate between the start and end colors
            r = int(start_rgb[0] + (end_rgb[0] - start_rgb[0]) * i / (num_colors - 1))
            g = int(start_rgb[1] + (end_rgb[1] - start_rgb[1]) * i / (num_colors - 1))
            b = int(start_rgb[2] + (end_rgb[2] - start_rgb[2]) * i / (num_colors - 1))

            # Convert RGB to hex format and append to the list
            color_hex = "#{:02x}{:02x}{:02x}".format(r, g, b)
            colors.append(color_hex)

    return(colors)

# %% [code]
count=0
non_quenched_O_36=0
non_quenched_O_34=0
check =[]

for i in backbone:
    #predict if a count above C20H32 is caused by quenching or not
    if(Quenching_pattern[count]==str(36)):
        tertiary_O = Chem.MolFromSmiles(backbone[count]).GetSubstructMatches(Chem.MolFromSmarts("[#6D4:1]-[#8:2]"))
        labdane_acception = Chem.MolFromSmiles(backbone[count]).GetSubstructMatches(Chem.MolFromSmarts("[#6D3:0]~[#6D2:1]~[#6D2:2]-[#8:3]"))
        if(Class_I_only==True):
            non_quenched_O_36+=1
            count+=1
            continue
        if((len(tertiary_O)+len(labdane_acception))>=2):
            count+=1
            continue
        if(len(tertiary_O)<2):
            non_quenched_O_36+=1   
            #check.append(count)
    elif(Quenching_pattern[count]==str(34)):
        tertiary_O = Chem.MolFromSmiles(backbone[count]).GetSubstructMatches(Chem.MolFromSmarts("[#6D4:1]-[#8:2]"))
        labdane_acception = Chem.MolFromSmiles(backbone[count]).GetSubstructMatches(Chem.MolFromSmarts("[#6D3:0]~[#6D2:1]~[#6D2:2]-[#8:3]"))
        if((len(tertiary_O)+len(labdane_acception))>=1):
            count+=1
            continue
        if(len(tertiary_O)<1):
            non_quenched_O_34+=1   
            check.append(count)
    count+=1

# %% [code]
print(non_quenched_O_34)
#print(CalcMolFormula(Chem.MolFromSmiles(backbone[count])))
count = 0
for i in check:
    if(count==1):
        print(i)
        break
    count+=1
#tertiary_O = Chem.MolFromSmiles(backbone[i]).GetSubstructMatches(Chem.MolFromSmarts("[#6D4:1]-[#8:2]"))
#labdane_acception = Chem.MolFromSmiles(backbone[i]).GetSubstructMatches(Chem.MolFromSmarts("[#6D3:0]~[#6D2:1]~[#6D2:2]-[#8:3]"))
# print(CalcMolFormula(Chem.MolFromSmiles(backbone[i])))
# Chem.MolFromSmiles(backbone[i])

# %% [code]
#Chem.MolFromSmiles(backbone[count])
Quenching_freq = np.unique(Quenching_pattern, return_counts=True)

# %% [code]
# Format for visualizing backbone patterns for a particular skeleton group
Quenching_freq = np.unique(Quenching_pattern, return_counts=True)
# Initialize 99999 as 0, just incase there are additional predicted saturation
if(Quenching_freq[0][len(Quenching_freq[0])-1]=="aromatic"):
    Quenching_freq[1][len(Quenching_freq[0])-2]=Quenching_freq[1][len(Quenching_freq[0])-2]-1
else:
    Quenching_freq[1][len(Quenching_freq[0])-1]=Quenching_freq[1][len(Quenching_freq[0])-1]-1
index = 0
print(Quenching_freq)
# create index for each compound class you want to look at (color with shades of pinks, greens, and gray)
Quenching_freq_32_34_36 = []
aromatic = []; aromatic_color = []
value_32 = []; color_32 = []
value_34 = []; color_34 = []
value_36 = []; color_36 = []
temp=[]
# get an idea for (de)saturation events
less_than = 0
more_than = 0
for i in Quenching_freq[0]:
    try:
        int_value = int(i)
    except ValueError:
        aromatic = [index]; aromatic_color = ["#B7B7B7"]
        continue
    if(int(i)==36):
        value_36 = [index]; color_36 = ["#a16abc"]
        Quenching_freq[1][index]= Quenching_freq[1][index]-non_quenched_O_36
    elif(int(i)==34):
        value_34 = [index]; color_34 = ["#d94f9b"]
        Quenching_freq[1][index]= Quenching_freq[1][index]-non_quenched_O_34
    elif(int(i)==32):
        value_32 = [index]; color_32 = ["#ff3e85"]
        #print(index)
    elif(int(i)<32):
        Quenching_freq_32_34_36.append(index)
        if(less_than==0):
            remember=index
        else:
            Quenching_freq[1][remember]=Quenching_freq[1][remember]+Quenching_freq[1][index]
            Quenching_freq[1][index]=0
        less_than +=1
    elif(int(i)>36):
        Quenching_freq_32_34_36.append(index)
        if(more_than==0):
            Quenching_freq[1][index]=Quenching_freq[1][index]+non_quenched_O_36+non_quenched_O_34
        more_than +=1
    index+=1
    
# Get list of colors needed
Quenching_freq_32_34_36 =value_32+value_34+value_36+Quenching_freq_32_34_36+aromatic
colors = color_32+color_34+color_36+color_range_list("#A4F799","#A4F799", less_than)+color_range_list("#236700","#236700", more_than)+ aromatic_color
                                     

# %% [code]
# Create even spaces for Pie Chart 1 and graph; Total number of compounds in that skeleton class is plotted on top of Pie
explosion = []; space=0.025
for i in range(len(Quenching_freq[0])):
    #space = space+0.005
    explosion.append(space)
print(explosion)

plt.figure(figsize=(8,8))
plt.text(-0.125*len(str(sum(Quenching_freq[1]))),-.1,str(sum(Quenching_freq[1])), size=70, color="Black")
plt.pie(Quenching_freq[1][Quenching_freq_32_34_36],startangle=90,colors=colors, explode=explosion)

# %% [code]
Xenon_Carbocation = AllChem.MolFromSmiles('[Xe]')
Tertiary_carbocation1 = AllChem.ReactionFromSmarts("[#6,#54:0]-[#6D3:1](-[#6:2])=[#6D2:3].[Xe:4]>>[#6,#54:0]-[XeH0:4](-[#6:2])-[#6D2:3].[#6:1]")
Tertiary_carbocation2 = AllChem.ReactionFromSmarts("[#6,#54:0]-[#6D3:1](-[#6:2])=[#6D1:3].[Xe:4]>>[#6,#54:0]-[XeH0:4](-[#6:2])-[#6D1:3].[#6:1]")
Tert_Tert_Neighbors = AllChem.ReactionFromSmarts("[#6:0]-[#6D3:1](-[#6:2])=[#6D3:3].[Xe:4]>>[#6:0]-[XeH0:4](-[#6:2])-[#6D3:3].[#6:1]")
def estimate_carbocation(compound):
    compound = [compound]; temp = []
    # create carbocation for all occurances of C3=C2
    while(compound[0].GetSubstructMatches(Chem.MolFromSmarts("[#6D3:0]=[#6D2:1]"))):
        compound[0] = Tertiary_carbocation1.RunReactants((compound[0],Xenon_Carbocation))[0][0]
        compound[0] = Chem.MolFromSmiles(Chem.MolToSmiles(compound[0]))
    # create carbocation for all occurances of C3=C1
    while(compound[0].GetSubstructMatches(Chem.MolFromSmarts("[#6,#54:0]-[#6D3:1](-[#6:2])=[#6D1:3]"))):
        compound[0] = Tertiary_carbocation2.RunReactants((compound[0],Xenon_Carbocation))[0][0]
        compound[0] = Chem.MolFromSmiles(Chem.MolToSmiles(compound[0]))
    # create all carbocation formations for C3=C3
    if(compound[0].GetSubstructMatches(Chem.MolFromSmarts("[#6D3:0]=[#6D3:1]"))):
        intermediate = Tert_Tert_Neighbors.RunReactants((compound[0],Xenon_Carbocation))
        for i in intermediate:
            temp.append(Chem.MolToSmiles(i[0]))
        compound = list(np.unique(temp))
    # Convert structure to a list if there is no C3=C3
    elif(len(compound)==1): compound[0] =Chem.MolToSmiles(compound[0])
        
    #FLATTEN COMPOUND(S) BEFORE RETURNING
    index = 0
    for i in compound:
        i = re.sub(r"\[C\@\@H\]","C",i)
        i = re.sub(r"\[C\@H\]","C",i)
        i = re.sub(r"\[C\@\@\]","C",i)
        i = re.sub(r"\[C\@\]","C",i)
        i = re.sub(r"\/","",i)
        compound[index] = re.sub(r"/","",i)
        compound[index] = Chem.CanonSmiles(compound[index]); index+=1
    # return tentative carbocation estimated structures
    return(compound)

# %% [code]
# Remove n number of oxygens based on conditions provided and provide all possible combinations that this can be done
remove_O = AllChem.ReactionFromSmarts("[#6:1][#8:2]>>[#6:1].[#8:2]")
def reduce_oxygen(compound, Remaining_Oxygen):
    iterations = len(compound.GetSubstructMatches(Chem.MolFromSmarts("[#6:0][#8:1]")))-Remaining_Oxygen
    final_compounds = [Chem.MolToSmiles(compound)]
    for i in range(iterations):
        new_compounds = []
        for j in final_compounds:
            j = Chem.MolFromSmiles(j)
            temp = remove_O.RunReactants((j,))
            for k in temp:
                new_compounds.append(Chem.MolToSmiles(Chem.MolFromSmiles(Chem.MolToSmiles(k[0]))))
        final_compounds = list(np.unique(new_compounds))
        if(iterations-i==1):
            break
    return final_compounds

# %% [code]
# convert cases of oxygen connected to tertiary carbons to carbocations (similar to "estimate_carbocations")
Quench_quad_O = AllChem.ReactionFromSmarts("[#8D1:0]-[#6D4:1](-[#6:2])(-[#6:3])-[#6:4].[Xe:5]>>[XeD3H0:5](-[#6:2])(-[#6:3])-[#6:4].[#8D1:0]-[#6D1:1]")
Quench_Tert_Neighbor_O = AllChem.ReactionFromSmarts("[#8D1:0]-[#6D2:2]-[#6D3:1](-[#6:3])-[#6:4].[Xe:5]>>[#6D1:2]-[XeD3H0:5](-[#6:3])-[#6:4].[#6:1][#8D1:0]")
Quench_Phosphate_Neighbor_O = AllChem.ReactionFromSmarts("[#8D1:0][#6D2:1][#6D2:2].[Xe:3]>>[#6D1:1][XeD2:3].[#6:2][#8:0]")
Xenon_Carbocation = AllChem.MolFromSmiles('[Xe]')
def estimate_carbocations_from_O(compound):
    compound = Chem.MolFromSmiles(compound)
    while(compound.GetSubstructMatches(Chem.MolFromSmarts("[#8D1:0]-[#6D4:1](-[#6:2])(-[#6:3])-[#6:4]"))):
        compound = Quench_quad_O.RunReactants((compound,Xenon_Carbocation))[0][0]
        compound = Chem.MolFromSmiles(Chem.MolToSmiles(compound))
    while(compound.GetSubstructMatches(Chem.MolFromSmarts("[#8D1:0]-[#6D2:2]-[#6D3:1](-[#6:3])-[#6:4]"))):
        compound = Quench_Tert_Neighbor_O.RunReactants((compound,Xenon_Carbocation))[0][0]
        compound = Chem.MolFromSmiles(Chem.MolToSmiles(compound))
    while(compound.GetSubstructMatches(Chem.MolFromSmarts("[#8D1:0][#6D2:1][#6D2:2]"))):
        compound = Quench_Phosphate_Neighbor_O.RunReactants((compound,Xenon_Carbocation))[0][0]
        compound = Chem.MolFromSmiles(Chem.MolToSmiles(compound))
    return compound

# %% [code]
# Run all possible combinations; structures with only one possible form are accumulated as unique and then those with
# multiple structures are checked against previous hits, prioritizing compounds with a higher frequency of abundance
print(len(Compounds2quench))
remove_O = AllChem.ReactionFromSmarts("[#6:1]-[#8:2]>>[#6:1].[#8:2]")
Predicted_Carbocations = []
Before_Times = 0
for i in Compounds2quench:
    if(Before_Times%50==0):print(Before_Times)
    original_compound = Chem.MolFromSmiles(i) 
    i = Chem.MolFromSmiles(i)
    TEMP_Oxygen = []; FINAL_Oxygen = []
    while(i.GetSubstructMatches(Chem.MolFromSmarts("[#6:1]-[#8:2]"))):
            i = remove_O.RunReactants((i,))[0][0]
            i = Chem.MolFromSmiles(Chem.MolToSmiles(i))
    
    if(int(re.findall("\d+",CalcMolFormula(i))[1])==32):
        x = estimate_carbocation(i)
        index=0
        for i in x:
            x[index] = Chem.CanonSmiles(x[index])
            index+=1
        Predicted_Carbocations.append(x)
        
    # CHECK FOR ALL POSSIBLE CARBOCATION PATTERNS FOR H34,H36,H38
    elif(int(re.findall("\d+",CalcMolFormula(i))[1])>32 and int(re.findall("\d+",CalcMolFormula(i))[1])%2==0):
        Oxygen2leave = int((int(re.findall("\d+",CalcMolFormula(i))[1])-32)/2)
        tentatitive_hits = reduce_oxygen(original_compound,Oxygen2leave)
        for i in tentatitive_hits:
            x = estimate_carbocations_from_O(i)
            if(re.search("O",Chem.MolToSmiles(x))==None):
                TEMP_Oxygen.append(Chem.MolToSmiles(x))
            TEMP_Oxygen = list(np.unique(TEMP_Oxygen))
        for i in TEMP_Oxygen:
            FINAL_Oxygen = FINAL_Oxygen + estimate_carbocation(Chem.MolFromSmiles(i))
        Predicted_Carbocations.append(FINAL_Oxygen)

### ODDITIES (LESS THAN)
    # CHECK FOR ALL POSSIBLE CARBOCATION PATTERNS FOR H30
    elif(int(re.findall("\d+",CalcMolFormula(i))[1])==30):
        tentatitive_hits = reduce_oxygen(original_compound,0)
        for i in tentatitive_hits:
            x = estimate_carbocations_from_O(i)
            if(re.search("O",Chem.MolToSmiles(x))==None):
                TEMP_Oxygen.append(Chem.MolToSmiles(x))
            TEMP_Oxygen = list(np.unique(TEMP_Oxygen))
        for i in TEMP_Oxygen:
            FINAL_Oxygen = FINAL_Oxygen + estimate_carbocation(Chem.MolFromSmiles(i))
        Predicted_Carbocations.append(FINAL_Oxygen)
    Before_Times+=1

# %% [code]
# Identifies how many structures can be resolved to a carbocation resolved structure (aromatics are ignored)
successfully_IDd = 0
Consensus_Quench = []
for i in Predicted_Carbocations:
    if(len(i)==1):
        Consensus_Quench =Consensus_Quench+i 
        successfully_IDd+=1
unique_elements, counts = np.unique(Consensus_Quench, return_counts=True)

# Get indices that would sort in descending order
sorted_indices = np.argsort(-counts)
sorted_unique_elements = unique_elements[sorted_indices]
sorted_counts = counts[sorted_indices]
sorted_result = (sorted_unique_elements, sorted_counts)[0]
for i in Predicted_Carbocations:
    if(len(i)>1):
        for j in sorted_result:
            if j in i:
                Consensus_Quench.append(j)
                successfully_IDd+=1
                break
print(successfully_IDd)

# %% [code]
# Orders carbocation groups based on the number of copmounds that are represented by a particular carbocation formation
unique_elements, counts = np.unique(Consensus_Quench,return_counts=True)
sorted_indices = np.argsort(-counts)
sorted_unique_elements = unique_elements[sorted_indices]
sorted_counts = counts[sorted_indices]
sorted_result = (sorted_unique_elements, sorted_counts)

# %% [code]
#You can create a Python list of colors ranging from "#3bd2f7" to "#1b5766" using a loop to generate the intermediate colors. Here's a code snippet to do that:
start_color = "#FF3E85"
end_color = "#319FFF"
num_colors = len(list(sorted_result)[0])

# Convert the hex colors to RGB tuples
start_rgb = tuple(int(start_color[i:i+2], 16) for i in (1, 3, 5))
end_rgb = tuple(int(end_color[i:i+2], 16) for i in (1, 3, 5))

# Generate a list of colors
colors = []
for i in range(num_colors):
    # Interpolate between the start and end colors
    r = int(start_rgb[0] + (end_rgb[0] - start_rgb[0]) * i / (num_colors - 1))
    g = int(start_rgb[1] + (end_rgb[1] - start_rgb[1]) * i / (num_colors - 1))
    b = int(start_rgb[2] + (end_rgb[2] - start_rgb[2]) * i / (num_colors - 1))
    
    # Convert RGB to hex format and append to the list
    color_hex = "#{:02x}{:02x}{:02x}".format(r, g, b)
    colors.append(color_hex)

if(successfully_IDd-len(backbone)!=0):
    #colors.append("#FDFDFD")
    colors.append("#B7B7B7")
color_list = colors
print(len(colors),"\t",colors)

# %% [code]
# Fractions of pie chart t ograph
Unique_quench_configs = list(sorted_result[1].astype(int))+[len(backbone)-successfully_IDd]
print(Unique_quench_configs)
explosion = []; space=0.01
for i in range(len(Unique_quench_configs)):
    space = space+0.005
    explosion.append(space)


# %% [code]
plt.figure(figsize=(8,8))
#plt.text(-0.125*len(str(successfully_IDd)),-.1,"[C+]", size=70, color="Black")
plt.pie(list(Unique_quench_configs),startangle=90,colors=colors,explode=explosion)

# %% [code]
## print(Skeletons["SMILES"].iloc[atom_value])
Chem.MolFromSmiles(Skeletons["SMILES"].iloc[atom_value])

# %% [code]


# %% [code]
```
