## Supplementary material for "Rule-Based Deconstruction and Reconstruction of Diterpene Libraries: Categorizing Patterns & Unravelling the Structural Landscape": Zip of all Code used: Supp_Code.5.Backbone_MSA.ipynb

```
# %% [code]
# IMPORTED LIBRARIES
import pandas as pd
import numpy as np
import re
import os
import scipy
import statsmodels
import matplotlib.pyplot as plt

import rdkit
from rdkit import Chem
from rdkit.Chem import AllChem
from rdkit import DataStructs
from rdkit.Chem.rdMolDescriptors import CalcMolFormula
from rdkit.Chem.Draw import IPythonConsole
from rdkit.Chem import Draw

# %% [code]
# PRINT LIBRARY VERSIONS
print("Pandas:",pd.__version__)
print("Numpy:",np.__version__)
print("RE:", re.__version__)
print("Scipy:", scipy.__version__)
print("Statsmodels:", statsmodels.__version__)
print("Matplotlib:",plt.matplotlib.__version__)
print("RDKit:", rdkit.__version__)
print("OS:",os.__file__)

# %% [markdown]
## Overview of program
## Supplemental data used and generated by this program

# %% [code]
#Supplemental Code 5: ATOMIC ALIGNMENT OF DITERPENE BACKBONES

### DATASET(S) INTENDED AS INPUT FOR THIS CODE
# PAIRED FOR DNP
# Supp.2a.DNP_Diterpene_Mining_v30.2_Updated_Skeleton_Backbone.tsv
# Supp.2e.DNP_Diterpene_Skeleton_Summary.tsv

# PAIRED FOR TEROKIT
# Supp.2b.TeroKit_Diterpene_v2.0_Updated_Skeleton_Backbone.tsv
# Supp.2f.Terokit_Diterpene_Skeleton_Summary.tsv 

### DATASET OUTPUT(S) (20 IN TOTAL)
# Figure 5
# Example SMILES alignment output 
# Supp.7.Backbone_IQV
#    Skeleton_X
#        X_consensus.txt --- Skeleton #X consensus SMILE
#        X_EDGE_IQV.tsv ---- Calculated IQV values for each bond (edge) 
#                            and description of node-node connections
#        X_MSA.txt --------- All aligned SMILES for a specific skeleton
#        X_NODE_IQV.tsv ---- Calculated IQV values for each atom (node)
#        X_ref_image.jpg --- Visualized structure of X skeleton

'''
### OVERVIEW OF FUNCTIONALITIY ###
### Davis Mathieu
### Jan 17, 2024

This program takes all deconstructed compounds with the same 
skeleton and aligns them so each atom and bond completely 
parrallels the reference compound skeleton SMILE. This is 
accomplished by converting all molecules within a skeleton group
to be the same number of atoms by replacing Carbons with residual
R-groups (represented by -OH) to Xenon. These Xenon containing
compounds are then subsequentely aligned to the reference and
stereocenters are changed to conform to the bonds that differ.
Occasionally SMILEs do not inherently align and two programs to 
make them align are implemented (attempted_consolidation and 
forced_consolidation). The first attempt is computationally
efficient but error prone, while the second is accurate but 
computationally and time intensive. With both of these programs
all diterpenes were able to be aligned with the kaurenes being the 
only major hinderance (over 24 hours to run).

After SMILEs are aligned, the index of qualitative variation 
calculation is used to determine how often a specific atom or 
bond position differs among compounds with that skeleton. SMILEs
are deconstructed to determine which Carbon are connected to other
Carbon (determining the bonds that represent them). These represent
the bonds (or edge) values for calculating variability. IQV values
for atoms are determined atom-by-atom indexed from 0-19 (for 
diterpenes).

Functions:
- Add_Xenon:               Replaces all occurances of COH with 
                           Xenon

- attempted_consolidation  Takes SMILEs that cannot be aligned to 
                           a reference SMILE and takes a "best 
                           guess" approach for alignment

- forced_consolidation     Takes SMILEs that cannot be aligned by
                           any other means and produces and compares
                           all possible ways that a compound can 
                           exist and compares these (computationally
                           demanding)

- convert_stereocenters    Takes all symbols representing [C@], [C@@],
                           [C@H], or [C@@H] and determines which bonds
                           they represent and identifies their chiral
                           directionality

- id_edge_intensity        Takes a SMILE and determines which bonds
                           correlate to which edge and determines IQV

- percent_diversity        given the position of a particular atom in 
                           a SMILEs string this program goes through
                           the MSA file(s) and calculates the occurances
                           of decoration (X) and divides by the total 
                           occurances.

- Primary_Carbon_stats     Investigates positional based variance for a 
                           compound depending on the neighboring carbon
                           connections for all occurances of primary methyl
                           groups.
                          
- Secondary_Carbon_stats   Same as Primary_Carbon_stats, but for all
                           occurances of Secondary carbons and their 
                           possible neighbors.
- Tertiary_Carbon_stats    Same as Primary_Carbon_stats, but for all
                           occurances of Tertiary carbons and their 
                           possible neighbors.

#-----------------------------------------------------------------#
This program produces an MSA for atoms with the same skeleton but
differing backbones/decoration and determines which areas of the 
molecule (atom and/or bonds) differ. This generates EDGE/NODE
tables that corresponds to that skeleton and variation that is 
present within the data. 

'''
x=0

# %% [markdown]
## Function for replacing Carbons with R-groups (-OH) with Xenon instead
---
##### This is done so all structures can have the same number of atoms

# %% [code]
def Add_Xenon(compound):
    Xenon=Chem.MolFromSmarts("[#54]")
    Carbon=Chem.MolFromSmarts("[#6]")
    formula = compound
    test = formula.GetSubstructMatches(Chem.MolFromSmarts("[#6:0]-[#6D2:1]-[#8:2]"))
    while(len(test)>0):
        formula = Chem.MolFromSmiles(Chem.MolToSmiles(formula))
        TEST = AllChem.ReactionFromSmarts("[#6:0]-[#6D2:1]-[#8:2].[#54:4]>>[#6:0][#54H0:4].[#6:1]-[#8:2]")
        test2 = TEST.RunReactants((formula,Xenon))
        for i in range(np.shape(test2)[0]):
            if(len(re.findall("C",Chem.MolToSmiles(formula),re.IGNORECASE))==len(re.findall("C",Chem.MolToSmiles(test2[i][0]),re.IGNORECASE))+1):
                formula = test2[i][0]
        test = formula.GetSubstructMatches(Chem.MolFromSmarts("[#6:0]-[#6D2:1]-[#8:2]"))

    test = formula.GetSubstructMatches(Chem.MolFromSmarts("[#6:0]=[#6D2:1]-[#8:2]"))
    while(len(test)>0):
        formula = Chem.MolFromSmiles(Chem.MolToSmiles(formula))
        TEST = AllChem.ReactionFromSmarts("[#6:0]=[#6D2:1]-[#8:2].[#54:4]>>[#6:0]=[#54H0:4].[#6:1]-[#8:2]")
        test2 = TEST.RunReactants((formula,Xenon))
        for i in range(np.shape(test2)[0]):
            if(len(re.findall("C",Chem.MolToSmiles(formula),re.IGNORECASE))==len(re.findall("C",Chem.MolToSmiles(test2[i][0]),re.IGNORECASE))+1):
                formula = test2[i][0]
        test = formula.GetSubstructMatches(Chem.MolFromSmarts("[#6:0]=[#6D2:1]-[#8:2]"))        

    test = formula.GetSubstructMatches(Chem.MolFromSmarts("[#6:0]-[#6D3:1](-[#6:3])-[#8:2]"))
    while(len(test)>0):
        formula = Chem.MolFromSmiles(Chem.MolToSmiles(formula))
        TEST = AllChem.ReactionFromSmarts("[#6:0]-[#6D3:1](-[#6:3])-[#8:2].[#54:5]>>[#6:0]-[#54D3H0:5](-[#6:3]).[#6:1][#8:2]")
        test2 = TEST.RunReactants((formula,Xenon))
        for i in range(np.shape(test2)[0]):
            if(len(re.findall("C",Chem.MolToSmiles(formula),re.IGNORECASE))==len(re.findall("C",Chem.MolToSmiles(test2[i][0]),re.IGNORECASE))+1):
                formula = test2[i][0]
        test = formula.GetSubstructMatches(Chem.MolFromSmarts("[#6:0]-[#6D3:1](-[#6:3])-[#8:2]"))        

    test = formula.GetSubstructMatches(Chem.MolFromSmarts("[#54:0]-[#6D3:1](-[#6:3])-[#8:2]"))
    while(len(test)>0):
        formula = Chem.MolFromSmiles(Chem.MolToSmiles(formula))
        TEST = AllChem.ReactionFromSmarts("[#54:0]-[#6D3:1](-[#6:3])-[#8:2].[#54:5]>>[#54:0]-[#54D3H0:5](-[#6:3]).[#6:1][#8:2]")
        test2 = TEST.RunReactants((formula,Xenon))
        for i in range(np.shape(test2)[0]):
            if(len(re.findall("C",Chem.MolToSmiles(formula),re.IGNORECASE))==len(re.findall("C",Chem.MolToSmiles(test2[i][0]),re.IGNORECASE))+1):
                formula = test2[i][0]
        test = formula.GetSubstructMatches(Chem.MolFromSmarts("[#54:0]-[#6D3:1](-[#6:3])-[#8:2]")) 

    test = formula.GetSubstructMatches(Chem.MolFromSmarts("[#54:0]-[#6D3:1](-[#54:3])-[#8:2]"))
    while(len(test)>0):
        formula = Chem.MolFromSmiles(Chem.MolToSmiles(formula))
        TEST = AllChem.ReactionFromSmarts("[#54:0]-[#6D3:1](-[#54:3])-[#8:2].[#54:5]>>[#54:0]-[#54D3H0:5](-[#54:3]).[#6:1][#8:2]")
        test2 = TEST.RunReactants((formula,Xenon))
        for i in range(np.shape(test2)[0]):
            if(len(re.findall("C",Chem.MolToSmiles(formula),re.IGNORECASE))==len(re.findall("C",Chem.MolToSmiles(test2[i][0]),re.IGNORECASE))+1):
                formula = test2[i][0]
        test = formula.GetSubstructMatches(Chem.MolFromSmarts("[#54:0]-[#6D3:1](-[#54:3])-[#8:2]"))         

    test = formula.GetSubstructMatches(Chem.MolFromSmarts("[#6:0]=[#6D3:1](-[#6:3])-[#8:2]"))
    while(len(test)>0):
        formula = Chem.MolFromSmiles(Chem.MolToSmiles(formula))
        TEST = AllChem.ReactionFromSmarts("[#6:0]=[#6D3:1](-[#6:3])-[#8:2].[#54:5]>>[#6:0]=[#54D3H0:5](-[#6:3]).[#6:1][#8:2]")
        test2 = TEST.RunReactants((formula,Xenon))
        for i in range(np.shape(test2)[0]):
            if(len(re.findall("C",Chem.MolToSmiles(formula),re.IGNORECASE))==len(re.findall("C",Chem.MolToSmiles(test2[i][0]),re.IGNORECASE))+1):
                formula = test2[i][0]
        test = formula.GetSubstructMatches(Chem.MolFromSmarts("[#6:0]=[#6D3:1]([#6:3])-[#8:2]"))         

    test = formula.GetSubstructMatches(Chem.MolFromSmarts("[#6:0]=[#6D3:1]([#8:2])-[#54:3]"))
    while(len(test)>0):
        formula = Chem.MolFromSmiles(Chem.MolToSmiles(formula))
        TEST = AllChem.ReactionFromSmarts("[#6:0]=[#6D3:1]([#8:2])-[#54:3].[#54:5]>>[#6:0]=[#54D2H0:5]-[#54:3].[#6D3:1]([#8:2])") 
        test2 = TEST.RunReactants((formula,Xenon))
        for i in range(np.shape(test2)[0]):
            if(len(re.findall("C",Chem.MolToSmiles(formula),re.IGNORECASE))==len(re.findall("C",Chem.MolToSmiles(test2[i][0]),re.IGNORECASE))+1):
                formula = test2[i][0]
        test = formula.GetSubstructMatches(Chem.MolFromSmarts("[#6:0]=[#6D3:1]([#8:2])-[#54:3]")) 
       
    
    test = formula.GetSubstructMatches(Chem.MolFromSmarts("[#6:0][#6D4:1]([#6:3])([#6:4])-[#8:2]"))
    while(len(test)>0):
        formula = Chem.MolFromSmiles(Chem.MolToSmiles(formula))
        TEST = AllChem.ReactionFromSmarts("[#6:0]-[#6D4:1](-[#6:3])(-[#6:4])-[#8:2].[#54:5]>>[#6:0]-[#54D3H0:5](-[#6:3])(-[#6:4]).[#6:1][#8:2]")
        test2 = TEST.RunReactants((formula,Xenon))
        for i in range(np.shape(test2)[0]):
            if(len(re.findall("C",Chem.MolToSmiles(formula),re.IGNORECASE))==len(re.findall("C",Chem.MolToSmiles(test2[i][0]),re.IGNORECASE))+1):
                formula = test2[i][0]
        test = formula.GetSubstructMatches(Chem.MolFromSmarts("[#6:0]-[#6D4:1](-[#6:3])(-[#6:4])-[#8:2]")) 
        
    test = formula.GetSubstructMatches(Chem.MolFromSmarts("[#54:0]-[#6D4:1](-[#6:3])(-[#6:4])-[#8:2]"))
    while(len(test)>0):
        formula = Chem.MolFromSmiles(Chem.MolToSmiles(formula))
        TEST = AllChem.ReactionFromSmarts("[#54:0]-[#6D4:1](-[#6:3])(-[#6:4])-[#8:2].[#54:5]>>[#54:0]-[#54D3H0:5](-[#6:3])(-[#6:4]).[#6:1][#8:2]")
        test2 = TEST.RunReactants((formula,Xenon))
        for i in range(np.shape(test2)[0]):
            if(len(re.findall("C",Chem.MolToSmiles(formula),re.IGNORECASE))==len(re.findall("C",Chem.MolToSmiles(test2[i][0]),re.IGNORECASE))+1):
                formula = test2[i][0]
        test = formula.GetSubstructMatches(Chem.MolFromSmarts("[#54:0]-[#6D4:1](-[#6:3])(-[#6:4])-[#8:2]")) 
    
    test = formula.GetSubstructMatches(Chem.MolFromSmarts("[#54:0]-[#6D4:1](-[#54:3])(-[#6:4])-[#8:2]"))
    while(len(test)>0):
        formula = Chem.MolFromSmiles(Chem.MolToSmiles(formula))
        TEST = AllChem.ReactionFromSmarts("[#54:0]-[#6D4:1](-[#54:3])(-[#6:4])-[#8:2].[#54:5]>>[#54:0]-[#54D3H0:5](-[#54:3])(-[#6:4]).[#6:1][#8:2]")
        test2 = TEST.RunReactants((formula,Xenon))
        for i in range(np.shape(test2)[0]):
            if(len(re.findall("C",Chem.MolToSmiles(formula),re.IGNORECASE))==len(re.findall("C",Chem.MolToSmiles(test2[i][0]),re.IGNORECASE))+1):
                formula = test2[i][0]
        test = formula.GetSubstructMatches(Chem.MolFromSmarts("[#54:0]-[#6D4:1](-[#54:3])(-[#6:4])-[#8:2]")) 

    test = formula.GetSubstructMatches(Chem.MolFromSmarts("[#54:0]-[#6D4:1](-[#54:3])(-[#54:4])-[#8:2]"))
    while(len(test)>0):
        formula = Chem.MolFromSmiles(Chem.MolToSmiles(formula))
        TEST = AllChem.ReactionFromSmarts("[#54:0]-[#6D4:1](-[#54:3])(-[#54:4])-[#8:2].[#54:5]>>[#54:0]-[#54D3H0:5](-[#54:3])(-[#54:4]).[#6:1][#8:2]")
        test2 = TEST.RunReactants((formula,Xenon))
        for i in range(np.shape(test2)[0]):
            if(len(re.findall("C",Chem.MolToSmiles(formula),re.IGNORECASE))==len(re.findall("C",Chem.MolToSmiles(test2[i][0]),re.IGNORECASE))+1):
                formula = test2[i][0]
        test = formula.GetSubstructMatches(Chem.MolFromSmarts("[#54:0]-[#6D4:1](-[#54:3])(-[#54:4])-[#8:2]")) 

    test = formula.GetSubstructMatches(Chem.MolFromSmarts("[#54:0]=[#6D3:1](-[#8:2])-[#54:3]"))
    while(len(test)>0):
        formula = Chem.MolFromSmiles(Chem.MolToSmiles(formula))
        TEST = AllChem.ReactionFromSmarts("[#54:0]=[#6D3:1](-[#8:2])-[#54:3].[#54:5]>>[#54:0]=[#54D2H0:5]-[#54:3].[#6:1]([#8:2])") 
        test2 = TEST.RunReactants((formula,Xenon))
        for i in range(np.shape(test2)[0]):
            if(len(re.findall("C",Chem.MolToSmiles(formula),re.IGNORECASE))==len(re.findall("C",Chem.MolToSmiles(test2[i][0]),re.IGNORECASE))+1):
                formula = test2[i][0]
        test = formula.GetSubstructMatches(Chem.MolFromSmarts("[#54:0]=[#6D3:1]([#8:2])-[#54:3]")) 
    
    #Dealing with Aromatics with R groups; Aromatics do not like Xenon
    Carbon=Chem.MolFromSmarts("[17C]")
    aromatic = formula.GetSubstructMatches(Chem.MolFromSmarts("[#6:0]:[#6:1](-[#8:2]):[#6:3]"))
    while(len(aromatic)>0):
        formula = Chem.MolFromSmiles(Chem.MolToSmiles(formula))
        TEST = AllChem.ReactionFromSmarts("[#6:0]:[#6:1](-[#8:2]):[#6:3].[17C:5]>>[#6:0]:[17C:5]:[#6:3].[#6:1](-[#8:2])")
        formula = TEST.RunReactants((formula,Carbon))[0][0]
        aromatic = formula.GetSubstructMatches(Chem.MolFromSmarts("[#6:0]:[#6:1](-[#8:2]):[#6:3]"))
    
    return(formula)

# %% [markdown]
## Function for attempting to align sequences that don't intuitively do so
---
##### This method of "consolidating" problematic SMILEs strings is the faster method and uses a best guess approach, however still isn't perfect

# %% [code]
#consensus_name = np.unique(temp,return_counts=True)#[0]#[list(np.unique(temp,return_counts=True)[1])#.index(max(np.unique(temp,return_counts=True)[1]))]
#consensus_name = np.unique(temp, return_counts=True)[0][0]
def attempted_consolidation(place_holder, consensus_name2):
    foo = consensus_name2; Best_Match=0

    #### This needs to add '-'' for every occurance of "#" there is in the molecule
    if(re.search(r"\#",place_holder)!=None):
        cycles = int(len(re.findall("\#",place_holder)))
        for k in range(int(cycles)):    
        #### This block will add a Xenon at all locations 
            counter = 0; placement=0; carbon_number = 1; temp3 =[] 
            for i in foo:
                #print(i, counter, placement, carbon_number)
                if(i=="-"):
                    placement= placement+1
                if(placement==carbon_number and i=="-"):
                    modified_smile = foo[:counter] + str("=") + foo[counter+1:]
                    temp3.append(modified_smile)
                    #print(modified_smile)
                    carbon_number=carbon_number+1
                counter=counter+1
            #### Need to now loop through and find compound with the best match to move forward with again
            Best_Match = 0
            for j in temp3:
                i = j
                i = Chem.MolFromSmiles(i)
                if(i!=None):
                    compare2 = re.sub("\[Xe\]","C",place_holder)
                    Comparison_Score = DataStructs.FingerprintSimilarity(Chem.RDKFingerprint(i),Chem.RDKFingerprint(Chem.MolFromSmiles(compare2)), metric=DataStructs.TanimotoSimilarity)
                    if(Comparison_Score>Best_Match):
                        Best_Match=Comparison_Score
                        foo = j

    #### This needs to add - for every occurance of = there is in the molecule
    if(re.search("\=",place_holder)!=None):
        cycles = int(len(re.findall("\=",place_holder)))
        for k in range(int(cycles)):    
        #### This block will add a Xenon at all locations 
            counter = 0; placement=0; carbon_number = 1; temp3 =[] 
            for i in foo:
                #print(i, counter, placement, carbon_number)
                if(i=="-"):
                    placement= placement+1
                if(placement==carbon_number and i=="-"):
                    modified_smile = foo[:counter] + str("=") + foo[counter+1:]
                    temp3.append(modified_smile)
                    #print(modified_smile)
                    carbon_number=carbon_number+1
                counter=counter+1
            #### Need to now loop through and find compound with the best match to move forward with again
            Best_Match = 0
            for j in temp3:
                i = j
                i = Chem.MolFromSmiles(i)
                if(i!=None):
                    compare2 = re.sub(r"\[Xe\]","C",place_holder)
                    Comparison_Score = DataStructs.FingerprintSimilarity(Chem.RDKFingerprint(i),Chem.RDKFingerprint(Chem.MolFromSmiles(compare2)), metric=DataStructs.TanimotoSimilarity)
                    if(Comparison_Score>Best_Match):
                        Best_Match=Comparison_Score
                        foo = j
    foo = [foo]

    #### This needs to add Xe for every occurance there is in the molecule
    if(re.search("Xe",CalcMolFormula(Chem.MolFromSmiles(place_holder)))!=None):
        cycles = re.findall("\d+",CalcMolFormula(Chem.MolFromSmiles(place_holder)))[-1]
        for k in range(int(cycles)):    
        #### This block will add a Xenon at all locations 
            #counter = 0; placement=0; carbon_number = 1; temp3 =[] 
            for j in foo:
                counter = 0; placement=0; carbon_number = 1; temp3 =[] 
                for i in j:
                #print(i, counter, placement, carbon_number)
                    if(i=="C"):
                        placement= placement+1
                    if(placement==carbon_number and i=="C"):
                        modified_smile = j[:counter] + str("Z") + j[counter+1:]
                        modified_smile = re.sub("Z", "[Xe]", modified_smile)
                        temp3.append(modified_smile)
                        carbon_number=carbon_number+1
                    counter=counter+1
            #### Need to now loop through and find compound with the best match to move forward with again
            Best_Match = 0; top_5 = []; top_5_score = []
            for j in temp3:
                i = j
                i = Chem.MolFromSmiles(i)

                #Comparison_Score = DataStructs.FingerprintSimilarity(Chem.RDKFingerprint(i),Chem.RDKFingerprint(Chem.MolFromSmiles(place_holder)),metric=DataStructs.TanimotoSimilarity)
                Comparison_Score = DataStructs.AllBitSimilarity(AllChem.GetMorganFingerprintAsBitVect(i, radius=4, nBits=1024),AllChem.GetMorganFingerprintAsBitVect(Chem.MolFromSmiles(place_holder), radius=4, nBits=1024))
                
                #Append all values here then compare after for which top ones are the best
                top_5.append(j)
                top_5_score.append(Comparison_Score)
                if(Comparison_Score==1):
                    Best_Match=Comparison_Score
                    foo = j
                    break

                    
            indices = sorted(range(len(top_5_score)), key=lambda i: top_5_score[i], reverse=True)[:5]
            if(Best_Match!=1):
                foo = []
            else:break
            for k in indices:
                foo.append(top_5[k])
           
    if(Best_Match==1):
        if(type(foo)==list):foo=foo[0]
        foo = re.sub("\[Xe\]","X",foo)
        foo = re.sub("17","",foo)
        foo = re.sub("N","X",foo, re.IGNORECASE)
        foo = re.sub("\[C\@\@H\]","U",foo)
        foo = re.sub("\[C\@H\]", "D", foo)
        foo = re.sub("\[C\]", "C", foo)
        foo = re.sub("\[CH\]", "C", foo)
        foo = re.sub("\[C\@H\]", "D", foo)
        foo = re.sub("\[C\@\]", "L", foo)
        foo = re.sub("\[C\@\@\]", "R", foo)
        return(foo, Best_Match)
    return("PROBLEM", 0)

# %% [markdown]
## Function for forcing SMILEs alignment that don't intuitively do so
---
##### This method of "consolidating" problematic SMILEs strings will do the job but is more time intensive. Most of the programs computational time is done from this program. It can make upwards of 100K compounds to compare for each case where consolidation doesn't happen.. Be aware, this would be worse with compounds containing more than 20C

# %% [code]
#consensus_name = np.unique(temp,return_counts=True)#[0]#[list(np.unique(temp,return_counts=True)[1])#.index(max(np.unique(temp,return_counts=True)[1]))]
#consensus_name = np.unique(temp, return_counts=True)[0][0]
def forced_consolidation(place_holder, consensus_name2, max_cycle=20):
    original = place_holder
    foo = consensus_name2
    place_holder = re.sub("\[17cH\]", "N", place_holder)
    place_holder = re.sub("c","C",place_holder)    
    #### This needs to add Xe for every occurance there is in the molecule
    if(re.search(r"\#",place_holder)!=None):
        cycles = int(len(re.findall("\#",place_holder)))
        for k in range(int(cycles)):    
        #### This block will add a # bond at all locations 
            counter = 0; placement=0; carbon_number = 1; temp3 =[] 
            for i in foo:
                if(i=="-"):
                    placement= placement+1
                if(placement==carbon_number and i=="-"):
                    modified_smile = foo[:counter] + str("=") + foo[counter+1:]
                    temp3.append(modified_smile)
                    carbon_number=carbon_number+1
                counter=counter+1
            #### Need to now loop through and find compound with the best match to move forward with again
            Best_Match = 0
            for j in temp3:
                i = j
                i = Chem.MolFromSmiles(i)
                if(i!=None):
                    compare2 = re.sub("\[Xe\]","C",place_holder)
                    compare2 = re.sub("N", "C", place_holder)
                    Comparison_Score = DataStructs.FingerprintSimilarity(Chem.RDKFingerprint(i),Chem.RDKFingerprint(Chem.MolFromSmiles(compare2)), metric=DataStructs.TanimotoSimilarity)
                    if(Comparison_Score>Best_Match):
                        Best_Match=Comparison_Score
                        foo = j

    #### This needs to add Xe for every occurance there is in the molecule
    if(re.search("\=",place_holder)!=None):
        compare2 = re.sub(r"\[Xe\]","C",place_holder)
        compare2 = re.sub("N", "C", compare2)
        compare2 = re.sub(":", "-", compare2)
        compare2 = re.sub("\[", "", compare2)
        compare2 = re.sub("\]", "", compare2)
        compare2 = re.sub("@", "", compare2)
        compare2 = re.sub("H", "", compare2)
        cycles = int(len(re.findall("\=",place_holder))); temp3=[]
        for k in range(int(cycles)):    
        #### This block will add a Xenon at all locations 
            counter = 0; placement=0; carbon_number = 1
            if(k==0): 
                for i in foo:
                    if(i=="-"):
                        placement= placement+1
                    if(placement==carbon_number and i=="-"):
                        modified_smile = foo[:counter] + str("=") + foo[counter+1:]
                        temp3.append(modified_smile)
                        carbon_number=carbon_number+1
                    counter=counter+1
                temp3 = np.unique(temp3)
            else:
                temp2 = []
                for i in temp3:
                    counter = 0; placement=0; carbon_number = 1; 
                    for j in i:
                        if(j=="-"):
                            placement= placement+1
                        if(placement==carbon_number and j=="-"):
                            modified_smile = i[:counter] + str("=") + i[counter+1:]
                            temp2.append(modified_smile)
                            carbon_number=carbon_number+1
                        counter=counter+1
                temp3 = np.unique(temp2)
        #### Need to now loop through and find compound with the best match to move forward with again
        Best_Match = 0
        for j in temp3:
            i = j
            i = Chem.MolFromSmiles(i)
            if(i!=None):
                #Comparison_Score = DataStructs.FingerprintSimilarity(Chem.RDKFingerprint(i),Chem.RDKFingerprint(Chem.MolFromSmiles(compare2)), metric=DataStructs.TanimotoSimilarity)
                Comparison_Score = DataStructs.AllBitSimilarity(AllChem.GetMorganFingerprintAsBitVect(i, radius=4, nBits=1024),AllChem.GetMorganFingerprintAsBitVect(Chem.MolFromSmiles(compare2), radius=4, nBits=1024))
                if(Comparison_Score==1):
                    foo = j
                    break

    if(DataStructs.FingerprintSimilarity(Chem.RDKFingerprint(Chem.MolFromSmiles(foo)), Chem.RDKFingerprint(Chem.MolFromSmiles(original)),metric=DataStructs.TanimotoSimilarity)==1):
        return(foo, 1.0)

##############################
    if(re.search(":",place_holder)!=None):
        #Identify locations that originally had "=" and remove those from being replace with ":"
        placement = 0; to_drop = []
        match = foo
        for i in foo:
            if(i=="="):
                to_drop.append(placement)
            placement+=1
            
        cycles=int(len(re.findall(":", place_holder))/2)
        for k in range(cycles):
            if(type(foo)!=list):
                placement = 0; temp = []
                for i in foo:
                    if(i=="-"):
                        temp.append(foo[0:placement]+"="+foo[placement+1:len(foo)])
                    placement+=1
            else:
                temp =[]
                for i in foo:
                    placement=0;
                    for j in i:
                        if(j=="-"):
                            temp.append(i[0:placement]+"="+i[placement+1:len(i)])
                        placement+=1
            foo = list(np.unique(temp))
        temp = [] 
        
        # Compare molecules to original only if they are physically possible with RDKit
        test = re.sub(r"\[Xe\]", "C", place_holder)
        test = re.sub("N", "C", test)
        k = Chem.MolFromSmiles(test)
        for i in foo:
            if(Chem.MolFromSmiles(i)!=None):
                Comparison_Score = DataStructs.FingerprintSimilarity(Chem.RDKFingerprint(Chem.MolFromSmiles(i)),Chem.RDKFingerprint(Chem.MolFromSmiles(test)), metric=DataStructs.TanimotoSimilarity)
                if(Comparison_Score== 1):
                    temp.append(i)
        #For those that match, identify all of the locations where "=" can be and match the aromatic
        placement = 0; aromatic_location=[]
        for i in temp:
            placement = 0
            for j in i:
                if(j=="="):
                    aromatic_location.append(placement)
                placement+=1
        aromatic_location = np.unique(aromatic_location)

        #Only keep aromatic locations
        result_list = [x for x in aromatic_location if x not in to_drop] ; placement=0; foo = ""
        #Rewrite new SMILEs with proper
        placement=0; foo =""
        for i in match:
            if(placement in result_list):
                foo =foo+":"
            else:
                foo=foo+i
            placement+=1
            
#######################
    if(re.search("N", CalcMolFormula(Chem.MolFromSmiles(place_holder)))!=None):
        x =0
        cycles = len(re.findall("N", place_holder, re.IGNORECASE))
        counter = 0; placement=0; carbon_number = 1; temp3 =[]
        for k in range(int(cycles)):
            for i in foo:
                if(type(foo)==list):
                    counter = 0
                    for j in i:
                        if(j=="C"):
                            placement= placement+1
                        if(placement==carbon_number and j=="C"):
                            modified_smile = i[:counter] + str("N") + i[counter+1:]
                            temp3.append(modified_smile)
                            carbon_number=carbon_number+1
                        counter=counter+1
                else:
                    if(i=="C"):
                        placement= placement+1
                    if(placement==carbon_number and i=="C"):
                        modified_smile = foo[:counter] + str("N") + foo[counter+1:]
                        temp3.append(modified_smile)
                        carbon_number=carbon_number+1
                    counter=counter+1
            foo = list(np.unique(temp3))
        to_match = Chem.MolFromSmiles(re.sub(r"\[Xe\]", "C", place_holder))
        for k in temp3:
            mol = Chem.MolFromSmiles(k)
            if(mol!=None):
                Comparison_Score = DataStructs.AllBitSimilarity(AllChem.GetMorganFingerprintAsBitVect(mol, radius=4, nBits=1024),AllChem.GetMorganFingerprintAsBitVect(to_match, radius=4, nBits=1024))
                if(Comparison_Score==1):
                    foo=k
                    break
                    
#######################################################
    #### This needs to add Xe for every occurance there is in the molecule
    if(re.search("Xe",CalcMolFormula(Chem.MolFromSmiles(place_holder)))!=None):
        cycles = len(re.findall("Xe",place_holder))
    else:cycles=0
    counter = 0; placement=0; carbon_number = 1; temp3 =[]
    if(int(cycles)>max_cycle):
        return(place_holder, 0)
    for k in range(int(cycles)):    
        #### This block will add a Xenon at all locations          
            for i in foo:
                if(type(foo)==list):
                    counter = 0
                    for j in i:
                        if(j=="C"):
                            placement= placement+1
                        if(placement==carbon_number and j=="C"):
                            modified_smile = i[:counter] + str("Z") + i[counter+1:]
                            modified_smile = re.sub("Z", "[Xe]", modified_smile)
                            temp3.append(modified_smile)
                            carbon_number=carbon_number+1
                        counter=counter+1
                else:
                    if(i=="C"):
                        placement= placement+1
                    if(placement==carbon_number and i=="C"):
                        modified_smile = foo[:counter] + str("Z") + foo[counter+1:]
                        modified_smile = re.sub("Z", "[Xe]", modified_smile)
                        temp3.append(modified_smile)
                        carbon_number=carbon_number+1
                    counter=counter+1
                    
            if(type(temp3)==list and k<1):

                Best_Match = 0; top_5 = []; top_5_score = []
                for j in temp3:
                    mol = j
                    mol = Chem.MolFromSmiles(mol)
                    if(mol!=None):
                        Comparison_Score = DataStructs.AllBitSimilarity(AllChem.GetMorganFingerprintAsBitVect(mol, radius=4, nBits=1024),AllChem.GetMorganFingerprintAsBitVect(Chem.MolFromSmiles(place_holder), radius=4, nBits=1024))
                    else: Comparison_Score=0
                    #Append all values here then compare after for which top ones are the best
                    top_5.append(j)
                    top_5_score.append(Comparison_Score)
                    if(Comparison_Score==1):
                        Best_Match=Comparison_Score
                        foo = j
                        foo = re.sub(r"\[Xe\]","X",foo)
                        foo = re.sub("N","X",foo, re.IGNORECASE)
                        foo = re.sub("\[C\@\@H\]","U",foo)
                        foo = re.sub("\[C\@H\]", "D", foo)
                        foo = re.sub("\[C\]", "C", foo)
                        foo = re.sub("\[CH\]", "C", foo)
                        foo = re.sub("\[C\@H\]", "D", foo)
                        foo = re.sub("\[C\@\]", "L", foo)
                        foo = re.sub("\[C\@\@\]", "R", foo)
                        return(foo, Best_Match)
                indices = sorted(range(len(top_5_score)), key=lambda i: top_5_score[i], reverse=True)[:5]
                if(Best_Match!=1):
                    foo = []
                else:break
                for k in indices:
                    foo.append(top_5[k])
                temp3 = foo
            temp3 = list(np.unique(temp3))
            #print(len(temp3))
            foo = temp3; temp3=[]
            #### Need to now loop through and find compound with the best match to move forward with again
    Best_Match = 0
    temp3 = foo
    if(type(temp3)==str):temp3=[temp3]
    for j in temp3:
        i = j
        i = Chem.MolFromSmiles(i)
        if(i!=None):
            Comparison_Score = DataStructs.FingerprintSimilarity(Chem.RDKFingerprint(i),Chem.RDKFingerprint(Chem.MolFromSmiles(place_holder)), metric=DataStructs.TanimotoSimilarity)
        else:Comparison_Score=0
        if(Comparison_Score>Best_Match):
            Best_Match=Comparison_Score
            foo = j
            if(Best_Match==1):
                foo = re.sub(r"\[Xe\]","X",foo)
                foo = re.sub("N","X",foo)
                foo = re.sub("\[C\@\@H\]","U",foo)
                foo = re.sub("\[C\@H\]", "D", foo)
                foo = re.sub("\[C\]", "C", foo)
                foo = re.sub("\[CH\]", "C", foo)
                foo = re.sub("\[C\@H\]", "D", foo)
                foo = re.sub("\[C\@\]", "L", foo)
                foo = re.sub("\[C\@\@\]", "R", foo)
                return(foo, Best_Match)
            

    return("PROBLEM", 0)

# %% [markdown]
## Takes stereocenters referenced by atoms and translates those to their respective bonds

# %% [code]
# convert stereochemical centers to reflect the stereochemistry at bond sites
def convert_stereocenters(stereo_compound):
#################### FOR THE RESOLUTION OF STEREOCHEMICAL PLACE HOLDER R ################
    while(re.search("R", stereo_compound)):
        if(re.search("R\d\d",stereo_compound)):
            temp = ""
            for i in range(len(stereo_compound)):
                if(i+2<len(stereo_compound) and re.search("R\d\d",stereo_compound[i]+stereo_compound[i+1]+stereo_compound[i+2])):
                    #print(stereo_compound[i]+stereo_compound[i+1]+stereo_compound[i+2])
                    temp +="C"; skip=1;skip1=1;skip2=1
                    for j in range(i+1,len(stereo_compound)):
                        if(j+1<len(stereo_compound) and stereo_compound[j+1]==stereo_compound[i+1] and skip1!=0):
                            temp+="^"
                            skip1=0
                        elif(j+2<len(stereo_compound) and stereo_compound[j+2]==stereo_compound[i+2] and skip2!=0):
                            temp+=stereo_compound[j]
                            skip2=0;skip=0
                        elif(skip==0):
                            temp+="*"
                            skip=1
                        else:
                            temp+=stereo_compound[j]
                    break
                else:temp+=stereo_compound[i]
            stereo_compound=temp
        elif(re.search("R\-\d",stereo_compound)):
            temp = ""; skip=0
            for i in range(len(stereo_compound)):
                if(i+2<len(stereo_compound) and re.search("R\-\d",stereo_compound[i]+stereo_compound[i+1]+stereo_compound[i+2])):
                    temp+="C"
                    skip=2
                elif(stereo_compound[i]=="-" and skip>=1):
                    if(skip==2):
                        temp+="^"
                    elif(skip==1):
                        temp+="*"
                    skip=skip-1
                else:
                    temp+=stereo_compound[i]
            stereo_compound=temp    
  
        elif(re.search("R\d\-",stereo_compound)):
            temp = ""; skip=0
            for i in range(len(stereo_compound)):
                if(i+2<len(stereo_compound) and re.search("R\d\-",stereo_compound[i]+stereo_compound[i+1]+stereo_compound[i+2])):
                    temp+="C"
                    skip=2
                elif(stereo_compound[i]=="-" and skip>=1):
                    if(skip==2):
                        temp+="^"
                    elif(skip==1):
                        temp+="*"
                    skip=skip-1
                else:
                    temp+=stereo_compound[i]
            stereo_compound=temp

        elif(re.search("R\(", stereo_compound)):
            temp = ""; skip=0;skip1=1;skip2=0;skip3=1;R_present=0
            for i in range(len(stereo_compound)):
                if(i+2<len(stereo_compound) and re.search("R\(",stereo_compound[i]+stereo_compound[i+1])):
                    temp+="C"; skip=0;skip1=1;skip2=0;skip3=1;R_present=1
                elif(stereo_compound[i]=="-" and skip1==1 and R_present==1):
                    temp+="^"
                    skip1=0
                elif(stereo_compound[i]=="(" and R_present==1):
                    temp+=stereo_compound[i]
                    skip2=skip2+1
                elif(stereo_compound[i]==")" and R_present==1):
                    temp+=stereo_compound[i]
                    skip2=skip2-1
                elif(stereo_compound[i]=="-" and skip2==1 and skip3==1):
                    temp+="*"
                    skip3=0
                else:
                    temp+=stereo_compound[i]
        elif(re.search("R\d\(",stereo_compound)):
            temp = ""; begin=0; R_search="N3V3R";skip=0
            for i in range(len(stereo_compound)):

                if(i+2<len(stereo_compound) and re.search("R\d\(",stereo_compound[i]+stereo_compound[i+1]+stereo_compound[i+2])):
                    temp+="C"; R_search="-"+stereo_compound[i+1]; skip=1
                elif(i+1<len(stereo_compound) and re.search(R_search, stereo_compound[i]+stereo_compound[i+1])):
                    temp+="*"
                elif(re.search("\(", stereo_compound[i]) and R_search!="N3V3R"):
                    temp+=stereo_compound[i]
                    begin+=1
                elif(re.search("\)", stereo_compound[i]) and R_search!="N3V3R"):
                    temp+=stereo_compound[i]
                    begin-=1
                elif(re.search("\-", stereo_compound[i])  and begin==0 and R_search!="N3V3R"):
                    temp+="^"; begin=100
                else:
                    temp+=stereo_compound[i]
        
        stereo_compound=temp
            
#########################################################################################

#################### FOR THE RESOLUTION OF STEREOCHEMICAL PLACE HOLDER L ################
    while(re.search("L", stereo_compound)):
        #print(stereo_compound)
        if(re.search("L\d\d",stereo_compound)):
            temp = ""
            for i in range(len(stereo_compound)):
                if(i+2<len(stereo_compound) and re.search("L\d\d",stereo_compound[i]+stereo_compound[i+1]+stereo_compound[i+2])):
                    #print(stereo_compound[i]+stereo_compound[i+1]+stereo_compound[i+2])
                    temp +="C"; skip=1;skip1=1;skip2=1
                    for j in range(i+1,len(stereo_compound)):
                        if(j+1<len(stereo_compound) and stereo_compound[j+1]==stereo_compound[i+1] and skip1!=0):
                            temp+="*"
                            skip1=0
                        elif(j+2<len(stereo_compound) and stereo_compound[j+2]==stereo_compound[i+2] and skip2!=0):
                            temp+=stereo_compound[j]
                            skip2=0;skip=0
                        elif(skip==0):
                            temp+="^"
                            skip=1
                        else:
                            temp+=stereo_compound[j]
                    break
                else:temp+=stereo_compound[i]
            
        elif(re.search("L\-\d",stereo_compound)):
            temp = ""; skip=0
            for i in range(len(stereo_compound)):
                if(i+2<len(stereo_compound) and re.search("L\-\d",stereo_compound[i]+stereo_compound[i+1]+stereo_compound[i+2])):
                    temp+="C"
                    skip=2
                elif(stereo_compound[i]=="-" and skip>=1):
                    if(skip==2):
                        temp+="*"
                    elif(skip==1):
                        temp+="^"
                    skip=skip-1
                else:
                    temp+=stereo_compound[i]  
            
        elif(re.search("L\d\D",stereo_compound)):
            temp = ""; skip=0
            for i in range(len(stereo_compound)):
                if(i+2<len(stereo_compound) and re.search("L\d\D",stereo_compound[i]+stereo_compound[i+1]+stereo_compound[i+2])):
                    temp+="C"
                    skip=2
                elif(stereo_compound[i]=="-" and skip>=1):
                    if(skip==2):
                        temp+="*"
                    elif(skip==1):
                        temp+="^"
                    skip=skip-1
                else:
                    temp+=stereo_compound[i]

        elif(re.search("L\(", stereo_compound)):
            temp = ""; skip=0;skip1=1;skip2=0;skip3=1;R_present=0
            for i in range(len(stereo_compound)):
                if(i+2<len(stereo_compound) and re.search("L\(",stereo_compound[i]+stereo_compound[i+1])):
                    temp+="C"; skip=0;skip1=1;skip2=0;skip3=1;R_present=1
                elif(stereo_compound[i]=="-" and skip1==1 and R_present==1):
                    temp+="*"
                    skip1=0
                elif(stereo_compound[i]=="(" and R_present==1):
                    temp+=stereo_compound[i]
                    skip2=skip2+1
                elif(stereo_compound[i]==")" and R_present==1):
                    temp+=stereo_compound[i]
                    skip2=skip2-1
                elif(stereo_compound[i]=="-" and skip2==1 and skip3==1):
                    temp+="^"
                    skip3=0
                else:
                    temp+=stereo_compound[i]
        elif(re.search("L\d\(",stereo_compound)):
            temp = ""; begin=0; R_search="N3V3R";skip=0
            for i in range(len(stereo_compound)):
                if(i+2<len(stereo_compound) and re.search("L\d\(",stereo_compound[i]+stereo_compound[i+1]+stereo_compound[i+2])):
                    temp+="C"; R_search="-"+stereo_compound[i+1]; skip=1
                elif(i+1<len(stereo_compound) and re.search(R_search, stereo_compound[i]+stereo_compound[i+1])):
                    temp+="^"
                elif(re.search("\(", stereo_compound[i]) and R_search!="N3V3R"):
                    temp+=stereo_compound[i]
                    begin+=1
                elif(re.search("\)", stereo_compound[i]) and R_search!="N3V3R"):
                    temp+=stereo_compound[i]
                    begin-=1
                elif(re.search("\-", stereo_compound[i])  and begin==0 and R_search!="N3V3R"):
                    temp+="*"; begin=100
                else:
                    temp+=stereo_compound[i]
        stereo_compound=temp


#########################################################################################

#################### FOR THE RESOLUTION OF STEREOCHEMICAL PLACE HOLDER U ################
    while(re.search("U", stereo_compound)):
        #print(stereo_compound)
        if(re.search("U\(",stereo_compound)):
            temp = ""; skip=0; U_present = True
            for i in range(len(stereo_compound)):
                if(i+3<len(stereo_compound) and re.search("U\(\-",stereo_compound[i]+stereo_compound[i+1]+stereo_compound[i+2])):
                    for j in range(i, len(stereo_compound)):
                        if(j+1<len(stereo_compound) and ")-"==stereo_compound[j]+stereo_compound[j+1]):
                            U_present=False
                            break
                        elif(j+1<len(stereo_compound) and ")("==stereo_compound[j]+stereo_compound[j+1]):
                            U_present = True
                            break
                    if(U_present==False):
                        temp +="C(*"; skip=2
                    else:
                        temp +="C(^"; skip=2
                elif(i+1<len(stereo_compound) and ")("==stereo_compound[i]+stereo_compound[i+1] and U_present):
                    temp+=stereo_compound[i]+stereo_compound[i+1]+"*"; skip=2; U_present=False
                elif(skip!=0):
                    skip=skip-1
                else:
                    temp+=stereo_compound[i]
            stereo_compound=temp
        elif(re.search("\-U",stereo_compound)):
            temp = ""; skip=0
            for i in range(len(stereo_compound)):
                if(i+1<len(stereo_compound) and re.search("\-U",stereo_compound[i]+stereo_compound[i+1])):
                    temp +="^C"; skip=1
                elif(skip==1):
                    skip=0
                else:
                    temp+=stereo_compound[i]      
        
        elif(re.search("U\d\d\-", stereo_compound)):
            temp = ""; skip=0; D_search="N3V3R"
            for i in range(len(stereo_compound)):
                if(i+3<len(stereo_compound) and re.search(r"U\d\d\-",stereo_compound[i]+stereo_compound[i+1]+stereo_compound[i+2]+stereo_compound[i+3])):
                    temp +="C"+stereo_compound[i+1]+stereo_compound[i+2]+"^"; D_search="-"+stereo_compound[i+1]; skip=3 
                elif(skip!=0):
                    skip=skip-1
                elif(i+1<len(stereo_compound) and re.search(D_search, stereo_compound[i]+stereo_compound[i+1])):
                    temp+="*"
                else:
                    temp+=stereo_compound[i] 

        elif(re.search("U\d\(", stereo_compound)):
            temp = ""; skip=0; D_search="N3V3R"; begin=0
            for i in range(len(stereo_compound)):
                if(i+3<len(stereo_compound) and re.search(r"U\d\(\-",stereo_compound[i]+stereo_compound[i+1]+stereo_compound[i+2]+stereo_compound[i+3])):
                    temp +="C"+stereo_compound[i+1]+stereo_compound[i+2]+"^"; D_search="-"+stereo_compound[i+1]; skip=3; begin = 1 
                elif(skip!=0):
                    skip=skip-1
                elif(re.search("\(", stereo_compound[i]) and D_search!="N3V3R"):
                    temp+=stereo_compound[i]
                    begin+=1
                elif(re.search("\)", stereo_compound[i]) and D_search!="N3V3R"):
                    temp+=stereo_compound[i]
                    begin-=1
                elif(re.search("\-", stereo_compound[i])  and begin==0 and D_search!="N3V3R"):
                    temp+="*"; begin=100
                else:
                    temp+=stereo_compound[i] 
        
        elif(re.search(r"\^U",str(stereo_compound))):
            temp = ""; skip=0
            for i in range(len(stereo_compound)):
                if(i+1<len(stereo_compound) and re.search(r"\^U",stereo_compound[i]+stereo_compound[i+1])):
                    temp +="^C"; skip=1
                elif(skip==1):
                    skip=0
                else:
                    temp+=stereo_compound[i]
        stereo_compound=temp
        
#########################################################################################

#################### FOR THE RESOLUTION OF STEREOCHEMICAL PLACE HOLDER D ################
    while(re.search("D", stereo_compound)):
        #print(stereo_compound)
        if(re.search("D\(",stereo_compound)):
            temp = ""; skip=0; U_present = True
            for i in range(len(stereo_compound)):
                if(i+3<len(stereo_compound) and re.search("D\(\-",stereo_compound[i]+stereo_compound[i+1]+stereo_compound[i+2])):
                    for j in range(i, len(stereo_compound)):
                        if(j+1<len(stereo_compound) and ")-"==stereo_compound[j]+stereo_compound[j+1]):
                            U_present=False
                            break
                        elif(j+1<len(stereo_compound) and ")("==stereo_compound[j]+stereo_compound[j+1]):
                            U_present = True
                            break
                    if(U_present==False):
                        temp +="C(^"; skip=2
                    else:
                        temp +="C(*"; skip=2
                elif(i+1<len(stereo_compound) and ")("==stereo_compound[i]+stereo_compound[i+1] and U_present):
                    temp+=stereo_compound[i]+stereo_compound[i+1]+"^"; skip=2; U_present=False
                elif(skip!=0):
                    skip=skip-1
                else:
                    temp+=stereo_compound[i]
            stereo_compound=temp
        elif(re.search("\-D",stereo_compound)):
            temp = ""; skip=0
            for i in range(len(stereo_compound)):
                if(i+1<len(stereo_compound) and re.search("\-D",stereo_compound[i]+stereo_compound[i+1])):
                    temp +="*C"; skip=1
                elif(skip==1):
                    skip=0
                else:
                    temp+=stereo_compound[i]
                           
        elif(re.search("D\d\d\-", stereo_compound)):
            temp = ""; skip=0; D_search="N3V3R"
            for i in range(len(stereo_compound)):
                if(i+3<len(stereo_compound) and re.search(r"D\d\d\-",stereo_compound[i]+stereo_compound[i+1]+stereo_compound[i+2]+stereo_compound[i+3])):
                    temp +="C"+stereo_compound[i+1]+stereo_compound[i+2]+"*"; D_search="-"+stereo_compound[i+1]; skip=3 #>>
                elif(skip!=0):
                    skip=skip-1
                elif(i+1<len(stereo_compound) and re.search(D_search, stereo_compound[i]+stereo_compound[i+1])):
                    temp+="^"
                else:
                    temp+=stereo_compound[i] 
                    
        elif(re.search("D\d\(", stereo_compound)):
            temp = ""; skip=0; D_search="N3V3R"; begin=0
            for i in range(len(stereo_compound)):
                if(i+3<len(stereo_compound) and re.search(r"D\d\(\-",stereo_compound[i]+stereo_compound[i+1]+stereo_compound[i+2]+stereo_compound[i+3])):
                    temp +="C"+stereo_compound[i+1]+stereo_compound[i+2]+"*"; D_search="-"+stereo_compound[i+1]; skip=3; begin = 1 
                elif(skip!=0):
                    skip=skip-1
                elif(re.search("\(", stereo_compound[i]) and D_search!="N3V3R"):
                    temp+=stereo_compound[i]
                    begin+=1
                elif(re.search("\)", stereo_compound[i]) and D_search!="N3V3R"):
                    temp+=stereo_compound[i]
                    begin-=1
                elif(re.search("\-", stereo_compound[i])  and begin==0 and D_search!="N3V3R"):
                    temp+="^"; begin=100
                else:
                    temp+=stereo_compound[i] 
                    
        elif(re.search(r"\^D",str(stereo_compound))):
            temp = ""; skip=0
            for i in range(len(stereo_compound)):
                if(i+1<len(stereo_compound) and re.search(r"\^D",stereo_compound[i]+stereo_compound[i+1])):
                    temp +="*C"; skip=1
                elif(skip==1):
                    skip=0
                else:
                    temp+=stereo_compound[i]
        stereo_compound=temp
    #print(stereo_compound)
    return(stereo_compound)
#x = convert_stereocenters("U(-C)(-C-X)-C-C-C-U(-C)-C-C-C-U(-C)-C-C-C-C(-C)-C")
#                   target: C(^C)(*C-X)-C-C-C-C(*C)-C-C-C-C(*C)-C-C-C-C(-C)-C
#print(x)

# %% [markdown]
## Load datasets and create Multiple Sequence Alignment (MSA)

# %% [code]
# IMPORT PAIRED DATASETS FOR RUNNING MSA ALIGNMENT
Backbones = pd.read_csv("Supp.2b.TeroKit_Diterpene_v2.0_Updated_Skeleton_Backbone.tsv", sep="\t")
Skeletons = pd.read_csv("Supp.2f.Terokit_Diterpene_Skeleton_Summary.tsv", sep="\t")

# Choose which skeleton you want to analyze (0 is most common)
atom_value = 19

# %% [markdown]
# V1.. Faster but does not match to the connonical SMILE

# %% [code]
# PERFORM MSA ALIGNMENT, USING SYMBOLS TO TAKE PLACE OF VARIOUS CONDITIONS BUT STRING LENGTH IS RETAINED

# We create compounds that can't exist, RDKit doesn't like this, best to turn warnings off
from rdkit import RDLogger                                                                                                                                                               
RDLogger.DisableLog('rdApp.*') 

# Which skeleton index would you like to investigate (0 is the most common skeleton) 
atom_value = 19
temp =[]; temp2 =[]

for i in eval(Skeletons["index"].iloc[atom_value]):
    formula = Chem.MolFromSmiles(Backbones["Backbone"].iloc[i])
    #formula, deconstruction = deconstruct2backbone(diterpene)
    formula = Chem.MolFromSmiles(Chem.MolToSmiles(Add_Xenon(formula)))
    if(len(formula.GetSubstructMatches(Chem.MolFromSmarts("[*D4]")))!=0):
        formula = Chem.MolToSmiles(formula, allBondsExplicit=True, rootedAtAtom=formula.GetSubstructMatches(Chem.MolFromSmarts("[*D4]"))[0][0])
    elif(len(formula.GetSubstructMatches(Chem.MolFromSmarts("[*D3]")))!=0):
        formula = Chem.MolToSmiles(formula, allBondsExplicit=True, rootedAtAtom=formula.GetSubstructMatches(Chem.MolFromSmarts("[*D3]"))[0][0])
    else:
        print("ALERT: Something is wrong with this compound", Chem.MolToSmiles(formula))
    
    intermediate = formula
    formula = re.sub("\[Xe\]","C",formula)
    formula = re.sub("\[17cH\]","C",formula)
    formula = re.sub(r"\\", "", formula)
    formula = re.sub(r"\/","", formula)
    formula = re.sub(r"\@","", formula)
    formula = re.sub(r"\[","", formula)
    formula = re.sub(r"\]","", formula)
    formula = re.sub(r"\=", "", formula)
    formula = re.sub(r"\-", "", formula)
    formula = re.sub("H", "", formula)
    formula = re.sub("#","", formula)
    formula = re.sub("\(\)","",formula)
    formula = re.sub(r"/", "-",formula)
    formula = re.sub(r"\\", "-",formula)
    temp.append(formula)

    intermediate = re.sub("\[Xe\]","C",intermediate)
    intermediate = re.sub("\[17cH\]","C",intermediate)
    intermediate = re.sub(r"\\", "-", intermediate)
    intermediate = re.sub(r"\/","-", intermediate)
    intermediate = re.sub(r"\@","", intermediate)
    intermediate = re.sub(r"\[","", intermediate)
    intermediate = re.sub(r"\]","", intermediate)
    intermediate = re.sub(r"\=", "-", intermediate)
    intermediate = re.sub("H", "", intermediate)
    intermediate = re.sub("#","-", intermediate)
    intermediate = re.sub("\(\)","",intermediate)
    intermediate = re.sub(r"/", "-",intermediate)
    intermediate = re.sub(r"\\", "-",intermediate)
    temp2.append(intermediate)
    
consensus_name = np.unique(temp,return_counts=True)[0][list(np.unique(temp,return_counts=True)[1]).index(max(np.unique(temp,return_counts=True)[1]))]
consensus_name2 = np.unique(temp2,return_counts=True)[0][list(np.unique(temp2,return_counts=True)[1]).index(max(np.unique(temp2,return_counts=True)[1]))]
alignment_smiles = []
print(consensus_name2,"\t",len(eval(Skeletons["index"].iloc[atom_value])),"\n")
print("----------------------------------------------------------------------")
#Align all smiles
iteration=0; counter=0
for i in eval(Skeletons["index"].iloc[atom_value]):
    counter+=1
    iteration+=1
    formula = Chem.MolFromSmiles(Backbones["Backbone"].iloc[i])
    #formula, deconstruction = deconstruct2backbone(diterpene)
    formula = Chem.MolFromSmiles(Chem.MolToSmiles(Add_Xenon(formula)))
    intermediate = formula
    canonical_atom_order = Chem.CanonicalRankAtoms(intermediate)
    for j in tuple(canonical_atom_order):
        test = 1
        place_holder = Chem.MolToSmiles(intermediate,rootedAtAtom=j, allBondsExplicit=True)
        place_holder = re.sub(r"/", "-",place_holder)
        place_holder = re.sub(r"\\", "-",place_holder)
        formula = Chem.MolToSmiles(intermediate,rootedAtAtom=j)
        formula = re.sub("\[Xe\]","C",formula)
        formula = re.sub("\[17cH\]","C",formula)
        formula = re.sub(r"\\", "", formula)
        formula = re.sub(r"\/","", formula)
        formula = re.sub(r"\@","", formula)
        formula = re.sub(r"\[","", formula)
        formula = re.sub(r"\]","", formula)
        formula = re.sub(r"\=", "", formula)
        formula = re.sub(r"\-", "", formula)
        formula = re.sub("H", "", formula)
        formula = re.sub("#","", formula)
        formula = re.sub("\(\)","",formula)
        formula = re.sub(r"/", "-",formula)
        formula = re.sub(r"\\", "-",formula)
        if(formula==consensus_name):
            #print(len(list(Chem.MolToSmiles(Chem.MolFromSmiles(consensus_name), allBondsExplicit=True))))
            place_holder = re.sub("\[Xe\]","X",place_holder)
            place_holder = re.sub("\[17CH\]","X",place_holder, re.IGNORECASE)
            place_holder = re.sub("N","X",place_holder, re.IGNORECASE)
            place_holder = re.sub("\[C\@\@H\]","U",place_holder)
            place_holder = re.sub("\[C\@H\]", "D", place_holder)
            place_holder = re.sub("\[C\]", "C", place_holder)
            place_holder = re.sub("\[CH\]", "C", place_holder)
            place_holder = re.sub("\[C\@H\]", "D", place_holder)
            place_holder = re.sub("\[C\@\]", "L", place_holder)
            place_holder = re.sub("\[C\@\@\]", "R", place_holder)
            place_holder = re.sub("\[Xe\]","X",place_holder)
            place_holder = re.sub(r"/", "-",place_holder)
            place_holder = re.sub(r"\\", "-",place_holder)
            alignment_smiles.append(place_holder)
            print(place_holder, "\t",iteration)
            test = 0
            break
        elif(DataStructs.FingerprintSimilarity(Chem.RDKFingerprint(intermediate),Chem.RDKFingerprint(Chem.MolFromSmiles(consensus_name2)), metric=DataStructs.TanimotoSimilarity)==1):
            alignment_smiles.append(consensus_name2)
            test =0
            print(consensus_name2, iteration)
            break
    if(test==1):
        attempt, value_match = attempted_consolidation(place_holder, consensus_name2)
        if(value_match==1):
            alignment_smiles.append(attempt)
            print(attempt,"\t", iteration)
        elif(value_match!=1):
            attempt, value_match = forced_consolidation(place_holder, consensus_name2)
            if(value_match==1):
                alignment_smiles.append(attempt)
                print(attempt,"\t", iteration)
        if(value_match!=1):
            test =1; counter = counter-1
            print("ALERT:",i, place_holder)
            #break
        
        
print("\n",len(list(eval(Skeletons["index"].iloc[atom_value])))-counter, "not aligning from ", len(list(eval(Skeletons["index"].iloc[atom_value]))))

# %% [markdown]
# V2.. Uses Skeleton Connonical SMILE (Slower)

# %% [code]
# PERFORM MSA ALIGNMENT, USING SYMBOLS TO TAKE PLACE OF VARIOUS CONDITIONS BUT STRING LENGTH IS RETAINED

# We create compounds that can't exist, RDKit doesn't like this, best to turn warnings off
from rdkit import RDLogger                                                                                                                                                               
RDLogger.DisableLog('rdApp.*') 

# Which skeleton index would you like to investigate (0 is the most common skeleton) 
atom_value = 19
temp =[]; temp2 =[]

consensus_name = Skeletons["SMILES"].iloc[atom_value]
consensus_name2 = Chem.MolToSmiles(Chem.MolFromSmiles(Skeletons["SMILES"].iloc[atom_value]), allBondsExplicit=True)

alignment_smiles = []
print(consensus_name2,"\t",len(eval(Skeletons["index"].iloc[atom_value])),"\n")
print("----------------------------------------------------------------------")
#Align all smiles
iteration=0; counter=0
for i in eval(Skeletons["index"].iloc[atom_value]):
    counter+=1
    iteration+=1
    formula = Chem.MolFromSmiles(Backbones["Backbone"].iloc[i])
    #formula, deconstruction = deconstruct2backbone(diterpene)
    formula = Chem.MolFromSmiles(Chem.MolToSmiles(Add_Xenon(formula)))
    intermediate = formula
    canonical_atom_order = Chem.CanonicalRankAtoms(intermediate)
    for j in tuple(canonical_atom_order):
        test = 1
        place_holder = Chem.MolToSmiles(intermediate,rootedAtAtom=j, allBondsExplicit=True)
        place_holder = re.sub(r"/", "-",place_holder)
        place_holder = re.sub(r"\\", "-",place_holder)
        formula = Chem.MolToSmiles(intermediate,rootedAtAtom=j)
        formula = re.sub("\[Xe\]","C",formula)
        formula = re.sub("\[17cH\]","C",formula)
        formula = re.sub(r"\\", "", formula)
        formula = re.sub(r"\/","", formula)
        formula = re.sub(r"\@","", formula)
        formula = re.sub(r"\[","", formula)
        formula = re.sub(r"\]","", formula)
        formula = re.sub(r"\=", "", formula)
        formula = re.sub(r"\-", "", formula)
        formula = re.sub("H", "", formula)
        formula = re.sub("#","", formula)
        formula = re.sub("\(\)","",formula)
        formula = re.sub(r"/", "-",formula)
        formula = re.sub(r"\\", "-",formula)
        if(formula==consensus_name):
            #print(len(list(Chem.MolToSmiles(Chem.MolFromSmiles(consensus_name), allBondsExplicit=True))))
            place_holder = re.sub("\[Xe\]","X",place_holder)
            place_holder = re.sub("\[17CH\]","X",place_holder, re.IGNORECASE)
            place_holder = re.sub("N","X",place_holder, re.IGNORECASE)
            place_holder = re.sub("\[C\@\@H\]","U",place_holder)
            place_holder = re.sub("\[C\@H\]", "D", place_holder)
            place_holder = re.sub("\[C\]", "C", place_holder)
            place_holder = re.sub("\[CH\]", "C", place_holder)
            place_holder = re.sub("\[C\@H\]", "D", place_holder)
            place_holder = re.sub("\[C\@\]", "L", place_holder)
            place_holder = re.sub("\[C\@\@\]", "R", place_holder)
            place_holder = re.sub("\[Xe\]","X",place_holder)
            place_holder = re.sub(r"/", "-",place_holder)
            place_holder = re.sub(r"\\", "-",place_holder)
            alignment_smiles.append(place_holder)
            print(place_holder, "\t",iteration)
            test = 0
            break
        elif(DataStructs.FingerprintSimilarity(Chem.RDKFingerprint(intermediate),Chem.RDKFingerprint(Chem.MolFromSmiles(consensus_name2)), metric=DataStructs.TanimotoSimilarity)==1):
            alignment_smiles.append(consensus_name2)
            test =0
            print(consensus_name2, iteration)
            break
    if(test==1):
        attempt, value_match = attempted_consolidation(place_holder, consensus_name2)
        if(value_match==1):
            alignment_smiles.append(attempt)
            print(attempt,"\t", iteration)
        elif(value_match!=1):
            attempt, value_match = forced_consolidation(place_holder, consensus_name2)
            if(value_match==1):
                alignment_smiles.append(attempt)
                print(attempt,"\t", iteration)
        if(value_match!=1):
            test =1; counter = counter-1
            print("ALERT:",i, place_holder)
            #break
        
        
print("\n",len(list(eval(Skeletons["index"].iloc[atom_value])))-counter, "not aligning from ", len(list(eval(Skeletons["index"].iloc[atom_value]))))

# %% [code]


# %% [code]
# CONVERT STEREOCENTERED ATOMS TO THEIR RESPECTIVE BONDS
counter = 1; temp = []; convert = ""
for i in alignment_smiles:
    temp.append(convert_stereocenters(i))
    
check = len(temp[0])
for i in temp:
    if(check!=len(i)):
        print("Uh oh, something doesn't seem quite right...")
        convert = "dont continue"
        break
        
if(convert!="dont continue"):
    print("good to go!")
    alignment_smiles=temp
    

# %% [markdown]
### Specific location of decoration consequently from the CYP701 clan

# %% [code]
# IMPORT PAIRED DATASETS FOR RUNNING MSA ALIGNMENT
Backbones = pd.read_csv("Supp.2b.TeroKit_Diterpene_v2.0_Updated_Skeleton_Backbone.tsv", sep="\t")
Skeletons = pd.read_csv("Supp.2f.Terokit_Diterpene_Skeleton_Summary.tsv", sep="\t")


# %% [code]
# Choose which skeleton you want to analyze (0 is most common)
atom_value = 1
temp = []

count=0
for i in eval(Skeletons["index"].iloc[atom_value]):
    formula = Chem.MolFromSmiles(Backbones["Backbone"].iloc[i])
    #formula, deconstruction = deconstruct2backbone(diterpene)
    formula = Chem.MolFromSmiles(Chem.MolToSmiles(Add_Xenon(formula)))
    temp.append(formula)
    if(count%100==0):
        print(count)
    count=count+1

# %% [code]
formula.GetSubstructMatches(Chem.MolFromSmarts("[#6D4:0](@[#6D2:4])(-[#6D1:1])[#6D3:2][#6D1:3]"))
formula

# %% [code]
#just for claradane
nothin = 0
one = 0
two = 0
check = 0
print(len(temp))
for i in temp:
    #no ring decoration
    no_dec = i.GetSubstructMatches(Chem.MolFromSmarts("[#6D4:0](@[#6D2:4])(~[#6D1:1])[#6D3:2](~[#6D1:3])"))
    one_dec = i.GetSubstructMatches(Chem.MolFromSmarts("[#6D4:0](@[#6D2:4])(~[#54D1:1])[#6D3:2](~[#6D1:3])"))
    also_one =i.GetSubstructMatches(Chem.MolFromSmarts("[#6D4:0](@[#6D2:4])(~[#6D1:1])[#6D3:2](~[#54D1:3])"))
    two_dec = i.GetSubstructMatches(Chem.MolFromSmarts("[#6D4:0](@[#6D2:4])(~[#54D1:1])[#6D3:2](~[#54D1:3])"))
    if(len(no_dec)==1):
        nothin=nothin+1; continue
    elif(len(one_dec)==1):
        one = one +1; continue
    elif(len(two_dec)==1):
        two = two +1; continue
    elif(len(also_one)==1):
        one=one+1; continue
    
    #ring_decoration (1)
    no_dec = i.GetSubstructMatches(Chem.MolFromSmarts("[#6D4:0](@[#54D2:4])(~[#6D1:1])[#6D3:2](~[#6D1:3])"))
    one_dec = i.GetSubstructMatches(Chem.MolFromSmarts("[#6D4:0](@[#54D2:4])(~[#54D1:1])[#6D3:2](~[#6D1:3])"))
    also_one =i.GetSubstructMatches(Chem.MolFromSmarts("[#6D4:0](@[#54D2:4])(~[#6D1:1])[#6D3:2](~[#54D1:3])"))
    two_dec = i.GetSubstructMatches(Chem.MolFromSmarts("[#6D4:0](@[#54D2:4])(~[#54D1:1])[#6D3:2](~[#54D1:3])"))
    if(len(no_dec)==1):
        nothin=nothin+1; continue
    elif(len(one_dec)==1):
        one = one +1; continue
    elif(len(two_dec)==1):
        two = two +1; continue
    elif(len(also_one)==1):
        one=one+1; continue
    
    #ring_decoration (1)
    no_dec = i.GetSubstructMatches(Chem.MolFromSmarts("[#6D4:0](@[#6D2:4])(~[#6D1:1])[#54D3:2](~[#6D1:3])"))
    one_dec = i.GetSubstructMatches(Chem.MolFromSmarts("[#6D4:0](@[#6D2:4])(~[#54D1:1])[#54D3:2](~[#6D1:3])"))
    also_one =i.GetSubstructMatches(Chem.MolFromSmarts("[#6D4:0](@[#6D2:4])(~[#6D1:1])[#54D3:2](~[#54D1:3])"))
    two_dec = i.GetSubstructMatches(Chem.MolFromSmarts("[#6D4:0](@[#6D2:4])(~[#54D1:1])[#54D3:2](~[#54D1:3])"))
    if(len(no_dec)==1):
        nothin=nothin+1; continue
    elif(len(one_dec)==1):
        one = one +1; continue
    elif(len(two_dec)==1):
        two = two +1; continue
    elif(len(also_one)==1):
        one=one+1; continue
        
    #ring_decoration (2)
    no_dec = i.GetSubstructMatches(Chem.MolFromSmarts("[#6D4:0](@[#54D2:4])(~[#6D1:1])[#54D3:2](~[#6D1:3])"))
    one_dec = i.GetSubstructMatches(Chem.MolFromSmarts("[#6D4:0](@[#54D2:4])(~[#54D1:1])[#54D3:2](~[#6D1:3])"))
    also_one =i.GetSubstructMatches(Chem.MolFromSmarts("[#6D4:0](@[#54D2:4])(~[#6D1:1])[#54D3:2](~[#54D1:3])"))
    two_dec = i.GetSubstructMatches(Chem.MolFromSmarts("[#6D4:0](@[#54D2:4])(~[#54D1:1])[#54D3:2](~[#54D1:3])"))
    if(len(no_dec)==1):
        nothin=nothin+1; continue
    elif(len(one_dec)==1):
        one = one +1; continue
    elif(len(two_dec)==1):
        two = two +1; continue
    elif(len(also_one)==1):
        one=one+1; continue
    break
    
print("zero:\t",nothin)
print("one:\t",one)
print("two:\t",two)
print(nothin+one+two,"\t\t",(one+two)/len(temp))

# %% [code]
#all labdane (non-claradane) moeities
nothin = 0
one = 0
two = 0
print(len(temp))
for i in temp:
    no_dec = i.GetSubstructMatches(Chem.MolFromSmarts("[#6D4:0](-[#6D1:1])-[#6D1:2]"))
    one_dec = i.GetSubstructMatches(Chem.MolFromSmarts("[#6D4:0](-[#54D1:1])-[#6D1:2]"))
    two_dec = i.GetSubstructMatches(Chem.MolFromSmarts("[#6D4:0](-[#54D1:1])-[#54D1:2]"))
    if(len(no_dec)==1):
        nothin=nothin+1
    elif(len(one_dec)==1):
        one = one +1
    elif(len(two_dec)==1):
        two = two +1
print("zero:\t",nothin)
print("one:\t",one)
print("two:\t",two)
print(nothin+one+two,"\t\t",(one+two)/len(temp))

# %% [code]
### RESULTS ###
#0:labdane: 0.37
#1:claradane:0.67
#2:kaurane: 0.51
#4:abietane:0.40
#6:pimirane:0.42
#9:cassane:0.42

# %% [markdown]
## Identify connection points based on SMILE format to convert SMILES to edge/node table

# %% [code]
def id_edge_intensity(match, edge_location_list, specific_location):
    temp = []
    # calculate IQV from bond variation list
    total = len(edge_location_list)
    total_unique = len(np.unique(edge_location_list))
    percent_square =0
    if(total_unique>1):
        for i in range(total_unique):
            catagory = ((np.unique(edge_location_list, return_counts=True)[1][i]))
            percent_square = percent_square +(((catagory/total)*100)**2)
        IQV = ((total_unique)*(10000-percent_square))/(10000*(total_unique-1))
    else:
        IQV = 0
    #print(total, total_unique, percent_square,IQV)
    return (round(IQV,4))


# %% [code]
# calculate edge/node locations and differential frequencies among a SMILEs MSA
atoms = re.findall("[A-Z]",consensus_name2, re.IGNORECASE)
bonds = re.findall("[\W\d]+", consensus_name2 )

print(consensus_name2,"\n")
if(len(atoms)==len(bonds)):
    cycles = len(atoms)
elif(len(atoms)>len(bonds)):
    cycles = len(bonds)

extra = len(atoms)-len(bonds)
    
print("Node1\tEdge_Intensity\tNode2")    
edge_intensity = pd.DataFrame(columns = ['Node1', 'Edge_Intensity',"Node2"])

for i in range(cycles):
    #print("\t\t\t\t\t\t",atoms[i],"\t",i,"\t", bonds[i])
    edge = re.findall("[\W\d]", bonds[i])
    counter = 0; skip=0
    
    edge_consensus = []
    for a in range(len(bonds)):
        edge_consensus2 = []
        for b in alignment_smiles:
            edge_consensus2.append(re.findall("[\W\d]+",b,re.IGNORECASE)[a])
        edge_consensus.append(edge_consensus2)
    

    for j in edge:
        
        if(skip==1):
            skip = 0
        #Rings complete + consensus
        elif(re.search("\d",j)!=None):
            for k in range(i,len(bonds)):
                if(str(j) in bonds[k] and i!=k):
                    consensus_edge_intensity = id_edge_intensity(bonds[k][bonds[k].find(str(j))-1], edge_consensus[k], [bonds[k].find(str(j))-1][0])
                    edge_intensity = pd.concat([edge_intensity, pd.DataFrame([{"Node1":i,"Edge_Intensity":consensus_edge_intensity,"Node2":k}])], ignore_index=True)
                    print(i,"\t",consensus_edge_intensity,"\t",k)
                    break
        elif(re.search("\-\d",bonds[i])!=None and bonds[i].find(j)==0 and counter+1!=len(bonds[i])):
            skip=1;
        elif(re.search("\-\d",bonds[i])!=None and re.search("\d",j)!=None and counter+1!=len(bonds[i])):
            skip=4      
        elif(re.search(r'^[-=#]',j)!=None and (counter+1)==len(edge) and skip!=2 and skip!=3):
            consensus_edge_intensity = id_edge_intensity(bonds[i][len(bonds[i])-1], edge_consensus[i], len(bonds[i])-1)
            edge_intensity = pd.concat([edge_intensity, pd.DataFrame([{"Node1":i,"Edge_Intensity":consensus_edge_intensity,"Node2":i+1}])], ignore_index=True)
            print(i,"\t",consensus_edge_intensity,"\t",i+1)
            

        # Print and match )(
        elif(re.search("\(",j)!=None and skip!=2 and skip!=3):
            counter1 = 0
            for k in range(i,len(bonds)):
                if(re.search("\)",bonds[k])!=None):
                    counter1=counter1-1
                if(re.search("\(",bonds[k])!=None):
                    counter1=counter1+1
                if(counter1==0):
                    consensus_edge_intensity = id_edge_intensity(bonds[k][len(bonds[k])-1], edge_consensus[k], len(bonds[k])-1)
                    edge_intensity = pd.concat([edge_intensity, pd.DataFrame([{"Node1":i,"Edge_Intensity":consensus_edge_intensity,"Node2":k+1}])], ignore_index=True)
                    print(i,"\t",consensus_edge_intensity,"\t",k+1)
                    if(re.search("\d+\(",bonds[i])!=None):
                        consensus_edge_intensity = id_edge_intensity(bonds[i][len(bonds[i])-1], edge_consensus[i], len(bonds[i])-1)
                        #print(i,"\t" ,consensus_edge_intensity,"\t", k, "TEST2")
                        #skip=1
                        break
                    break

        elif(re.search("\)",j)!=None and skip!=3):
            if(re.search("\)\(", bonds[i])!=None):
                counter1 = 1
            else: break
            skip=2
            for k in range(i,-1,-1):
                if(re.search("\(",bonds[k])!=None):
                    counter1=counter1-1
                if(re.search("\)",bonds[k])!=None):
                    counter1=counter1+1
                if(counter1==0):
                    consensus_edge_intensity = id_edge_intensity(bonds[k][len(bonds[k])-1], edge_consensus[k], len(bonds[k])-1)
                    edge_intensity = pd.concat([edge_intensity, pd.DataFrame([{"Node1":k,"Edge_Intensity":consensus_edge_intensity,"Node2":i+1}])], ignore_index=True)
                    print(k,"\t",consensus_edge_intensity,"\t",i+1)
                    skip=3
                    break
            break

        elif(re.search("\)",j)!=None):
            skip=2
        counter=counter+1
    skip=0
# for j in range(extra):
#     print(int(len(bonds))+j,"\t", atoms[int(len(bonds))+j])

    

# NODE INTENSITIES
print("\nNode\tIntensity")
'''EMPTY PD.DATAFRAME HERE WITH ABOVE FOR HEADERS'''
node_intensity = pd.DataFrame(columns = ['Node', 'Intensity'])
for i in range(len(atoms)):
    
    temp = []; percent_square=0; total=0
    for j in alignment_smiles:
        #j=re.sub("[LDUR]","C",j)
        temp.append(re.findall("[A-Z]",j,re.IGNORECASE)[i])
    
    if(len(np.unique(temp))==1):
        IQV_atom = 0
    else:
        for k in range(len(np.unique(temp))):
            if(re.search("C", np.unique(temp, return_counts=True)[0][k], re.IGNORECASE)):
                catagory = (np.unique(temp, return_counts=True)[1][k])*100
                percent_square = percent_square + (catagory/len(temp))**2
                total = total +1
            else:
                percent_square = percent_square + ((np.unique(temp,return_counts=True)[1][k]*100)/len(temp))
                total = total + np.unique(temp,return_counts=True)[1][k]
        IQV_atom = (len(temp)*(10000-percent_square))/(10000*(len(temp)-1))
        #print(IQV_atom)
    node_intensity = pd.concat([node_intensity, pd.DataFrame([{"Node":i,"Intensity":round(IQV_atom,3)}])], ignore_index=True)
    print(i, "\t", round(IQV_atom,3))
    
### Directory names depending on skeleton abundance (indexed at 1; not 0)
directory_name = str(re.sub(r'\\', r'/',str(os.getcwd())))+"/Supp.7.Backbone_IQV/Skeleton_"+str(atom_value+1)+"/"
# Check if the directory already exists
if not os.path.exists(directory_name):
    os.makedirs(directory_name)
    
node_file = directory_name+str(atom_value+1)+"_"+"NODE_IQV.tsv"
edge_file = directory_name+str(atom_value+1)+"_"+"EDGE_IQV.tsv"
MSA_file  = directory_name+str(atom_value+1)+"_"+"MSA.txt"
consensus_file = directory_name+str(atom_value+1)+"_"+"consensus.txt"

'''
WRITE OUTPUTS
'''
# Write Node IQV scores as an output .tsv
node_intensity.to_csv(node_file, sep='\t', index=False)

# Write Edge IQV scores as an output .tsv
edge_intensity.to_csv(edge_file, sep='\t', index=False)

# Write Consensus as an output .txt
with open(consensus_file, 'w') as file:
    file.write(consensus_name2)

# Write MSA as an output .txt
with open(MSA_file, 'w') as file:
    for item in alignment_smiles:
        file.write(str(item) + '\n')


# %% [markdown]
## Positional Variability Analysis Functions

# %% [code]
# FUNCTION FOR QUANTIFYING POSITIONAL VARIANCE
def percent_diversity(atom_number, MSA_library):
    location_diversity = []
    for i in MSA_library:
        location_diversity.append(re.findall("[a-z]", i, re.IGNORECASE)[atom_number])
    variability = location_diversity.count("X")/len(location_diversity)
    return round(variability,3)

# %% [code]
def Primary_Carbon_stats(consensus_mol, MSA_library):
    # [1]
    temp = []
    for i in consensus_mol.GetSubstructMatches(Chem.MolFromSmarts("[#6D1]")):
        print(percent_diversity(i[0], MSA_library))
        temp.append(percent_diversity(i[0], MSA_library))
        
    Possible_Primary_Neighbors = [2,3,4]
    
    for i in Possible_Primary_Neighbors:
        Primary_SMARTs = "[#6D1][#6D"+str(i)+"]"
        message = "1,"+str(i)
        print("\n\t--",message)
        for j in consensus_mol.GetSubstructMatches(Chem.MolFromSmarts(Primary_SMARTs)):
            print("\t", percent_diversity(j[0], MSA_library))
    return temp


def Secondary_Carbon_stats(consensus_mol, MSA_library):
    # [2]
    temp = []
    for i in consensus_mol.GetSubstructMatches(Chem.MolFromSmarts("[#6D2]")):
        print(percent_diversity(i[0], MSA_library))
        temp.append(percent_diversity(i[0], MSA_library))
        
    Possible_Secondary_Neighbors = [[1,2],
                                    [1,3],
                                    [1,4],
                                    [2,2],
                                    [2,3],
                                    [2,4],
                                    [3,3],
                                    [3,4],
                                    [4,4],
                                   ]
    
    for i in Possible_Secondary_Neighbors:
        Secondary_SMARTs = "[#6D2]([#6D"+str(i[0])+"])[#6D"+str(i[1])+"]"
        message = "2,"+str(i[0])+","+str(i[1])
        print("\n\t--",message)
        for j in consensus_mol.GetSubstructMatches(Chem.MolFromSmarts(Secondary_SMARTs)):
            print("\t", percent_diversity(j[0], MSA_library))
    return temp

def Tertiary_Carbon_stats(consensus_mol, MSA_library):
    # [3]
    temp = []
    for i in consensus_mol.GetSubstructMatches(Chem.MolFromSmarts("[#6D3]")):
        print(percent_diversity(i[0], MSA_library))
        temp.append(percent_diversity(i[0], MSA_library))
        
    Possible_Tertiary_Neighbors = [[1,1,2],
                                    [1,1,3],
                                    [1,1,4],
                                    [1,2,2],
                                    [1,2,3],
                                    [1,2,4],
                                    [1,3,3],
                                    [1,3,4],
                                    [1,4,4],
                                    [2,2,2],
                                    [2,2,3],
                                    [2,2,4],
                                    [2,3,3],
                                    [2,3,4],
                                    [2,4,4],
                                    [3,3,3],
                                    [3,3,4],
                                    [3,4,4],
                                    [4,4,4],
                                   ]
    
    for i in Possible_Tertiary_Neighbors:
        Tertiary_SMARTs = "[#6D3]([#6D"+str(i[0])+"])([#6D"+str(i[1])+"])[#6D"+str(i[2])+"]"
        message = "3,"+str(i[0])+","+str(i[1])+","+str(i[2])
        print("\n\t--",message)
        for j in consensus_mol.GetSubstructMatches(Chem.MolFromSmarts(Tertiary_SMARTs)):
            print("\t", percent_diversity(j[0], MSA_library))
    return temp

# %% [markdown]
## Code below submits and generates statistics for the top 20 compound variabilities and groups either by Labdane like or macrocyclic like

# %% [code]
# LOAD IN MSA LIBRARY

# LABDANE BASED BACKBONES 
Skeleton_group = [1,2,3,5,7,10,21]
Labdane_1 = []
Labdane_2 = []
Labdane_3 = []


for Skeleton_ID in Skeleton_group:
    MSA_library = []
    # FOR CYCLING BETWEEN MSA LIBRARIES EASILY
    MSA_file       ="Supp.7.Backbone_IQV/Skeleton_"+str(Skeleton_ID)+"/"+str(Skeleton_ID)+"_MSA.txt"
    consensus_file ="Supp.7.Backbone_IQV/Skeleton_"+str(Skeleton_ID)+"/"+str(Skeleton_ID)+"_consensus.txt"

    # LOAD IN MSA FILE
    with open(MSA_file, 'r') as file:
        for line in file:
            MSA_library.append(line.strip())

    # LOAD IN CONSENSUS 
    with open(consensus_file, "r") as file:
        consensus_name = file.read()
        consensus_mol = Chem.MolFromSmiles(consensus_name)

    # Primary Carbon Info
    print("#############################################################")
    print("######################     ",Skeleton_ID,"     ##########################")
    print("#############################################################")
    print("------------------------- Primary ---------------------------")
    Labdane_1 = Labdane_1+Primary_Carbon_stats(consensus_mol, MSA_library)
    print("------------------------- Secondary -------------------------")
    Labdane_2 = Labdane_2+Secondary_Carbon_stats(consensus_mol, MSA_library)
    print("------------------------- Tertiary --------------------------")
    Labdane_3 = Labdane_3+Tertiary_Carbon_stats(consensus_mol, MSA_library)


# %% [code]
# LOAD IN MSA LIBRARY
    
# MACROCYCLIC BASED BACKBONES
Skeleton_group = [4,6,8,11,12,13,14,15,16,18]
Macro_1 = []
Macro_2 = []
Macro_3 = []

for Skeleton_ID in Skeleton_group:
    MSA_library = []
    # FOR CYCLING BETWEEN MSA LIBRARIES EASILY
    MSA_file       ="Supp.7.Backbone_IQV/Skeleton_"+str(Skeleton_ID)+"/"+str(Skeleton_ID)+"_MSA.txt"
    consensus_file ="Supp.7.Backbone_IQV/Skeleton_"+str(Skeleton_ID)+"/"+str(Skeleton_ID)+"_consensus.txt"

    # LOAD IN MSA FILE
    with open(MSA_file, 'r') as file:
        for line in file:
            MSA_library.append(line.strip())

    # LOAD IN CONSENSUS 
    with open(consensus_file, "r") as file:
        consensus_name = file.read()
        consensus_mol = Chem.MolFromSmiles(consensus_name)

    # Primary Carbon Info
    print("#############################################################")
    print("######################     ",Skeleton_ID,"     ##########################")
    print("#############################################################")
    print("------------------------- Primary ---------------------------")
    Macro_1 = Macro_1+Primary_Carbon_stats(consensus_mol, MSA_library)
    print("------------------------- Secondary -------------------------")
    Macro_2 = Macro_2+Secondary_Carbon_stats(consensus_mol, MSA_library)
    print("------------------------- Tertiary --------------------------")
    Macro_3 = Macro_3+Tertiary_Carbon_stats(consensus_mol, MSA_library)


# %% [code]
from scipy.stats import f_oneway

# Perform one-way ANOVA
f_statistic, p_value = f_oneway(Labdane_1, Labdane_2, Labdane_3, Macro_1, Macro_2, Macro_3)

# Print the results
print("F-statistic:", f_statistic)
print("P-value:", p_value)

# Interpret the results
if p_value < 0.05:
    print("The means are significantly different. Reject the null hypothesis.")
else:
    print("Fail to reject the null hypothesis. There is no significant difference in means.")


# %% [code]
from scipy.stats import f_oneway
from statsmodels.stats.multicomp import pairwise_tukeyhsd

# Perform one-way ANOVA
f_statistic, p_value = f_oneway(Labdane_1, Labdane_2, Labdane_3, Macro_1, Macro_2, Macro_3)

# Print the results
print("F-statistic:", f_statistic)
print("P-value:", p_value)

# Perform Tukey's HSD test for post-hoc analysis
data = Labdane_1 + Labdane_2 + Labdane_3 + Macro_1 + Macro_2 + Macro_3
groups = ['Labdane_1'] * len(Labdane_1) + ['Labdane_2'] * len(Labdane_2) + ['Labdane_3'] * len(Labdane_3) + ['Macro_1'] * len(Macro_1) + ['Macro_2'] * len(Macro_2) + ['Macro_3'] * len(Macro_3)

tukey_results = pairwise_tukeyhsd(data, groups)

# Print the Tukey HSD results
print(tukey_results)


# %% [code]
import matplotlib.pyplot as plt
from scipy.stats import sem
means = [np.mean(Labdane_1), np.mean(Labdane_2), np.mean(Labdane_3), np.mean(Macro_1), np.mean(Macro_2), np.mean(Macro_3)]
errors = [sem(Labdane_1), sem(Labdane_2), sem(Labdane_3), sem(Macro_1), sem(Macro_2), sem(Macro_3)]

# Bar graph with error bars
categories = ['Labdane 1C', 'Labdane 2C', 'Labdane 3C', 'Macro 1C', 'Macro 2C', 'Macro 3C']

plt.bar(categories, means, yerr=errors, capsize=5, color= ["red","red","red","blue","blue","blue"])
plt.title('Bar Graph with Error Bars')
plt.xlabel('Categories')
plt.ylabel('Mean Value')
plt.gcf().set_size_inches(14, 5)
plt.show()

# %% [code]
Labdane_1

# %% [code]
# LABDANE CARBON TRENDS

# %% [code]
# LABDANE TRENDS
Labdane_1_2 = [0.6,0.489,0.245,0.913]
Labdane_1_3 = [0.281,0.098,0.302,0.138,0.641,0.23,0.066, 0.201,0.154,0.194]
Labdane_1_4 = [0.017,0.355,0.028,0.136, 0.375,0.039,0.495,0.248,0.101,0.303,0.153,0.024,0.076,0.352,0.062,0.078,0.415,0.093,0.263,0.0,0.505]

Labdane_2_3_3 = [0.055]
Labdane_2_1_4 = [0.223]

Labdane_2_1_3 = [0.065,0.019,0.012]
Labdane_2_2_2 = [0.098,0.325,0.073,0.054,0.216,0.079,0.055]
Labdane_2_2_3 = [0.113,0.05,0.18,0.215,0.098,0.233,0.209,0.141,0.305,0.248,0.087,0.5,0.387,0.462,0.21,0.173,0.322,0.102,0.668,0.066,0.561,0.464,0.145,0.104]
Labdane_2_2_4 = [0.278,0.047,0.086,0.434,0.49,0.267,0.245,0.191,0.07,0.129,0.422,0.169,0.24,0.292,0.211,0.021,0.284,0.356]
Labdane_2_3_4 = [0.592,0.232,0.276,0.202,0.398]


Labdane_3_1_1_3 = [0.089]
Labdane_3_1_3_3 = [0.096]
Labdane_3_1_2_2 = [0.275,0.072]
Labdane_3_1_2_4 = [0.086,0.139]

Labdane_3_1_2_3 = [0.253,0.173,0.26]
Labdane_3_2_2_3 = [0.091,0.076,0.069,0.173,0.006,0.002,0.017]
Labdane_3_2_3_4 = [0.152,0.015,0.082]
Labdane_3_2_4_4 = [0.007,0.026,0.0,0.039,0.009,0.012,0.0,0.4,0.01,0.007]

from scipy.stats import f_oneway
from statsmodels.stats.multicomp import pairwise_tukeyhsd
# Perform one-way ANOVA
f_statistic, p_value = f_oneway(Labdane_1_2,Labdane_1_3,Labdane_1_4,Labdane_2_1_3,Labdane_2_1_4,Labdane_2_2_2,Labdane_2_2_3,Labdane_2_2_4,Labdane_2_3_3,Labdane_2_3_4,Labdane_3_1_1_3,Labdane_3_1_2_2,Labdane_3_1_2_3,Labdane_3_1_2_4,Labdane_3_1_3_3,Labdane_3_2_2_3,Labdane_3_2_3_4,Labdane_3_2_4_4,)

# Print the results
print("F-statistic:", f_statistic)
print("P-value:", p_value)

# Perform Tukey's HSD test for post-hoc analysis
data = Labdane_1_2 + Labdane_1_3 + Labdane_1_4 + Labdane_2_1_3 + Labdane_2_2_2 + Labdane_2_2_3 + Labdane_2_2_4+Labdane_2_3_4+Labdane_3_1_2_2+Labdane_3_1_2_3+Labdane_3_1_2_4+Labdane_3_2_2_3+Labdane_3_2_3_4+Labdane_3_2_4_4#+Labdane_2_3_3+ Labdane_2_1_4 +Labdane_3_1_1_3+Labdane_3_1_3_3
groups = ["1_2"]* len(Labdane_1_2) + ["1_3"]* len(Labdane_1_3) + ["1_4"]* len(Labdane_1_4) + ["2_1_3"]* len(Labdane_2_1_3)  + ["2_2_2"]* len(Labdane_2_2_2) + ["2_2_3"]* len(Labdane_2_2_3) + ["2_2_4"]* len(Labdane_2_2_4)+["2_3_4"]* len(Labdane_2_3_4)+["3_1_2_2"]* len(Labdane_3_1_2_2)+["3_1_2_3"]* len(Labdane_3_1_2_3)+["3_1_2_4"]* len(Labdane_3_1_2_4)+["3_2_2_3"]* len(Labdane_3_2_2_3)+["3_2_3_4"]* len(Labdane_3_2_3_4)+["3_2_4_4"]* len(Labdane_3_2_4_4)#+["2_3_3"]* len(Labdane_2_3_3)+ ["2_1_4"]* len(Labdane_2_1_4)+["3_1_1_3"]* len(Labdane_3_1_1_3)+["3_1_3_3"]* len(Labdane_3_1_3_3)

data = Labdane_1_2 + Labdane_1_3 + Labdane_1_4 + Labdane_2_1_3 + Labdane_2_2_2 + Labdane_2_2_3 + Labdane_2_2_4+Labdane_2_3_4+Labdane_3_1_2_3+Labdane_3_2_2_3+Labdane_3_2_3_4+Labdane_3_2_4_4#+Labdane_2_3_3+ Labdane_2_1_4 +Labdane_3_1_1_3+Labdane_3_1_3_3
groups = ["1_2"]* len(Labdane_1_2) + ["1_3"]* len(Labdane_1_3) + ["1_4"]* len(Labdane_1_4) + ["2_1_3"]* len(Labdane_2_1_3)  + ["2_2_2"]* len(Labdane_2_2_2) + ["2_2_3"]* len(Labdane_2_2_3) + ["2_2_4"]* len(Labdane_2_2_4)+["2_3_4"]* len(Labdane_2_3_4)+["3_1_2_3"]* len(Labdane_3_1_2_3)+["3_2_2_3"]* len(Labdane_3_2_2_3)+["3_2_3_4"]* len(Labdane_3_2_3_4)+["3_2_4_4"]* len(Labdane_3_2_4_4)#+["2_3_3"]* len(Labdane_2_3_3)+ ["2_1_4"]* len(Labdane_2_1_4)+["3_1_1_3"]* len(Labdane_3_1_1_3)+["3_1_3_3"]* len(Labdane_3_1_3_3)


tukey_results = pairwise_tukeyhsd(data, groups)

# Print the Tukey HSD results
print(tukey_results)

means = [np.mean(Labdane_1_2),     np.mean(Labdane_1_3),     np.mean(Labdane_1_4),0, 
         np.mean(Labdane_2_1_3),   np.mean(Labdane_2_2_2),   np.mean(Labdane_2_2_3),   np.mean(Labdane_2_2_4), np.mean(Labdane_2_3_4), 0,
         np.mean(Labdane_3_1_2_3), np.mean(Labdane_3_2_2_3), np.mean(Labdane_3_2_3_4), np.mean(Labdane_3_2_4_4)]

errors = [sem(Labdane_1_2),     sem(Labdane_1_3),     sem(Labdane_1_4),0,
          sem(Labdane_2_1_3),   sem(Labdane_2_2_2),   sem(Labdane_2_2_3),   sem(Labdane_2_2_4),   sem(Labdane_2_3_4), 0,
          sem(Labdane_3_1_2_3), sem(Labdane_3_2_2_3), sem(Labdane_3_2_3_4), sem(Labdane_3_2_4_4)]

# Bar graph with error bars
categories = ["C1 (2)","C1 (3)","C1 (4)","",
              "C2 (1_3)","C2 (2_2)","C2 (2_3)","C2 (2_4)","C2 (3_4)", " ",
              "C3 (1_2_3)","C3 (2_2_3)","C3 (2_3_4)","C3 (2_4_4)"]

labdane_colors = ["#FFDB91","#FFDB91","#FFDB91","White",
                  "#F5BA41","#F5BA41","#F5BA41","#F5BA41","#F5BA41","White",
                  "#F57041","#F57041","#F57041","#F57041"]

plt.bar(categories, means, yerr=errors, capsize=5, color= labdane_colors )
#plt.title('Labdane Carbon Decoration and Position')
#plt.xlabel('Carbon Position (& Neighbors)')
plt.ylabel('Percent Decoration')
plt.xticks(rotation='vertical')
plt.gcf().set_size_inches(9.66, 2)
plt.ylim(0,1)
#3.22:1

ax = plt.gca()
ax.spines['top'].set_visible(False)
ax.spines['right'].set_visible(False)
plt.show()

# %% [code]
# MACROCYCLIC TRENDS 
Macro_1_3 = [0.096,0.16,0.009,0.406,0.149,0.65,0.001,0.209,0.0,0.243,0.999,0.026,0.0,0.0,0.0,0.956,0.0,0.124,0.026,0.029,0.843,0.013,0.005,0.0,0.005,0.047,0.153,0.006,0.033,0.006,0.074,0.0,0.122,0.03,0.759,0.0,0.0]
Macro_1_4 = [0.021,0.0,0.024,0.0,0.001,0.0,0.0,0.107,0.0,0.048,0.052,0.0,0.213]

Macro_2_2_2 = [0.137,0.149,0.013,0.271,0.194,0.631,0.159,0.004]
Macro_2_2_3 = [0.002,0.117,0.245,0.121,0.218,0.251,0.186,0.265,0.987,0.988,0.931,0.074,0.257,0.673,0.426,0.283,0.908,0.053,0.539,0.396,0.234,0.041,0.145,0.153,0.283,0.134,0.694,0.192,0.312,0.246,0.868,0.17]
Macro_2_2_4 = [0.732,0.947,0.971,0.994,0.06,0.203,0.109]
Macro_2_3_3 = [0.907,0.773,0.93,1.0,0.044,0.991,0.622,0.997,0.093,0.11,0.662,0.0,0.394,0.998,0.007,0.99,0.379, 0.856,0.996,0.0,0.556,0.61,0.231,0.86,0.011,0.989,0.989,0.003]


Macro_3_1_1_3 = [0.328,0.0,0.353,0.068]
Macro_3_1_2_2 = [0.022,0.047,0.038,0.338,0.0,0.013,0.048,0.002,0.017,0.854,0.002,0.126,0.064,0.486,0.848,0.008,0.0]
Macro_3_1_2_3 = [0.556,0.021,0.278,0.0,0.0,0.45]

Macro_3_2_2_3 = [0.598,0.983,0.005,0.768,0.89,0.071,0.998,0.563,0.0,0.0]
Macro_3_2_3_3 = [0.002,0.002,0.0,0.149,0.0,0.0,0.002]
Macro_3_2_3_4 = [0.033,0.0,0.0,0.981,0.0,0.0,0.016,0.21]


from scipy.stats import f_oneway
from statsmodels.stats.multicomp import pairwise_tukeyhsd
# Perform one-way ANOVA
f_statistic, p_value = f_oneway(Macro_1_3,Macro_1_4,Macro_2_2_2,Macro_2_2_3,
                                Macro_2_2_4,Macro_2_3_3,Macro_3_1_1_3,Macro_3_1_2_2,
                                Macro_3_1_2_3,Macro_3_2_2_3,Macro_3_2_3_3,Macro_3_2_3_4)


# Print the results
print("F-statistic:", f_statistic)
print("P-value:", p_value)

# Perform Tukey's HSD test for post-hoc analysis
data =               Macro_1_3+              Macro_1_4+               Macro_2_2_2+               Macro_2_2_3+               Macro_2_2_4+               Macro_2_3_3+                 Macro_3_1_1_3+                Macro_3_1_2_2+                  Macro_3_1_2_3+                 Macro_3_2_2_3+                 Macro_3_2_3_3+                 Macro_3_2_3_4
groups = ["1_3"]*len(Macro_1_3)+ ["1_4"]*len(Macro_1_4)+["2_2_2"]*len(Macro_2_2_2)+["2_2_3"]*len(Macro_2_2_3)+["2_2_4"]*len(Macro_2_2_4)+["2_3_3"]*len(Macro_2_3_3)+["3_1_1_3"]*len(Macro_3_1_1_3)+["3_1_2_2"]*len(Macro_3_1_2_2)+["3_1_2_3"]*len(Macro_3_1_2_3)+["3_2_2_3"]*len(Macro_3_2_2_3)+["3_2_3_3"]*len(Macro_3_2_3_3)+["3_2_3_4"]*len(Macro_3_2_3_4)

data =Macro_1_3+Macro_1_4+ Macro_2_2_2+ Macro_2_2_3+ Macro_2_2_4+ Macro_2_3_3+ Macro_3_1_1_3+ Macro_3_1_2_2+ Macro_3_1_2_3+Macro_3_2_2_3 +Macro_3_2_3_3+Macro_3_2_3_4
groups = ["1_3"]*len(Macro_1_3)+ ["1_4"]*len(Macro_1_4)+["2_2_2"]*len(Macro_2_2_2)+["2_2_3"]*len(Macro_2_2_3)+["2_2_4"]*len(Macro_2_2_4)+["2_3_3"]*len(Macro_2_3_3)+["3_1_1_3"]*len(Macro_3_1_1_3)+["3_1_2_2"]*len(Macro_3_1_2_2)+["3_1_2_3"]*len(Macro_3_1_2_3)+["3_2_2_3"]*len(Macro_3_2_2_3)+["3_2_3_3"]*len(Macro_3_2_3_3)+["3_2_3_4"]*len(Macro_3_2_3_4)


tukey_results = pairwise_tukeyhsd(data, groups)

# Print the Tukey HSD results
print(tukey_results)

#Macro_3_2_2_4 = [0.625,0.989]

means = [np.mean(Macro_1_3),     np.mean(Macro_1_4),0, 
         np.mean(Macro_2_2_2), np.mean(Macro_2_2_3), np.mean(Macro_2_2_4),np.mean(Macro_2_3_3),0, 
         np.mean(Macro_3_1_1_3),np.mean(Macro_3_1_2_2), np.mean(Macro_3_1_2_3), np.mean(Macro_3_2_2_3),np.mean(Macro_3_2_3_3), np.mean(Macro_3_2_3_4)]

errors = [sem(Macro_1_3),     sem(Macro_1_4),0, sem(Macro_2_2_2), sem(Macro_2_2_3), sem(Macro_2_2_4),
         sem(Macro_2_3_3),0, sem(Macro_3_1_1_3),sem(Macro_3_1_2_2), 
         sem(Macro_3_1_2_3), sem(Macro_3_2_2_3),sem(Macro_3_2_3_3), sem(Macro_3_2_3_4)]

# Bar graph with error bars
categories = ["C1 (3)","C1 (4)","",
              "C2 (2_2)","C2 (2_3)","C2 (2_4)","C2 (3_3)", " ",
              "C3 (1_1_3)","C3 (1_2_2)","C3 (1_2_3)","C3 (2_2_3)","C3 (2_3_3)","C3 (2_3_4)"]

Macro_colors = ["#9FCEFF","#9FCEFF","White",
                "#5D95EF","#5D95EF","#5D95EF","#5D95EF","White",
                "#3C63A2","#3C63A2","#3C63A2","#3C63A2","#3C63A2","#3C63A2"]

plt.bar(categories, means, yerr=errors, capsize=3, color= Macro_colors )
#plt.title('Labdane Carbon Decoration and Position')
#plt.xlabel('Carbon Position (& Neighbors)')
plt.xticks(rotation='vertical')
plt.gcf().set_size_inches(9.66, 2)
#3.22:1
plt.tick_params(axis='y', which='both', left=False, right=False, labelleft=False)

plt.ylim(0,1)
ax = plt.gca()
ax.spines['top'].set_visible(False)
ax.spines['right'].set_visible(False)
plt.show()

# %% [code]
data = Labdane_1_2 + Labdane_1_3 + Labdane_1_4 + Labdane_2_1_3 + Labdane_2_2_2 + Labdane_2_2_3 + Labdane_2_2_4+Labdane_2_3_4+Labdane_3_1_2_2+Labdane_3_1_2_3+Labdane_3_1_2_4+Labdane_3_2_2_3+Labdane_3_2_3_4+Labdane_3_2_4_4+Macro_1_3+Macro_1_4+Macro_2_2_2+Macro_2_2_3+Macro_2_2_4+ Macro_2_3_3+ Macro_3_1_1_3+ Macro_3_1_2_2+Macro_3_1_2_3+Macro_3_2_2_3+Macro_3_2_3_3+Macro_3_2_3_4
groups = ["L1_2"]* len(Labdane_1_2) + ["L1_3"]* len(Labdane_1_3) + ["L1_4"]* len(Labdane_1_4) + ["L2_1_3"]* len(Labdane_2_1_3)  + ["L2_2_2"]* len(Labdane_2_2_2) + ["L2_2_3"]* len(Labdane_2_2_3) + ["L2_2_4"]* len(Labdane_2_2_4)+["L2_3_4"]* len(Labdane_2_3_4)+["L3_1_2_2"]* len(Labdane_3_1_2_2)+["L3_1_2_3"]* len(Labdane_3_1_2_3)+["L3_1_2_4"]* len(Labdane_3_1_2_4)+["L3_2_2_3"]* len(Labdane_3_2_2_3)+["L3_2_3_4"]* len(Labdane_3_2_3_4)+["L3_2_4_4"]* len(Labdane_3_2_4_4)+["M1_3"]*len(Macro_1_3)+ ["M1_4"]*len(Macro_1_4)+["M2_2_2"]*len(Macro_2_2_2)+["M2_2_3"]*len(Macro_2_2_3)+["M2_2_4"]*len(Macro_2_2_4)+["M2_3_3"]*len(Macro_2_3_3)+["M3_1_1_3"]*len(Macro_3_1_1_3)+["M3_1_2_2"]*len(Macro_3_1_2_2)+["M3_1_2_3"]*len(Macro_3_1_2_3)+["M3_2_2_3"]*len(Macro_3_2_2_3)+["M3_2_3_3"]*len(Macro_3_2_3_3)+["M3_2_3_4"]*len(Macro_3_2_3_4)

tukey_results = pairwise_tukeyhsd(data, groups)
print(tukey_results)


# %% [code]
data = Labdane_1_2 + Labdane_1_3 + Labdane_1_4 + Labdane_2_1_3 + Labdane_2_2_2 + Labdane_2_2_3 + Labdane_2_2_4+Labdane_2_3_4+Labdane_3_1_2_3+Labdane_3_2_2_3+Labdane_3_2_3_4+Labdane_3_2_4_4+Macro_1_3+Macro_1_4+ Macro_2_2_2+ Macro_2_2_3+ Macro_2_2_4+ Macro_2_3_3+ Macro_3_1_1_3+ Macro_3_1_2_2+ Macro_3_1_2_3+Macro_3_2_2_3 +Macro_3_2_3_3+Macro_3_2_3_4
groups = ["L1_2"]* len(Labdane_1_2) + ["L1_3"]* len(Labdane_1_3) + ["L1_4"]* len(Labdane_1_4) + ["L2_1_3"]* len(Labdane_2_1_3)  + ["L2_2_2"]* len(Labdane_2_2_2) + ["L2_2_3"]* len(Labdane_2_2_3) + ["L2_2_4"]* len(Labdane_2_2_4)+["L2_3_4"]* len(Labdane_2_3_4)+["L3_1_2_3"]* len(Labdane_3_1_2_3)+["L3_2_2_3"]* len(Labdane_3_2_2_3)+["L3_2_3_4"]* len(Labdane_3_2_3_4)+["L3_2_4_4"]* len(Labdane_3_2_4_4)+ ["M1_3"]*len(Macro_1_3)+ ["M1_4"]*len(Macro_1_4)+["M2_2_2"]*len(Macro_2_2_2)+["M2_2_3"]*len(Macro_2_2_3)+["M2_2_4"]*len(Macro_2_2_4)+["M2_3_3"]*len(Macro_2_3_3)+["M3_1_1_3"]*len(Macro_3_1_1_3)+["M3_1_2_2"]*len(Macro_3_1_2_2)+["M3_1_2_3"]*len(Macro_3_1_2_3)+["M3_2_2_3"]*len(Macro_3_2_2_3)+["M3_2_3_3"]*len(Macro_3_2_3_3)+["M3_2_3_4"]*len(Macro_3_2_3_4)
tukey_results = pairwise_tukeyhsd(data, groups, alpha=0.4)
print(tukey_results)

# %% [code]
pairwise_tukeyhsd?

# %% [code]
data2 = Labdane_1_2 + Labdane_1_3 + Labdane_1_4 + Labdane_2_1_3 + Labdane_2_2_2 + Labdane_2_2_3 + Labdane_2_2_4+Labdane_2_3_4+Labdane_3_1_2_2+Labdane_3_1_2_3+Labdane_3_1_2_4+Labdane_3_2_2_3+Labdane_3_2_3_4+Labdane_3_2_4_4+Macro_1_3+Macro_1_4+Macro_2_2_2+Macro_2_2_3+Macro_2_2_4+ Macro_2_3_3+ Macro_3_1_1_3+ Macro_3_1_2_2+Macro_3_1_2_3+Macro_3_2_2_3+Macro_3_2_3_3+Macro_3_2_3_4


# %% [code]
np.mean(data)

# %% [code]
np.mean(Macro_2_2_2+Macro_2_3_3+Macro_2_3_3+Macro_2_2_4)


# %% [code]
np.mean(Macro_2_2_4)

# %% [code]
```
