## Supplementary material for "Rule-Based Deconstruction and Reconstruction of Diterpene Libraries: Categorizing Patterns & Unravelling the Structural Landscape": Zip of all Code used: Supp_Code.6.Phylogenetic_Skeleton_Abundance_Heatmap.ipynb

```
# %% [code]
# IMPORT ALL RELEVENT MODULES
import pandas as pd
import numpy as np
import seaborn as sns
import matplotlib.pyplot as plt
import rdkit
from rdkit import Chem
from rdkit.Chem import Draw

# %% [code]
print("Pandas:",pd.__version__)
print("Numpy:",np.__version__)
print("Seaborn:", sns.__version__)
print("Matplotlib:",plt.matplotlib.__version__)
print("RDKit:", rdkit.__version__)

# %% [markdown]
# Overview of program
## Supplemental data used and generated by this program

# %% [code]
#Supplemental Code 6: PHYLOGENETIC DISTRIBUTION OF THE MOST COMMON DITERPENE SKELETONS IN VIRIDIPLANTAE AND ALGAE

### DATASET(S) INTENDED AS INPUT FOR THIS CODE
# Supp.2e.DNP_Diterpene_Skeleton_Summary.tsv
# Supp.2a.DNP_Diterpene_Mining_v30.2_Updated_Skeleton_Backbone.tsv


### SUPPLEMENTAL DATASETS GENERATED BY THIS FUNCTION 
# Figure 5
# Supp.8a.Family_Compound_Phylogeny_Clustered.png
# Supp.8b.Family_Species_count_V_compound.png
# Supp.8c.Family_Species_count_V_compound_ratio.png


'''
### OVERVIEW OF FUNCTIONALITIY ###
### Davis Mathieu
### Jan 31, 2024

###################################################################
This program functions to take the DNP dataset and analyze the 
physiology information provided. DNP families are converted also 
to phylum and kingdom for further seperation. These lists were 
generated by ChatGPT and manually checked. Family entries and 
compound skeletons were compared and when overlapping (based on 
index) were summed and used to quantify the abundance of compounds
in each class. This list is manually organized by Phylum, as 
diterpenes do not cluster correlating to evolutionary trends. The
dataset is then filtered to only include Plantae, the top 50 diterpene
skeletons, and families with 10 or more compounds total. The top 50
skeletons were manually annotated to literature common name. The 
natural log of compound abundance was taken and allowed to have 
hierarchal clustering. The trends, families, and clusters were 
manually annotated post. 


This code produces a heatmap of skeleton abundance based on phylogeny
'''
x=0

# %% [markdown]
## Conversion from family to Kingdom Dictionary
---
##### compliments to ChatGPT for helping with the ease of this (yes, still manually checked)

# %% [code]
# CONVERSION OF FAMILY DATA TO KINGDOM AND PHYLUM (thank you ChatGPT)
family_to_kingdom = {
    'Acanthaceae': 'Plantae',
    'Achariaceae': 'Plantae',
    'Acoraceae': 'Plantae',
    'Adiantaceae': 'Plantae',
    'Agelasida': 'Animalia',
    'Aizoaceae': 'Plantae',
    'Alangiaceae': 'Plantae',
    'Alismataceae': 'Plantae',
    'Amaranthaceae': 'Plantae',
    'Amphibians': 'Animalia',
    'Anacardiaceae': 'Plantae',
    'Annonaceae': 'Plantae',
    'Apocynaceae': 'Plantae',
    'Aquifoliaceae': 'Plantae',
    'Araceae': 'Plantae',
    'Arachnids': 'Animalia',
    'Araliaceae': 'Plantae',
    'Araucariaceae': 'Plantae',
    'Archaebacteria': 'Archaea',
    'Aristolochiaceae': 'Plantae',
    'Asclepiadaceae': 'Plantae',
    'Ascomycetes': 'Fungi',
    'Asparagaceae': 'Plantae',
    'Avicenniaceae': 'Plantae',
    'Axinellida': 'Animalia',
    'Basidiomycetes': 'Fungi',
    'Berberidaceae': 'Plantae',
    'Bignoniaceae': 'Plantae',
    'Bixaceae': 'Plantae',
    'Boraginaceae': 'Plantae',
    'Bromeliaceae': 'Plantae',
    'Bryozoans': 'Animalia',
    'Bubarida': 'Animalia',
    'Buddlejaceae': 'Plantae',
    'Burseraceae': 'Plantae',
    'Calycanthaceae': 'Plantae',
    'Cannabaceae': 'Plantae',
    'Capparaceae': 'Plantae',
    'Caprifoliaceae': 'Plantae',
    'Caryophyllaceae': 'Plantae',
    'Celastraceae': 'Plantae',
    'Cephalopods': 'Animalia',
    'Cephalotaxaceae': 'Plantae',
    'Chloranthaceae': 'Plantae',
    'Chlorophytes (green algae)': 'Plantae',
    'Chrysobalanaceae': 'Plantae',
    'Cistaceae': 'Plantae',
    'Clionaida': 'Animalia',
    'Cneoraceae': 'Plantae',
    'Coleoptera': 'Animalia',
    'Combretaceae': 'Plantae',
    'Compositae': 'Plantae',
    'Convolvulaceae': 'Plantae',
    'Crassulaceae': 'Plantae',
    'Crinoids': 'Animalia',
    'Cruciferae': 'Plantae',
    'Crustaceans': 'Animalia',
    'Cucurbitaceae': 'Plantae',
    'Cupressaceae': 'Plantae',
    'Cymodoceaceae': 'Plantae',
    'Cyperaceae': 'Plantae',
    'Dendroceratida': 'Animalia',
    'Dennstaedtiaceae': 'Plantae',
    'Deuteromycetes (fungi imperfecti)': 'Fungi',
    'Dichapetalaceae': 'Plantae',
    'Dictyoceratida': 'Animalia',
    'Dilleniaceae': 'Plantae',
    'Dioscoreaceae': 'Plantae',
    'Diptera': 'Animalia',
    'Dipteridaceae': 'Plantae',
    'Dryopteridaceae': 'Plantae',
    'Ebenaceae': 'Plantae',
    'Epacridaceae': 'Plantae',
    'Ericaceae': 'Plantae',
    'Erythroxylaceae': 'Plantae',
    'Euphorbiaceae': 'Plantae',
    'Fagaceae': 'Plantae',
    'Fish': 'Animalia',
    'Flacourtiaceae': 'Plantae',
    'Gastropods; Ophisthobranchs': 'Animalia',
    'Gastropods; Prosobranchs': 'Animalia',
    'Gastropods; Pulmonates': 'Animalia',
    'Gentianaceae': 'Plantae',
    'Geraniaceae': 'Plantae',
    'Gesneriaceae': 'Plantae',
    'Ginkgoaceae': 'Plantae',
    'Gleicheniaceae': 'Plantae',
    'Goodeniaceae': 'Plantae',
    'Gramineae': 'Plantae',
    'Grossulariaceae': 'Plantae',
    'Guttiferae': 'Plantae',
    'Hamamelidaceae': 'Plantae',
    'Haplosclerida': 'Animalia',
    'Hemerocallidaceae': 'Plantae',
    'Hemiptera': 'Animalia',
    'Hernandiaceae': 'Plantae',
    'Hornworts (Anthocerotophyta)': 'Plantae',
    'Hydrocharitaceae': 'Plantae',
    'Hymenophyllaceae': 'Plantae',
    'Hymenoptera': 'Animalia',
    'Icacinaceae': 'Plantae',
    'Illiciaceae': 'Plantae',
    'Iridaceae': 'Plantae',
    'Juncaceae': 'Plantae',
    'Lamiaceae': 'Plantae',
    'Lauraceae': 'Plantae',
    'Lecythidaceae': 'Plantae',
    'Leguminosae': 'Plantae',
    'Lemnaceae': 'Plantae',
    'Lepidoptera': 'Animalia',
    'Lichens': 'Fungi',
    'Liliaceae': 'Plantae',
    'Linaceae': 'Plantae',
    'Liverworts': 'Plantae',
    'Loganiaceae': 'Plantae',
    'Loranthaceae': 'Plantae',
    'Lycopodiaceae': 'Plantae',
    'Lythraceae': 'Plantae',
    'Magnoliaceae': 'Plantae',
    'Malpighiaceae': 'Plantae',
    'Malvaceae': 'Plantae',
    'Mammals': 'Animalia',
    'Meliaceae': 'Plantae',
    'Menispermaceae': 'Plantae',
    'Miscellaneous and unclassified Cnidarians': 'Animalia',
    'Miscellaneous and unclassified sponges': 'Animalia',
    'Miscellaneous and unidentified fungi': 'Fungi',
    'Moraceae': 'Plantae',
    'Mosses': 'Plantae',
    'Musaceae': 'Plantae',
    'Myoporaceae': 'Plantae',
    'Myristicaceae': 'Plantae',
    'Myrtaceae': 'Plantae',
    'Nematodes': 'Animalia',
    'Octocorals': 'Animalia',
    'Olacaceae': 'Plantae',
    'Oleaceae': 'Plantae',
    'Onagraceae': 'Plantae',
    'Oomycetes': 'Fungi',
    'Orchidaceae': 'Plantae',
    'Osmundaceae': 'Plantae',
    'Other insects': 'Animalia',
    'Other non-insect arthropods': 'Animalia',
    'Pedaliaceae': 'Plantae',
    'Phaeophytes (brown algae)': 'Plantae',
    'Phyllocladaceae': 'Plantae',
    'Pinaceae': 'Plantae',
    'Podocarpaceae': 'Plantae',
    'Poecilosclerida': 'Animalia',
    'Polemoniaceae': 'Plantae',
    'Polygonaceae': 'Plantae',
    'Polypodiaceae': 'Plantae',
    'Portulacaceae': 'Plantae',
    'Potamogetonaceae': 'Plantae',
    'Protochordates': 'Animalia',
    'Protozoa': 'Protista',
    'Pteridaceae': 'Plantae',
    'Ranunculaceae': 'Plantae',
    'Reptiles': 'Animalia',
    'Rhamnaceae': 'Plantae',
    'Rhizophoraceae': 'Plantae',
    'Rhodophytes (red algae)': 'Plantae',
    'Rosaceae': 'Plantae',
    'Rubiaceae': 'Plantae',
    'Rutaceae': 'Plantae',
    'Salicaceae': 'Plantae',
    'Salviniaceae': 'Plantae',
    'Sapindaceae': 'Plantae',
    'Saururaceae': 'Plantae',
    'Saxifragaceae': 'Plantae',
    'Schisandraceae': 'Plantae',
    'Scopalinida': 'Animalia',
    'Scrophulariaceae': 'Plantae',
    'Sea cucumbers (Holothuroidea)': 'Animalia',
    'Selaginellaceae': 'Plantae',
    'Simaroubaceae': 'Plantae',
    'Solanaceae': 'Plantae',
    'Strelitziaceae': 'Plantae',
    'Suberitida': 'Animalia',
    'Taxaceae': 'Plantae',
    'Taxodiaceae': 'Plantae',
    'Tetractinellida': 'Animalia',
    'Theaceae': 'Plantae',
    'Thymelaeaceae': 'Plantae',
    'Tiliaceae': 'Plantae',
    'Trigoniaceae': 'Plantae',
    'Umbelliferae': 'Plantae',
    'Unassigned': 'Unknown',
    'Valerianaceae': 'Plantae',
    'Velloziaceae': 'Plantae',
    'Verbenaceae': 'Plantae',
    'Verongiida': 'Animalia',
    'Vitaceae': 'Plantae',
    'Zingiberaceae': 'Plantae',
    'Zygomycetes': 'Fungi',
    'Zygophyllaceae': 'Plantae'
}

family_to_phylum = {
    'Acanthaceae': 'Eudicot',
    'Achariaceae': 'Eudicot',
    'Acoraceae': 'Monocot',
    'Adiantaceae': 'Ferns and Allies',
    'Agelasida': 'Porifera',
    'Aizoaceae': 'Eudicot',
    'Alangiaceae': 'Eudicot',
    'Alismataceae': 'Monocot',
    'Amaranthaceae': 'Eudicot',
    'Amphibians': 'Chordates',
    'Anacardiaceae': 'Eudicot',
    'Annonaceae': 'Eudicot',
    'Apocynaceae': 'Eudicot',
    'Aquifoliaceae': 'Eudicot',
    'Araceae': 'Monocot',
    'Arachnids': 'Arthropods',
    'Araliaceae': 'Eudicot',
    'Araucariaceae': 'Conifers',
    'Archaebacteria': 'Bacteria',
    'Aristolochiaceae': 'Eudicot',
    'Asclepiadaceae': 'Eudicot',
    'Ascomycetes': 'Fungi',
    'Asparagaceae': 'Monocot',
    'Avicenniaceae': 'Eudicot',
    'Axinellida': 'Porifera',
    'Basidiomycetes': 'Fungi',
    'Berberidaceae': 'Eudicot',
    'Bignoniaceae': 'Eudicot',
    'Bixaceae': 'Eudicot',
    'Boraginaceae': 'Eudicot',
    'Bromeliaceae': 'Monocot',
    'Bryozoans': 'Bryozoans',
    'Bubarida': 'Porifera',
    'Buddlejaceae': 'Eudicot',
    'Burseraceae': 'Eudicot',
    'Calycanthaceae': 'Eudicot',
    'Cannabaceae': 'Eudicot',
    'Capparaceae': 'Eudicot',
    'Caprifoliaceae': 'Eudicot',
    'Caryophyllaceae': 'Eudicot',
    'Celastraceae': 'Eudicot',
    'Cephalopods': 'Mollusks',
    'Cephalotaxaceae': 'Conifers',
    'Chloranthaceae': 'Magnoliods',
    'Chlorophytes (green algae)': 'Chlorophyta',
    'Chrysobalanaceae': 'Eudicot',
    'Cistaceae': 'Eudicot',
    'Clionaida': 'Porifera',
    'Cneoraceae': 'Eudicot',
    'Coleoptera': 'Arthropods',
    'Combretaceae': 'Eudicot',
    'Compositae': 'Eudicot',
    'Convolvulaceae': 'Eudicot',
    'Crassulaceae': 'Eudicot',
    'Crinoids': 'Echinoderms',
    'Cruciferae': 'Eudicot',
    'Crustaceans': 'Arthropods',
    'Cucurbitaceae': 'Eudicot',
    'Cupressaceae': 'Conifers',
    'Cymodoceaceae': 'Eudicot',
    'Cyperaceae': 'Monocot',
    'Dendroceratida': 'Porifera',
    'Dennstaedtiaceae': 'Ferns and Allies',
    'Deuteromycetes (fungi imperfecti)': 'Fungi',
    'Dichapetalaceae': 'Eudicot',
    'Dictyoceratida': 'Porifera',
    'Dilleniaceae': 'Eudicot',
    'Dioscoreaceae': 'Monocot',
    'Diptera': 'Arthropods',
    'Dipteridaceae': 'Ferns and Allies',
    'Dryopteridaceae': 'Ferns and Allies',
    'Ebenaceae': 'Eudicot',
    'Epacridaceae': 'Eudicot',
    'Ericaceae': 'Eudicot',
    'Erythroxylaceae': 'Eudicot',
    'Euphorbiaceae': 'Eudicot',
    'Fagaceae': 'Eudicot',
    'Fish': 'Chordates',
    'Flacourtiaceae': 'Eudicot',
    'Gastropods; Ophisthobranchs': 'Mollusks',
    'Gastropods; Prosobranchs': 'Mollusks',
    'Gastropods; Pulmonates': 'Mollusks',
    'Gentianaceae': 'Eudicot',
    'Geraniaceae': 'Eudicot',
    'Gesneriaceae': 'Eudicot',
    'Ginkgoaceae': 'Ginkgoophyta',
    'Gleicheniaceae': 'Ferns and Allies',
    'Goodeniaceae': 'Eudicot',
    'Gramineae': 'Monocot',
    'Grossulariaceae': 'Eudicot',
    'Guttiferae': 'Eudicot',
    'Hamamelidaceae': 'Eudicot',
    'Haplosclerida': 'Porifera',
    'Hemerocallidaceae': 'Eudicot',
    'Hemiptera': 'Arthropods',
    'Hernandiaceae': 'Eudicot',
    'Hornworts (Anthocerotophyta)': 'Bryophytes',
    'Hydrocharitaceae': 'Angiosperms',
    'Hymenophyllaceae': 'Ferns and Allies',
    'Hymenoptera': 'Arthropods',
    'Icacinaceae': 'Eudicot',
    'Illiciaceae': 'Eudicot',
    'Iridaceae': 'Monocot',
    'Juncaceae': 'Monocot',
    'Lamiaceae': 'Eudicot',
    'Lauraceae': 'Eudicot',
    'Lecythidaceae': 'Eudicot',
    'Leguminosae': 'Eudicot',
    'Lemnaceae': 'Monocot',
    'Lepidoptera': 'Arthropods',
    'Lichens': 'Fungi',
    'Liliaceae': 'Monocot',
    'Linaceae': 'Eudicot',
    'Liverworts': 'Bryophytes',
    'Loganiaceae': 'Eudicot',
    'Loranthaceae': 'Eudicot',
    'Lycopodiaceae': 'Lycophytes',
    'Lythraceae': 'Eudicot',
    'Magnoliaceae': 'Magnoliods',
    'Malpighiaceae': 'Eudicot',
    'Malvaceae': 'Eudicot',
    'Mammals': 'Chordates',
    'Meliaceae': 'Eudicot',
    'Menispermaceae': 'Eudicot',
    'Miscellaneous and unclassified Cnidarians': 'Cnidarians',
    'Miscellaneous and unclassified sponges': 'Porifera',
    'Miscellaneous and unidentified fungi': 'Fungi',
    'Moraceae': 'Eudicot',
    'Mosses': 'Bryophytes',
    'Musaceae': 'Eudicot',
    'Myoporaceae': 'Eudicot',
    'Myristicaceae': 'Eudicot',
    'Myrtaceae': 'Eudicot',
    'Nematodes': 'Nematoda',
    'Octocorals': 'Cnidarians',
    'Olacaceae': 'Eudicot',
    'Oleaceae': 'Eudicot',
    'Onagraceae': 'Eudicot',
    'Oomycetes': 'Chromista',
    'Orchidaceae': 'Monocot',
    'Osmundaceae': 'Ferns and Allies',
    'Other insects': 'Arthropods',
    'Other non-insect arthropods': 'Arthropods',
    'Pedaliaceae': 'Eudicot',
    'Phaeophytes (brown algae)': 'Stramenopiles',
    'Phyllocladaceae': 'Gnetophyta',
    'Pinaceae': 'Conifers',
    'Podocarpaceae': 'Conifers',
    'Poecilosclerida': 'Porifera',
    'Polemoniaceae': 'Eudicot',
    'Polygonaceae': 'Eudicot',
    'Polypodiaceae': 'Ferns and Allies',
    'Portulacaceae': 'Eudicot',
    'Potamogetonaceae': 'Eudicot',
    'Protochordates': 'Chordates',
    'Protozoa': 'Protists',
    'Pteridaceae': 'Ferns and Allies',
    'Ranunculaceae': 'Eudicot',
    'Reptiles': 'Chordates',
    'Rhamnaceae': 'Eudicot',
    'Rhizophoraceae': 'Eudicot',
    'Rhodophytes (red algae)': 'Rhodophyta',
    'Rosaceae': 'Eudicot',
    'Rubiaceae': 'Eudicot',
    'Rutaceae': 'Eudicot',
    'Salicaceae': 'Eudicot',
    'Salviniaceae': 'Ferns and Allies',
    'Sapindaceae': 'Eudicot',
    'Saururaceae': 'Eudicot',
    'Saxifragaceae': 'Eudicot',
    'Schisandraceae': 'Eudicot',
    'Scopalinida': 'Porifera',
    'Scrophulariaceae': 'Eudicot',
    'Sea cucumbers (Holothuroidea)': 'Echinoderms',
    'Selaginellaceae': 'Lycophytes',
    'Simaroubaceae': 'Eudicot',
    'Solanaceae': 'Eudicot',
    'Strelitziaceae': 'Eudicot',
    'Suberitida': 'Porifera',
    'Taxaceae': 'Conifers',
    'Taxodiaceae': 'Conifers',
    'Tetractinellida': 'Porifera',
    'Theaceae': 'Eudicot',
    'Thymelaeaceae': 'Eudicot',
    'Tiliaceae': 'Eudicot',
    'Trigoniaceae': 'Eudicot',
    'Umbelliferae': 'Eudicot',
    'Unassigned': 'Unknown',
    'Valerianaceae': 'Eudicot',
    'Velloziaceae': 'Eudicot',
    'Verbenaceae': 'Eudicot',
    'Verongiida': 'Porifera',
    'Vitaceae': 'Eudicot',
    'Zingiberaceae': 'Eudicot',
    'Zygomycetes': 'Fungi',
    'Zygophyllaceae': 'Eudicot'
}

# %% [code]
# IMPORT RELEVANT DATA FOR CREATING PHYLOGENY
Skeleton_Summary = pd.read_csv("Supp.2e.DNP_Diterpene_Skeleton_Summary.tsv",sep="\t")
DNP_updated = pd.read_csv("Supp.2a.DNP_Diterpene_Mining_v30.2_Updated_Skeleton_Backbone.tsv",sep="\t")

# %% [code]
# percent of backbones represented in the first 50 compounds [0:49]
print("Total number of diterpenes represented by the top 50 skeletons:")
print(round(sum(list(Skeleton_Summary["frequency"][Skeleton_Summary["Carbon Number"]==20][0:49]))/sum(list(Skeleton_Summary["frequency"][Skeleton_Summary["Carbon Number"]==20]))*100,2),"%")

# %% [code]
# MAKE A DATAFRAME CONTAINING THE NUMBER OF COMPOUNDS REPORTED FOR EACH PHYLOGENETIC FAMILY 
family_data = pd.DataFrame()
for i in np.unique(list(DNP_updated["Smaller_Clade"])):
    temp = pd.DataFrame({"Frequency":np.shape(DNP_updated[DNP_updated["Smaller_Clade"]==i])[0], 
                         "Family":i, 
                         "index":[list(DNP_updated[DNP_updated["Smaller_Clade"]==i].index)]})
    family_data = pd.concat([family_data, temp])
family_data = family_data[family_data['Frequency']!= 0]
family_data = family_data.reset_index()
family_data.drop('level_0', axis=1, inplace=True)
#family_data

# MAKE A DATAFRAME CONTAINING THE NUMBER OF SKELETONS AND THEIR INDEX FOR EACH SKELETON TYPE
Skeleton_Summary.drop('Carbon Number', axis=1, inplace=True)
#Skeleton_Summary

# %% [code]
# CREATE A NEW DATAFRAME TO TAKE THE SUM OF COMPOUNDS WITH EACH SKELETON AND
# QUANTIFY THEM FOR EACH FAMILY
# create empty dataframe with column headers to include kingdom, phylum, family, and numbers 0-50
column_headers = ["Kingdom ID"]+["Phylum ID"]+["Family ID"] + list(range(len(Skeleton_Summary["frequency"])))
skeleton_per_family = pd.DataFrame(columns=column_headers)
counter = 0

# loop through each family and sum the overlap in index for each skeleton position
for i in family_data["index"]:
    i = list(i)
    abundance = []
    for j in Skeleton_Summary["index"]:
        j = list(eval(j))
        temp = len(list(set(i).intersection(set(j))))
        # raw counts
        abundance.append(temp)
        # corrected for frequency
        # abundance.append(temp/len(j))
    row =[family_to_kingdom[family_data["Family"].iloc[counter]]]+[family_to_phylum[family_data["Family"].iloc[counter]]]+[family_data['Family'].iloc[counter]]+abundance
    skeleton_per_family.loc[len(skeleton_per_family["Family ID"])] = row
    
    counter+=1

# dataframe containing the abundance of each skeleton for each phylogeny
skeleton_per_family.head()

# %% [code]
print("All diterpene dataframe shape:",np.shape(skeleton_per_family))
print("Plant diterpene dataframe shape:",np.shape(skeleton_per_family[skeleton_per_family["Kingdom ID"]=="Plantae"]))

# %% [code]
numeric_columns = Skeleton_Summary.select_dtypes(include=np.number)
total_sum_numeric = numeric_columns.sum().sum()
print("Total overall:", total_sum_numeric)
numeric_columns = skeleton_per_family.select_dtypes(include=np.number)
total_sum_numeric = numeric_columns.sum().sum()
print("Total affiliated with a phyla:",total_sum_numeric)
numeric_columns = skeleton_per_family[skeleton_per_family["Kingdom ID"]=="Plantae"].select_dtypes(include=np.number)
total_sum_numeric = numeric_columns.sum().sum()
print("Total affiliated with plants:", total_sum_numeric)

# %% [code]
# ORGANIZE PHYLUMS BASED ON EVOLUTIONARY DIVERGENCE
skeleton_per_family = skeleton_per_family.sort_values(by="Family ID")
skeleton_per_family = skeleton_per_family[skeleton_per_family["Kingdom ID"]=="Plantae"]
print(np.unique(list(skeleton_per_family["Phylum ID"])))

categories=['Stramenopiles','Rhodophyta', 'Chlorophyta', 'Bryophytes', 'Ferns and Allies',
            'Lycophytes', 'Ginkgoophyta', 'Gnetophyta', 'Conifers', 'Magnoliods', 'Monocot', 'Eudicot']

Family_sorted = skeleton_per_family[skeleton_per_family['Phylum ID'].isin(categories)].sort_values(by='Phylum ID', key=lambda x: x.map({val: i for i, val in enumerate(categories)}))
#np.unique(list(Family_sorted["Family ID"]))
Family_sorted

# %% [code]
# CREATE A NEW DATAFRAME TO ONLY CONTAIN THE TOP 50 SKELETONS AND REMOVE
# ALL FAMILIES WITH LESS THAN 10 SKELETONS

Abridged = pd.DataFrame(columns=column_headers); counter = 0
for i in range(len(Family_sorted["Phylum ID"])):
    if(sum(list(Family_sorted.iloc[i,3:52]))>=10):
        Abridged = Abridged.append(Family_sorted.iloc[counter], ignore_index=True)
    counter+=1
Abridged.set_index('Family ID', inplace=True)   
Abridged

# %% [code]
print("Euphorbia:",sum(Abridged.iloc[61,2:52]))
print("Lamiaceae:",sum(Abridged.iloc[56,2:52]))
print("Asteraceae:",sum(Abridged.iloc[55,2:52]))

# %% [code]
Clade0 = Abridged[Abridged["Phylum ID"]!="Eudicot"]
Clade0 = Clade0[Clade0["Phylum ID"]!="Monocot"]
Clade0 = Clade0[Clade0["Phylum ID"]!="Magnoliods"]
Clade0 = Clade0[Clade0["Phylum ID"]!="Conifers"]


Clade1 = Abridged[Abridged["Phylum ID"]=="Conifers"]

Clade2 = Abridged[Abridged["Phylum ID"]=="Monocot"]

Clade3 = Abridged[Abridged["Phylum ID"]=="Eudicot"]

Gymnosperm_order = [
"Podocarpaceae",
"Araucariaceae",
"Taxodiaceae","Cephalotaxaceae","Cupressaceae",
"Taxaceae",
"Pinaceae"]

Monocot_order = [
"Alismataceae",
"Liliaceae","Dioscoreaceae","Orchidaceae","Gramineae"
]

Eudicot_order = ["Illiciaceae",
"Potamogetonaceae",
"Velloziaceae","Zingiberaceae",
"Annonaceae","Aristolochiaceae",
"Ranunculaceae","Menispermaceae",
"Olacaceae",
"Euphorbiaceae","Linaceae","Erythroxylaceae","Chrysobalanaceae","Rhizophoraceae","Achariaceae","Flacourtiaceae",
"Rhamnaceae","Rosaceae",
"Cucurbitaceae",
"Leguminosae",
"Celastraceae",
"Zygophyllaceae",
"Capparaceae",
"Burseraceae","Rutaceae","Sapindaceae","Meliaceae",
"Thymelaeaceae","Cistaceae",
"Ericaceae",
"Icacinaceae",
"Umbelliferae","Araliaceae",
"Caprifoliaceae",
"Compositae",
"Rubiaceae","Loganiaceae",
"Solanaceae","Convolvulaceae",
"Lamiaceae","Oleaceae","Myoporaceae","Buddlejaceae","Acanthaceae","Scrophulariaceae",
"Portulacaceae"]


Clade1 = pd.DataFrame(Clade1, index=Gymnosperm_order)
Clade2 = pd.DataFrame(Clade2, index=Monocot_order)
Clade3 = pd.DataFrame(Clade3, index=Eudicot_order)

result = pd.concat([Clade0, Clade1], ignore_index=False)
result = pd.concat([result, Abridged[Abridged["Phylum ID"]=="Magnoliods"]], ignore_index=False)
result = pd.concat([result, Clade2], ignore_index=False)
Abridged = pd.concat([result, Clade3], ignore_index=False)

# %% [code]
# Function for annotating TaxID's for NCBI phylogeny
# Not necessary, but used here to manually organize
import requests

plant_families = list(Clade1.index)
plant_families = list(Clade2.index)
plant_families = list(Clade3.index)


def get_taxid(family):
    base_url = 'https://eutils.ncbi.nlm.nih.gov/entrez/eutils/'
    search_url = f'{base_url}esearch.fcgi?db=taxonomy&term={family}&retmode=json'
    
    try:
        response = requests.get(search_url)
        data = response.json()
        taxid = data['esearchresult']['idlist'][0]
        return taxid
    except Exception as e:
        print(f"Error retrieving TaxID for {family}: {e}")
        return None

# Get TaxIDs for each family
taxids = {family: get_taxid(family) for family in plant_families}

# Print the results
for family, taxid in taxids.items():
    print(f"{family}: {taxid}")

# %% [code]
# CORRELATE THE VARIOUS PLANT PHYLUMS WITH A COLOR BLIND FRIENDLY COLOR PALETTE
# note: This is used for the right side of the y-axis
row_colors = Abridged["Phylum ID"].map({    'Eudicot':'#7D2284',
                                             'Monocot':"#B446BB",
                                             'Magnoliods':'#F560FF',
                                             'Arthropods':'black', 
                                             'Bacteria':'black', 
                                             'Bryophytes':'#158705', 
                                             'Bryozoans':'black',
                                             'Chlorophyta':'#0F6A02', 
                                             'Chordates':'black', 
                                             'Chromista':'black', 
                                             'Cnidarians':'black', 
                                             'Conifers':'#62F04F',
                                             'Echinoderms':'black', 
                                             'Ferns and Allies':'#17AA03', 
                                             'Fungi':'orange', 
                                             'Ginkgoophyta':'#62F04F',
                                             'Gnetophyta':'#62F04F', 
                                             'Lycophytes':'#17AA03', 
                                             'Mollusks':'black', 
                                             'Nematoda':'black', 
                                             'Porifera':'black',
                                             'Protists':'black', 
                                             'Rhodophyta': '#FF3030',#'#1c381c', 
                                             'Stramenopiles':'#909090', 
                                             'Unknown':'black'})
#np.unique(list(family_matrix["Phylum ID"]))

# %% [code]
# Subset the dataframe for graphing 
To_graph = Abridged.iloc[:,2:52].astype(np.int64)
# Take the natural log of total compounds so graph is not dwarfed by Euphorbia and Lamiaceae
To_graph = np.log(To_graph+1)

# %% [code]
# MANUALLY IDENTIFY AND LABEL ALL SKELETONS REPORTED IN THE DNP AND IN THE ORDER THEY APPEAR
diterpene_skeleton_titles = ["Clerodanes","Labdanes", "Kauranes", "Abietanes", "Cembranes",
            "Pimaranes","Briaranes","Cassanes","Phytanes (GGDP)","Taxanes",
            "Jatrophanes","Eunicellanes","Lathyranes","Tiglianes","Neodolabellanes",
            "Daphnanes","Grayananes","Enmeins","Xenicanes","Isocopalanes",
            "Halimanes","Bifloranes","Ingenanes","Cyathanes","Abeotaxanes",
            "Atisanes","Beyeranes","Dolabellanes","Rosanes","Verticillanes",
            "Cyclophytanes","Ryanodines","Myrsinane","Trachylobanes","Amphilectanes",
            "Triptolides","Pachydictyanes","Erythoxalanes","Jatrophanes","Casbanes",
            "Icetexanes","Cleistanthanes","Bis-Abeo-Abietanes","Guanacastanes","Dolastanes",
            "Seco-Abietanes","Abeo-Abietanes","Totaranes","Tricalysiosades","Prenylbisabolanes"
           ]
To_graph.columns = diterpene_skeleton_titles


# %% [code]
# To_graph
# Skeleton_Summary
# count = 0
# for name, df in To_graph.items():
    
#     print(name,"\t",Skeleton_Summary["SMILES"].iloc[count],"\t",Skeleton_Summary["frequency"].iloc[count])
#     count+=1


# %% [markdown]
## Create a heatmap of top 50 most common skeleton structures, organized by phylogeny and heirarchally* clustered based on skeleton appearance

# %% [code]
#plt.figure(figsize=(20, 6))  # Set the size of the heatmap
#plt.figure().set_figwidth(15)
cluster = sns.clustermap(To_graph, cmap='BuGn', row_cluster=False, col_cluster=True, dendrogram_ratio=0.05,
                         row_colors=row_colors, yticklabels=True, xticklabels=True, cbar_pos=(-0.03, 0.4, 0.03, 0.2))
#cbar_pos=(0.8, 0.02, 0.05, 0.5)
#sns.clustermap(df_sorted.iloc[:,2:], cmap='Oranges', row_cluster=False, col_cluster=True, row_colors=row_colors, yticklabels=True, xticklabels=False)#, cbar_pos=(0.05, .2, .03, .4))
ax = cluster.ax_heatmap
# ax.yaxis.tick_left()

### HORIZONTAL LINES TO SEPERATE PHYLOGENY
# Phylogeny
ax.axhline(y=1, color = "gray", linewidth=1)
ax.axhline(y=2, color = "gray", linewidth=1)
ax.axhline(y=3, color = "gray", linewidth=1)
ax.axhline(y=5, color = "gray", linewidth=1)
ax.axhline(y=8, color = "gray", linewidth=1)
ax.axhline(y=15, color = "gray", linewidth=1)


#euphorbia
ax.axhline(y=30, color = "gray", linewidth=1)
ax.axhline(y=31, color = "gray", linewidth=1)

#composite
ax.axhline(y=55, color = "gray", linewidth=1)
ax.axhline(y=56, color = "gray", linewidth=1)
#lameaceae

ax.axhline(y=60, color = "gray", linewidth=1)
ax.axhline(y=61, color = "gray", linewidth=1)


### VERTICAL LINES FOR INDICATING VARIOUS COMPOUND CLUSTERS
#GGDP, Pimarane, cembrene (common precursors for many other diterpene skeletons)
#ax.axvline(x=3, color='black', linestyle='-', linewidth=1.5)

#pimarane derived
ax.axvline(x=5, color='black', linestyle='-', linewidth=1.5)
ax.axvline(x=8, color='black', linestyle='-', linewidth=1.5)

#Casbene/Cembrene derived
ax.axvline(x=11, color='black', linestyle='-', linewidth=1.5)
ax.axvline(x=19,color='black', linestyle='-', linewidth=1.5)

#Taxane and Taxane like derivatives
ax.axvline(x=24, color='black', linestyle='-', linewidth=1.5)
ax.axvline(x=21, color='black', linestyle='-', linewidth=1.5)

# Fungal and Marine Diterpenes
ax.axvline(x=29, color='black', linestyle='-', linewidth=1.5)

# Abietane derived
ax.axvline(x=39, color='black', linestyle='-', linewidth=1.5)

# Labdane-like
ax.axvline(x=45, color='black', linestyle='-', linewidth=1.5)


# cmap: color map for the heatmap
# annot: annotate each cell with its value
# fmt: format for the annotation (e.g., '.2f' for 2 decimal places)
# linewidths: set line width between cells

# ax.ax_heatmap.set(xticklabels=[])
# ax.ax_col_dendrogram.set(visible=False)

#plt.title('Heatmap Example')  # Set a title for the heatmap
plt.gcf().set_size_inches(13, 14)
plt.show()

# %% [code]
#plt.figure(figsize=(20, 6))  # Set the size of the heatmap
#plt.figure().set_figwidth(15)
cluster = sns.clustermap(To_graph, cmap='BuGn', row_cluster=True, col_cluster=True, dendrogram_ratio=0.05,
                         row_colors=row_colors, yticklabels=True, xticklabels=True, cbar_pos=(-0.03, 0.4, 0.03, 0.2))
#cbar_pos=(0.8, 0.02, 0.05, 0.5)
#sns.clustermap(df_sorted.iloc[:,2:], cmap='Oranges', row_cluster=False, col_cluster=True, row_colors=row_colors, yticklabels=True, xticklabels=False)#, cbar_pos=(0.05, .2, .03, .4))
ax = cluster.ax_heatmap
# ax.yaxis.tick_left()

### HORIZONTAL LINES TO SEPERATE PHYLOGENY
# Phylogeny
ax.axhline(y=1, color = "gray", linewidth=1)
ax.axhline(y=2, color = "gray", linewidth=1)
ax.axhline(y=3, color = "gray", linewidth=1)
ax.axhline(y=5, color = "gray", linewidth=1)
ax.axhline(y=8, color = "gray", linewidth=1)
ax.axhline(y=15, color = "gray", linewidth=1)

#composite
ax.axhline(y=55, color = "gray", linewidth=1)
ax.axhline(y=56, color = "gray", linewidth=1)
#lameaceae
ax.axhline(y=57, color = "gray", linewidth=1)
#euphorbia
ax.axhline(y=61, color = "gray", linewidth=1)
ax.axhline(y=62, color = "gray", linewidth=1)


### VERTICAL LINES FOR INDICATING VARIOUS COMPOUND CLUSTERS
#GGDP, Pimarane, cembrene (common precursors for many other diterpene skeletons)
#ax.axvline(x=3, color='black', linestyle='-', linewidth=1.5)

#pimarane derived
ax.axvline(x=5, color='black', linestyle='-', linewidth=1.5)
ax.axvline(x=8, color='black', linestyle='-', linewidth=1.5)

#Casbene/Cembrene derived
ax.axvline(x=11, color='black', linestyle='-', linewidth=1.5)
ax.axvline(x=19,color='black', linestyle='-', linewidth=1.5)

#Taxane and Taxane like derivatives
ax.axvline(x=24, color='black', linestyle='-', linewidth=1.5)
ax.axvline(x=21, color='black', linestyle='-', linewidth=1.5)

# Fungal and Marine Diterpenes
ax.axvline(x=29, color='black', linestyle='-', linewidth=1.5)

# Abietane derived
ax.axvline(x=39, color='black', linestyle='-', linewidth=1.5)

# Labdane-like
ax.axvline(x=45, color='black', linestyle='-', linewidth=1.5)


# cmap: color map for the heatmap
# annot: annotate each cell with its value
# fmt: format for the annotation (e.g., '.2f' for 2 decimal places)
# linewidths: set line width between cells

# ax.ax_heatmap.set(xticklabels=[])
# ax.ax_col_dendrogram.set(visible=False)

#plt.title('Heatmap Example')  # Set a title for the heatmap
plt.gcf().set_size_inches(13, 14)
plt.show()
cluster.savefig("Supp.XXa.Family_Compound_Phylogeny_Clustered.png", dpi='figure', bbox_inches="tight")


# %% [code]
# SKELETON APPEARANCE IN THE CLUSTERED GRAPH; USED FOR MANUALLY NAMING 
temp = [8,7,4,14,20,33,25,26,32,37,16,39,10,31,22,38,15,12,13,28,41,9,24,30,47,27,48,49,23,34,19,43,6,11,21,36,18,44,35,40,17,45,42,46,0,2,1,3,5]
#temp = [48]

from rdkit import Chem
from rdkit.Chem import Draw

img = []
j=0
for i in Skeleton_Summary["SMILES"].iloc[temp]:
    if(i!="Unknown"):
        print(temp[j],"\t",i)
        img.append(Chem.MolFromSmiles(i))
    j+=1
Draw.MolsToGridImage(img, molsPerRow=1,)

# %% [code]
# REFERENCE SPECIFIC COMPOUNDS WITH A SPECIFIC SKELETON (FOR IDENTIFYING 
# FORMAL CHARACTERIZATION OF THOSE CARBON STRUCTURES)
DNP_updated[DNP_updated["Skeleton"]=="CCC1CCCC2(C)C1CCC13CC(C)C(CCC12)C3"].head(10)

# %% [code]
quant = [4150,155,1,1100,1800,24,600,2500,860,2300,
 4555,400,3600,1500,40,600,2000,2900,8,700,
 860,25,2000,3400,125,860,3,170,860,860,
 2200,1500,5,80,6829,500,170,6,600,23000,
 1400,1450,3700,965,140,17,5500,260,170,500,845,
 12,1700,768,3000,4000,250,7800,900,
 1500,1700,860,3200,1,150,400,12075,150,1400,80,105,80,300,120,600,80,42,
 66,2000,7200,2500,3500, 19500,38,600,250,9500,400,1000,1200,620,330,1300,4225, 600,520,1100,13500,91,190,150,5600,60,600,650,25000,15,
 1750,1750,5,250,200,300,1200,1200,500,100,950,1000,900,
 150,5200,3000,13150,1500,350,10,1800,10,1500,90,5000,
 700,170,2800,10,30,20,600,900,430,25,3700,315,250,3100,
 800,1300,285]

# %% [code]
new=[]
linear_regression = pd.DataFrame(columns=["Family","Compounds","Species"])
for i in range(len(skeleton_per_family["Family ID"])):
    new.append(skeleton_per_family["Family ID"].iloc[i])
    print(skeleton_per_family["Family ID"].iloc[i], "\t",quant[i])#"\t", sum(skeleton_per_family.iloc[i,3:]),"\t",quant[i])
    row = [skeleton_per_family["Family ID"].iloc[i], sum(skeleton_per_family.iloc[i,3:]),quant[i]]
    linear_regression.loc[len(linear_regression["Family"])] = row

# %% [code]
linear_regression
x_values=list(linear_regression["Species"])
y_values=list(linear_regression["Compounds"])
labels=list(linear_regression["Family"])
plt.scatter(x_values,y_values, alpha=0.5)
plt.xlabel("Species Count per Family")
plt.ylabel("Reported Diterpenes")

plt.xlim(-300,27000)
plt.ylim(-50,4500)
# for label, x, y in zip(labels, x_values, y_values):
#     plt.annotate(label, (x, y), textcoords="offset points", xytext=(0,3), ha='center', fontsize=6)

# plt.savefig("Supp.8.Family_Species_count_V_compound.png", dpi='figure', bbox_inches="tight")


# %% [code]
correlation_coefficient = linear_regression['Compounds'].corr(linear_regression['Species'])
print("P-value for correlation between species number and compound:",correlation_coefficient)

# %% [code]
#Ratio of compounds reported normalized to how many species are present in said species
linear_regression
x_values=list(linear_regression["Species"])
y_values=list(linear_regression["Compounds"]/linear_regression["Species"])
labels=list(linear_regression["Family"])
plt.scatter(x_values,y_values)
for label, x, y in zip(labels, x_values, y_values):
    plt.annotate(label, (x, y), textcoords="offset points", xytext=(0,1), ha='center')
plt.savefig("Supp.XXc.Family_Species_count_V_compound_ratio.png", dpi='figure', bbox_inches="tight")


# %% [code]
for i in range(len(linear_regression["Species"])):
    print(linear_regression["Family"].iloc[i],"\t", linear_regression["Compounds"].iloc[i]/linear_regression["Species"].iloc[i])

# %% [code]


# %% [code]
```
