## Supplementary material for "Rule-Based Deconstruction and Reconstruction of Diterpene Libraries: Categorizing Patterns & Unravelling the Structural Landscape": Zip of all data and Metadata: Supp.4.Pickaxe_Carbocation_Source.pptx

### Slide 1
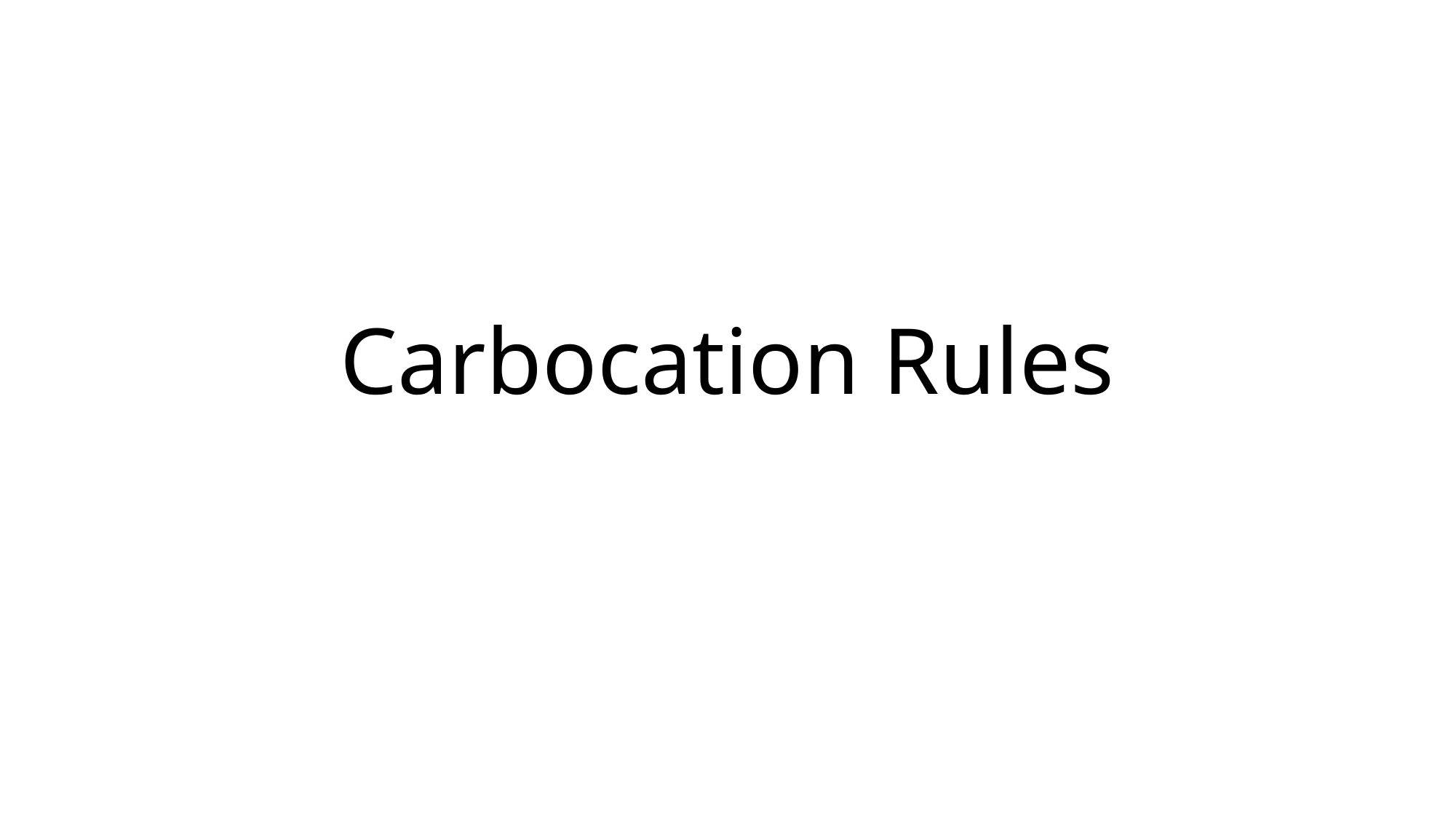

#### Carbocation Rules

### Slide 2
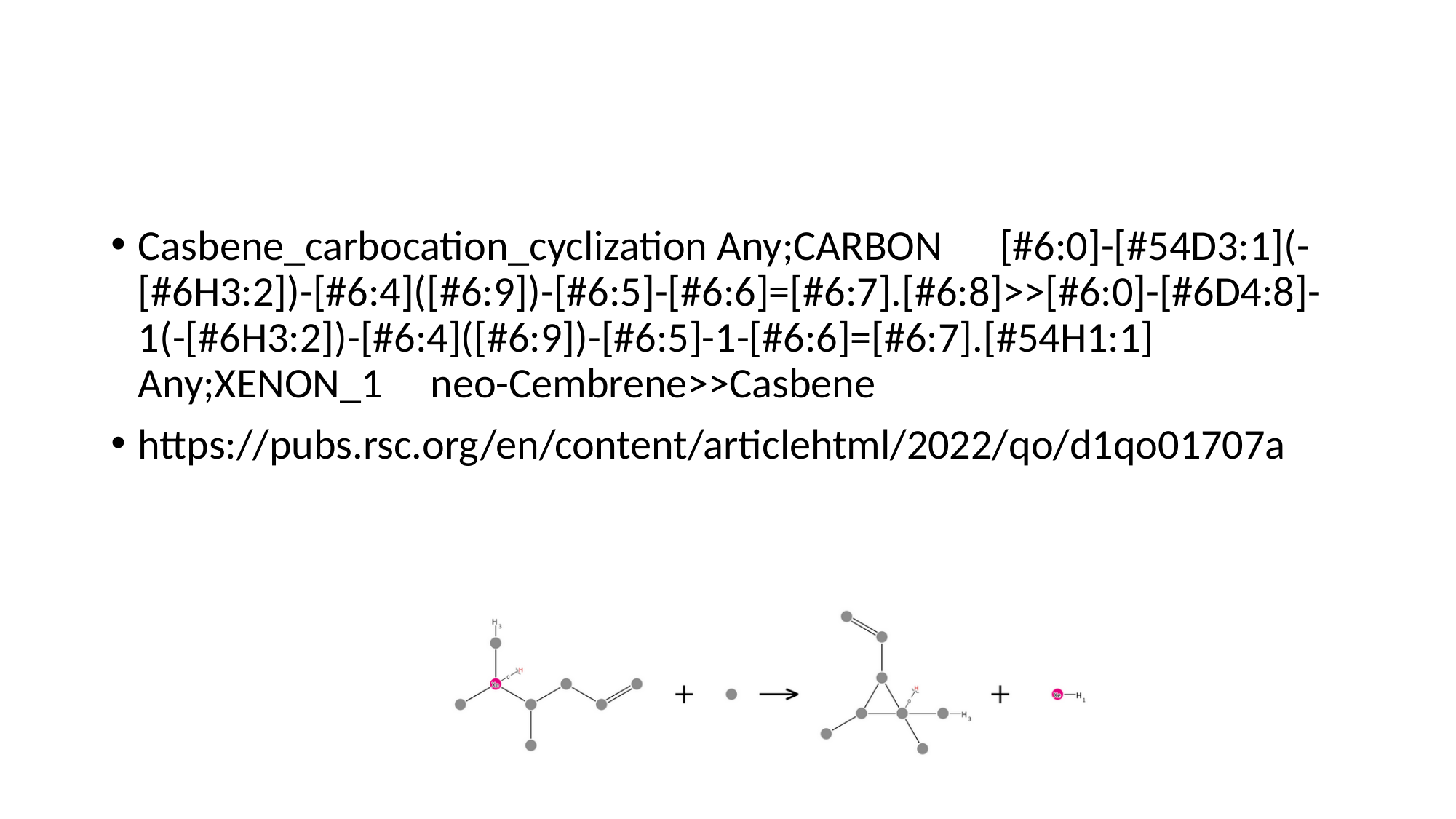

#
Casbene_carbocation_cyclization Any;CARBON [#6:0]-[#54D3:1](-[#6H3:2])-[#6:4]([#6:9])-[#6:5]-[#6:6]=[#6:7].[#6:8]>>[#6:0]-[#6D4:8]-1(-[#6H3:2])-[#6:4]([#6:9])-[#6:5]-1-[#6:6]=[#6:7].[#54H1:1] Any;XENON_1 neo-Cembrene>>Casbene
https://pubs.rsc.org/en/content/articlehtml/2022/qo/d1qo01707a

### Slide 3
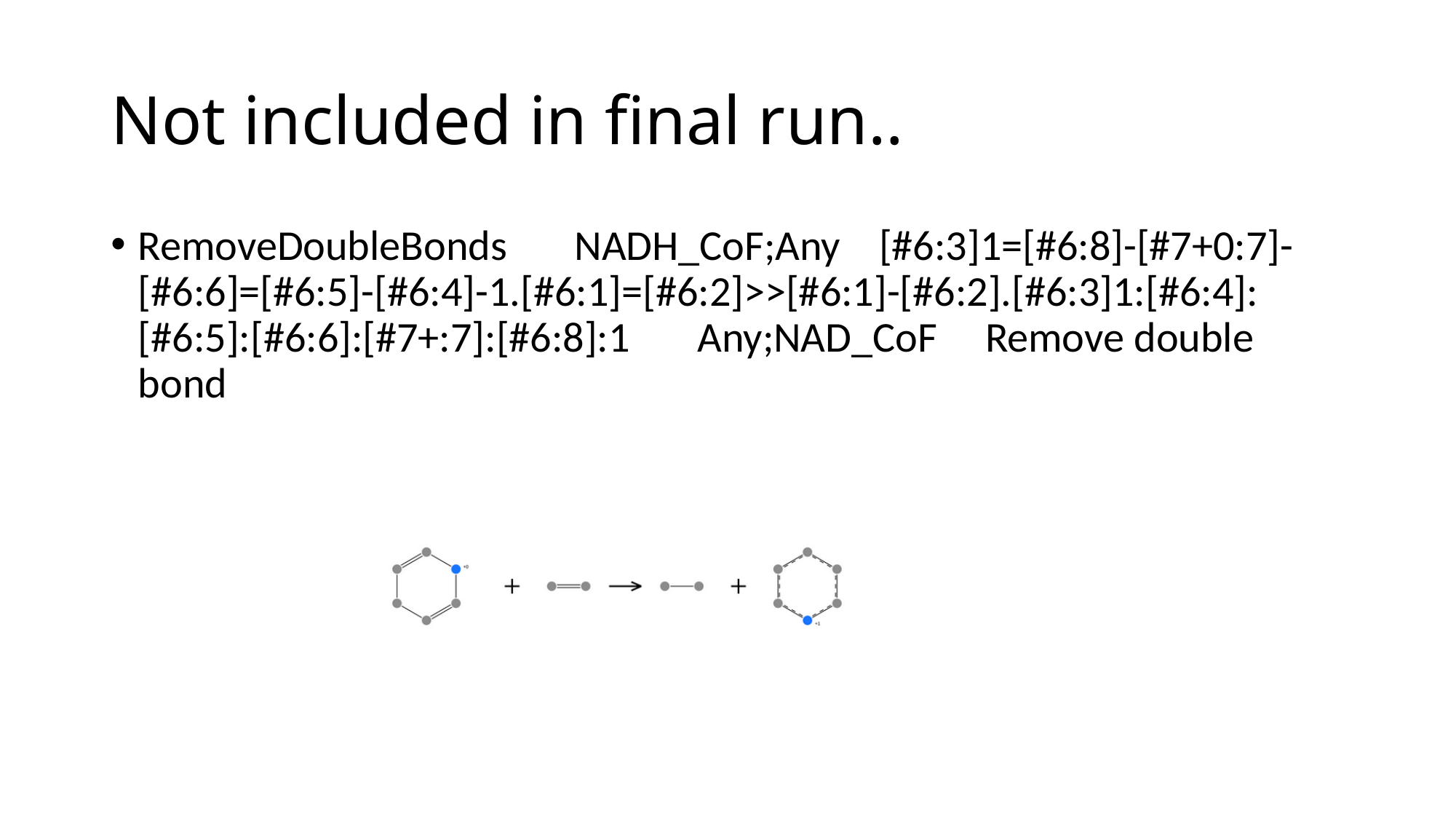

#### Not included in final run..
RemoveDoubleBonds NADH_CoF;Any [#6:3]1=[#6:8]-[#7+0:7]-[#6:6]=[#6:5]-[#6:4]-1.[#6:1]=[#6:2]>>[#6:1]-[#6:2].[#6:3]1:[#6:4]:[#6:5]:[#6:6]:[#7+:7]:[#6:8]:1 Any;NAD_CoF Remove double bond

### Slide 4
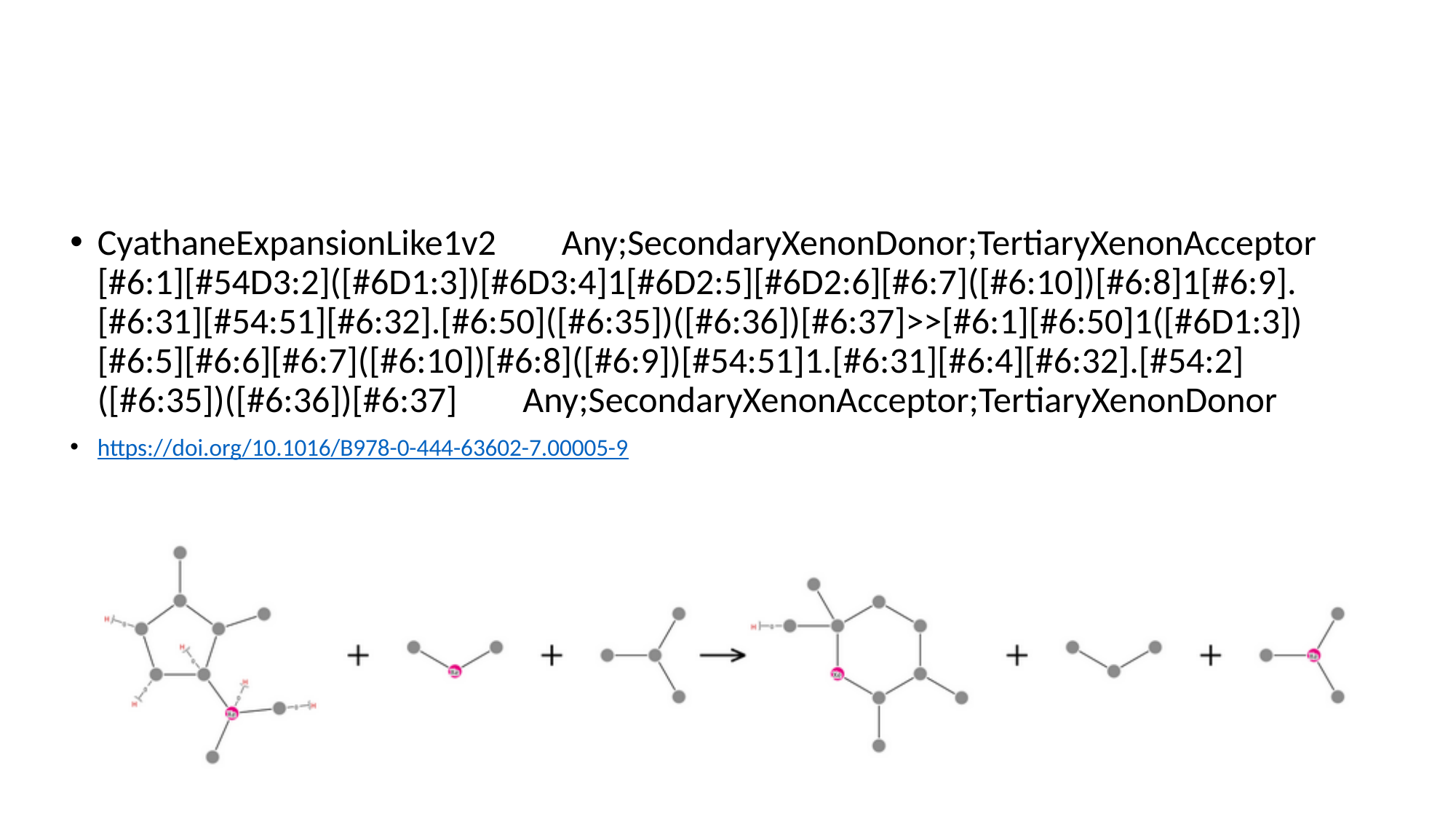

#
CyathaneExpansionLike1v2 Any;SecondaryXenonDonor;TertiaryXenonAcceptor [#6:1][#54D3:2]([#6D1:3])[#6D3:4]1[#6D2:5][#6D2:6][#6:7]([#6:10])[#6:8]1[#6:9].[#6:31][#54:51][#6:32].[#6:50]([#6:35])([#6:36])[#6:37]>>[#6:1][#6:50]1([#6D1:3])[#6:5][#6:6][#6:7]([#6:10])[#6:8]([#6:9])[#54:51]1.[#6:31][#6:4][#6:32].[#54:2]([#6:35])([#6:36])[#6:37] Any;SecondaryXenonAcceptor;TertiaryXenonDonor
https://doi.org/10.1016/B978-0-444-63602-7.00005-9

### Slide 5
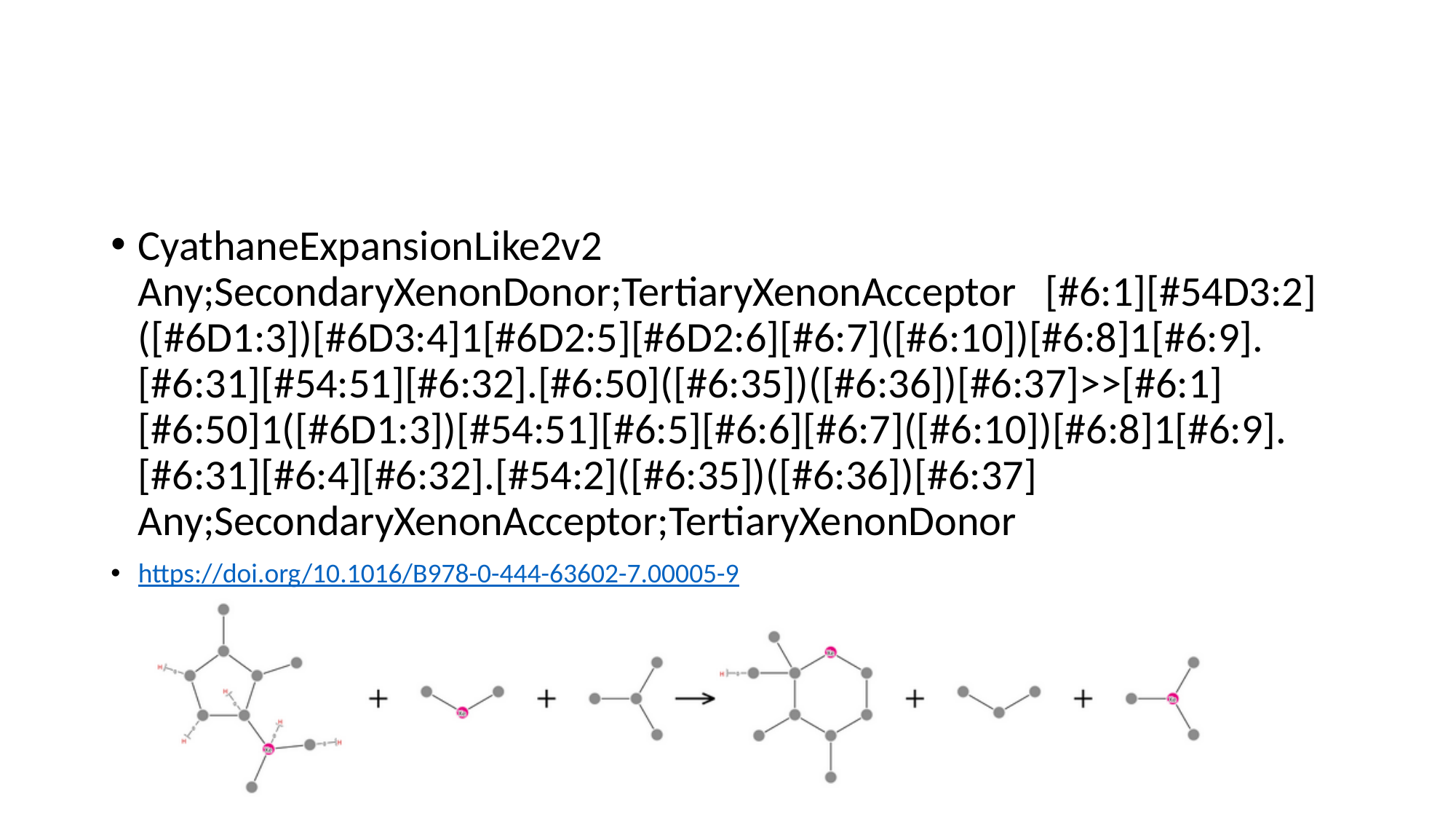

#
CyathaneExpansionLike2v2 Any;SecondaryXenonDonor;TertiaryXenonAcceptor [#6:1][#54D3:2]([#6D1:3])[#6D3:4]1[#6D2:5][#6D2:6][#6:7]([#6:10])[#6:8]1[#6:9].[#6:31][#54:51][#6:32].[#6:50]([#6:35])([#6:36])[#6:37]>>[#6:1][#6:50]1([#6D1:3])[#54:51][#6:5][#6:6][#6:7]([#6:10])[#6:8]1[#6:9].[#6:31][#6:4][#6:32].[#54:2]([#6:35])([#6:36])[#6:37] Any;SecondaryXenonAcceptor;TertiaryXenonDonor
https://doi.org/10.1016/B978-0-444-63602-7.00005-9

### Slide 6
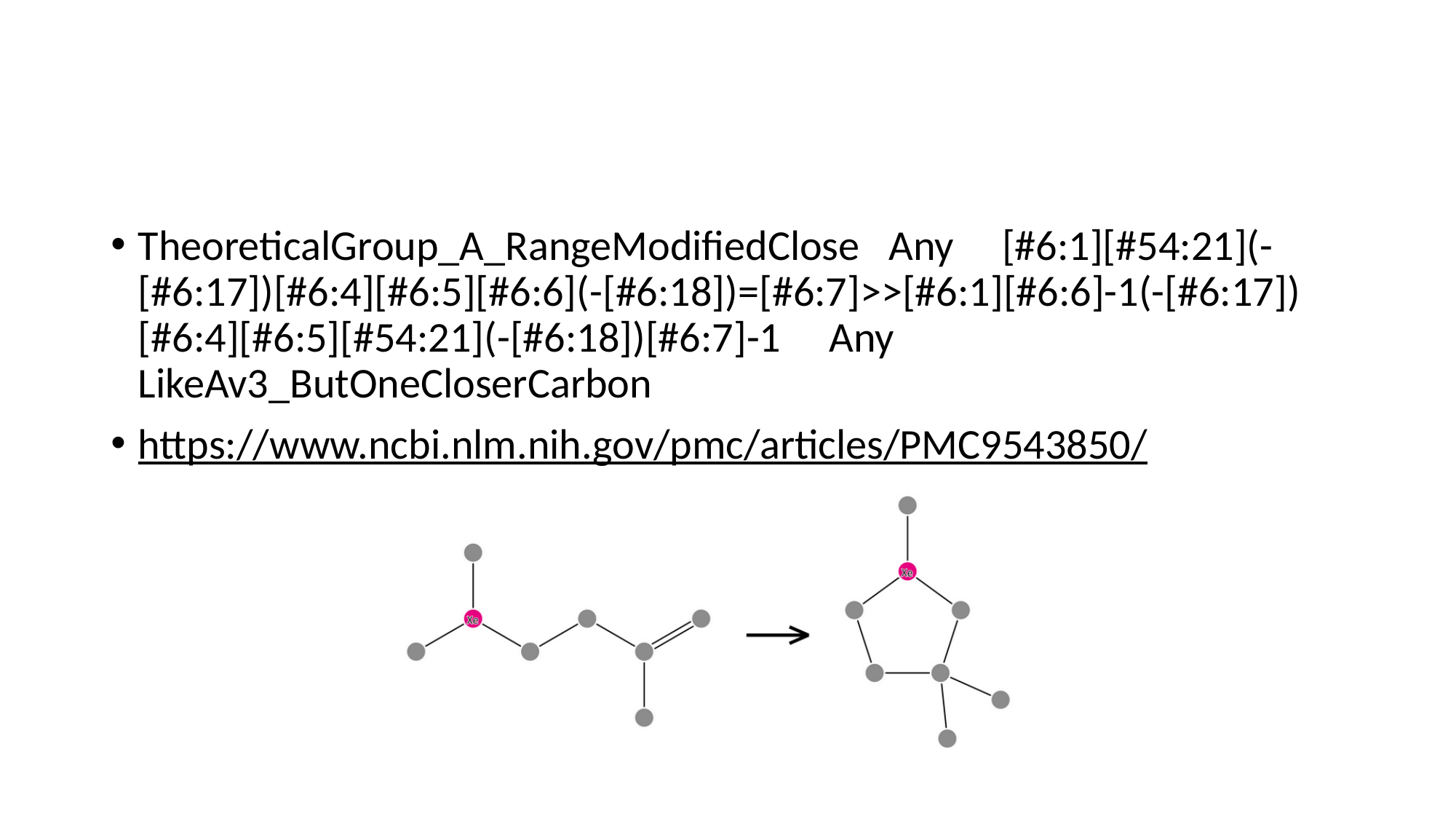

#
TheoreticalGroup_A_RangeModifiedClose Any [#6:1][#54:21](-[#6:17])[#6:4][#6:5][#6:6](-[#6:18])=[#6:7]>>[#6:1][#6:6]-1(-[#6:17])[#6:4][#6:5][#54:21](-[#6:18])[#6:7]-1 Any LikeAv3_ButOneCloserCarbon
https://www.ncbi.nlm.nih.gov/pmc/articles/PMC9543850/

### Slide 7
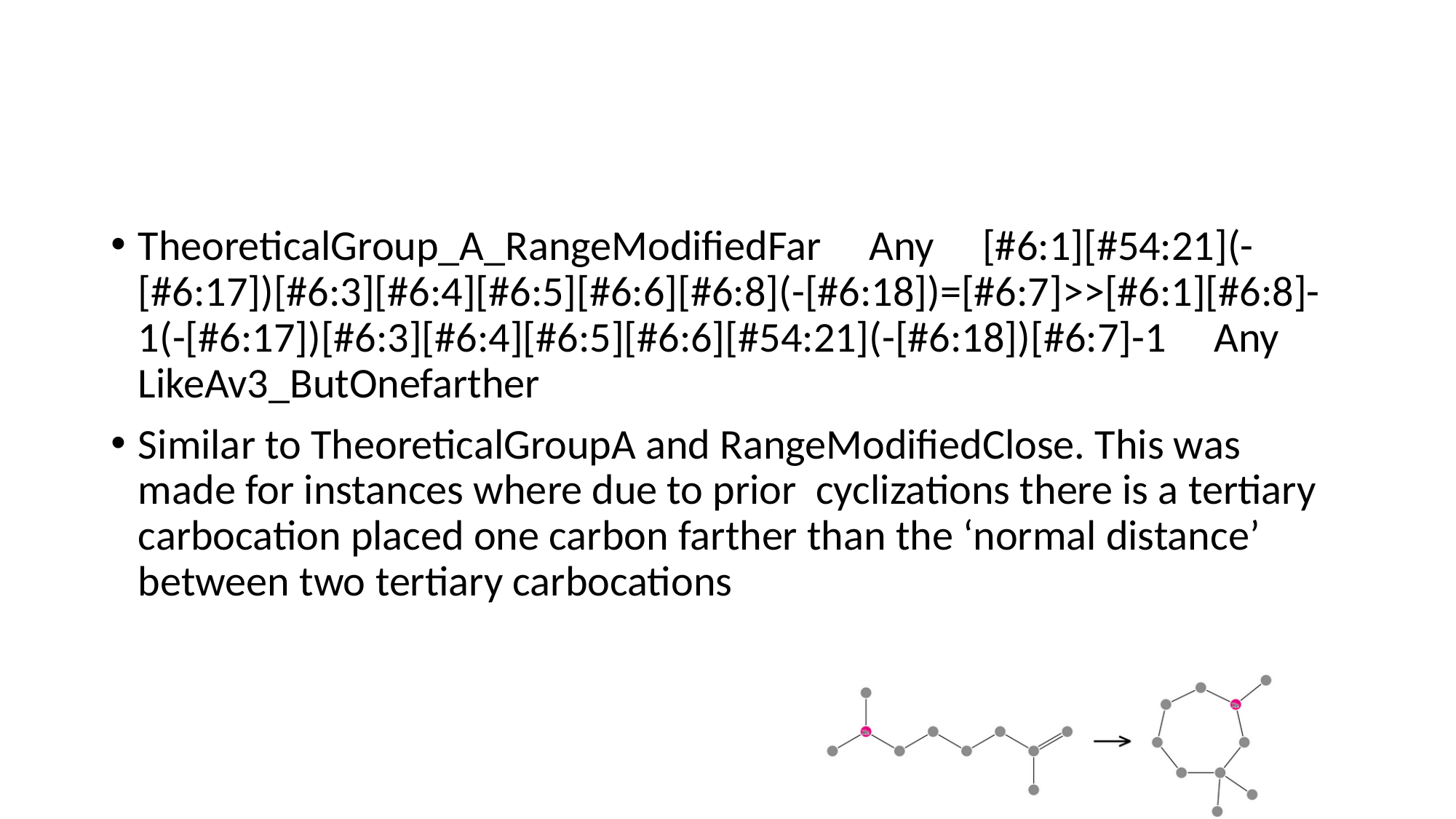

TheoreticalGroup_A_RangeModifiedFar Any [#6:1][#54:21](-[#6:17])[#6:3][#6:4][#6:5][#6:6][#6:8](-[#6:18])=[#6:7]>>[#6:1][#6:8]-1(-[#6:17])[#6:3][#6:4][#6:5][#6:6][#54:21](-[#6:18])[#6:7]-1 Any LikeAv3_ButOnefarther
Similar to TheoreticalGroupA and RangeModifiedClose. This was made for instances where due to prior cyclizations there is a tertiary carbocation placed one carbon farther than the ‘normal distance’ between two tertiary carbocations

### Slide 8
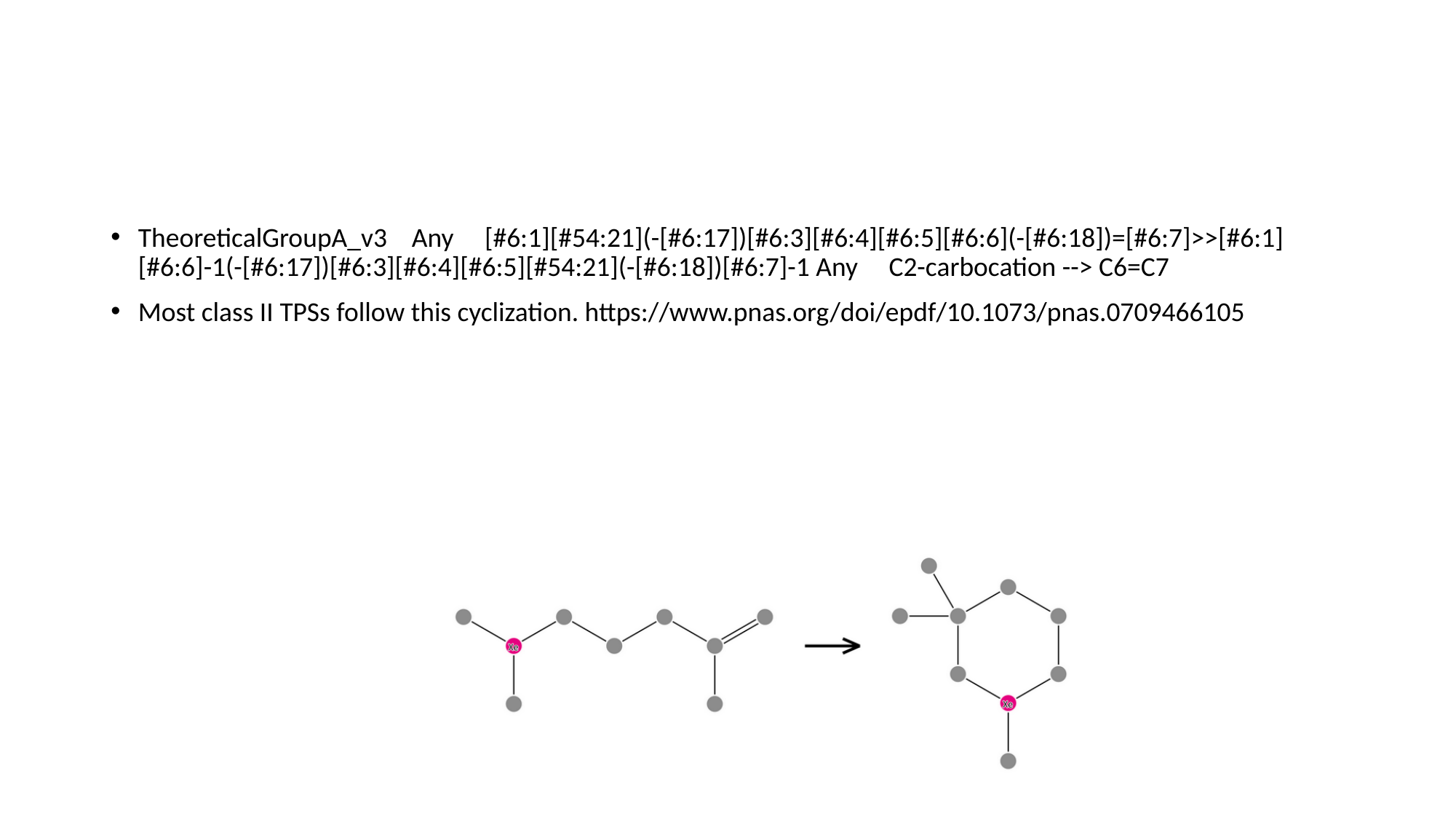

#
TheoreticalGroupA_v3 Any [#6:1][#54:21](-[#6:17])[#6:3][#6:4][#6:5][#6:6](-[#6:18])=[#6:7]>>[#6:1][#6:6]-1(-[#6:17])[#6:3][#6:4][#6:5][#54:21](-[#6:18])[#6:7]-1 Any C2-carbocation --> C6=C7
Most class II TPSs follow this cyclization. https://www.pnas.org/doi/epdf/10.1073/pnas.0709466105

### Slide 9
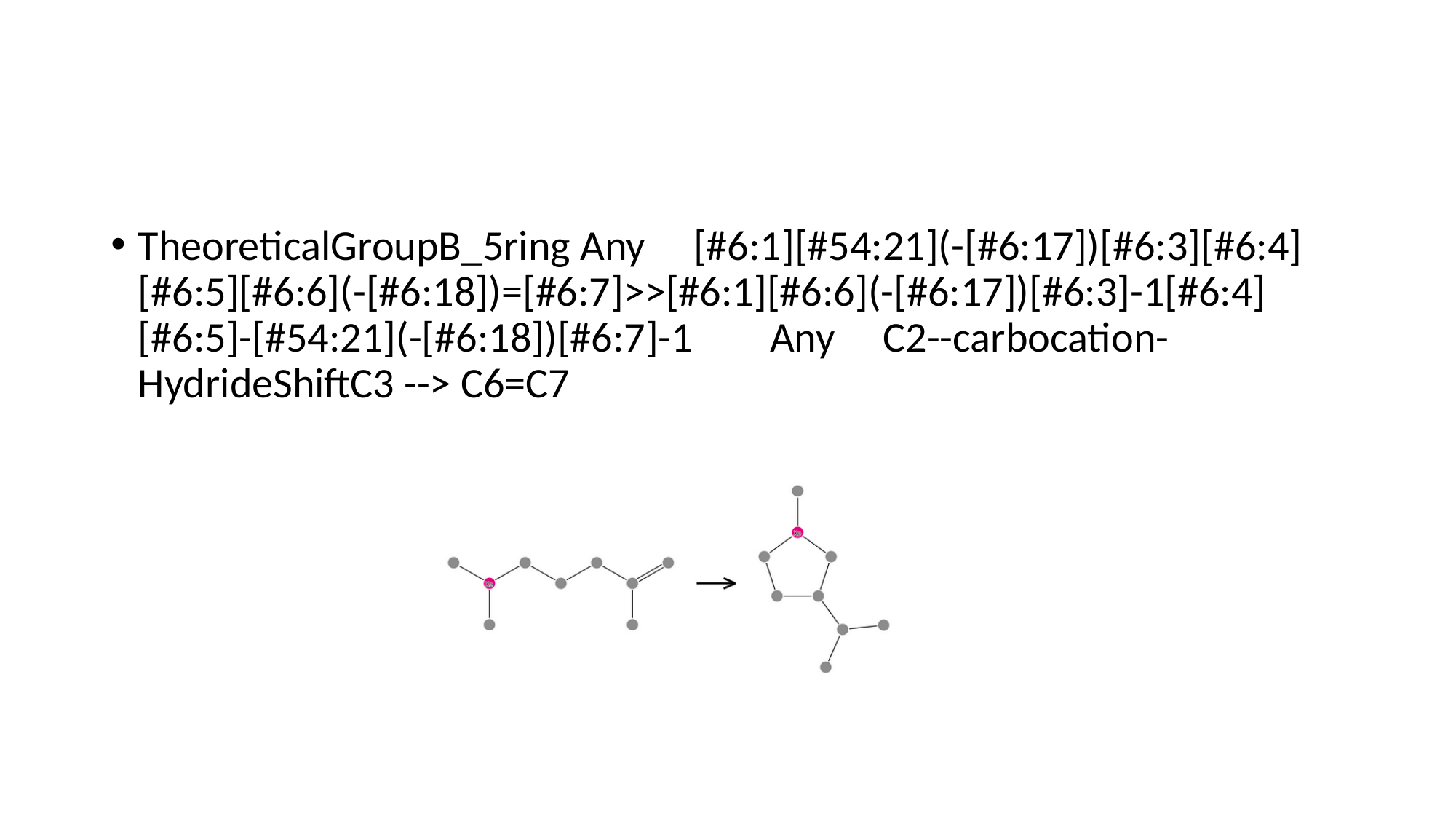

#
TheoreticalGroupB_5ring Any [#6:1][#54:21](-[#6:17])[#6:3][#6:4][#6:5][#6:6](-[#6:18])=[#6:7]>>[#6:1][#6:6](-[#6:17])[#6:3]-1[#6:4][#6:5]-[#54:21](-[#6:18])[#6:7]-1 Any C2--carbocation-HydrideShiftC3 --> C6=C7

### Slide 10
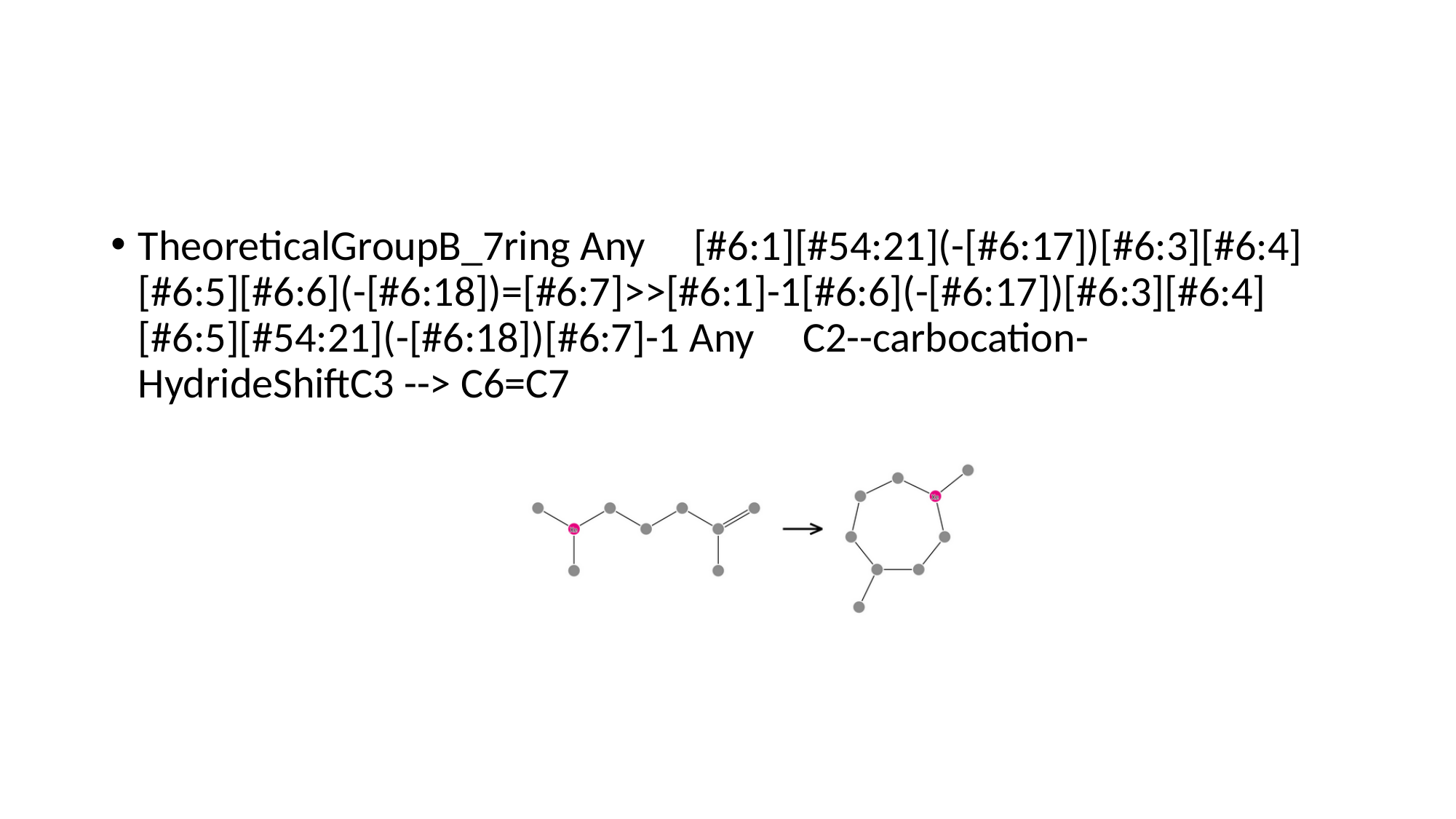

#
TheoreticalGroupB_7ring Any [#6:1][#54:21](-[#6:17])[#6:3][#6:4][#6:5][#6:6](-[#6:18])=[#6:7]>>[#6:1]-1[#6:6](-[#6:17])[#6:3][#6:4][#6:5][#54:21](-[#6:18])[#6:7]-1 Any C2--carbocation-HydrideShiftC3 --> C6=C7

### Slide 11
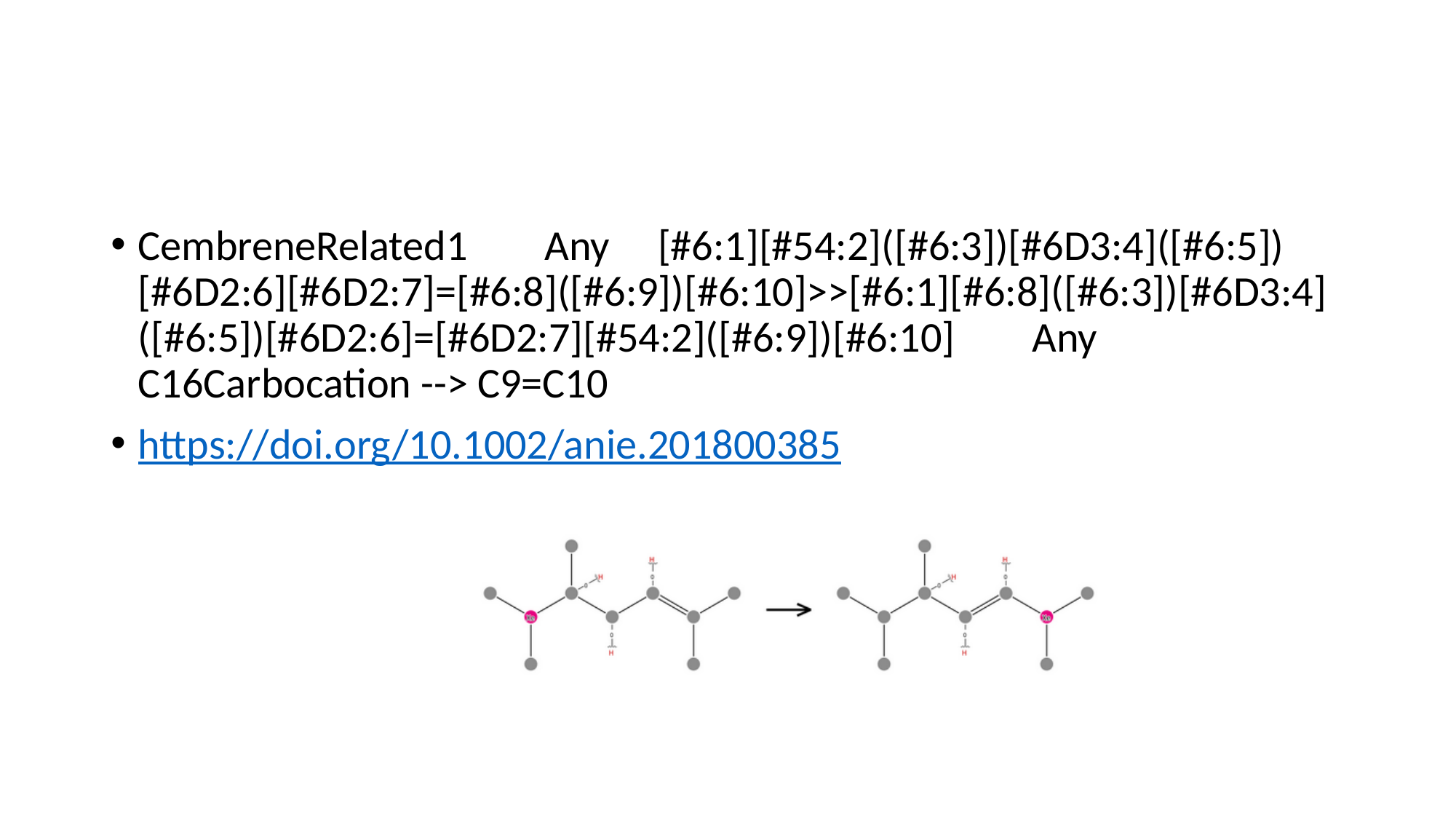

#
CembreneRelated1 Any [#6:1][#54:2]([#6:3])[#6D3:4]([#6:5])[#6D2:6][#6D2:7]=[#6:8]([#6:9])[#6:10]>>[#6:1][#6:8]([#6:3])[#6D3:4]([#6:5])[#6D2:6]=[#6D2:7][#54:2]([#6:9])[#6:10] Any C16Carbocation --> C9=C10
https://doi.org/10.1002/anie.201800385

### Slide 12
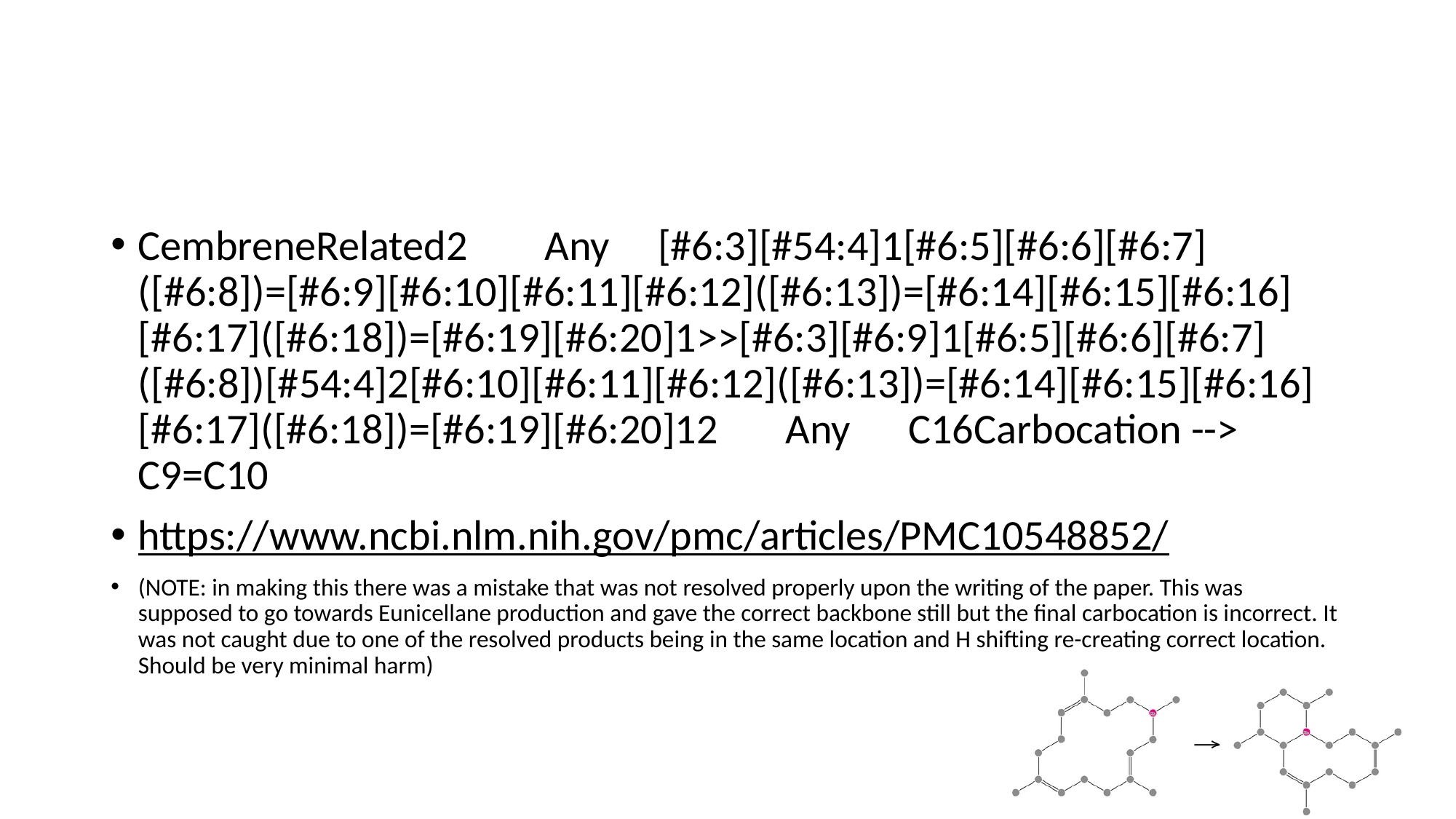

#
CembreneRelated2 Any [#6:3][#54:4]1[#6:5][#6:6][#6:7]([#6:8])=[#6:9][#6:10][#6:11][#6:12]([#6:13])=[#6:14][#6:15][#6:16][#6:17]([#6:18])=[#6:19][#6:20]1>>[#6:3][#6:9]1[#6:5][#6:6][#6:7]([#6:8])[#54:4]2[#6:10][#6:11][#6:12]([#6:13])=[#6:14][#6:15][#6:16][#6:17]([#6:18])=[#6:19][#6:20]12 Any C16Carbocation --> C9=C10
https://www.ncbi.nlm.nih.gov/pmc/articles/PMC10548852/
(NOTE: in making this there was a mistake that was not resolved properly upon the writing of the paper. This was supposed to go towards Eunicellane production and gave the correct backbone still but the final carbocation is incorrect. It was not caught due to one of the resolved products being in the same location and H shifting re-creating correct location. Should be very minimal harm)

### Slide 13
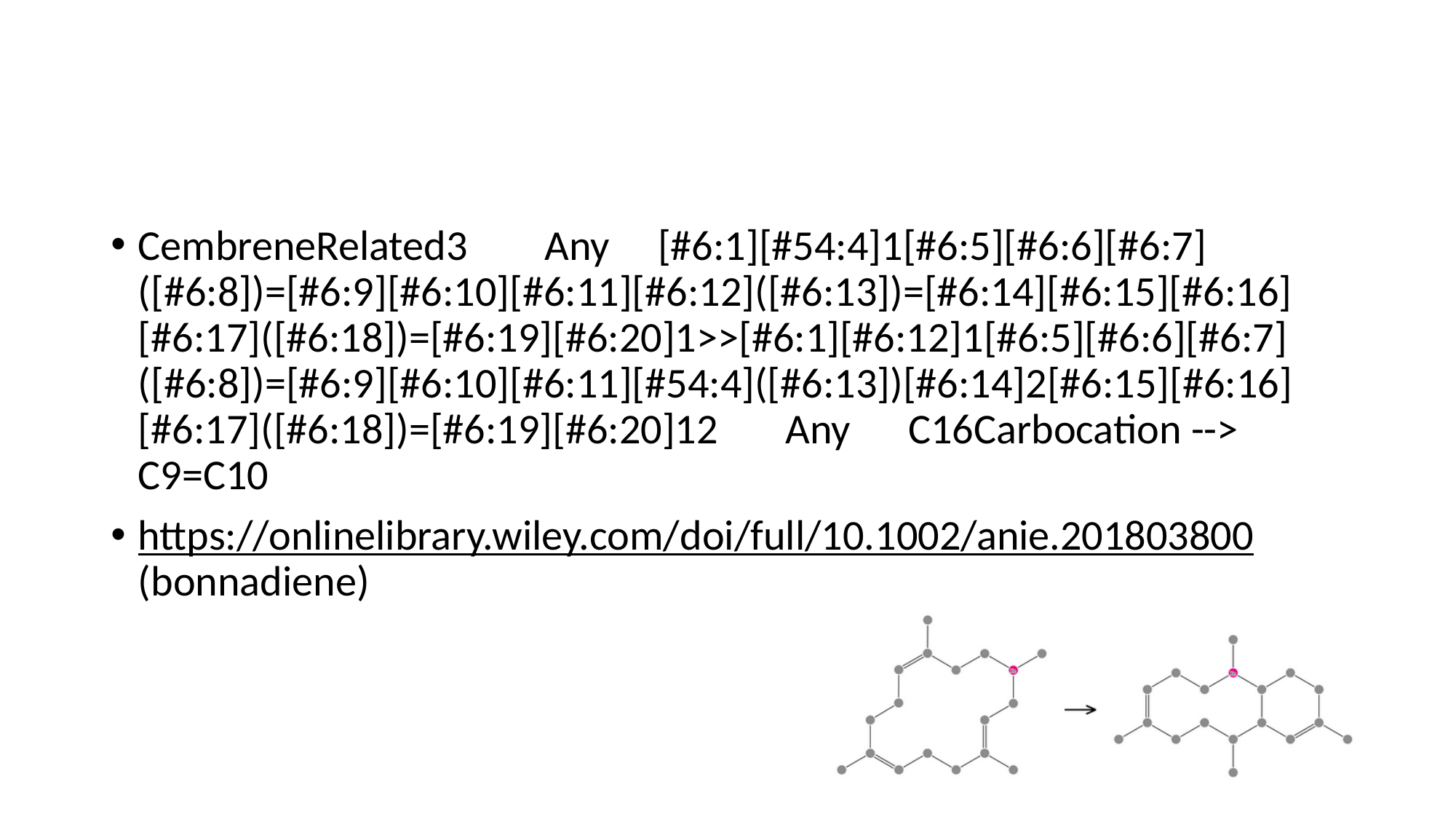

#
CembreneRelated3 Any [#6:1][#54:4]1[#6:5][#6:6][#6:7]([#6:8])=[#6:9][#6:10][#6:11][#6:12]([#6:13])=[#6:14][#6:15][#6:16][#6:17]([#6:18])=[#6:19][#6:20]1>>[#6:1][#6:12]1[#6:5][#6:6][#6:7]([#6:8])=[#6:9][#6:10][#6:11][#54:4]([#6:13])[#6:14]2[#6:15][#6:16][#6:17]([#6:18])=[#6:19][#6:20]12 Any C16Carbocation --> C9=C10
https://onlinelibrary.wiley.com/doi/full/10.1002/anie.201803800 (bonnadiene)

### Slide 14
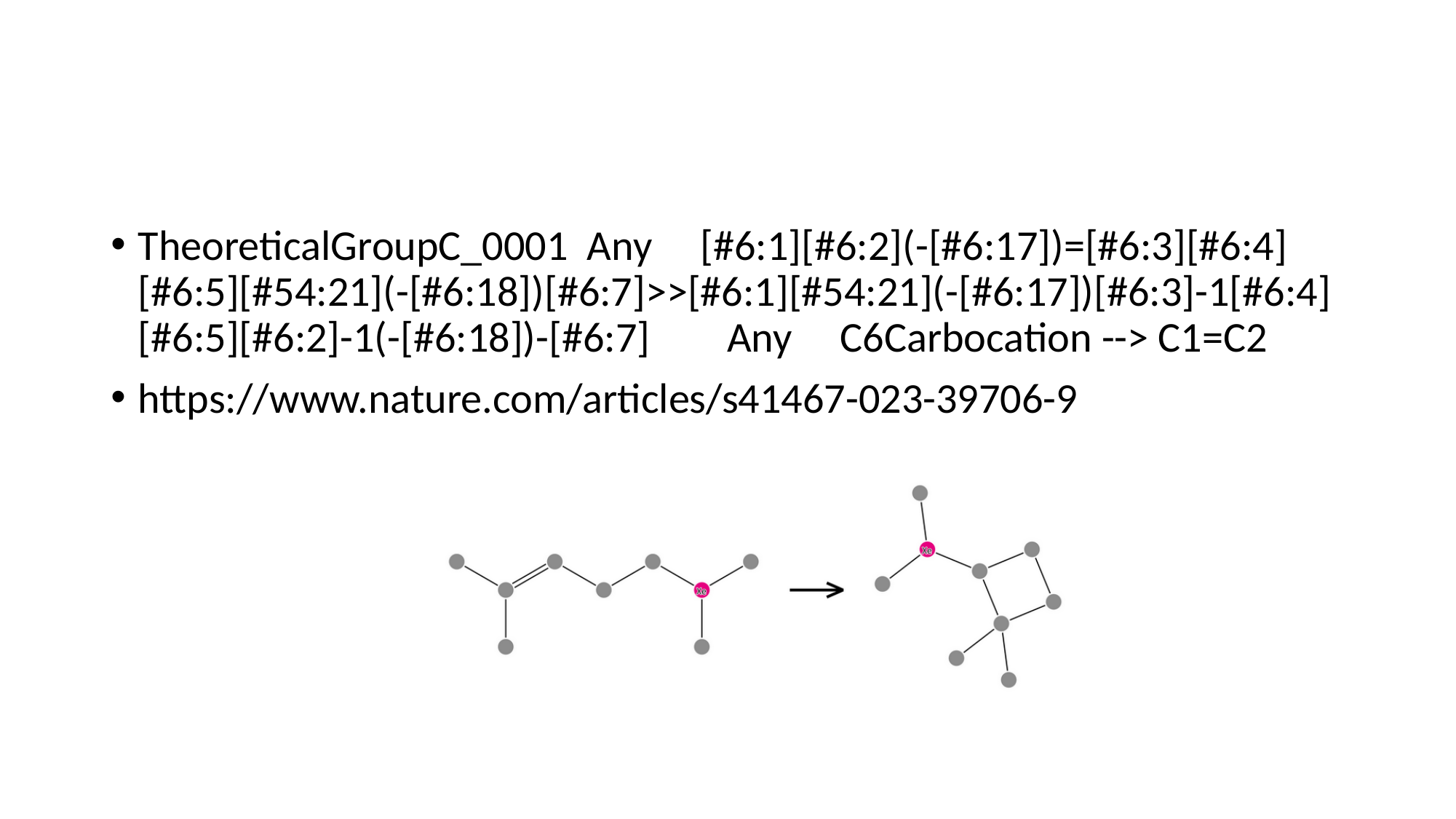

#
TheoreticalGroupC_0001 Any [#6:1][#6:2](-[#6:17])=[#6:3][#6:4][#6:5][#54:21](-[#6:18])[#6:7]>>[#6:1][#54:21](-[#6:17])[#6:3]-1[#6:4][#6:5][#6:2]-1(-[#6:18])-[#6:7] Any C6Carbocation --> C1=C2
https://www.nature.com/articles/s41467-023-39706-9

### Slide 15
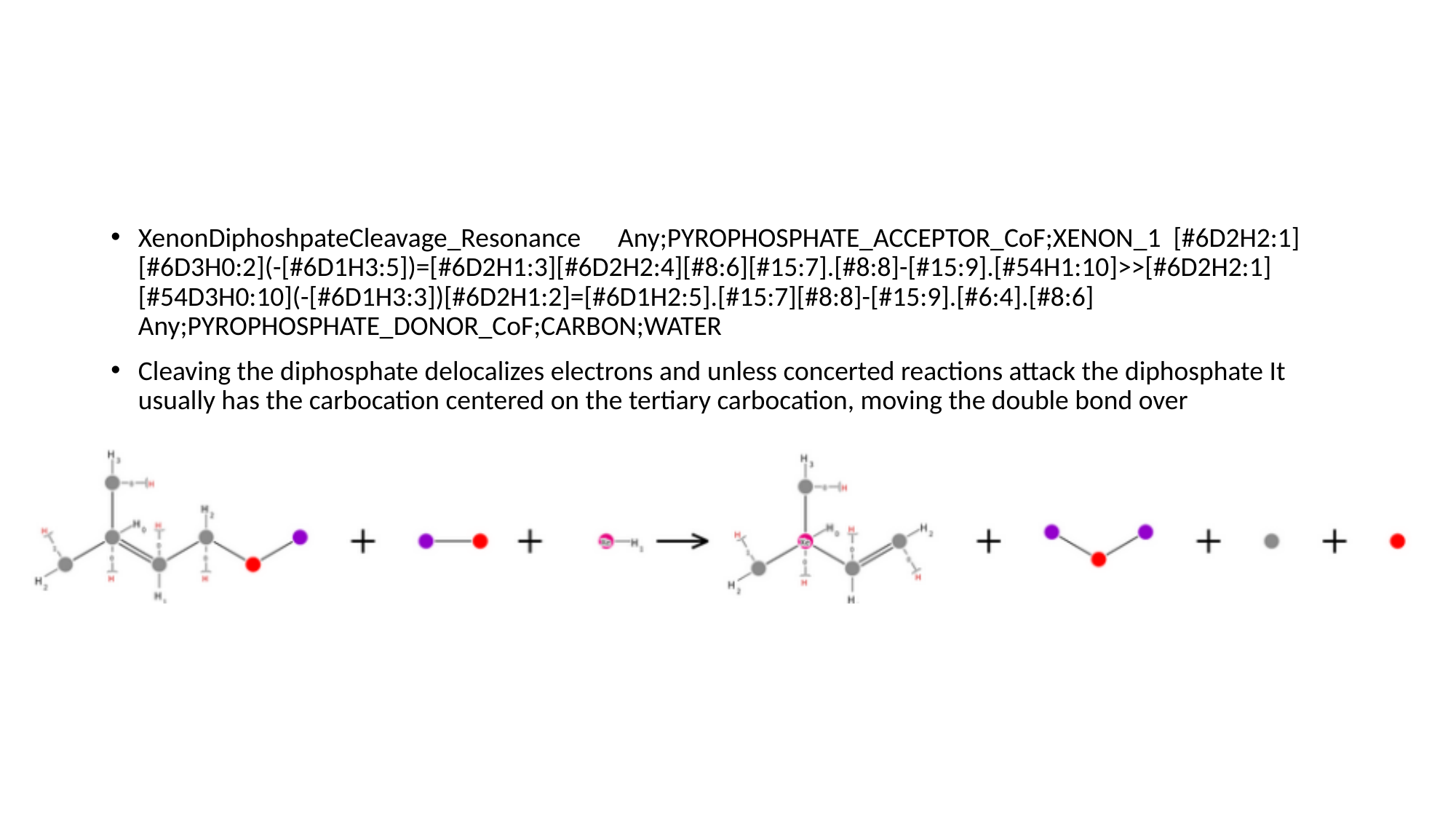

#
XenonDiphoshpateCleavage_Resonance Any;PYROPHOSPHATE_ACCEPTOR_CoF;XENON_1 [#6D2H2:1][#6D3H0:2](-[#6D1H3:5])=[#6D2H1:3][#6D2H2:4][#8:6][#15:7].[#8:8]-[#15:9].[#54H1:10]>>[#6D2H2:1][#54D3H0:10](-[#6D1H3:3])[#6D2H1:2]=[#6D1H2:5].[#15:7][#8:8]-[#15:9].[#6:4].[#8:6] Any;PYROPHOSPHATE_DONOR_CoF;CARBON;WATER
Cleaving the diphosphate delocalizes electrons and unless concerted reactions attack the diphosphate It usually has the carbocation centered on the tertiary carbocation, moving the double bond over

### Slide 16
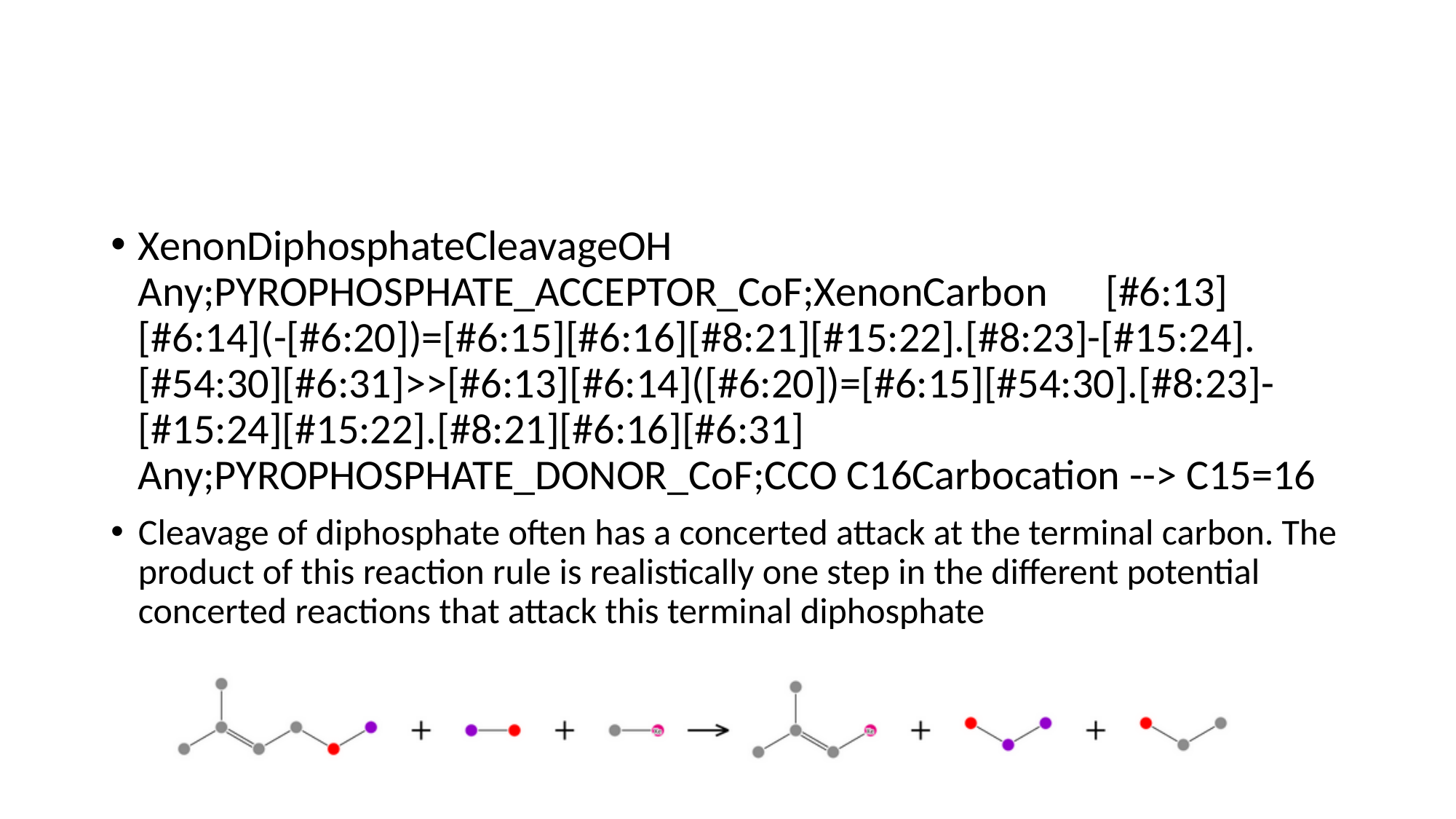

#
XenonDiphosphateCleavageOH Any;PYROPHOSPHATE_ACCEPTOR_CoF;XenonCarbon [#6:13][#6:14](-[#6:20])=[#6:15][#6:16][#8:21][#15:22].[#8:23]-[#15:24].[#54:30][#6:31]>>[#6:13][#6:14]([#6:20])=[#6:15][#54:30].[#8:23]-[#15:24][#15:22].[#8:21][#6:16][#6:31] Any;PYROPHOSPHATE_DONOR_CoF;CCO C16Carbocation --> C15=16
Cleavage of diphosphate often has a concerted attack at the terminal carbon. The product of this reaction rule is realistically one step in the different potential concerted reactions that attack this terminal diphosphate

### Slide 17
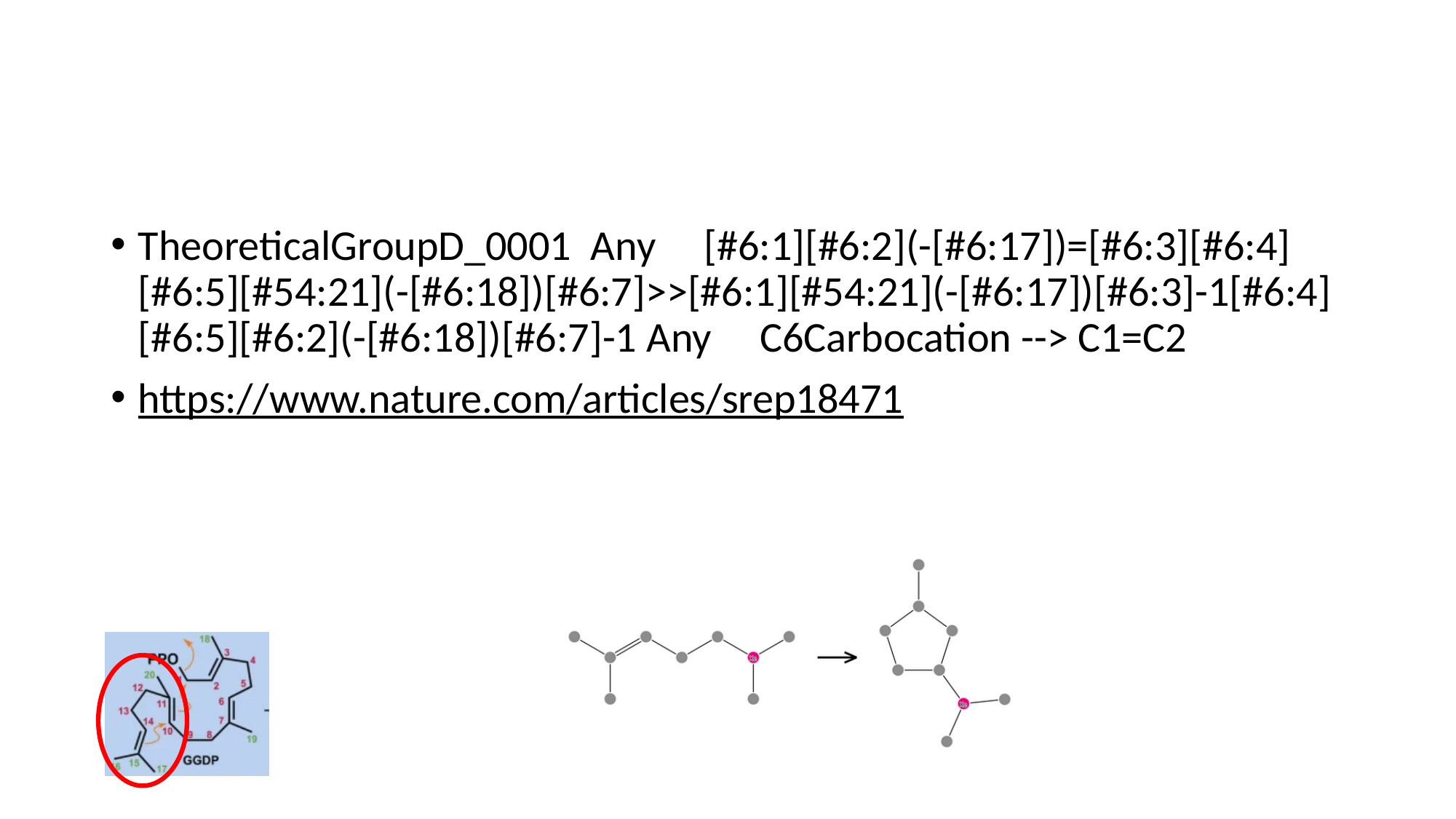

#
TheoreticalGroupD_0001 Any [#6:1][#6:2](-[#6:17])=[#6:3][#6:4][#6:5][#54:21](-[#6:18])[#6:7]>>[#6:1][#54:21](-[#6:17])[#6:3]-1[#6:4][#6:5][#6:2](-[#6:18])[#6:7]-1 Any C6Carbocation --> C1=C2
https://www.nature.com/articles/srep18471

### Slide 18
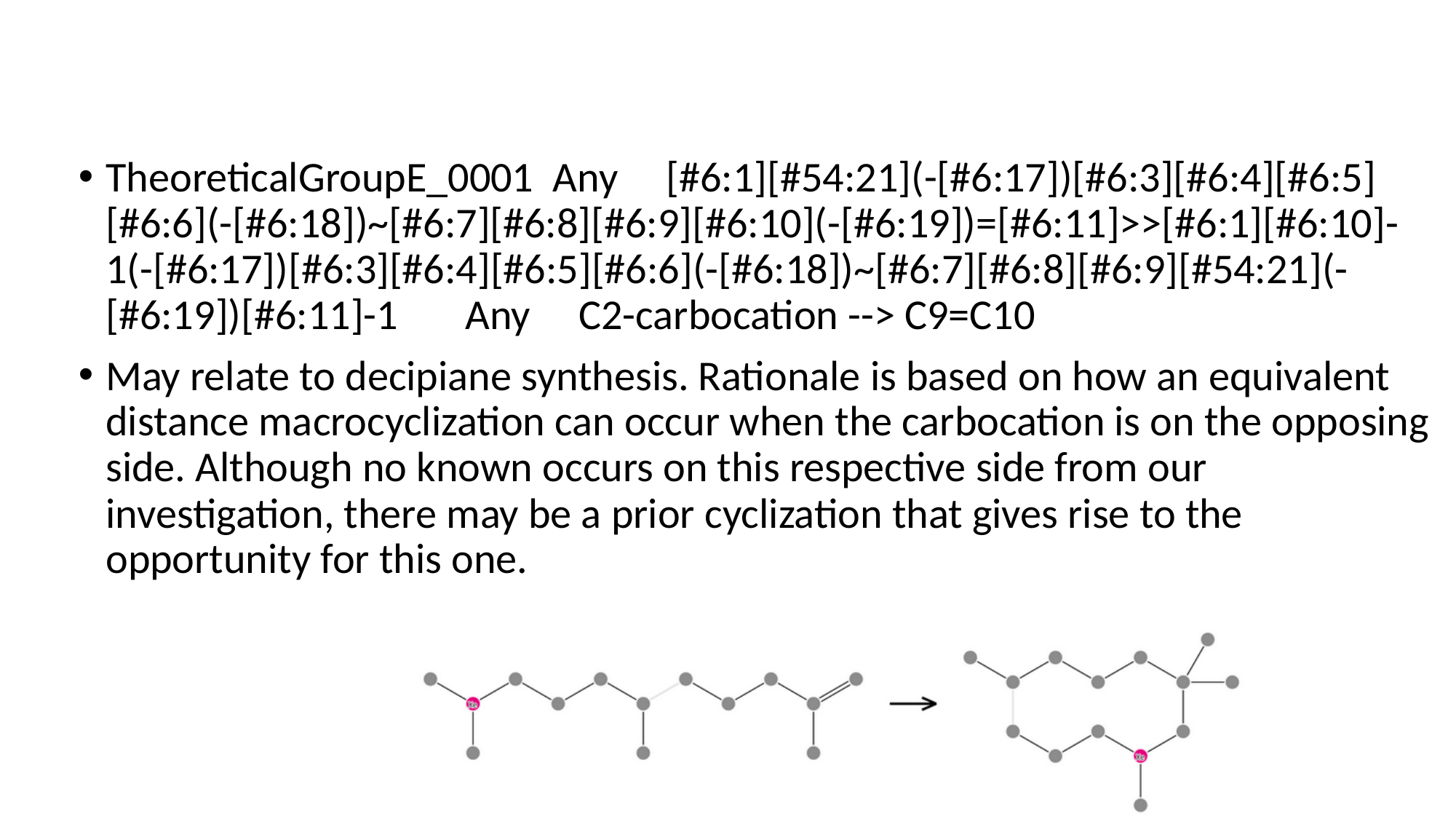

#
TheoreticalGroupE_0001 Any [#6:1][#54:21](-[#6:17])[#6:3][#6:4][#6:5][#6:6](-[#6:18])~[#6:7][#6:8][#6:9][#6:10](-[#6:19])=[#6:11]>>[#6:1][#6:10]-1(-[#6:17])[#6:3][#6:4][#6:5][#6:6](-[#6:18])~[#6:7][#6:8][#6:9][#54:21](-[#6:19])[#6:11]-1 Any C2-carbocation --> C9=C10
May relate to decipiane synthesis. Rationale is based on how an equivalent distance macrocyclization can occur when the carbocation is on the opposing side. Although no known occurs on this respective side from our investigation, there may be a prior cyclization that gives rise to the opportunity for this one.

### Slide 19
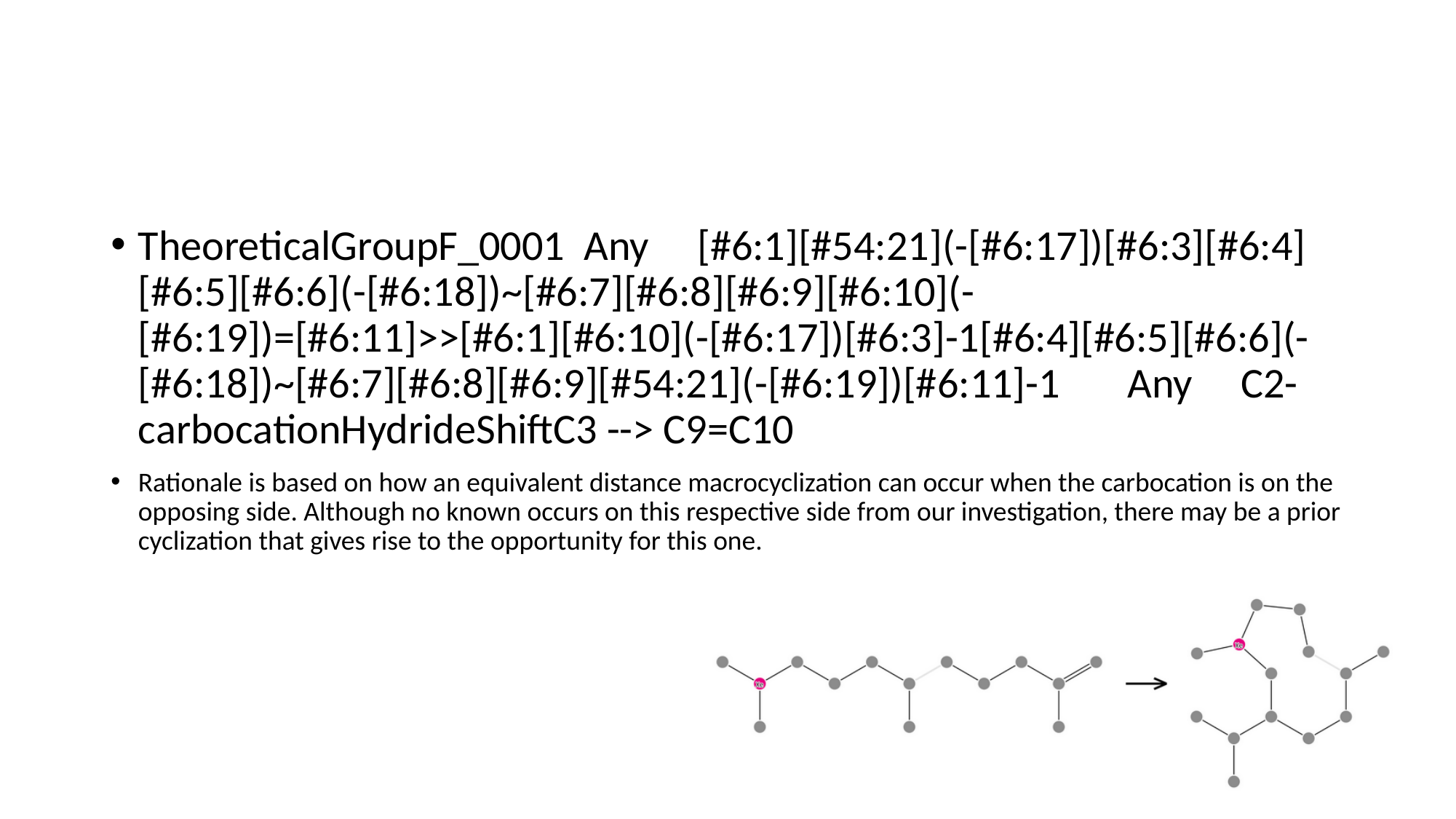

#
TheoreticalGroupF_0001 Any [#6:1][#54:21](-[#6:17])[#6:3][#6:4][#6:5][#6:6](-[#6:18])~[#6:7][#6:8][#6:9][#6:10](-[#6:19])=[#6:11]>>[#6:1][#6:10](-[#6:17])[#6:3]-1[#6:4][#6:5][#6:6](-[#6:18])~[#6:7][#6:8][#6:9][#54:21](-[#6:19])[#6:11]-1 Any C2-carbocationHydrideShiftC3 --> C9=C10
Rationale is based on how an equivalent distance macrocyclization can occur when the carbocation is on the opposing side. Although no known occurs on this respective side from our investigation, there may be a prior cyclization that gives rise to the opportunity for this one.

### Slide 20
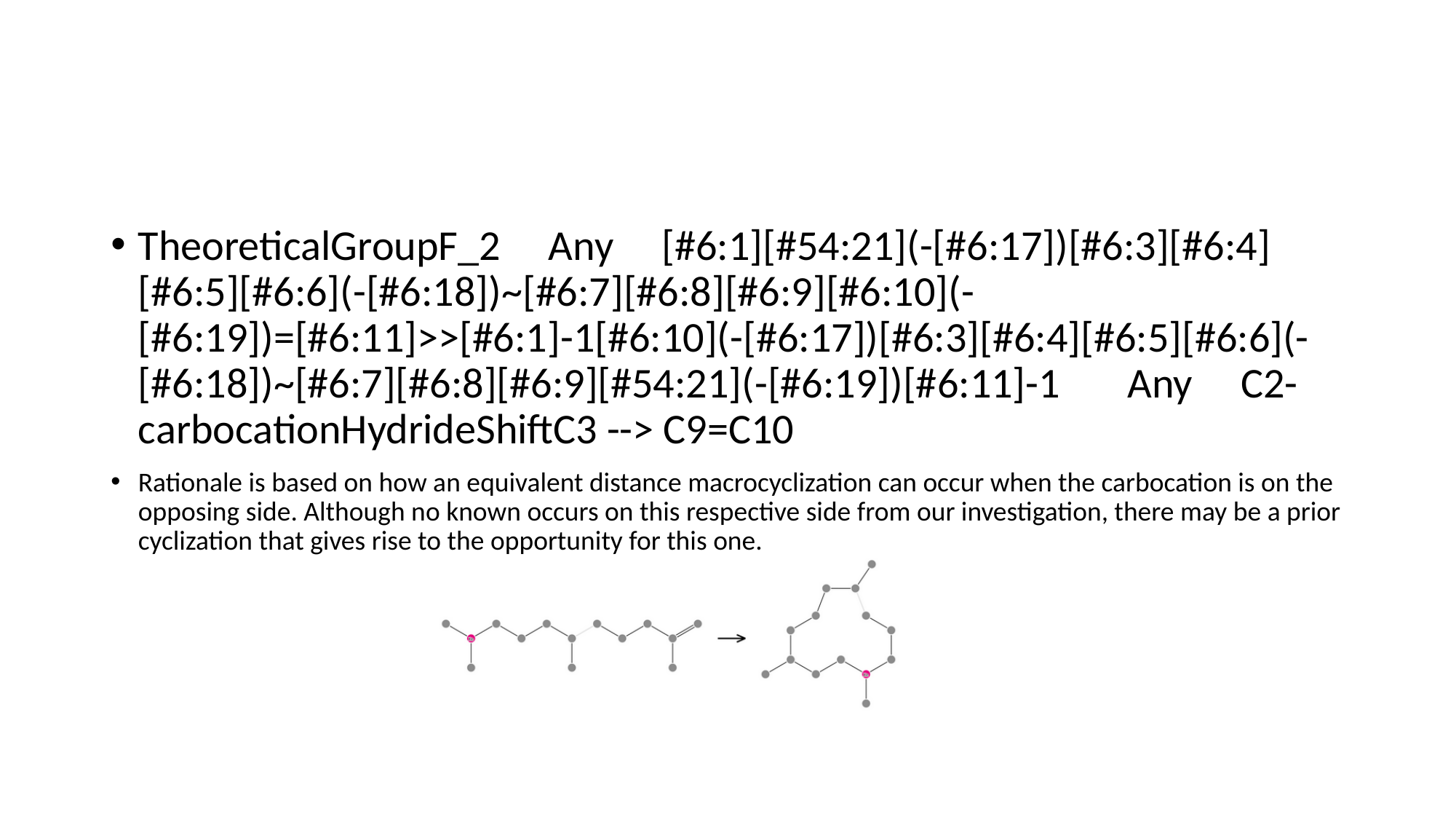

#
TheoreticalGroupF_2 Any [#6:1][#54:21](-[#6:17])[#6:3][#6:4][#6:5][#6:6](-[#6:18])~[#6:7][#6:8][#6:9][#6:10](-[#6:19])=[#6:11]>>[#6:1]-1[#6:10](-[#6:17])[#6:3][#6:4][#6:5][#6:6](-[#6:18])~[#6:7][#6:8][#6:9][#54:21](-[#6:19])[#6:11]-1 Any C2-carbocationHydrideShiftC3 --> C9=C10
Rationale is based on how an equivalent distance macrocyclization can occur when the carbocation is on the opposing side. Although no known occurs on this respective side from our investigation, there may be a prior cyclization that gives rise to the opportunity for this one.

### Slide 21
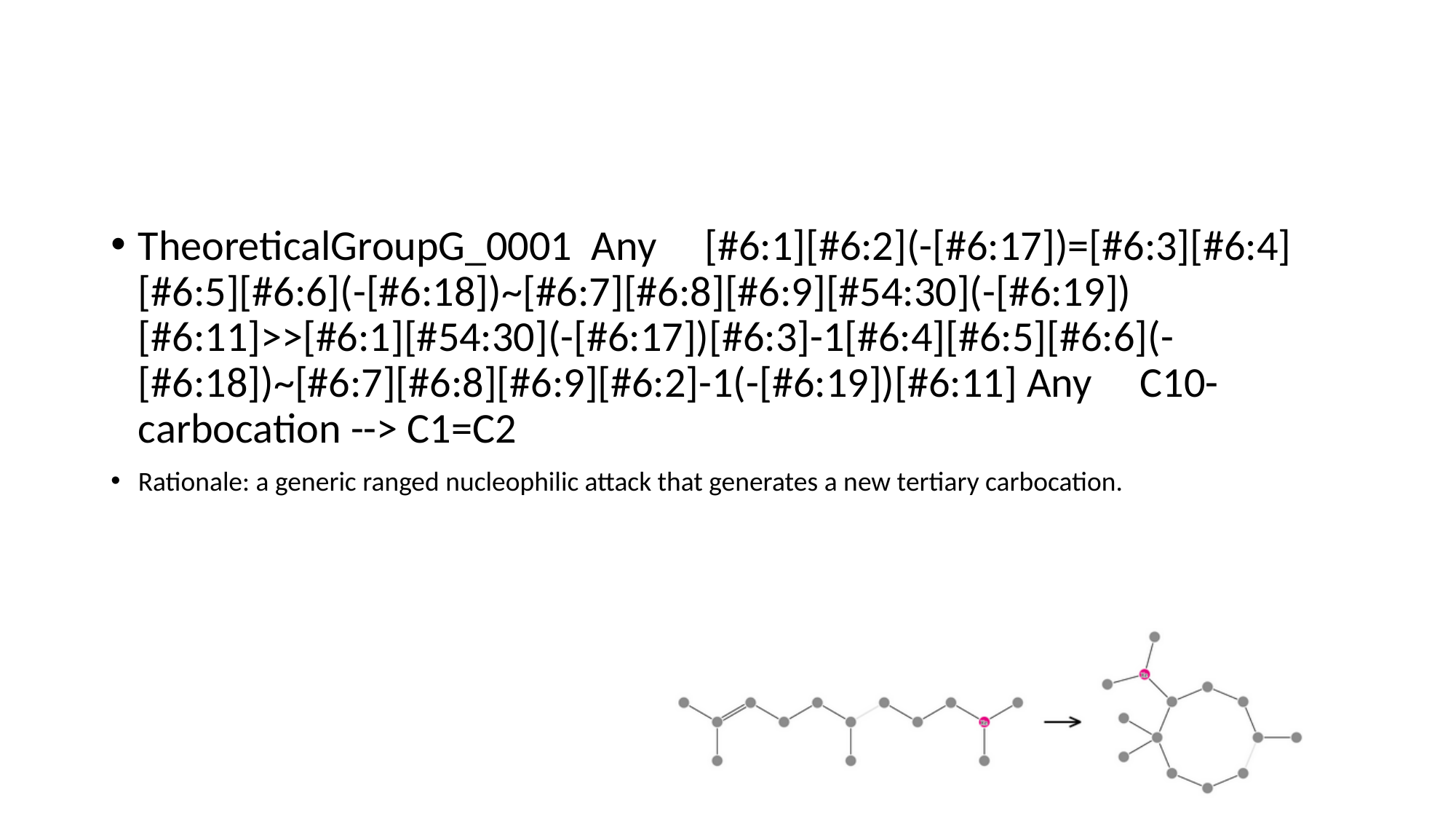

#
TheoreticalGroupG_0001 Any [#6:1][#6:2](-[#6:17])=[#6:3][#6:4][#6:5][#6:6](-[#6:18])~[#6:7][#6:8][#6:9][#54:30](-[#6:19])[#6:11]>>[#6:1][#54:30](-[#6:17])[#6:3]-1[#6:4][#6:5][#6:6](-[#6:18])~[#6:7][#6:8][#6:9][#6:2]-1(-[#6:19])[#6:11] Any C10-carbocation --> C1=C2
Rationale: a generic ranged nucleophilic attack that generates a new tertiary carbocation.

### Slide 22
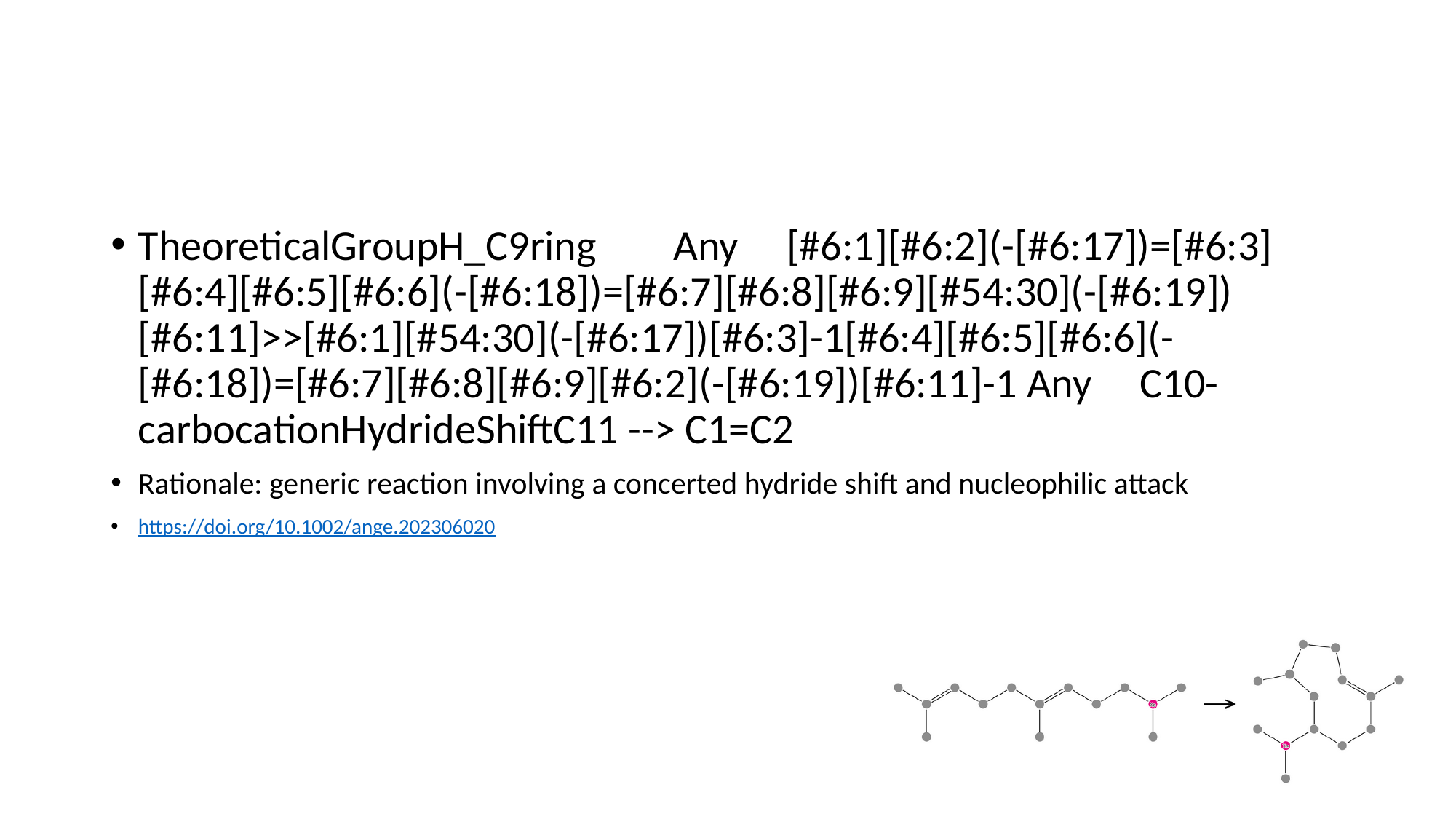

#
TheoreticalGroupH_C9ring Any [#6:1][#6:2](-[#6:17])=[#6:3][#6:4][#6:5][#6:6](-[#6:18])=[#6:7][#6:8][#6:9][#54:30](-[#6:19])[#6:11]>>[#6:1][#54:30](-[#6:17])[#6:3]-1[#6:4][#6:5][#6:6](-[#6:18])=[#6:7][#6:8][#6:9][#6:2](-[#6:19])[#6:11]-1 Any C10-carbocationHydrideShiftC11 --> C1=C2
Rationale: generic reaction involving a concerted hydride shift and nucleophilic attack
https://doi.org/10.1002/ange.202306020

### Slide 23
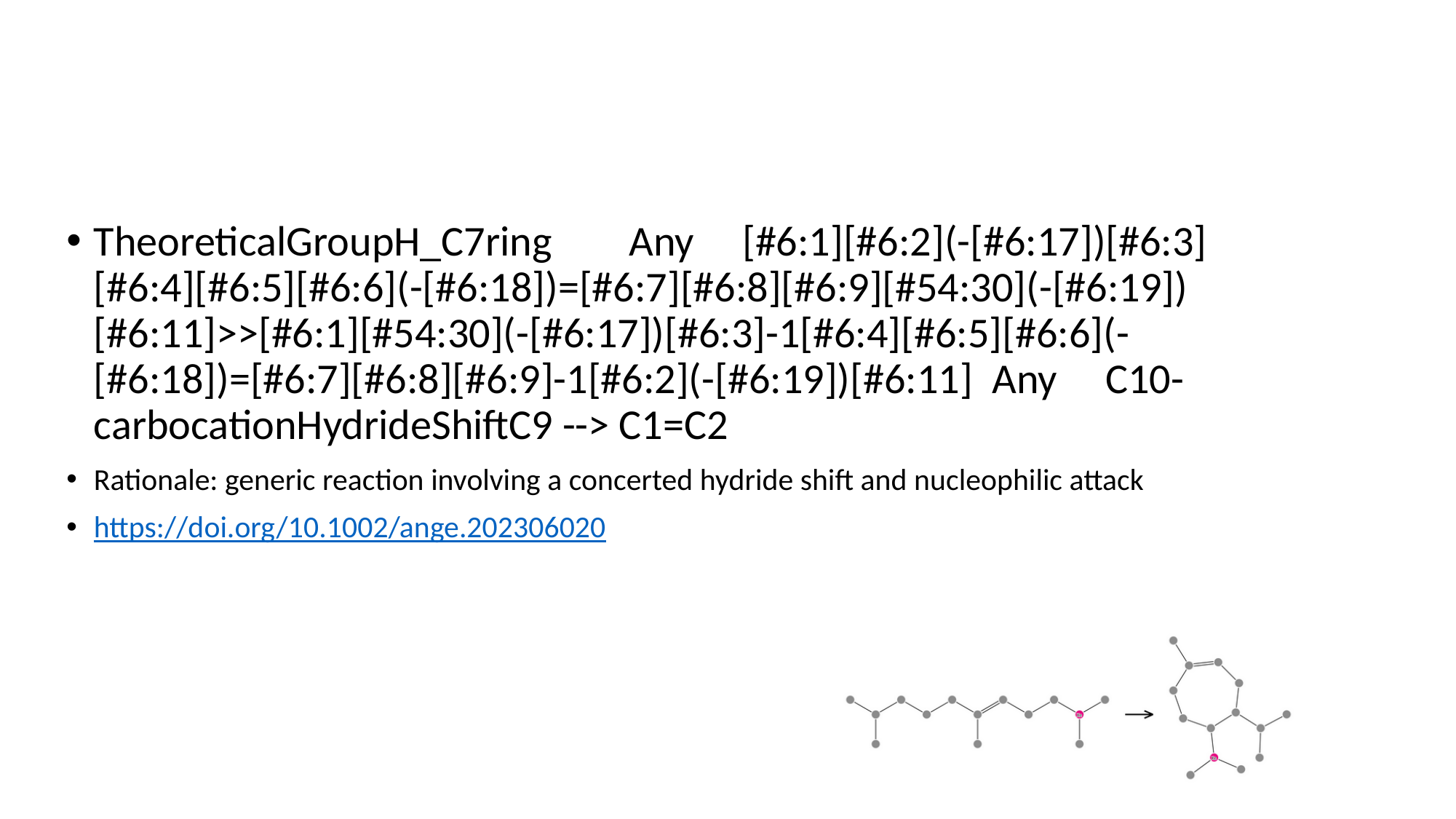

#
TheoreticalGroupH_C7ring Any [#6:1][#6:2](-[#6:17])[#6:3][#6:4][#6:5][#6:6](-[#6:18])=[#6:7][#6:8][#6:9][#54:30](-[#6:19])[#6:11]>>[#6:1][#54:30](-[#6:17])[#6:3]-1[#6:4][#6:5][#6:6](-[#6:18])=[#6:7][#6:8][#6:9]-1[#6:2](-[#6:19])[#6:11] Any C10-carbocationHydrideShiftC9 --> C1=C2
Rationale: generic reaction involving a concerted hydride shift and nucleophilic attack
https://doi.org/10.1002/ange.202306020

### Slide 24
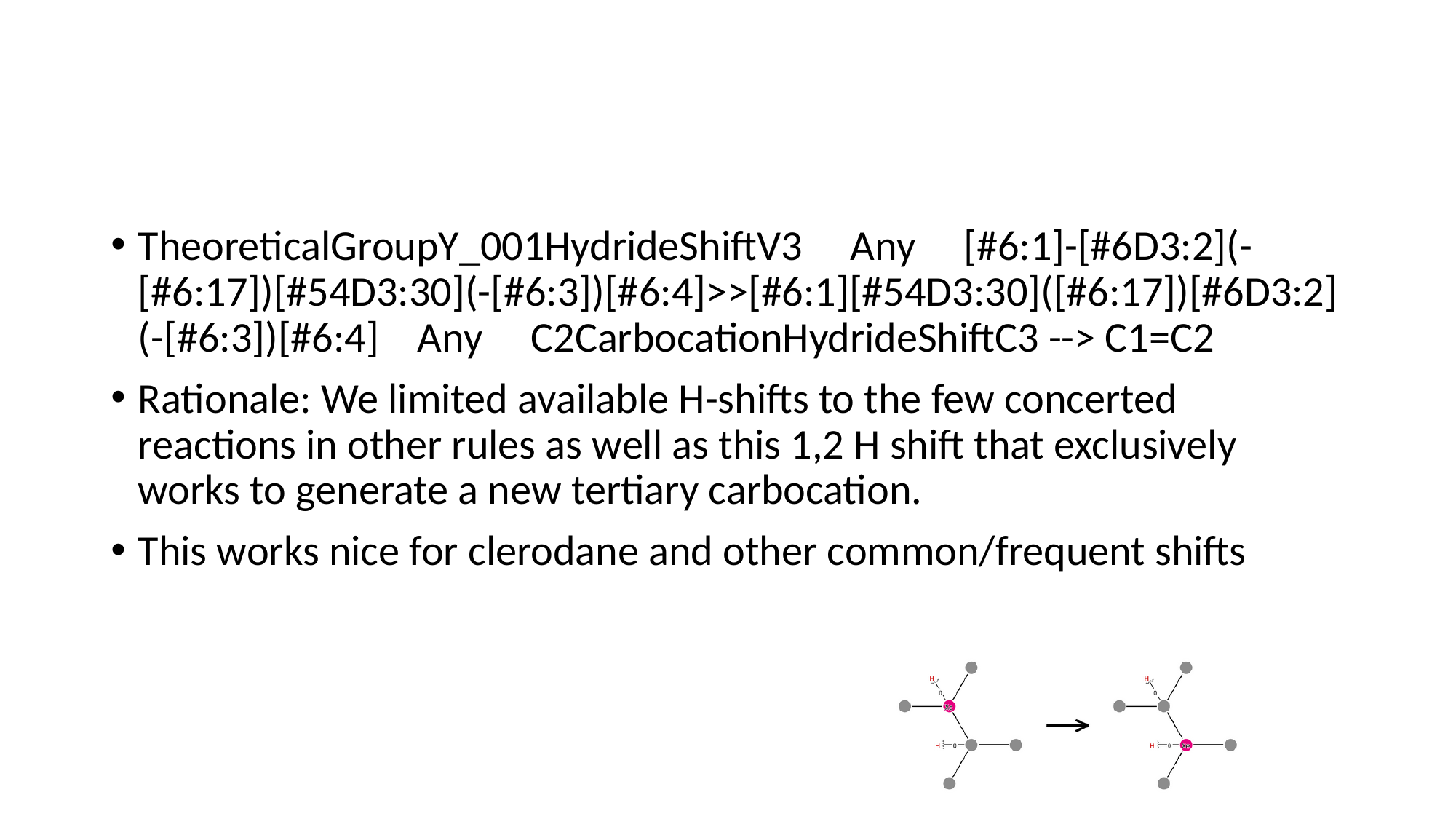

#
TheoreticalGroupY_001HydrideShiftV3 Any [#6:1]-[#6D3:2](-[#6:17])[#54D3:30](-[#6:3])[#6:4]>>[#6:1][#54D3:30]([#6:17])[#6D3:2](-[#6:3])[#6:4] Any C2CarbocationHydrideShiftC3 --> C1=C2
Rationale: We limited available H-shifts to the few concerted reactions in other rules as well as this 1,2 H shift that exclusively works to generate a new tertiary carbocation.
This works nice for clerodane and other common/frequent shifts

### Slide 25
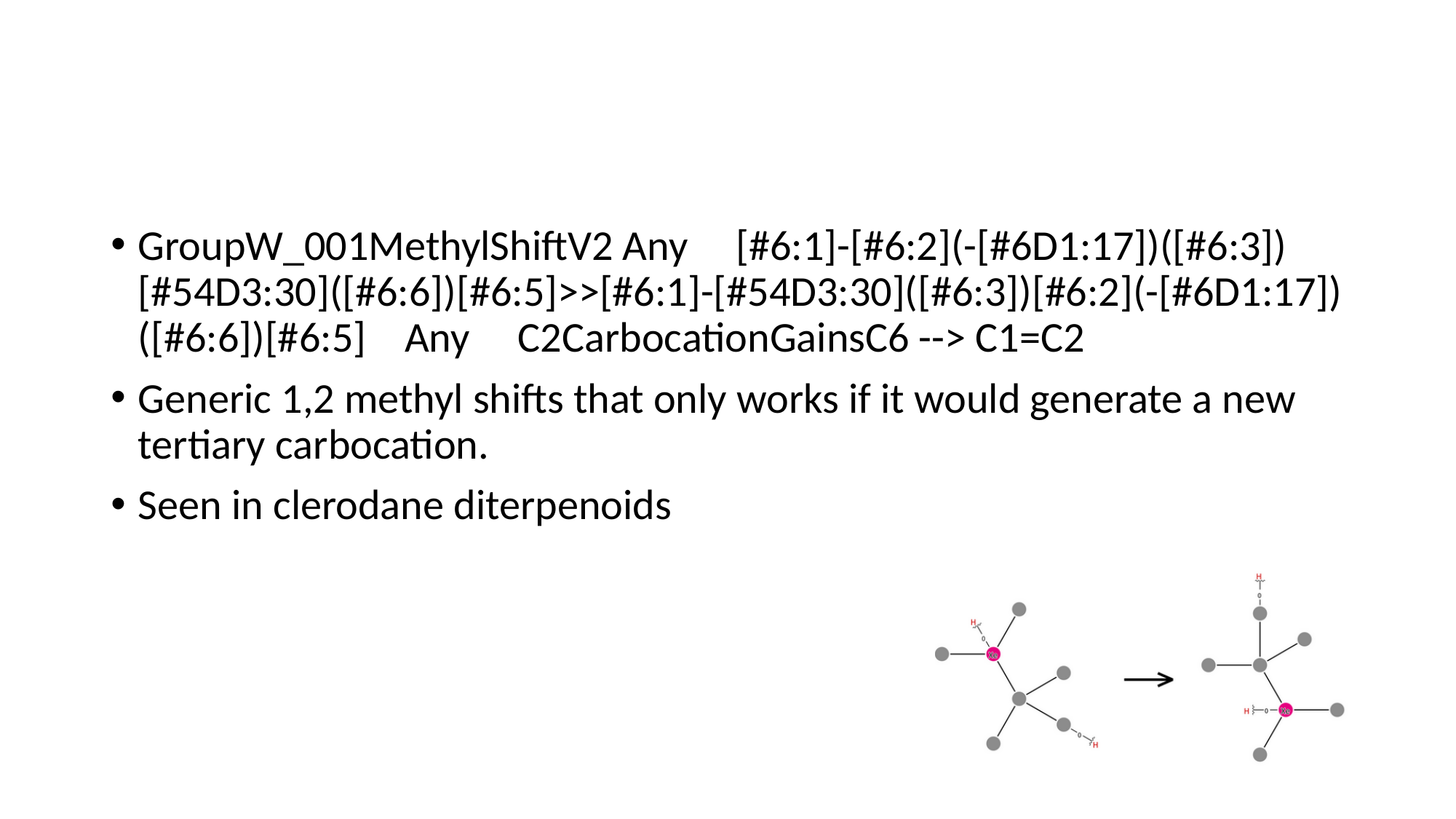

#
GroupW_001MethylShiftV2 Any [#6:1]-[#6:2](-[#6D1:17])([#6:3])[#54D3:30]([#6:6])[#6:5]>>[#6:1]-[#54D3:30]([#6:3])[#6:2](-[#6D1:17])([#6:6])[#6:5] Any C2CarbocationGainsC6 --> C1=C2
Generic 1,2 methyl shifts that only works if it would generate a new tertiary carbocation.
Seen in clerodane diterpenoids

### Slide 26
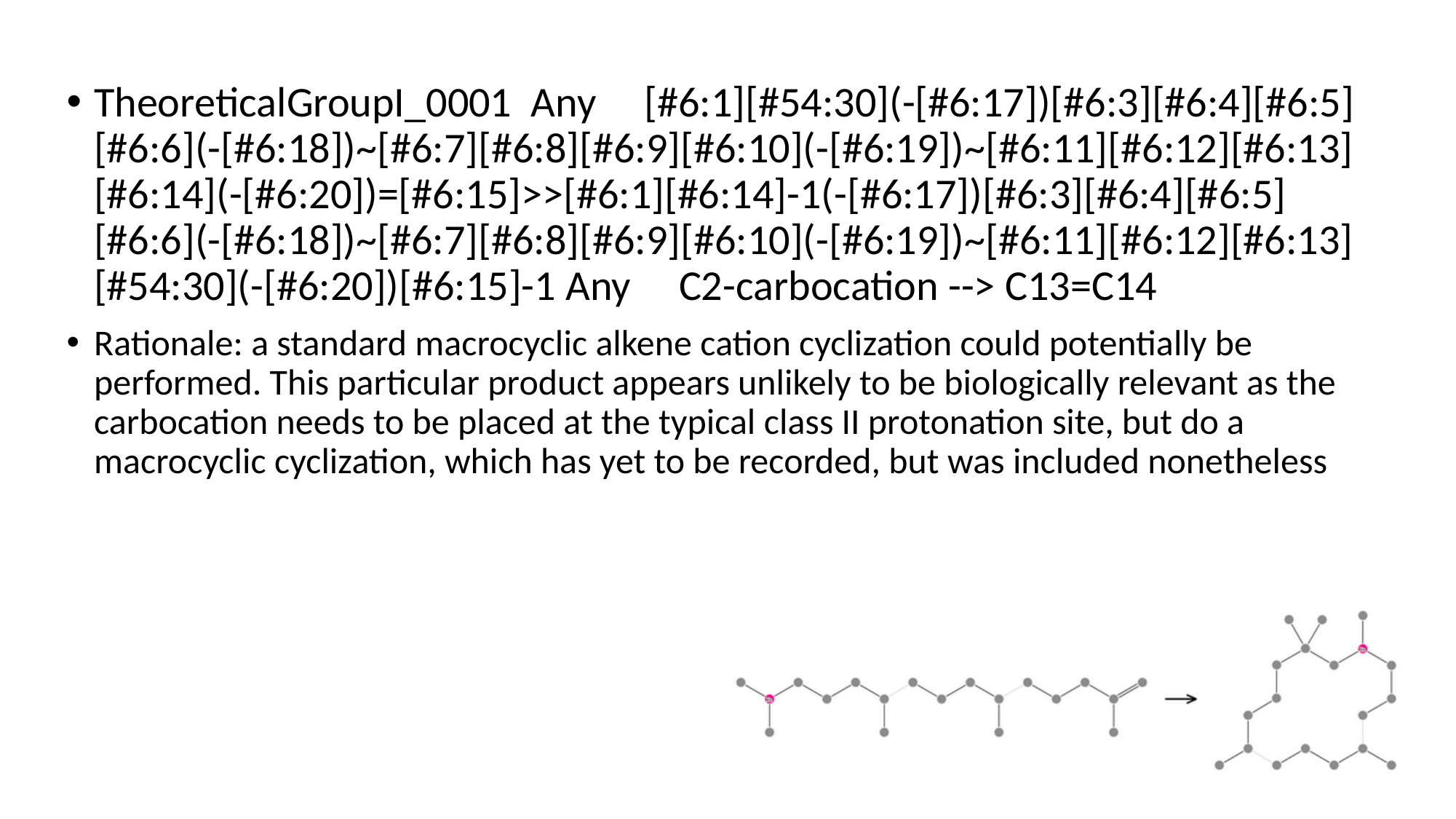

TheoreticalGroupI_0001 Any [#6:1][#54:30](-[#6:17])[#6:3][#6:4][#6:5][#6:6](-[#6:18])~[#6:7][#6:8][#6:9][#6:10](-[#6:19])~[#6:11][#6:12][#6:13][#6:14](-[#6:20])=[#6:15]>>[#6:1][#6:14]-1(-[#6:17])[#6:3][#6:4][#6:5][#6:6](-[#6:18])~[#6:7][#6:8][#6:9][#6:10](-[#6:19])~[#6:11][#6:12][#6:13][#54:30](-[#6:20])[#6:15]-1 Any C2-carbocation --> C13=C14
Rationale: a standard macrocyclic alkene cation cyclization could potentially be performed. This particular product appears unlikely to be biologically relevant as the carbocation needs to be placed at the typical class II protonation site, but do a macrocyclic cyclization, which has yet to be recorded, but was included nonetheless

### Slide 27
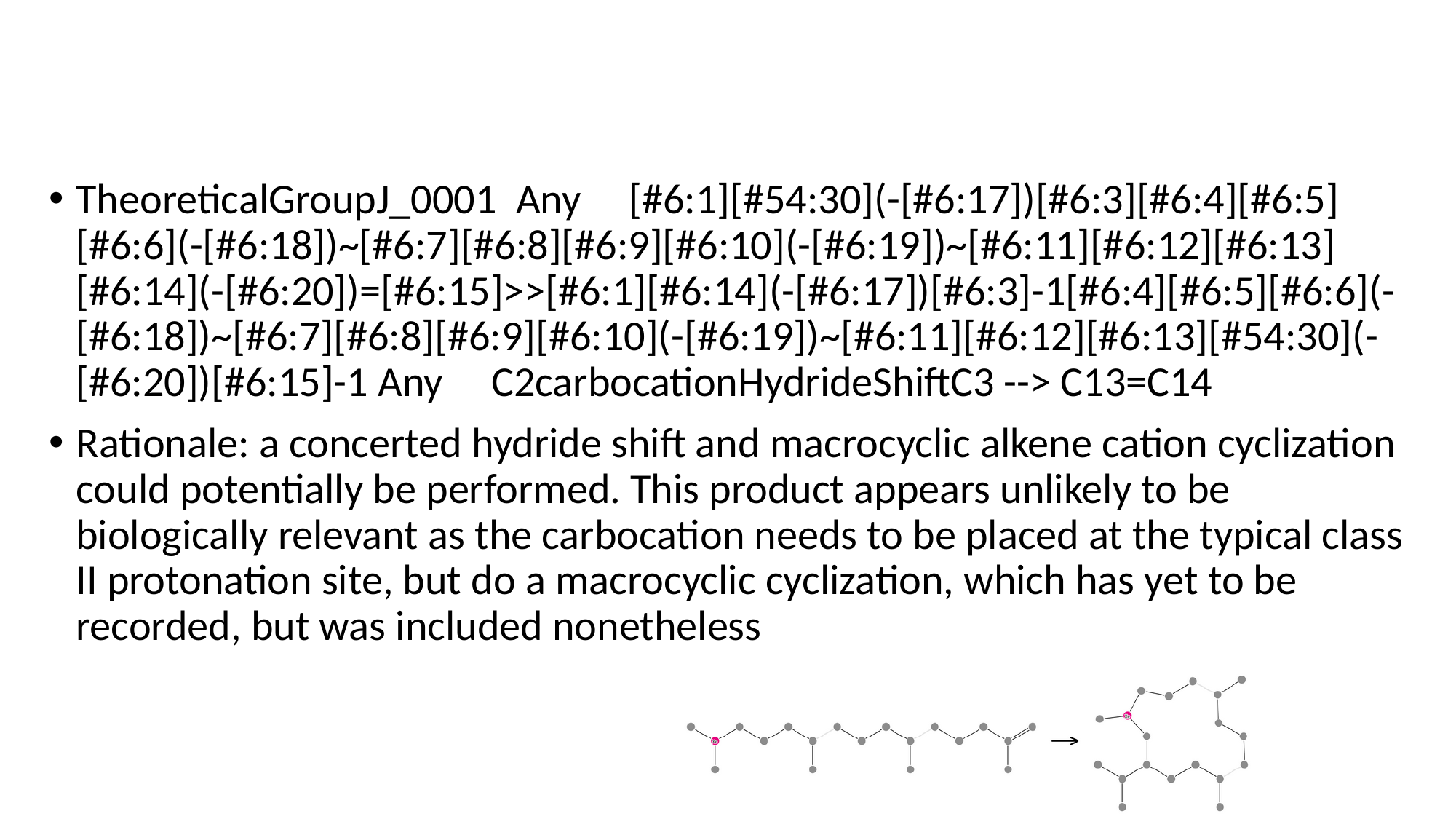

#
TheoreticalGroupJ_0001 Any [#6:1][#54:30](-[#6:17])[#6:3][#6:4][#6:5][#6:6](-[#6:18])~[#6:7][#6:8][#6:9][#6:10](-[#6:19])~[#6:11][#6:12][#6:13][#6:14](-[#6:20])=[#6:15]>>[#6:1][#6:14](-[#6:17])[#6:3]-1[#6:4][#6:5][#6:6](-[#6:18])~[#6:7][#6:8][#6:9][#6:10](-[#6:19])~[#6:11][#6:12][#6:13][#54:30](-[#6:20])[#6:15]-1 Any C2carbocationHydrideShiftC3 --> C13=C14
Rationale: a concerted hydride shift and macrocyclic alkene cation cyclization could potentially be performed. This product appears unlikely to be biologically relevant as the carbocation needs to be placed at the typical class II protonation site, but do a macrocyclic cyclization, which has yet to be recorded, but was included nonetheless

### Slide 28
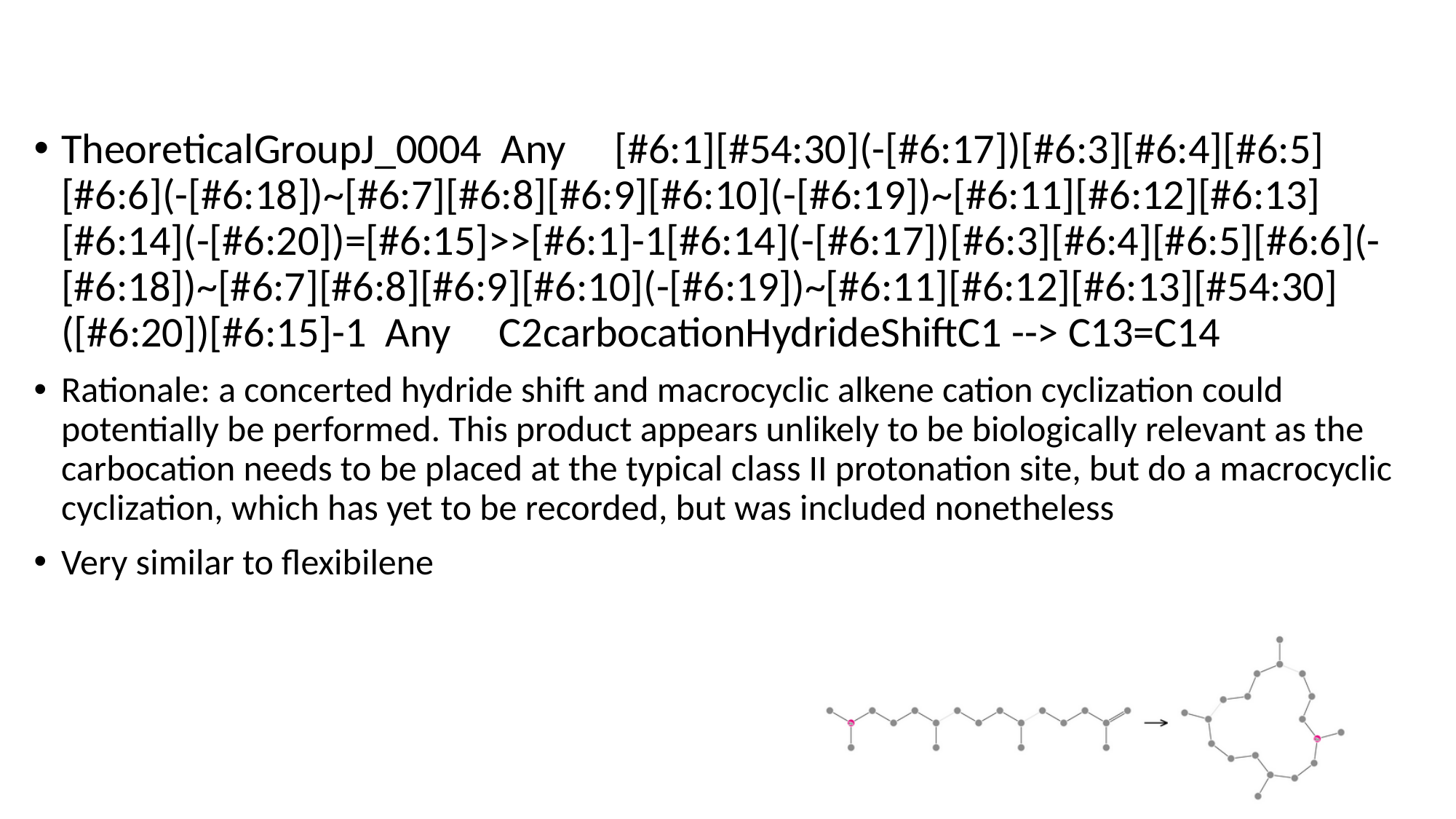

TheoreticalGroupJ_0004 Any [#6:1][#54:30](-[#6:17])[#6:3][#6:4][#6:5][#6:6](-[#6:18])~[#6:7][#6:8][#6:9][#6:10](-[#6:19])~[#6:11][#6:12][#6:13][#6:14](-[#6:20])=[#6:15]>>[#6:1]-1[#6:14](-[#6:17])[#6:3][#6:4][#6:5][#6:6](-[#6:18])~[#6:7][#6:8][#6:9][#6:10](-[#6:19])~[#6:11][#6:12][#6:13][#54:30]([#6:20])[#6:15]-1 Any C2carbocationHydrideShiftC1 --> C13=C14
Rationale: a concerted hydride shift and macrocyclic alkene cation cyclization could potentially be performed. This product appears unlikely to be biologically relevant as the carbocation needs to be placed at the typical class II protonation site, but do a macrocyclic cyclization, which has yet to be recorded, but was included nonetheless
Very similar to flexibilene

### Slide 29
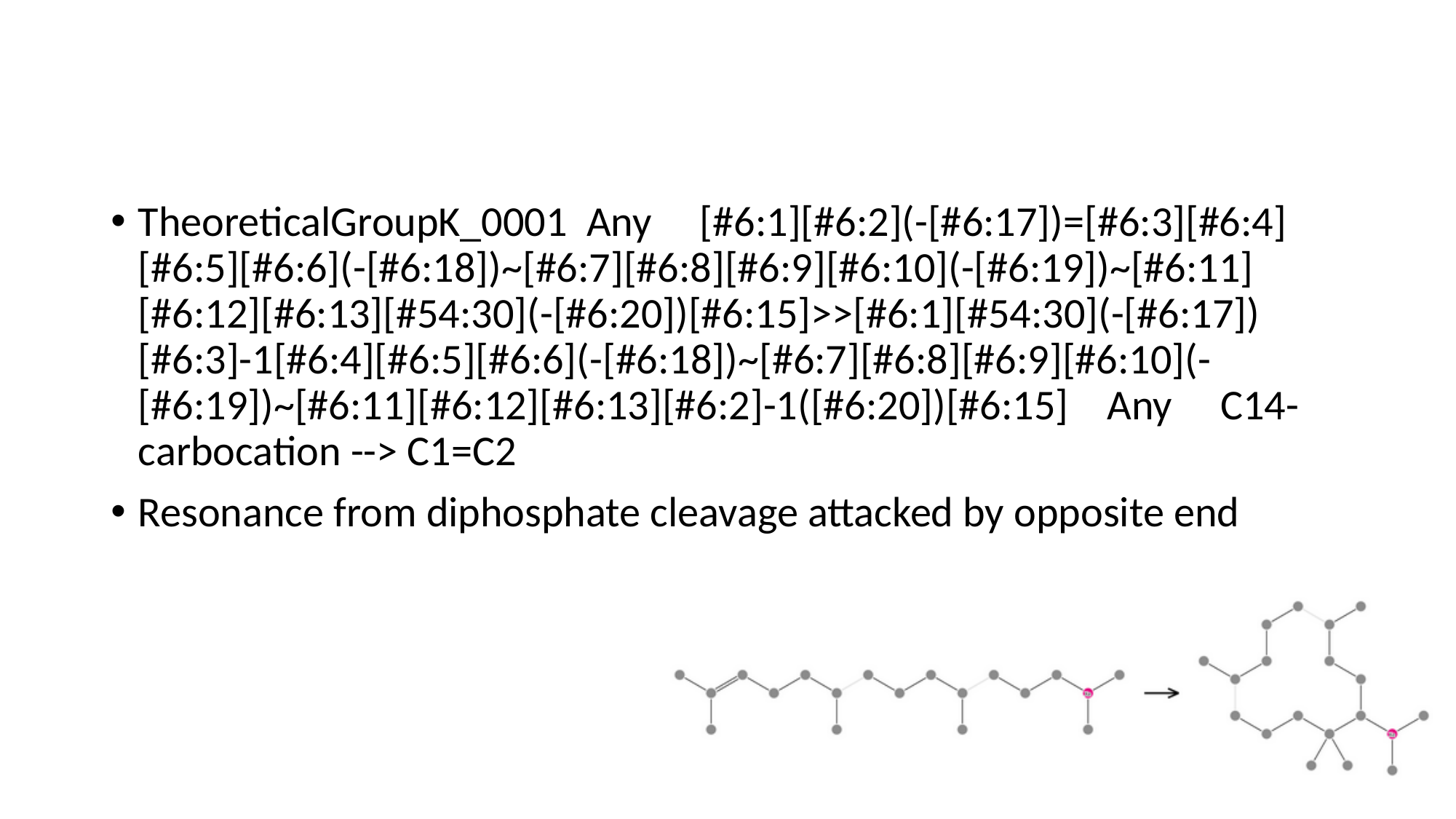

#
TheoreticalGroupK_0001 Any [#6:1][#6:2](-[#6:17])=[#6:3][#6:4][#6:5][#6:6](-[#6:18])~[#6:7][#6:8][#6:9][#6:10](-[#6:19])~[#6:11][#6:12][#6:13][#54:30](-[#6:20])[#6:15]>>[#6:1][#54:30](-[#6:17])[#6:3]-1[#6:4][#6:5][#6:6](-[#6:18])~[#6:7][#6:8][#6:9][#6:10](-[#6:19])~[#6:11][#6:12][#6:13][#6:2]-1([#6:20])[#6:15] Any C14-carbocation --> C1=C2
Resonance from diphosphate cleavage attacked by opposite end

### Slide 30
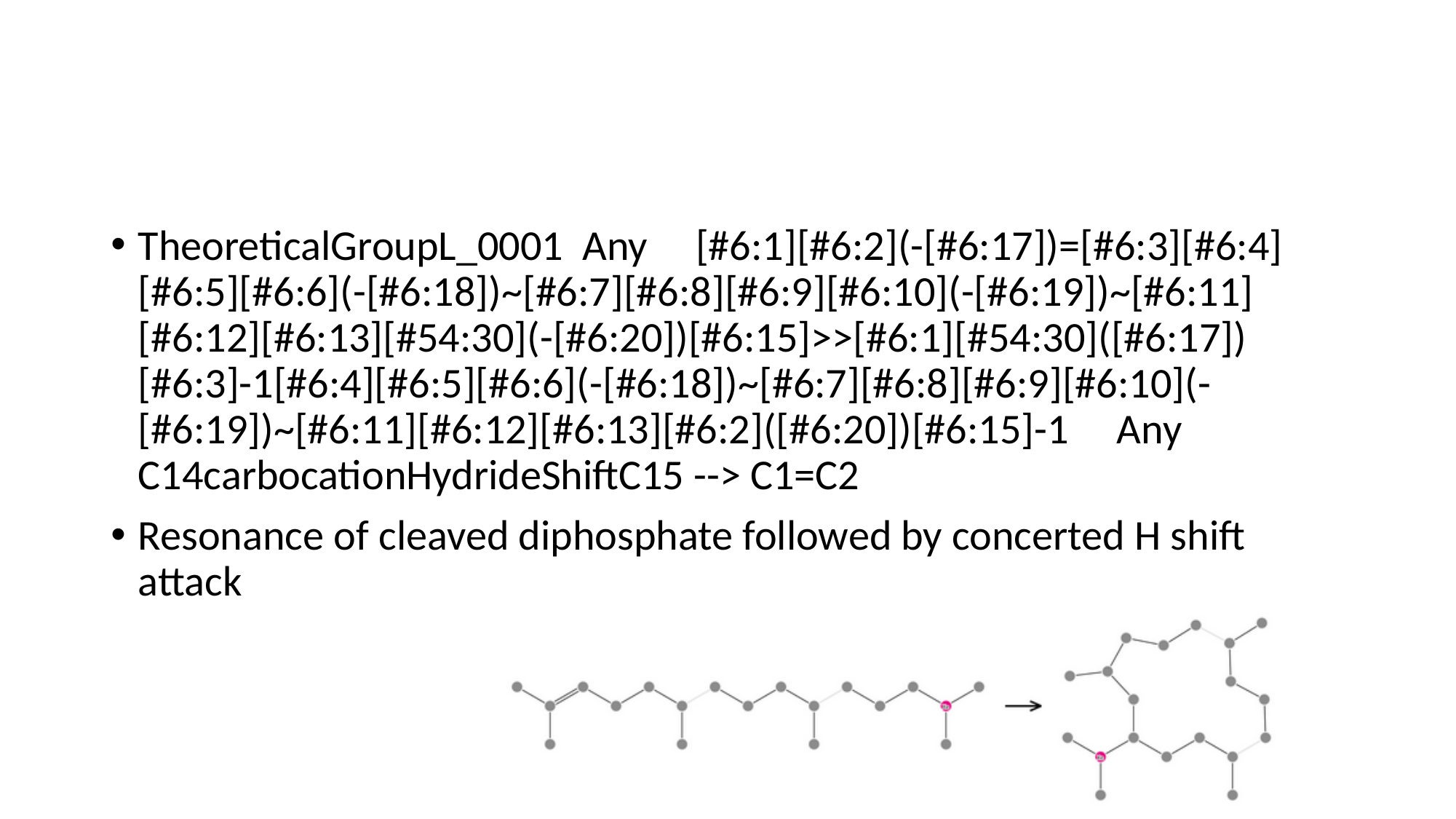

#
TheoreticalGroupL_0001 Any [#6:1][#6:2](-[#6:17])=[#6:3][#6:4][#6:5][#6:6](-[#6:18])~[#6:7][#6:8][#6:9][#6:10](-[#6:19])~[#6:11][#6:12][#6:13][#54:30](-[#6:20])[#6:15]>>[#6:1][#54:30]([#6:17])[#6:3]-1[#6:4][#6:5][#6:6](-[#6:18])~[#6:7][#6:8][#6:9][#6:10](-[#6:19])~[#6:11][#6:12][#6:13][#6:2]([#6:20])[#6:15]-1 Any C14carbocationHydrideShiftC15 --> C1=C2
Resonance of cleaved diphosphate followed by concerted H shift attack

### Slide 31
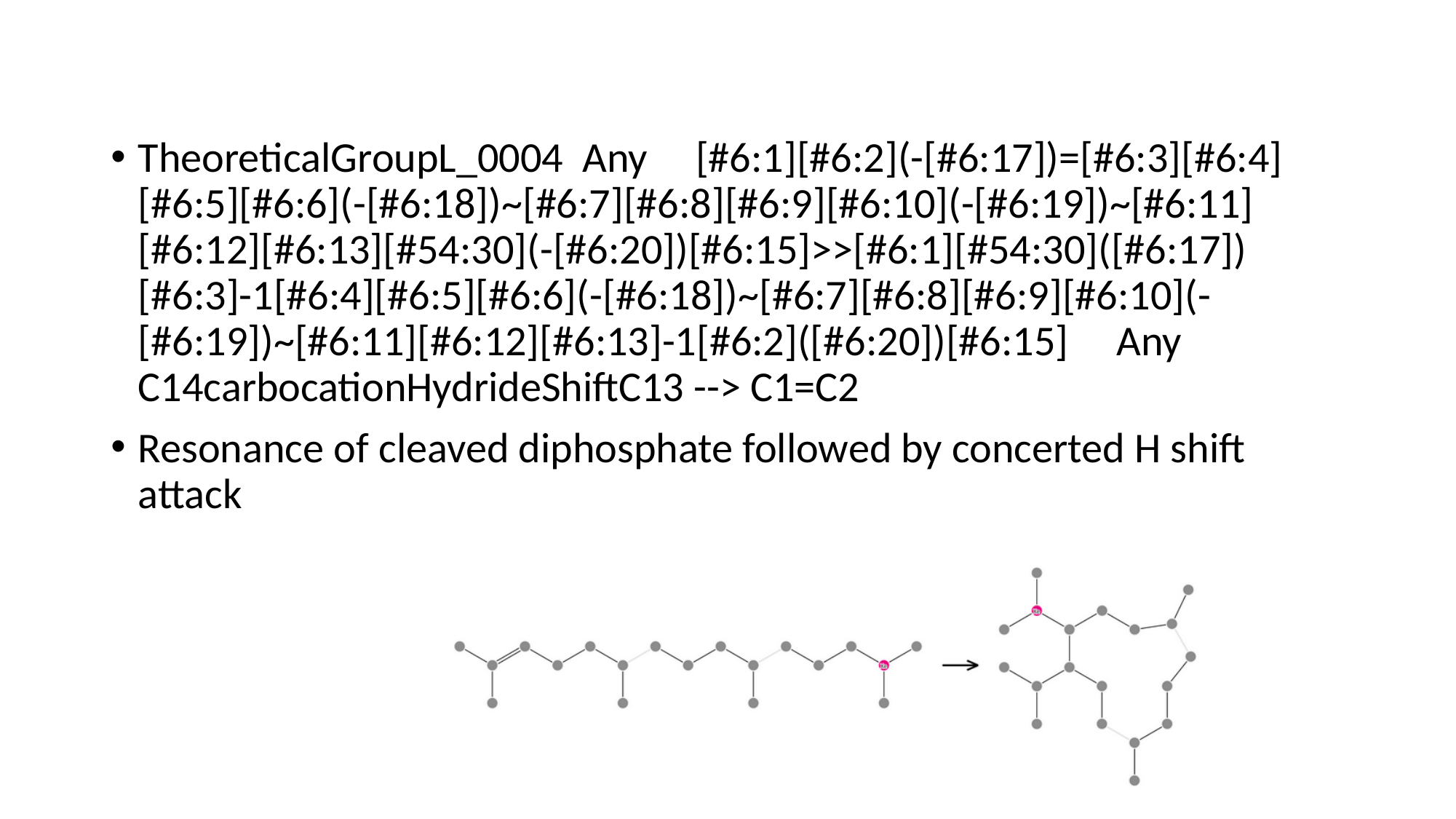

#
TheoreticalGroupL_0004 Any [#6:1][#6:2](-[#6:17])=[#6:3][#6:4][#6:5][#6:6](-[#6:18])~[#6:7][#6:8][#6:9][#6:10](-[#6:19])~[#6:11][#6:12][#6:13][#54:30](-[#6:20])[#6:15]>>[#6:1][#54:30]([#6:17])[#6:3]-1[#6:4][#6:5][#6:6](-[#6:18])~[#6:7][#6:8][#6:9][#6:10](-[#6:19])~[#6:11][#6:12][#6:13]-1[#6:2]([#6:20])[#6:15] Any C14carbocationHydrideShiftC13 --> C1=C2
Resonance of cleaved diphosphate followed by concerted H shift attack

### Slide 32
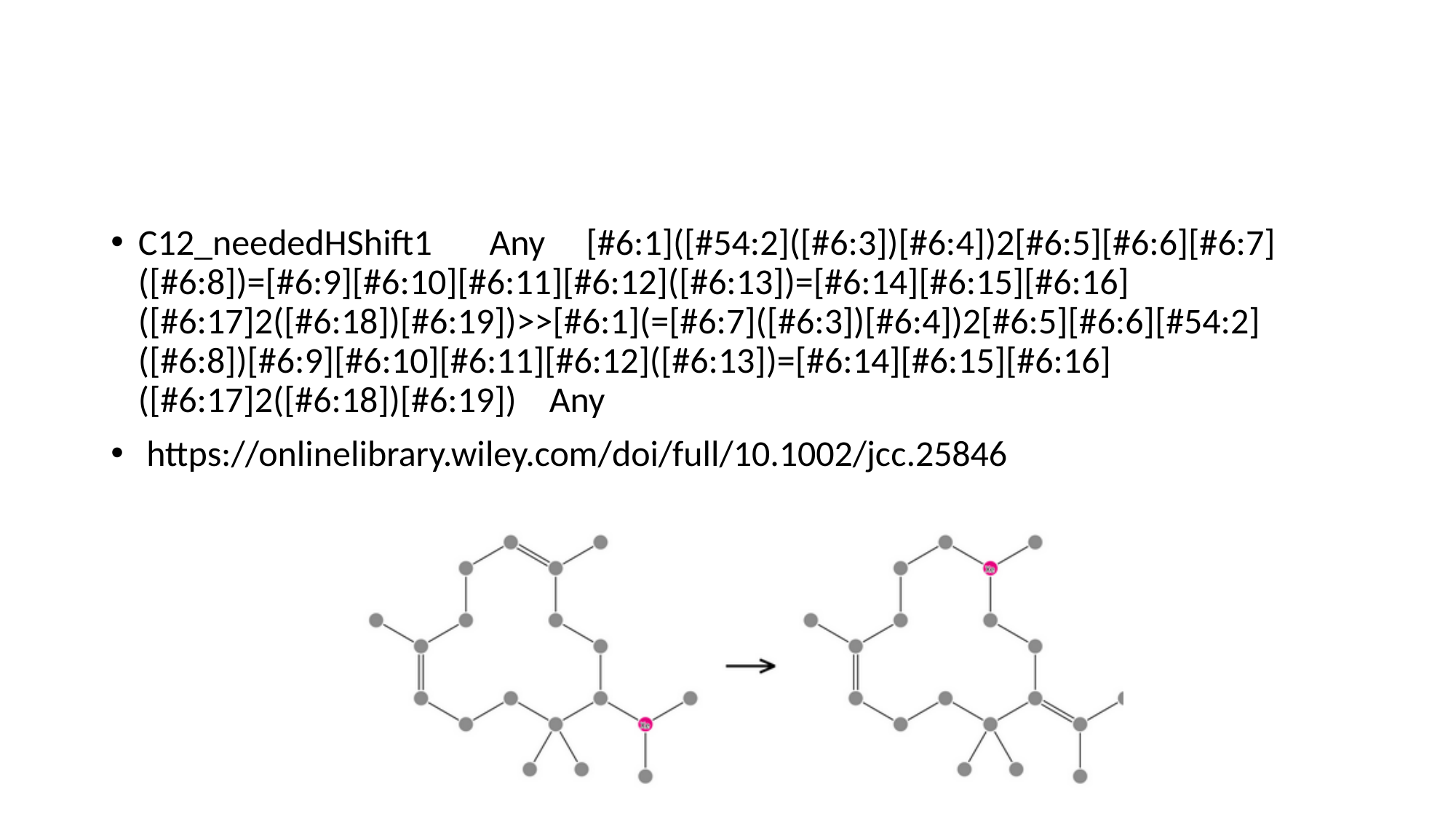

#
C12_neededHShift1 Any [#6:1]([#54:2]([#6:3])[#6:4])2[#6:5][#6:6][#6:7]([#6:8])=[#6:9][#6:10][#6:11][#6:12]([#6:13])=[#6:14][#6:15][#6:16]([#6:17]2([#6:18])[#6:19])>>[#6:1](=[#6:7]([#6:3])[#6:4])2[#6:5][#6:6][#54:2]([#6:8])[#6:9][#6:10][#6:11][#6:12]([#6:13])=[#6:14][#6:15][#6:16]([#6:17]2([#6:18])[#6:19]) Any
 https://onlinelibrary.wiley.com/doi/full/10.1002/jcc.25846

### Slide 33
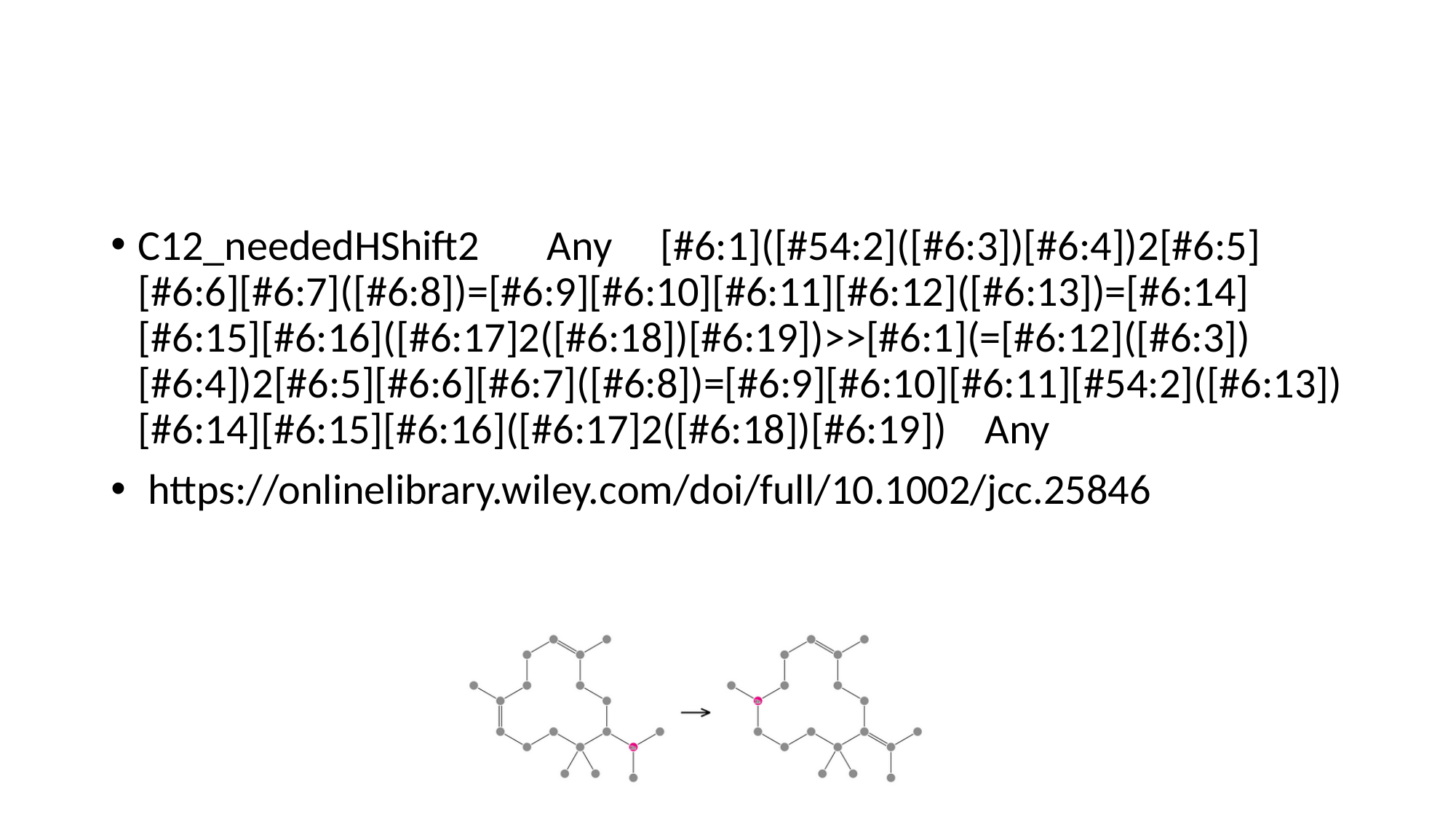

#
C12_neededHShift2 Any [#6:1]([#54:2]([#6:3])[#6:4])2[#6:5][#6:6][#6:7]([#6:8])=[#6:9][#6:10][#6:11][#6:12]([#6:13])=[#6:14][#6:15][#6:16]([#6:17]2([#6:18])[#6:19])>>[#6:1](=[#6:12]([#6:3])[#6:4])2[#6:5][#6:6][#6:7]([#6:8])=[#6:9][#6:10][#6:11][#54:2]([#6:13])[#6:14][#6:15][#6:16]([#6:17]2([#6:18])[#6:19]) Any
 https://onlinelibrary.wiley.com/doi/full/10.1002/jcc.25846

### Slide 34
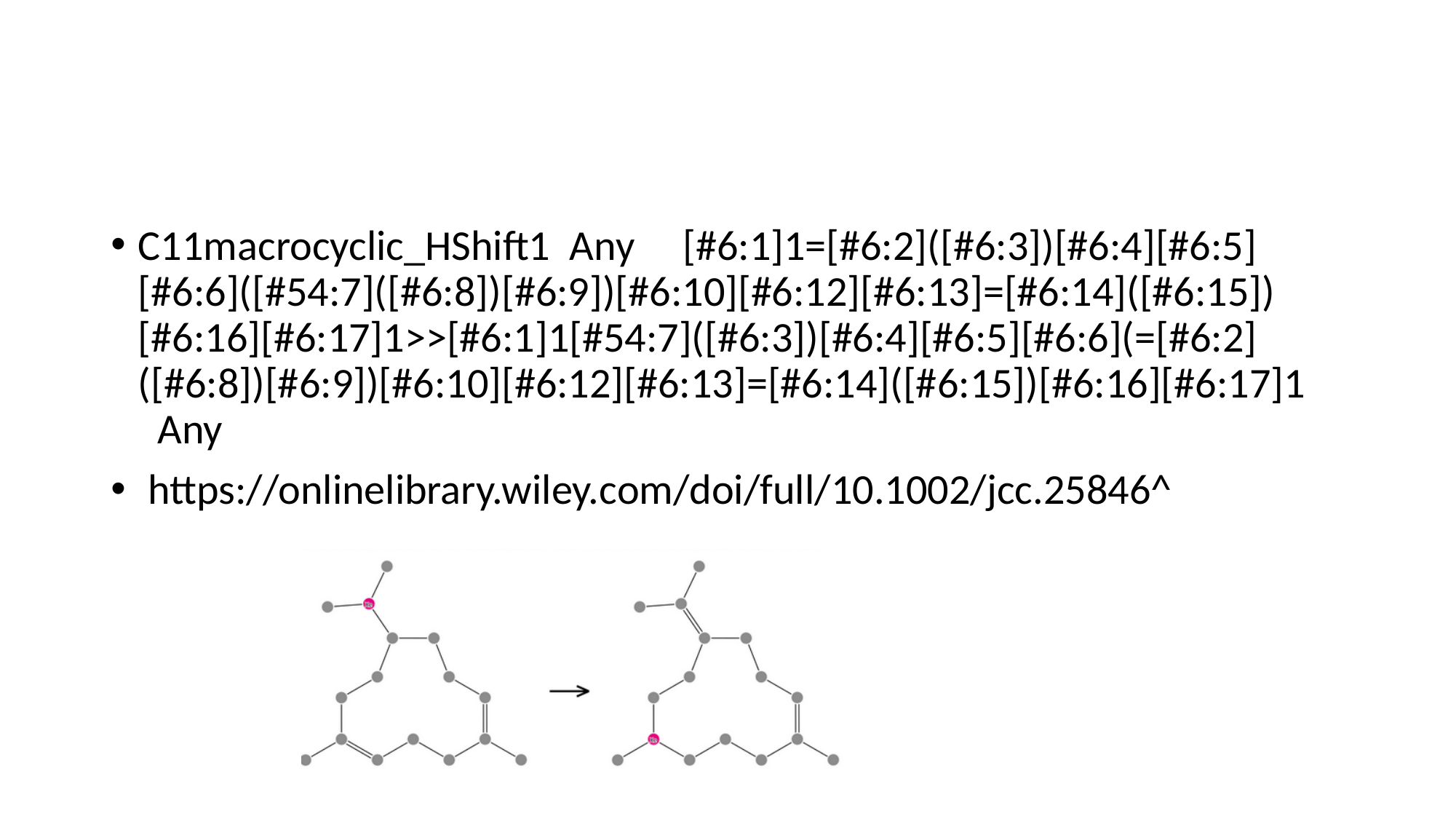

#
C11macrocyclic_HShift1 Any [#6:1]1=[#6:2]([#6:3])[#6:4][#6:5][#6:6]([#54:7]([#6:8])[#6:9])[#6:10][#6:12][#6:13]=[#6:14]([#6:15])[#6:16][#6:17]1>>[#6:1]1[#54:7]([#6:3])[#6:4][#6:5][#6:6](=[#6:2]([#6:8])[#6:9])[#6:10][#6:12][#6:13]=[#6:14]([#6:15])[#6:16][#6:17]1 Any
 https://onlinelibrary.wiley.com/doi/full/10.1002/jcc.25846^

### Slide 35
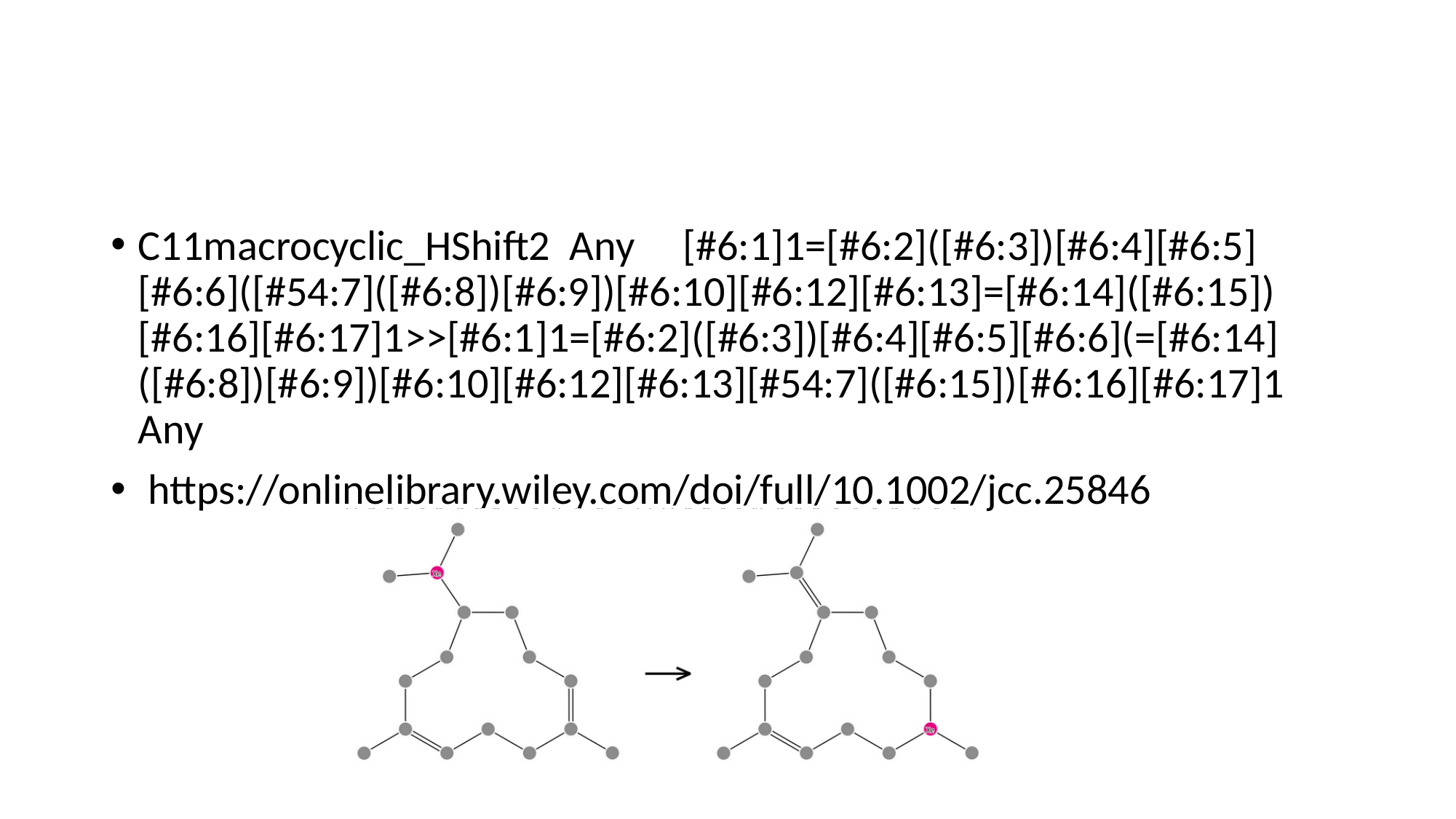

#
C11macrocyclic_HShift2 Any [#6:1]1=[#6:2]([#6:3])[#6:4][#6:5][#6:6]([#54:7]([#6:8])[#6:9])[#6:10][#6:12][#6:13]=[#6:14]([#6:15])[#6:16][#6:17]1>>[#6:1]1=[#6:2]([#6:3])[#6:4][#6:5][#6:6](=[#6:14]([#6:8])[#6:9])[#6:10][#6:12][#6:13][#54:7]([#6:15])[#6:16][#6:17]1 Any
 https://onlinelibrary.wiley.com/doi/full/10.1002/jcc.25846

### Slide 36
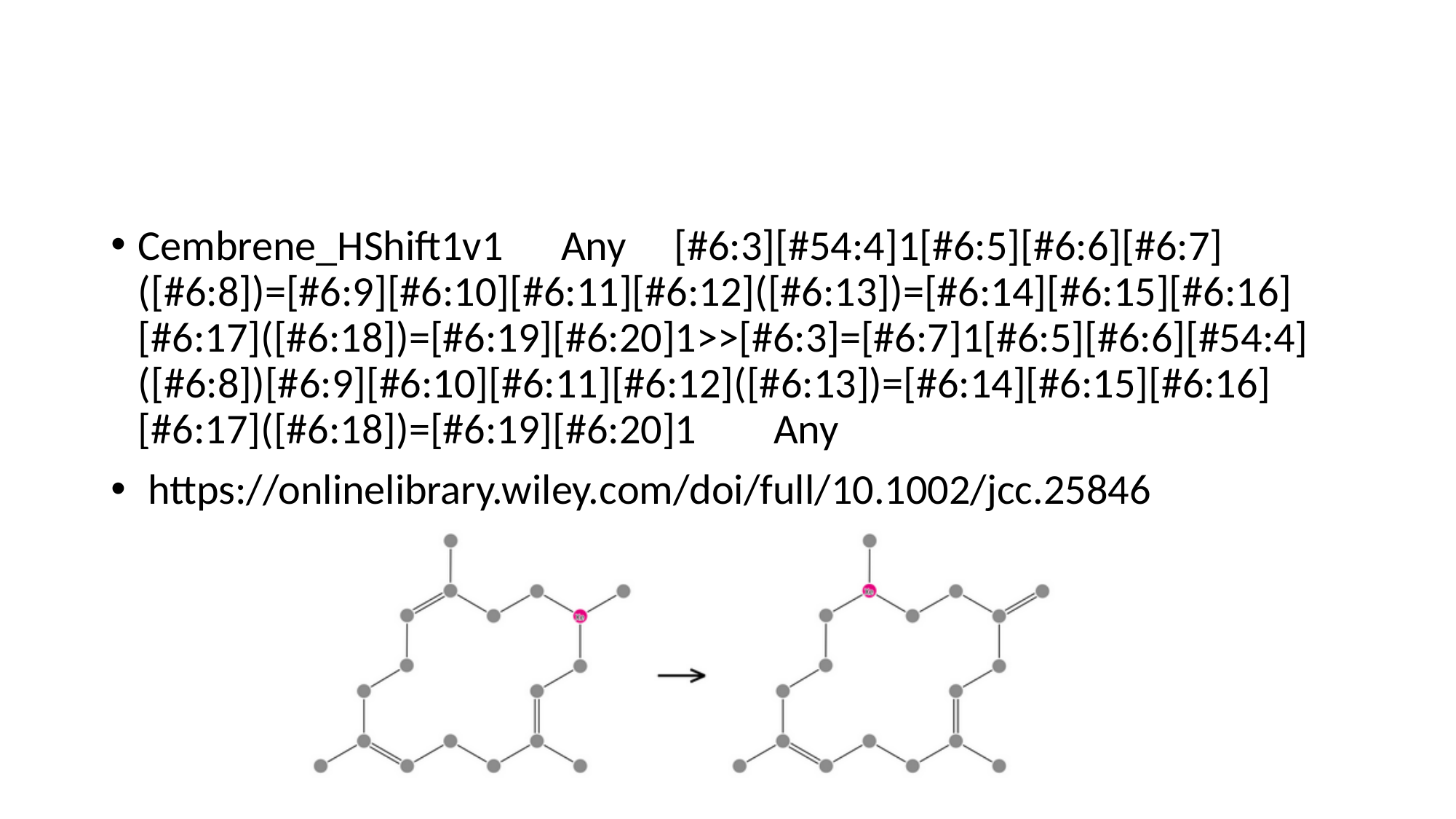

#
Cembrene_HShift1v1 Any [#6:3][#54:4]1[#6:5][#6:6][#6:7]([#6:8])=[#6:9][#6:10][#6:11][#6:12]([#6:13])=[#6:14][#6:15][#6:16][#6:17]([#6:18])=[#6:19][#6:20]1>>[#6:3]=[#6:7]1[#6:5][#6:6][#54:4]([#6:8])[#6:9][#6:10][#6:11][#6:12]([#6:13])=[#6:14][#6:15][#6:16][#6:17]([#6:18])=[#6:19][#6:20]1 Any
 https://onlinelibrary.wiley.com/doi/full/10.1002/jcc.25846

### Slide 37
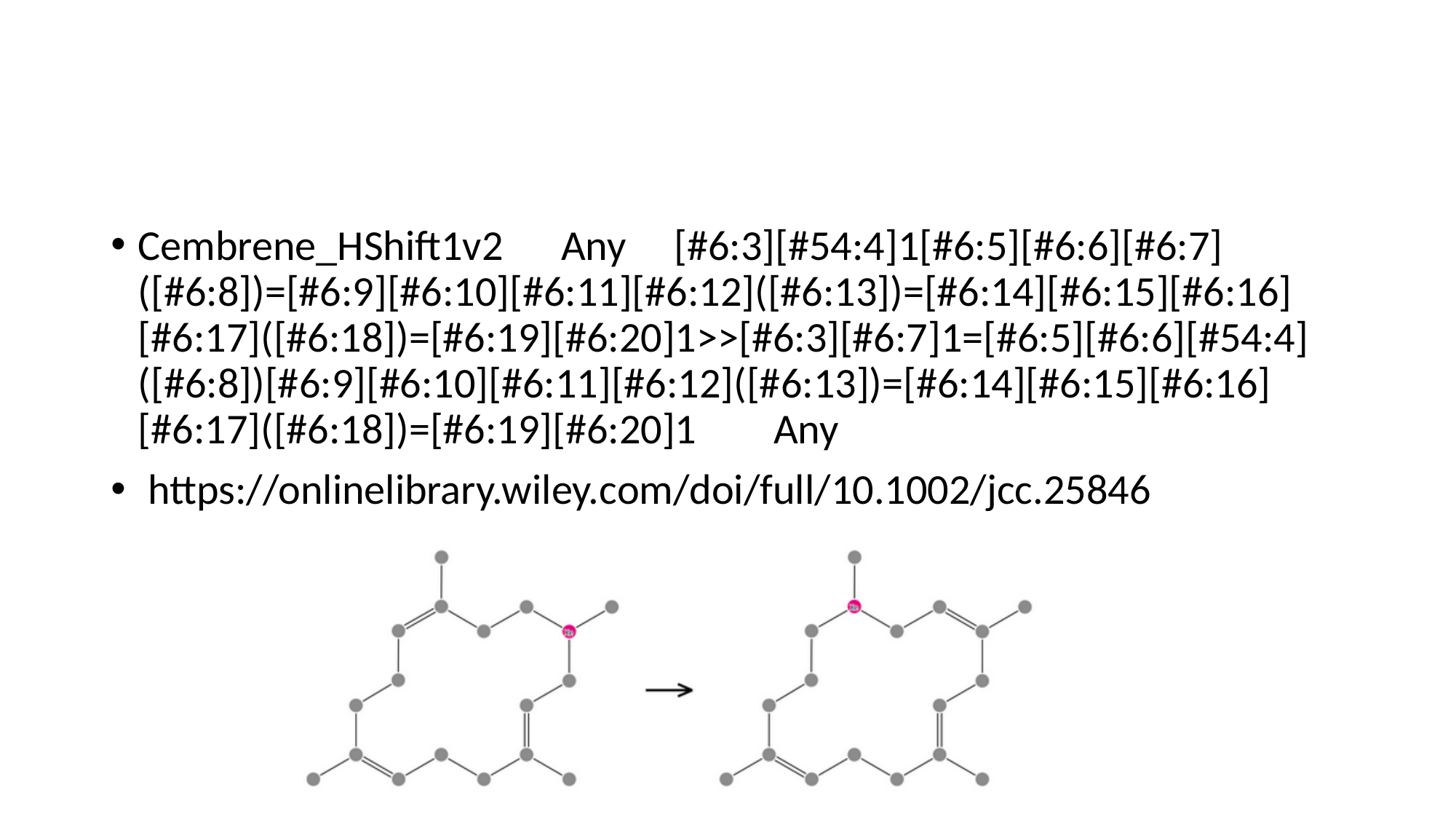

#
Cembrene_HShift1v2 Any [#6:3][#54:4]1[#6:5][#6:6][#6:7]([#6:8])=[#6:9][#6:10][#6:11][#6:12]([#6:13])=[#6:14][#6:15][#6:16][#6:17]([#6:18])=[#6:19][#6:20]1>>[#6:3][#6:7]1=[#6:5][#6:6][#54:4]([#6:8])[#6:9][#6:10][#6:11][#6:12]([#6:13])=[#6:14][#6:15][#6:16][#6:17]([#6:18])=[#6:19][#6:20]1 Any
 https://onlinelibrary.wiley.com/doi/full/10.1002/jcc.25846

### Slide 38
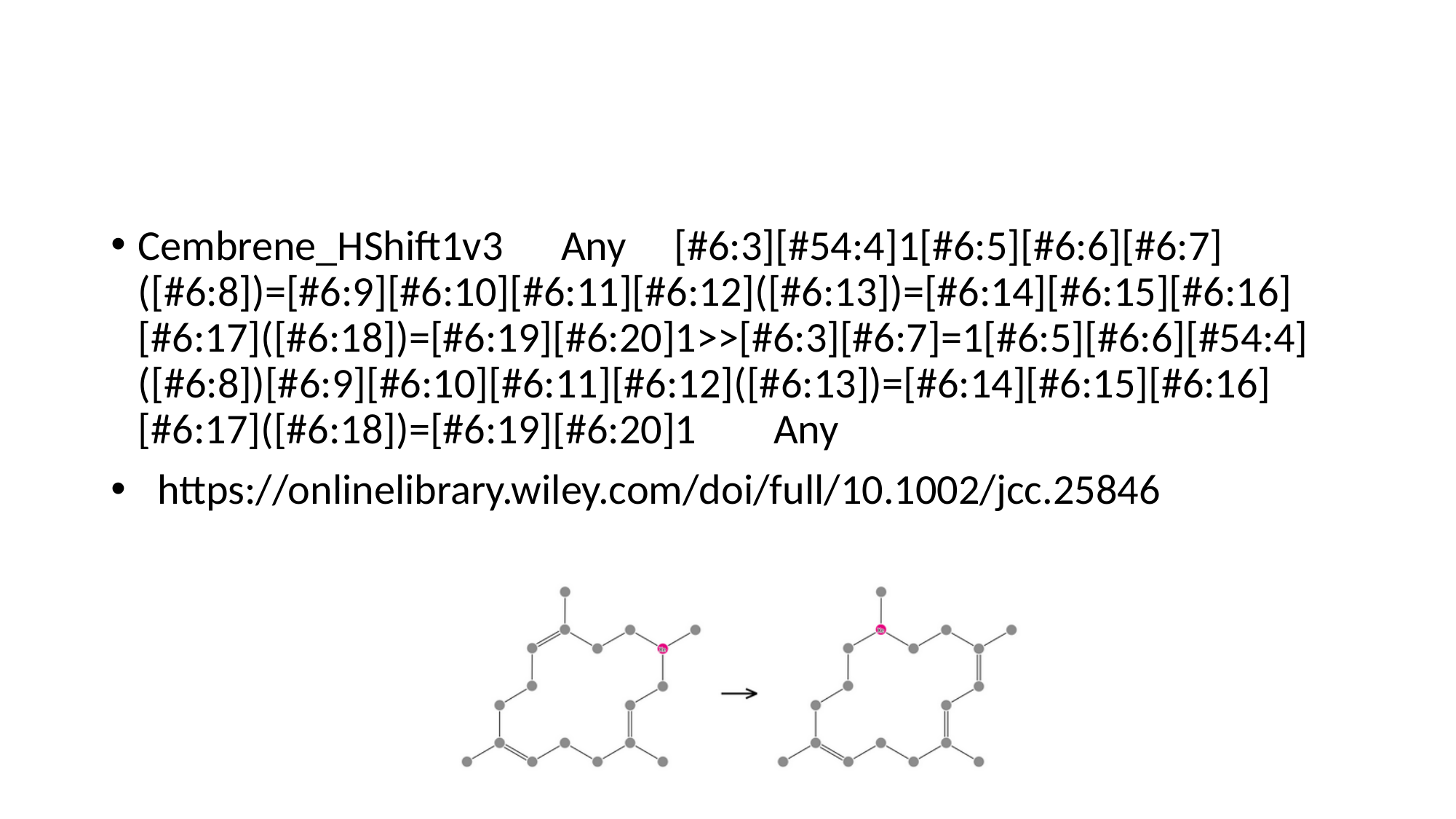

#
Cembrene_HShift1v3 Any [#6:3][#54:4]1[#6:5][#6:6][#6:7]([#6:8])=[#6:9][#6:10][#6:11][#6:12]([#6:13])=[#6:14][#6:15][#6:16][#6:17]([#6:18])=[#6:19][#6:20]1>>[#6:3][#6:7]=1[#6:5][#6:6][#54:4]([#6:8])[#6:9][#6:10][#6:11][#6:12]([#6:13])=[#6:14][#6:15][#6:16][#6:17]([#6:18])=[#6:19][#6:20]1 Any
 https://onlinelibrary.wiley.com/doi/full/10.1002/jcc.25846

### Slide 39
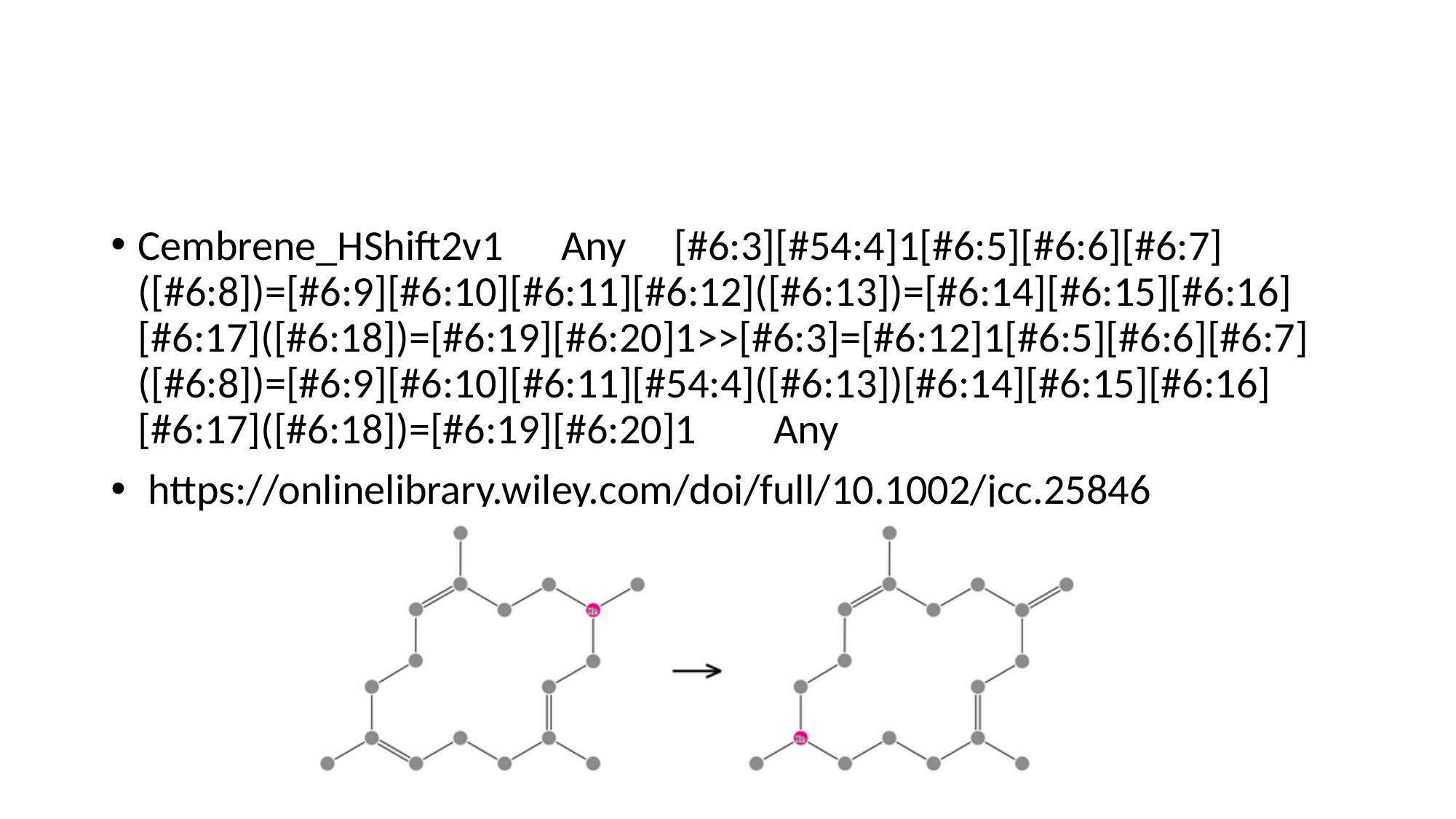

#
Cembrene_HShift2v1 Any [#6:3][#54:4]1[#6:5][#6:6][#6:7]([#6:8])=[#6:9][#6:10][#6:11][#6:12]([#6:13])=[#6:14][#6:15][#6:16][#6:17]([#6:18])=[#6:19][#6:20]1>>[#6:3]=[#6:12]1[#6:5][#6:6][#6:7]([#6:8])=[#6:9][#6:10][#6:11][#54:4]([#6:13])[#6:14][#6:15][#6:16][#6:17]([#6:18])=[#6:19][#6:20]1 Any
 https://onlinelibrary.wiley.com/doi/full/10.1002/jcc.25846

### Slide 40
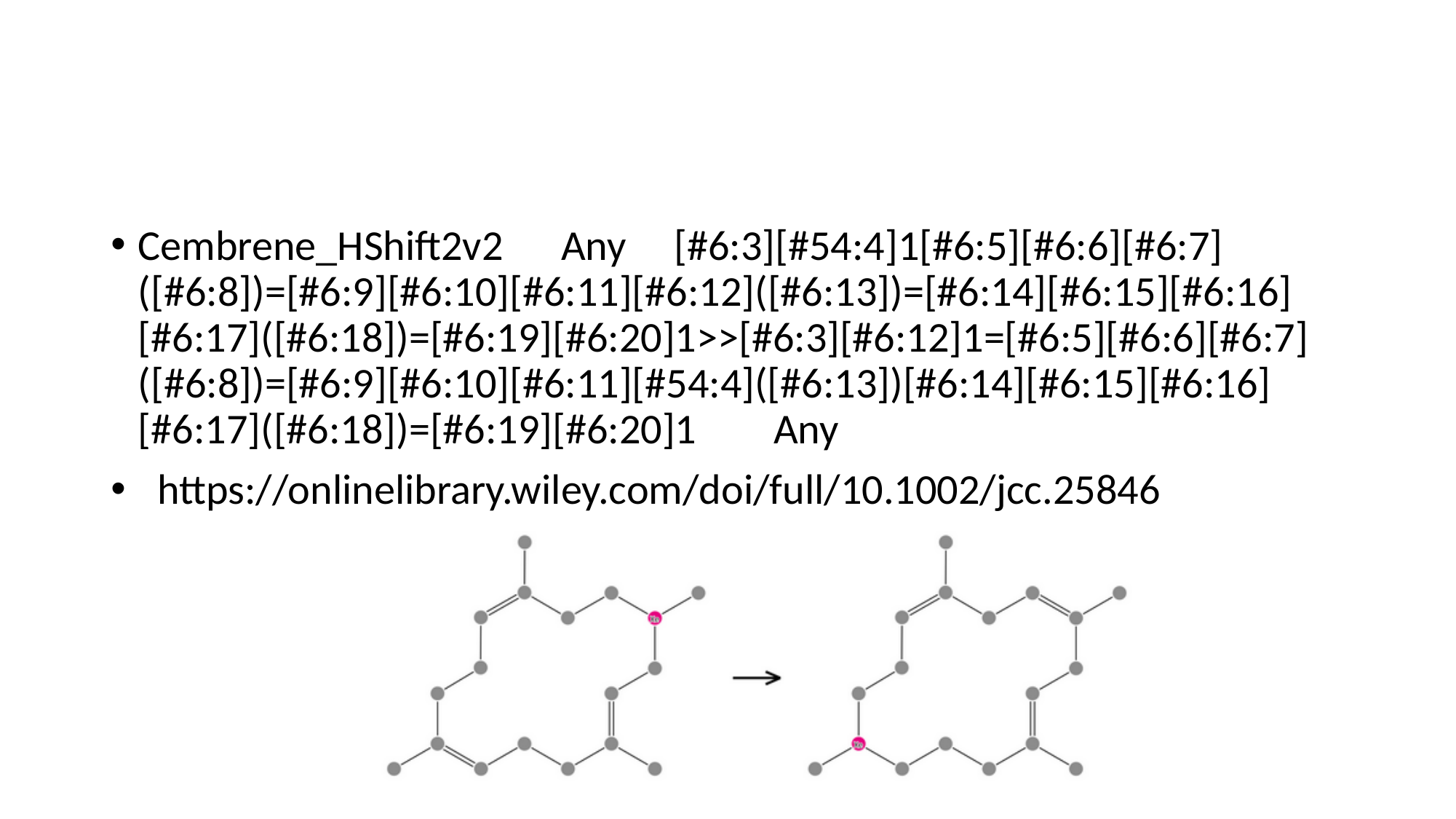

#
Cembrene_HShift2v2 Any [#6:3][#54:4]1[#6:5][#6:6][#6:7]([#6:8])=[#6:9][#6:10][#6:11][#6:12]([#6:13])=[#6:14][#6:15][#6:16][#6:17]([#6:18])=[#6:19][#6:20]1>>[#6:3][#6:12]1=[#6:5][#6:6][#6:7]([#6:8])=[#6:9][#6:10][#6:11][#54:4]([#6:13])[#6:14][#6:15][#6:16][#6:17]([#6:18])=[#6:19][#6:20]1 Any
 https://onlinelibrary.wiley.com/doi/full/10.1002/jcc.25846

### Slide 41
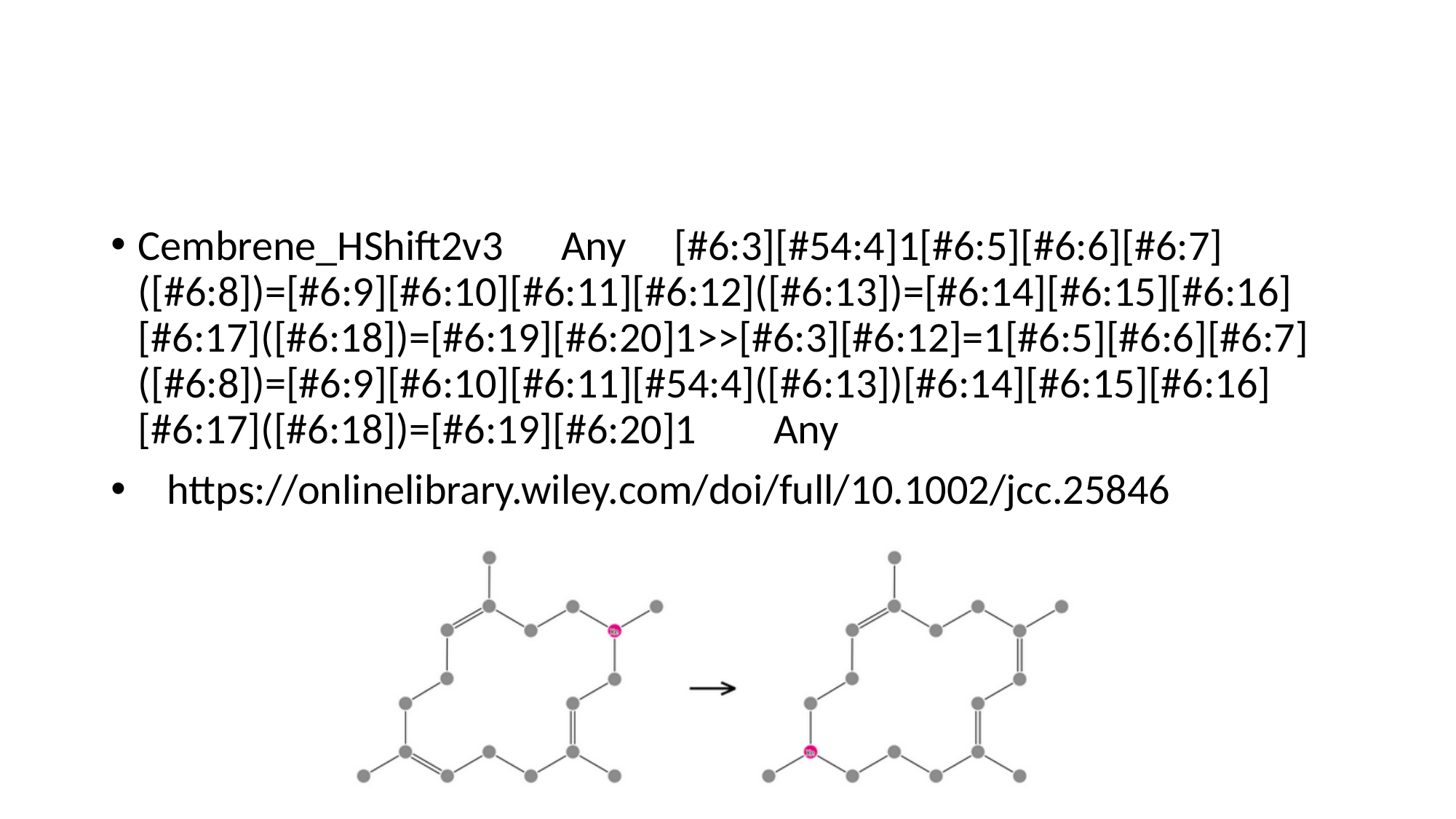

#
Cembrene_HShift2v3 Any [#6:3][#54:4]1[#6:5][#6:6][#6:7]([#6:8])=[#6:9][#6:10][#6:11][#6:12]([#6:13])=[#6:14][#6:15][#6:16][#6:17]([#6:18])=[#6:19][#6:20]1>>[#6:3][#6:12]=1[#6:5][#6:6][#6:7]([#6:8])=[#6:9][#6:10][#6:11][#54:4]([#6:13])[#6:14][#6:15][#6:16][#6:17]([#6:18])=[#6:19][#6:20]1 Any
 https://onlinelibrary.wiley.com/doi/full/10.1002/jcc.25846

### Slide 42
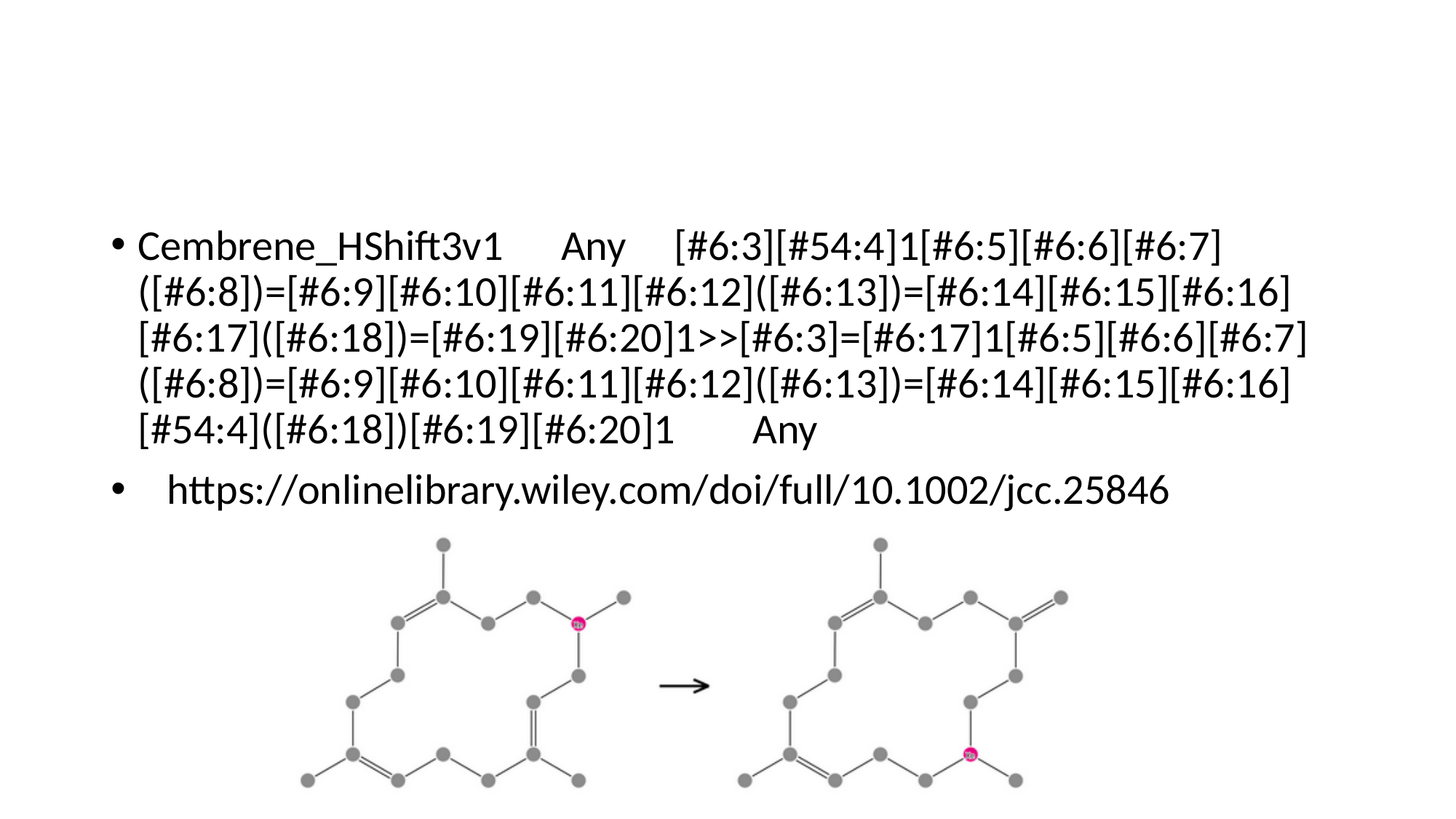

#
Cembrene_HShift3v1 Any [#6:3][#54:4]1[#6:5][#6:6][#6:7]([#6:8])=[#6:9][#6:10][#6:11][#6:12]([#6:13])=[#6:14][#6:15][#6:16][#6:17]([#6:18])=[#6:19][#6:20]1>>[#6:3]=[#6:17]1[#6:5][#6:6][#6:7]([#6:8])=[#6:9][#6:10][#6:11][#6:12]([#6:13])=[#6:14][#6:15][#6:16][#54:4]([#6:18])[#6:19][#6:20]1 Any
 https://onlinelibrary.wiley.com/doi/full/10.1002/jcc.25846

### Slide 43
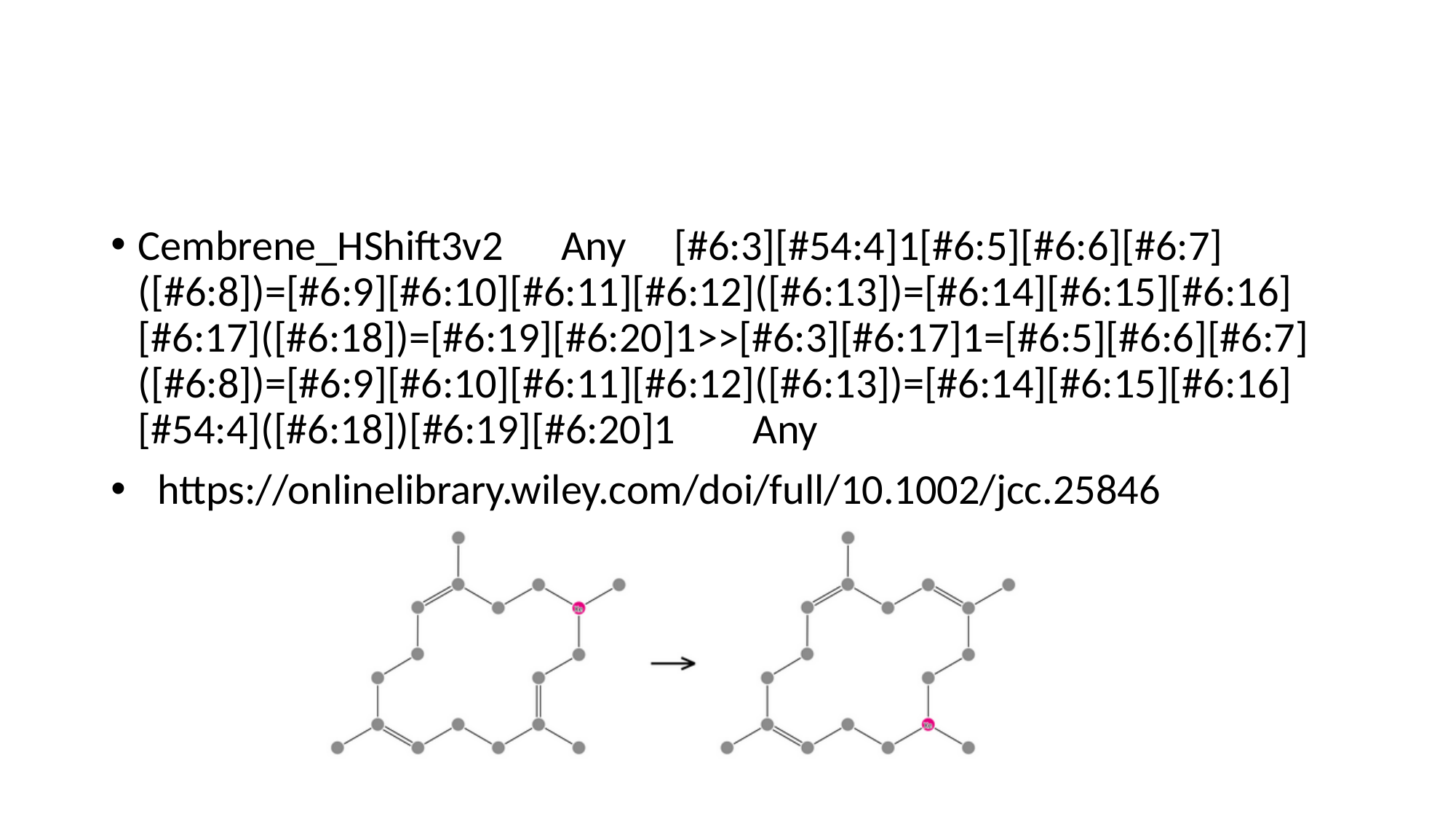

#
Cembrene_HShift3v2 Any [#6:3][#54:4]1[#6:5][#6:6][#6:7]([#6:8])=[#6:9][#6:10][#6:11][#6:12]([#6:13])=[#6:14][#6:15][#6:16][#6:17]([#6:18])=[#6:19][#6:20]1>>[#6:3][#6:17]1=[#6:5][#6:6][#6:7]([#6:8])=[#6:9][#6:10][#6:11][#6:12]([#6:13])=[#6:14][#6:15][#6:16][#54:4]([#6:18])[#6:19][#6:20]1 Any
 https://onlinelibrary.wiley.com/doi/full/10.1002/jcc.25846

### Slide 44
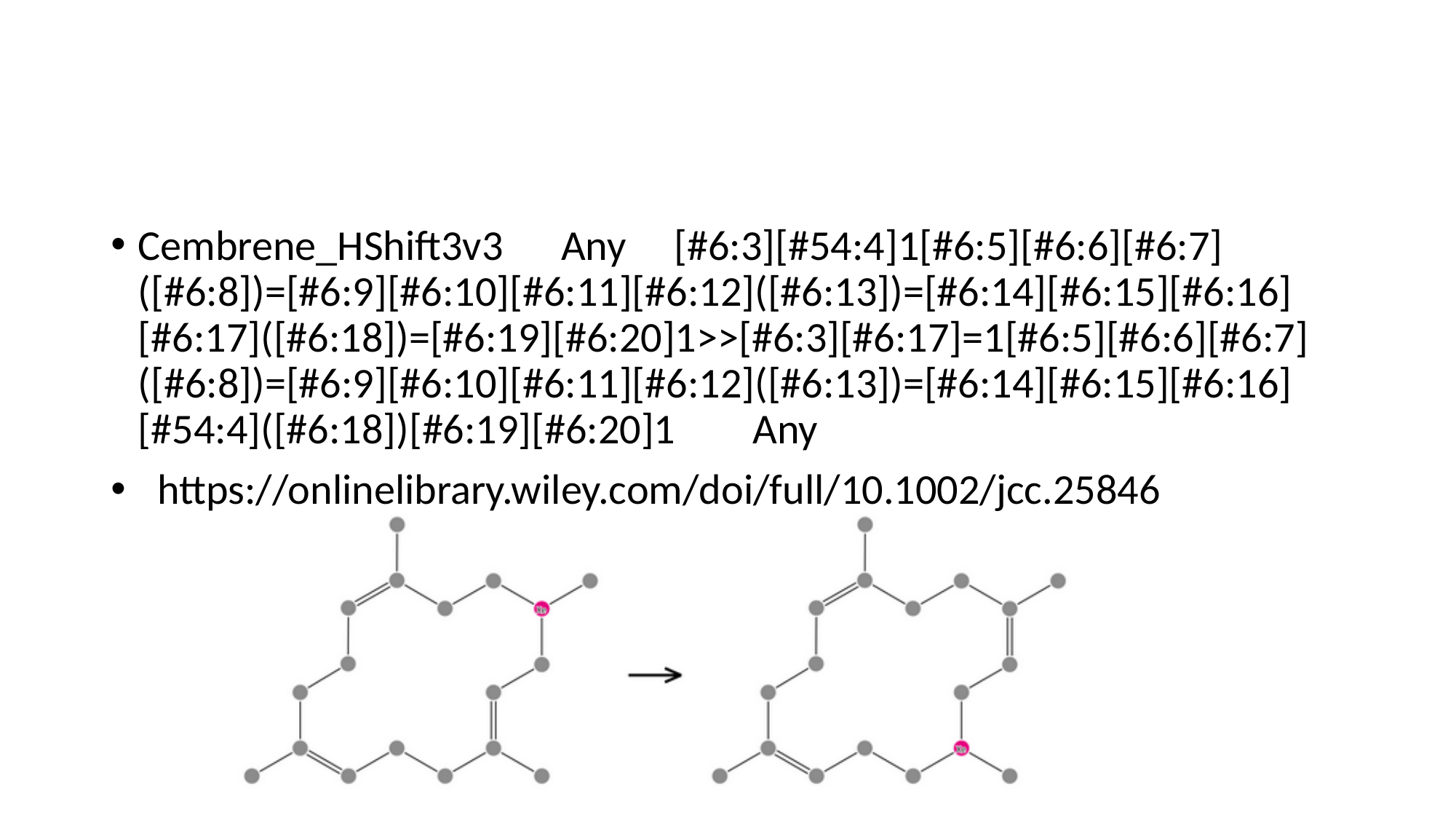

#
Cembrene_HShift3v3 Any [#6:3][#54:4]1[#6:5][#6:6][#6:7]([#6:8])=[#6:9][#6:10][#6:11][#6:12]([#6:13])=[#6:14][#6:15][#6:16][#6:17]([#6:18])=[#6:19][#6:20]1>>[#6:3][#6:17]=1[#6:5][#6:6][#6:7]([#6:8])=[#6:9][#6:10][#6:11][#6:12]([#6:13])=[#6:14][#6:15][#6:16][#54:4]([#6:18])[#6:19][#6:20]1 Any
 https://onlinelibrary.wiley.com/doi/full/10.1002/jcc.25846

### Slide 45
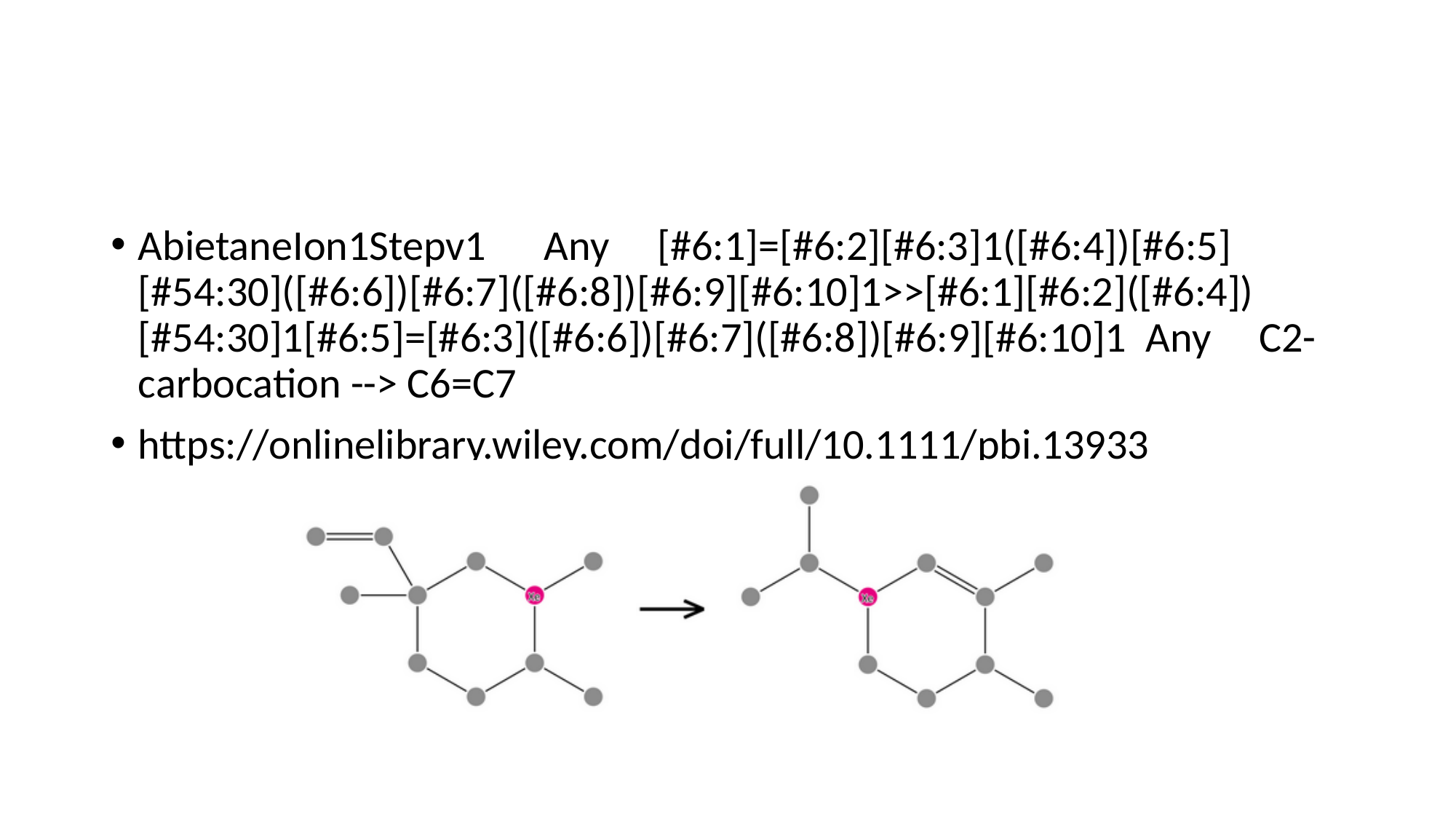

#
AbietaneIon1Stepv1 Any [#6:1]=[#6:2][#6:3]1([#6:4])[#6:5][#54:30]([#6:6])[#6:7]([#6:8])[#6:9][#6:10]1>>[#6:1][#6:2]([#6:4])[#54:30]1[#6:5]=[#6:3]([#6:6])[#6:7]([#6:8])[#6:9][#6:10]1 Any C2-carbocation --> C6=C7
https://onlinelibrary.wiley.com/doi/full/10.1111/pbi.13933

### Slide 46
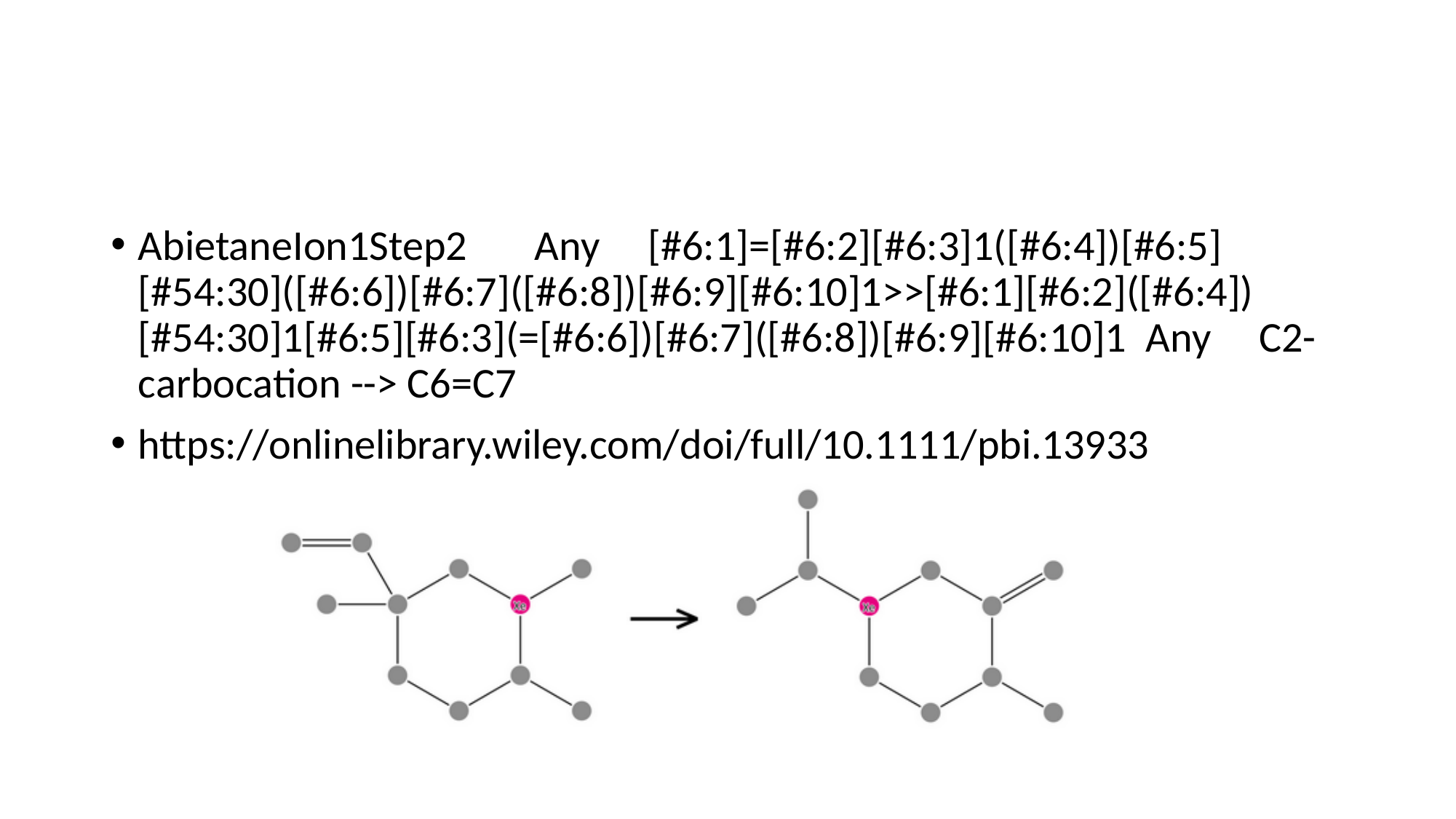

#
AbietaneIon1Step2 Any [#6:1]=[#6:2][#6:3]1([#6:4])[#6:5][#54:30]([#6:6])[#6:7]([#6:8])[#6:9][#6:10]1>>[#6:1][#6:2]([#6:4])[#54:30]1[#6:5][#6:3](=[#6:6])[#6:7]([#6:8])[#6:9][#6:10]1 Any C2-carbocation --> C6=C7
https://onlinelibrary.wiley.com/doi/full/10.1111/pbi.13933

### Slide 47
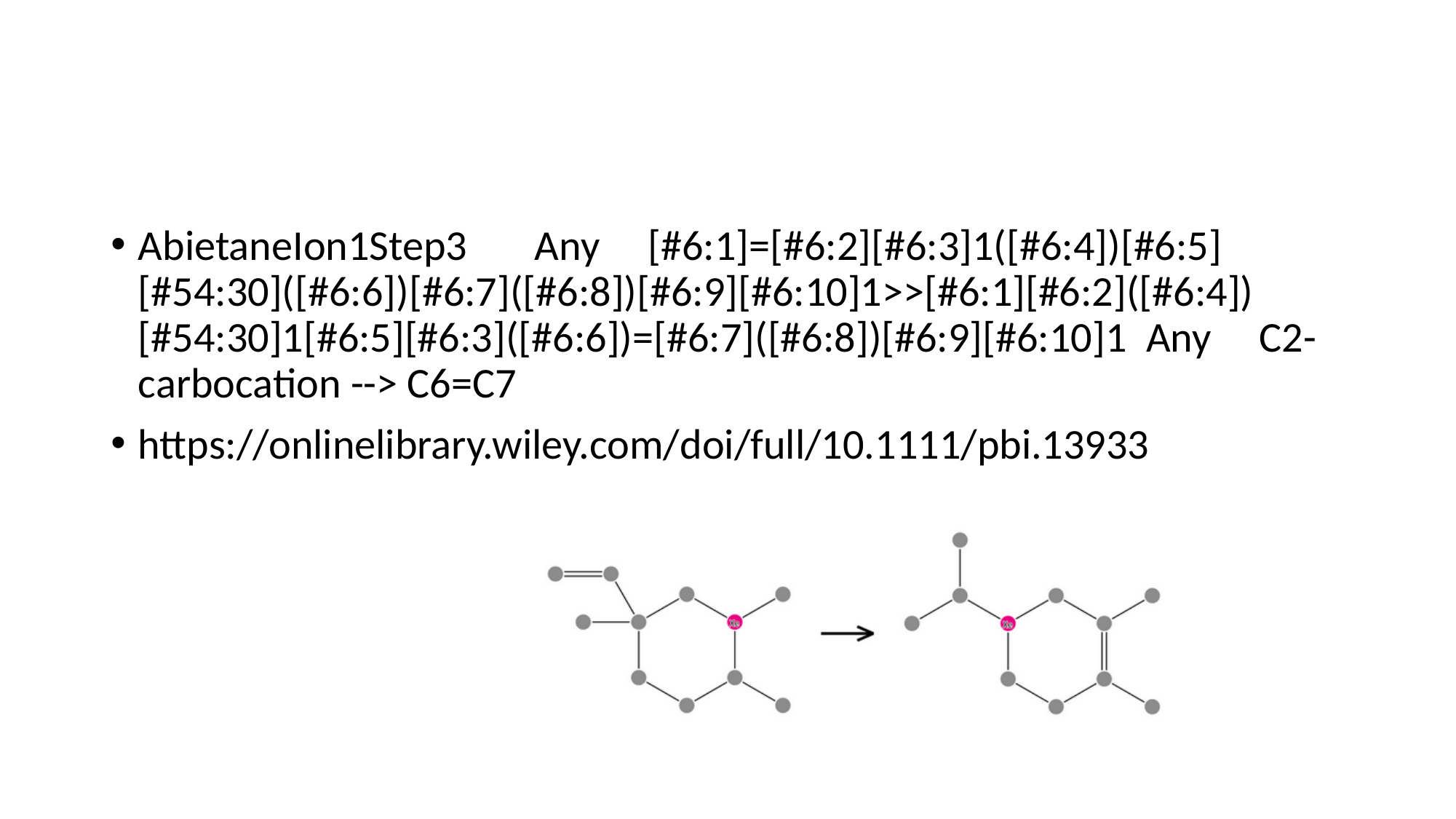

#
AbietaneIon1Step3 Any [#6:1]=[#6:2][#6:3]1([#6:4])[#6:5][#54:30]([#6:6])[#6:7]([#6:8])[#6:9][#6:10]1>>[#6:1][#6:2]([#6:4])[#54:30]1[#6:5][#6:3]([#6:6])=[#6:7]([#6:8])[#6:9][#6:10]1 Any C2-carbocation --> C6=C7
https://onlinelibrary.wiley.com/doi/full/10.1111/pbi.13933

### Slide 48
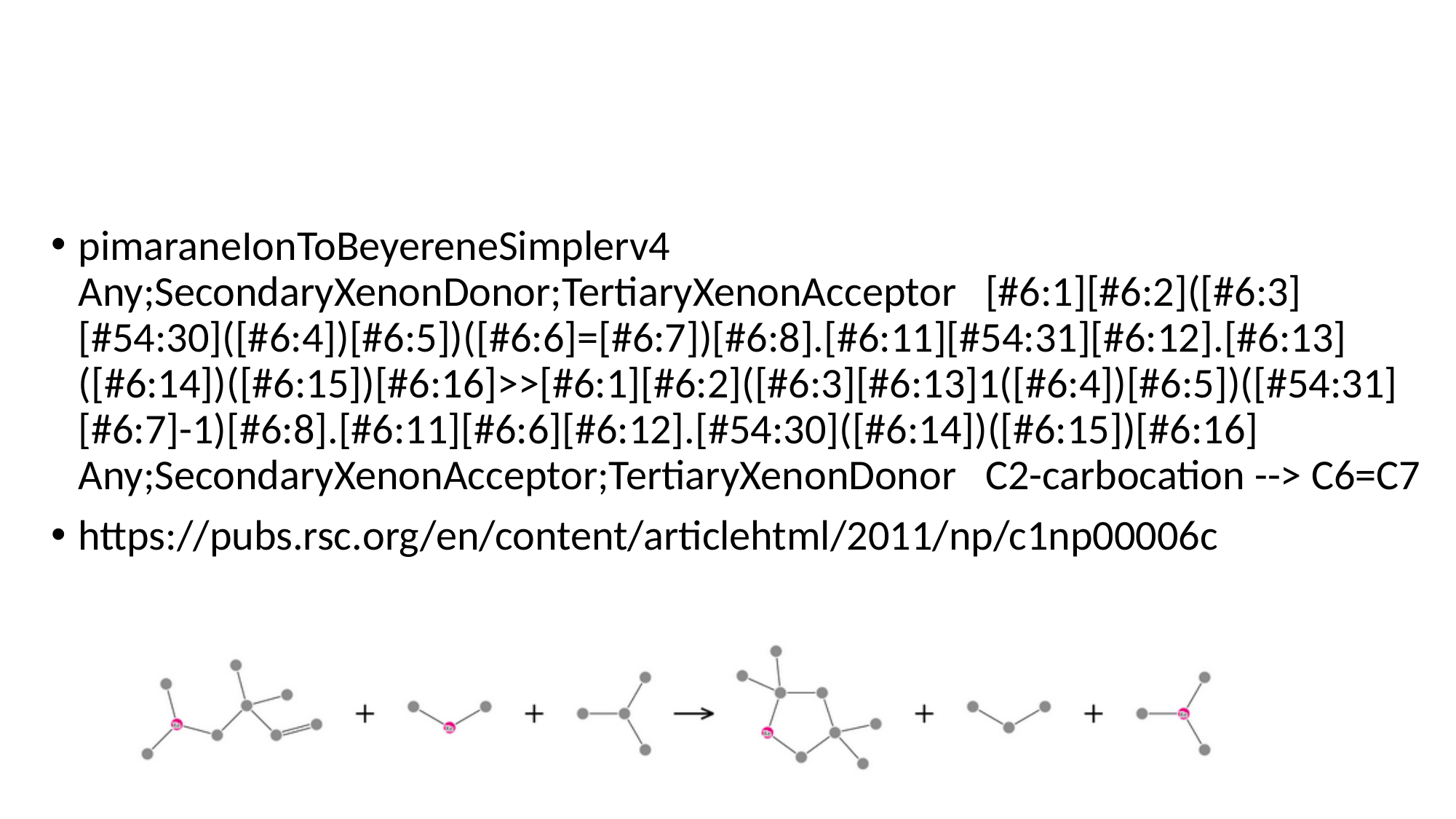

#
pimaraneIonToBeyereneSimplerv4 Any;SecondaryXenonDonor;TertiaryXenonAcceptor [#6:1][#6:2]([#6:3][#54:30]([#6:4])[#6:5])([#6:6]=[#6:7])[#6:8].[#6:11][#54:31][#6:12].[#6:13]([#6:14])([#6:15])[#6:16]>>[#6:1][#6:2]([#6:3][#6:13]1([#6:4])[#6:5])([#54:31][#6:7]-1)[#6:8].[#6:11][#6:6][#6:12].[#54:30]([#6:14])([#6:15])[#6:16] Any;SecondaryXenonAcceptor;TertiaryXenonDonor C2-carbocation --> C6=C7
https://pubs.rsc.org/en/content/articlehtml/2011/np/c1np00006c

### Slide 49
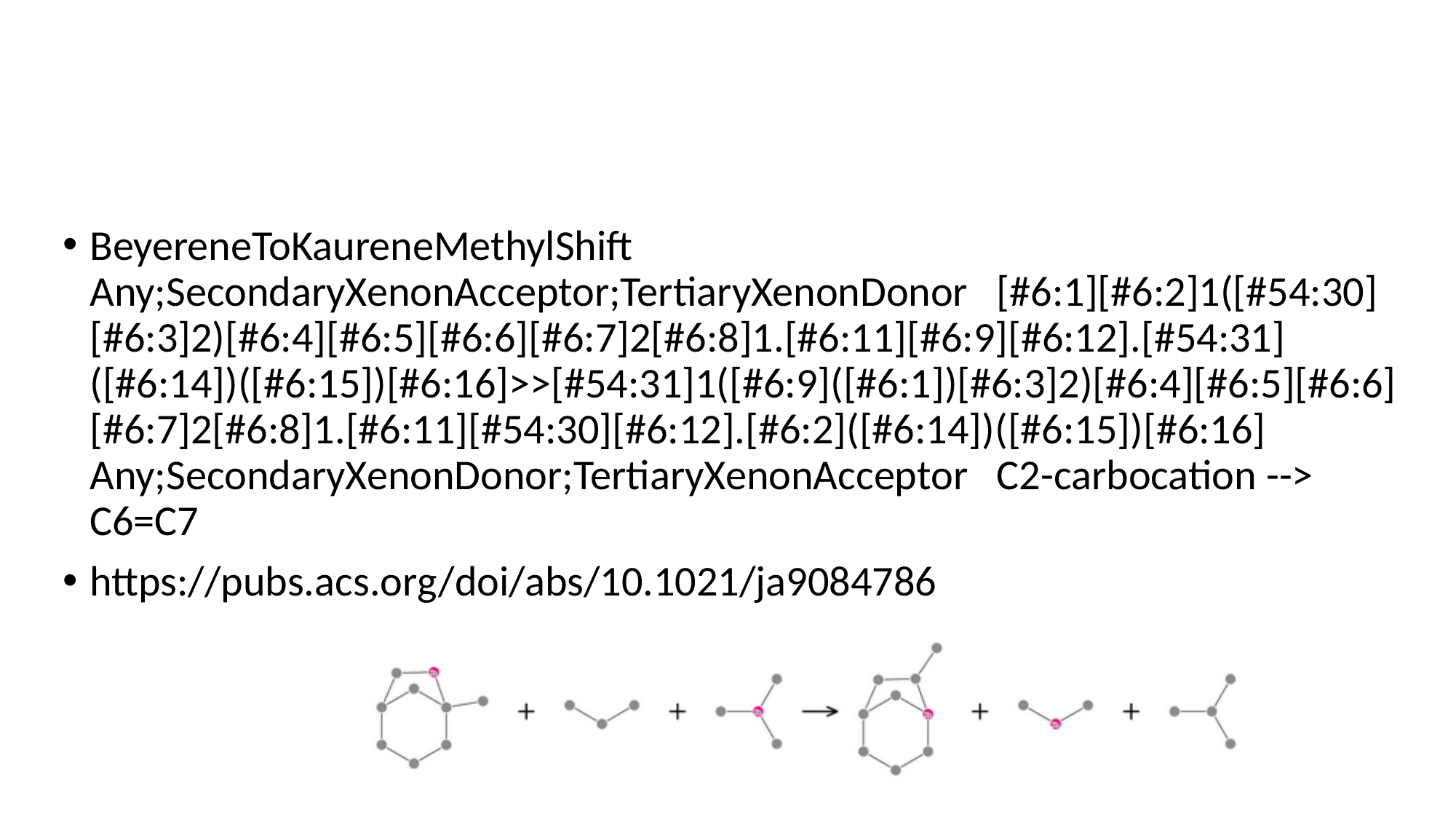

#
BeyereneToKaureneMethylShift Any;SecondaryXenonAcceptor;TertiaryXenonDonor [#6:1][#6:2]1([#54:30][#6:3]2)[#6:4][#6:5][#6:6][#6:7]2[#6:8]1.[#6:11][#6:9][#6:12].[#54:31]([#6:14])([#6:15])[#6:16]>>[#54:31]1([#6:9]([#6:1])[#6:3]2)[#6:4][#6:5][#6:6][#6:7]2[#6:8]1.[#6:11][#54:30][#6:12].[#6:2]([#6:14])([#6:15])[#6:16] Any;SecondaryXenonDonor;TertiaryXenonAcceptor C2-carbocation --> C6=C7
https://pubs.acs.org/doi/abs/10.1021/ja9084786

### Slide 50
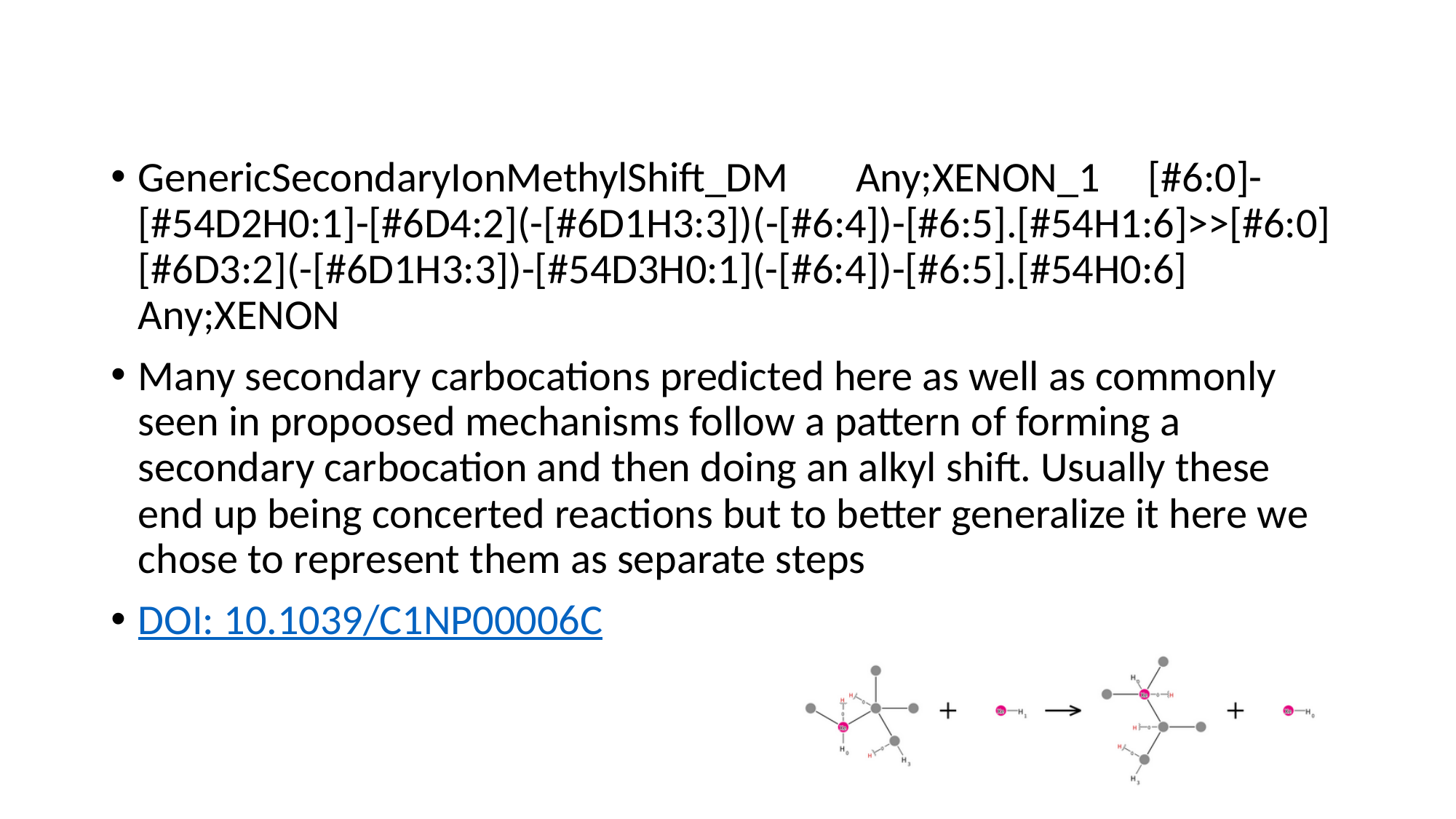

#
GenericSecondaryIonMethylShift_DM Any;XENON_1 [#6:0]-[#54D2H0:1]-[#6D4:2](-[#6D1H3:3])(-[#6:4])-[#6:5].[#54H1:6]>>[#6:0][#6D3:2](-[#6D1H3:3])-[#54D3H0:1](-[#6:4])-[#6:5].[#54H0:6] Any;XENON
Many secondary carbocations predicted here as well as commonly seen in propoosed mechanisms follow a pattern of forming a secondary carbocation and then doing an alkyl shift. Usually these end up being concerted reactions but to better generalize it here we chose to represent them as separate steps
DOI: 10.1039/C1NP00006C

### Slide 51
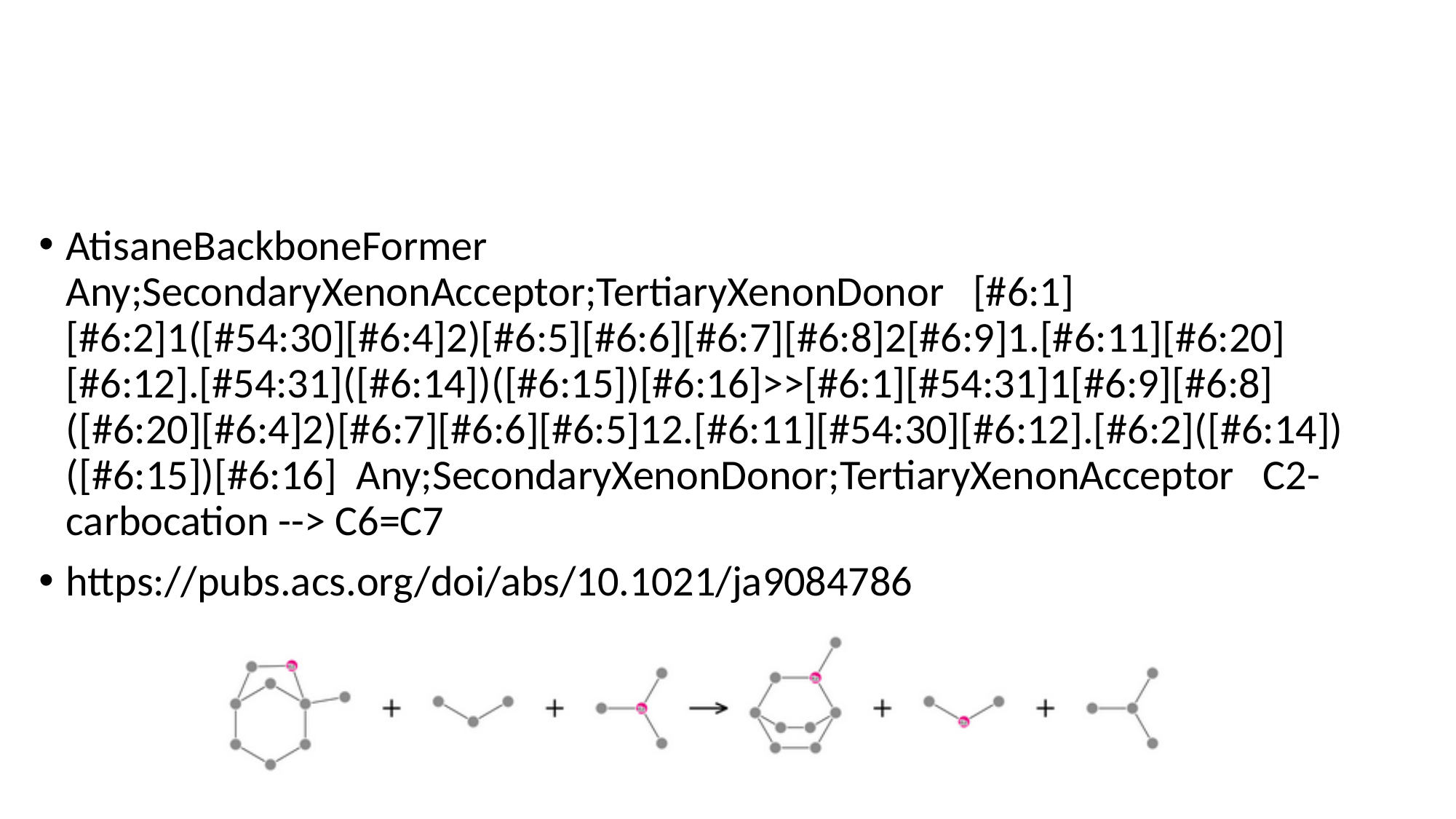

#
AtisaneBackboneFormer Any;SecondaryXenonAcceptor;TertiaryXenonDonor [#6:1][#6:2]1([#54:30][#6:4]2)[#6:5][#6:6][#6:7][#6:8]2[#6:9]1.[#6:11][#6:20][#6:12].[#54:31]([#6:14])([#6:15])[#6:16]>>[#6:1][#54:31]1[#6:9][#6:8]([#6:20][#6:4]2)[#6:7][#6:6][#6:5]12.[#6:11][#54:30][#6:12].[#6:2]([#6:14])([#6:15])[#6:16] Any;SecondaryXenonDonor;TertiaryXenonAcceptor C2-carbocation --> C6=C7
https://pubs.acs.org/doi/abs/10.1021/ja9084786

### Slide 52
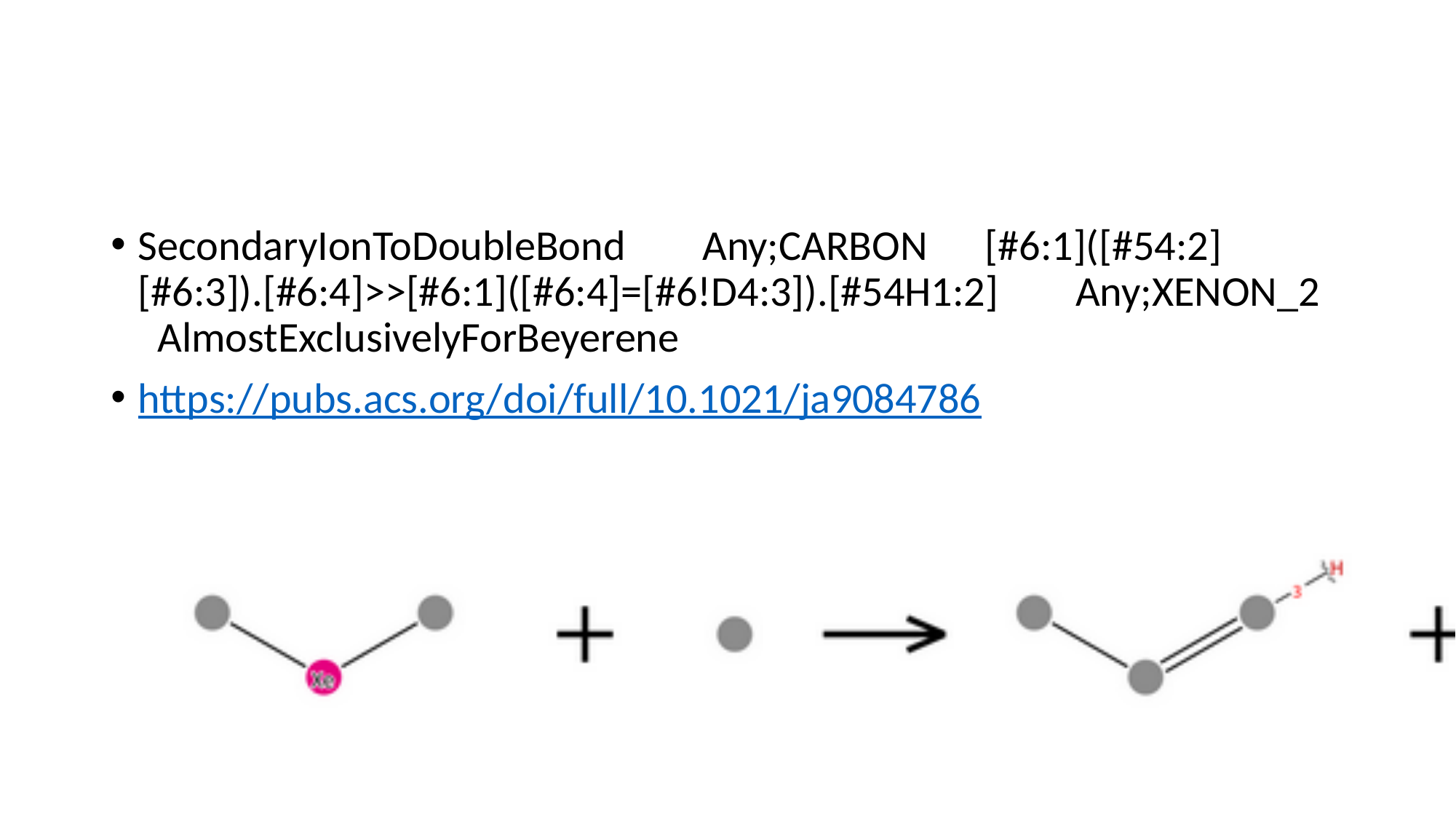

#
SecondaryIonToDoubleBond Any;CARBON [#6:1]([#54:2][#6:3]).[#6:4]>>[#6:1]([#6:4]=[#6!D4:3]).[#54H1:2] Any;XENON_2 AlmostExclusivelyForBeyerene
https://pubs.acs.org/doi/full/10.1021/ja9084786

### Slide 53
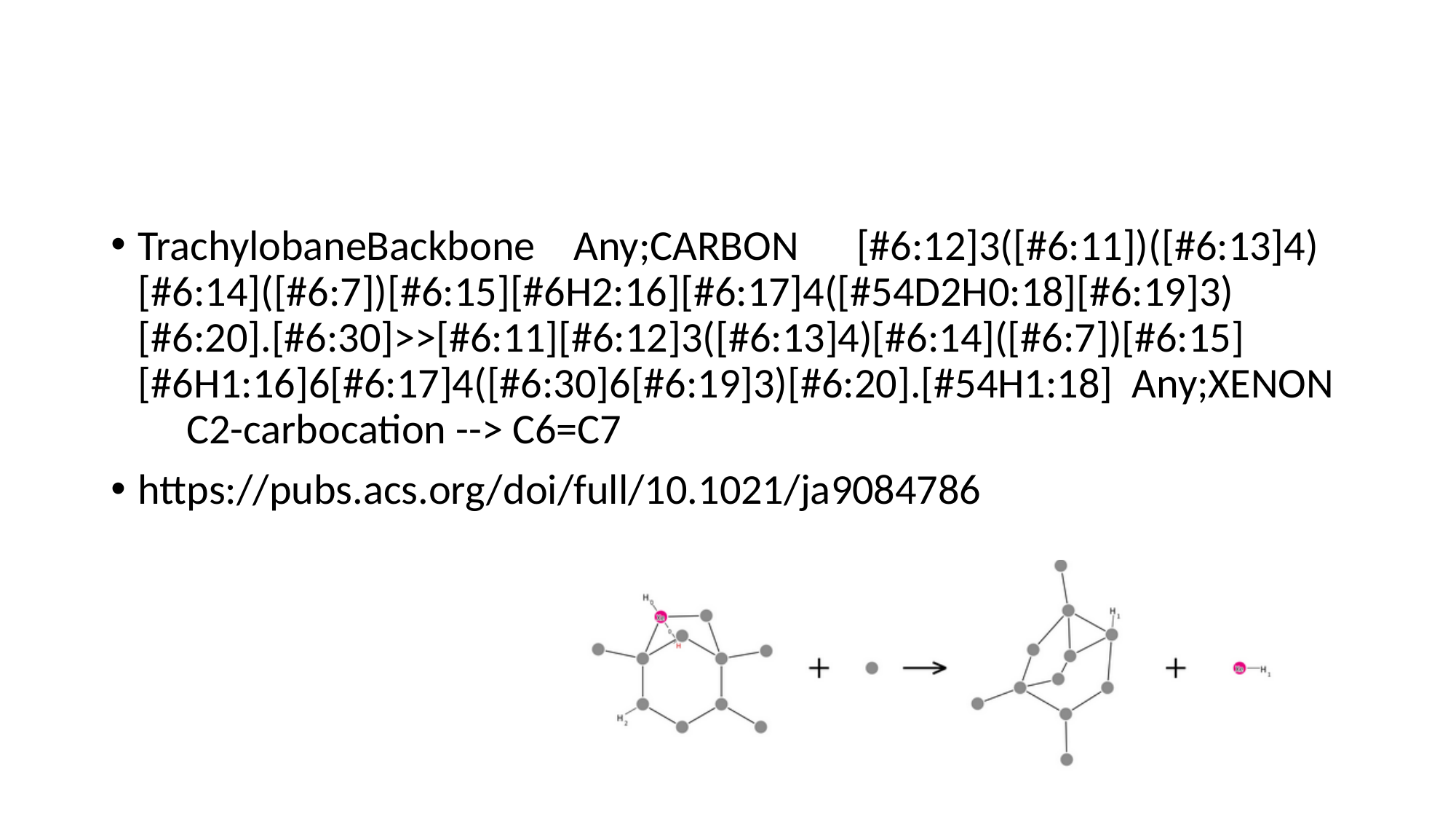

#
TrachylobaneBackbone Any;CARBON [#6:12]3([#6:11])([#6:13]4)[#6:14]([#6:7])[#6:15][#6H2:16][#6:17]4([#54D2H0:18][#6:19]3)[#6:20].[#6:30]>>[#6:11][#6:12]3([#6:13]4)[#6:14]([#6:7])[#6:15][#6H1:16]6[#6:17]4([#6:30]6[#6:19]3)[#6:20].[#54H1:18] Any;XENON C2-carbocation --> C6=C7
https://pubs.acs.org/doi/full/10.1021/ja9084786

### Slide 54
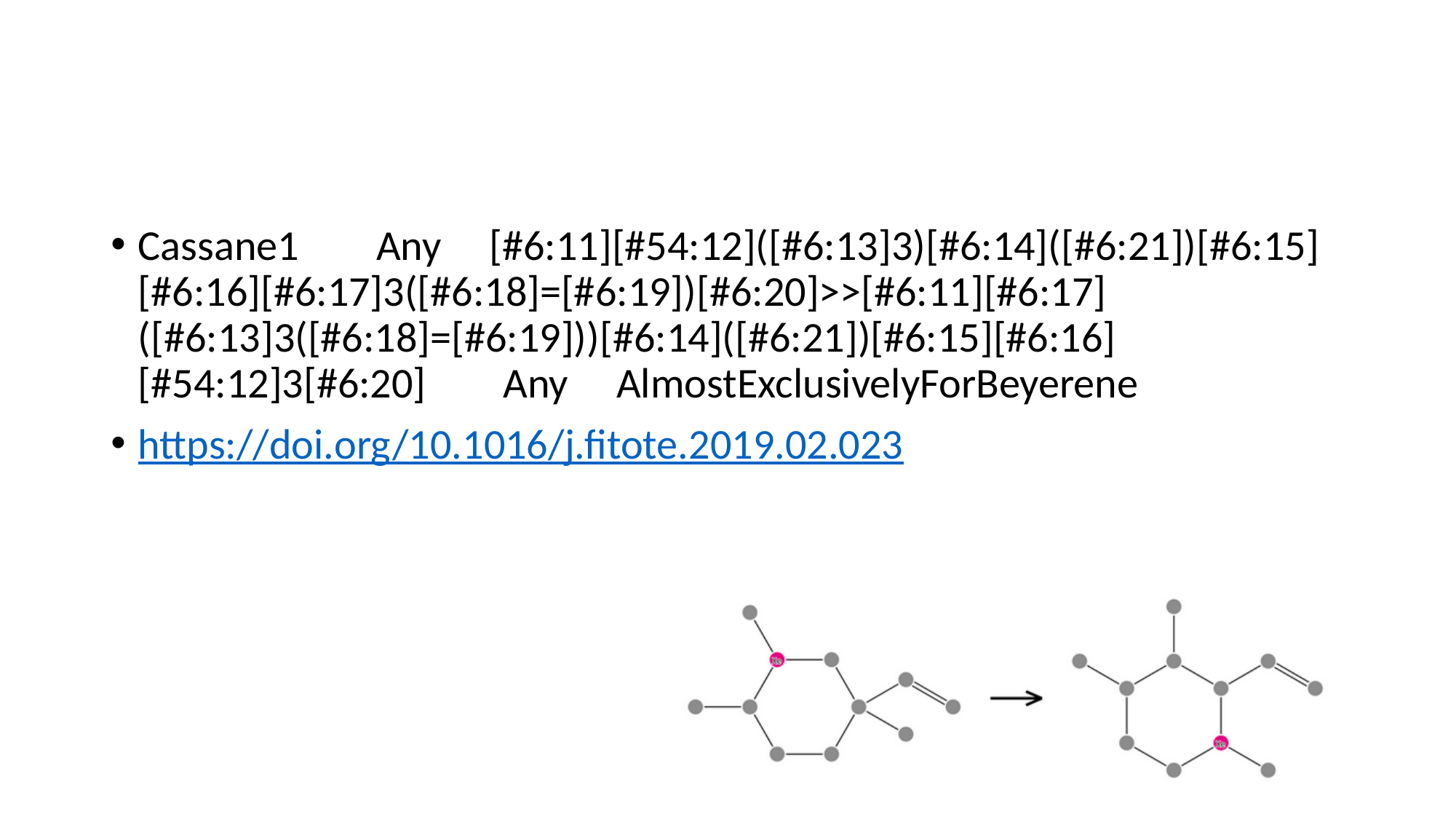

#
Cassane1 Any [#6:11][#54:12]([#6:13]3)[#6:14]([#6:21])[#6:15][#6:16][#6:17]3([#6:18]=[#6:19])[#6:20]>>[#6:11][#6:17]([#6:13]3([#6:18]=[#6:19]))[#6:14]([#6:21])[#6:15][#6:16][#54:12]3[#6:20] Any AlmostExclusivelyForBeyerene
https://doi.org/10.1016/j.fitote.2019.02.023

### Slide 55
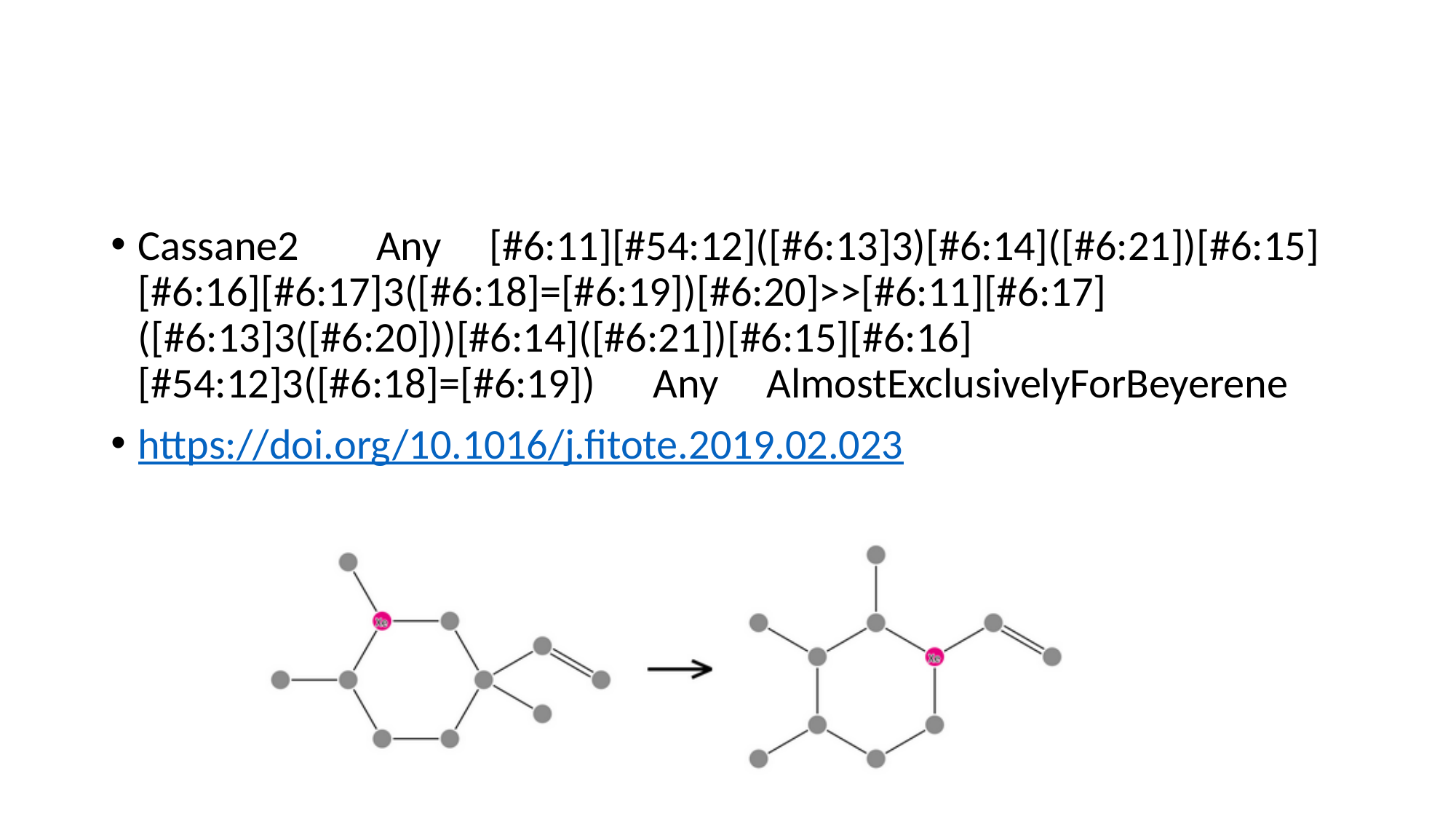

#
Cassane2 Any [#6:11][#54:12]([#6:13]3)[#6:14]([#6:21])[#6:15][#6:16][#6:17]3([#6:18]=[#6:19])[#6:20]>>[#6:11][#6:17]([#6:13]3([#6:20]))[#6:14]([#6:21])[#6:15][#6:16][#54:12]3([#6:18]=[#6:19]) Any AlmostExclusivelyForBeyerene
https://doi.org/10.1016/j.fitote.2019.02.023

### Slide 56
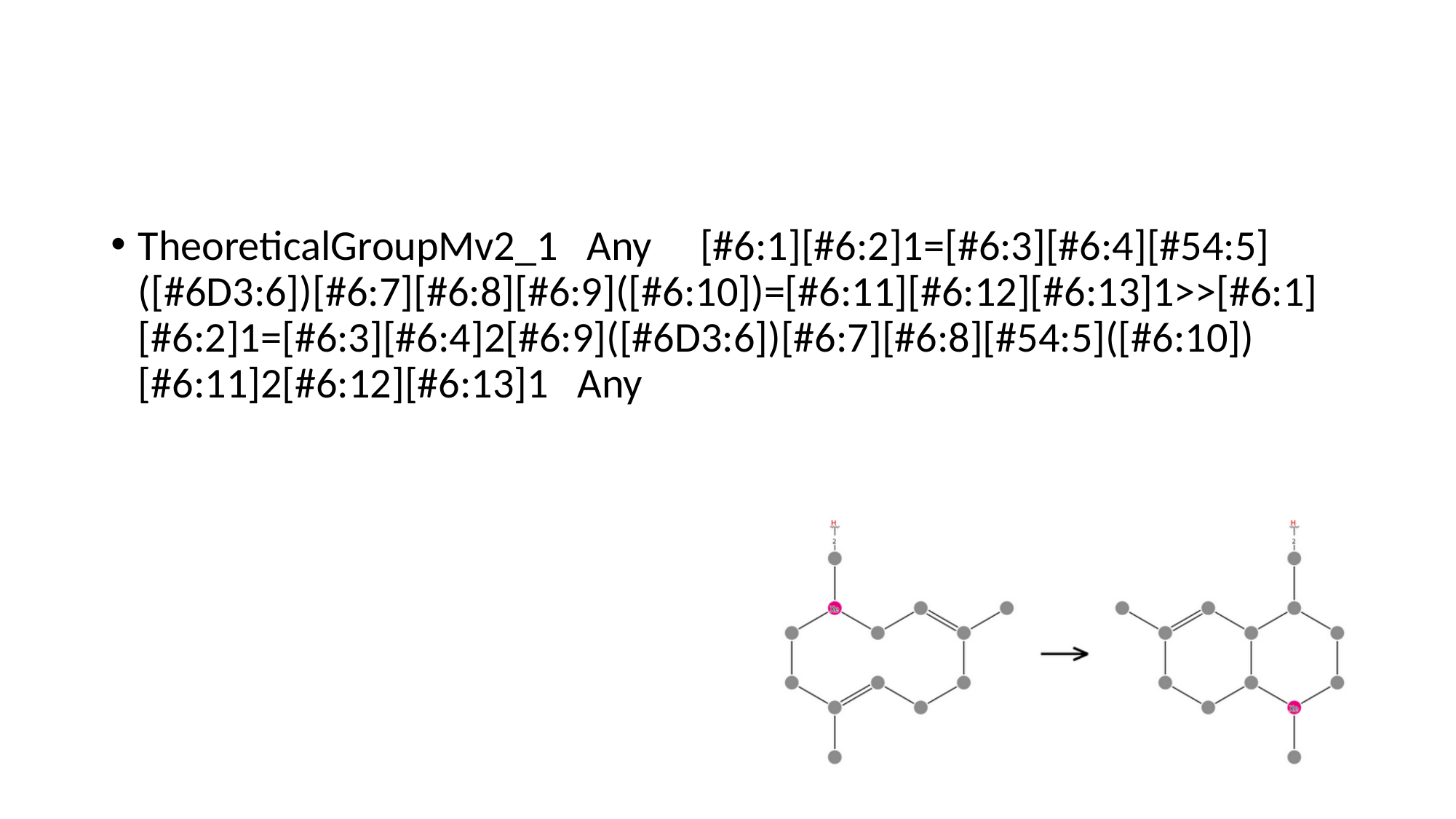

#
TheoreticalGroupMv2_1 Any [#6:1][#6:2]1=[#6:3][#6:4][#54:5]([#6D3:6])[#6:7][#6:8][#6:9]([#6:10])=[#6:11][#6:12][#6:13]1>>[#6:1][#6:2]1=[#6:3][#6:4]2[#6:9]([#6D3:6])[#6:7][#6:8][#54:5]([#6:10])[#6:11]2[#6:12][#6:13]1 Any

### Slide 57
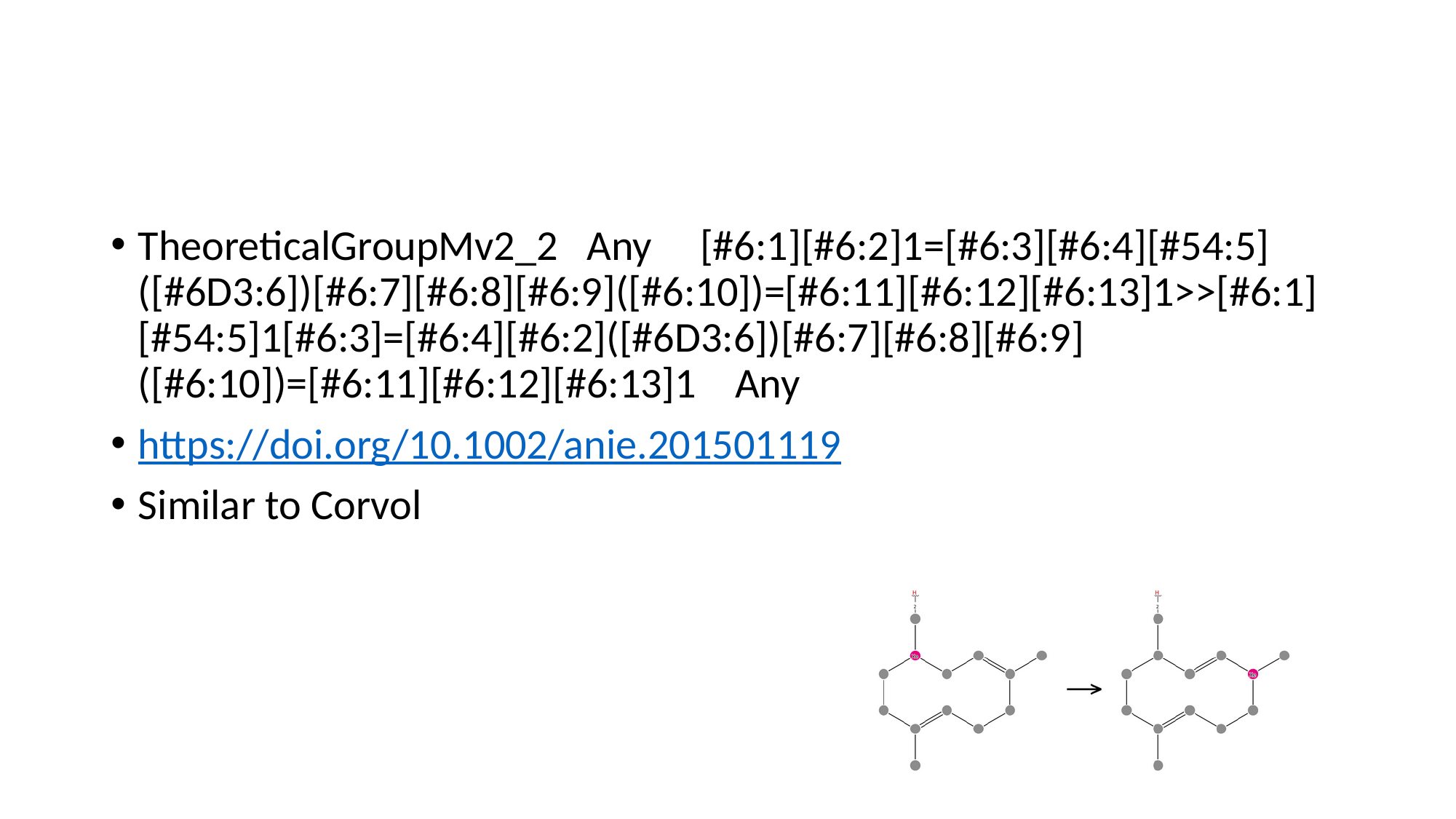

#
TheoreticalGroupMv2_2 Any [#6:1][#6:2]1=[#6:3][#6:4][#54:5]([#6D3:6])[#6:7][#6:8][#6:9]([#6:10])=[#6:11][#6:12][#6:13]1>>[#6:1][#54:5]1[#6:3]=[#6:4][#6:2]([#6D3:6])[#6:7][#6:8][#6:9]([#6:10])=[#6:11][#6:12][#6:13]1 Any
https://doi.org/10.1002/anie.201501119
Similar to Corvol

### Slide 58
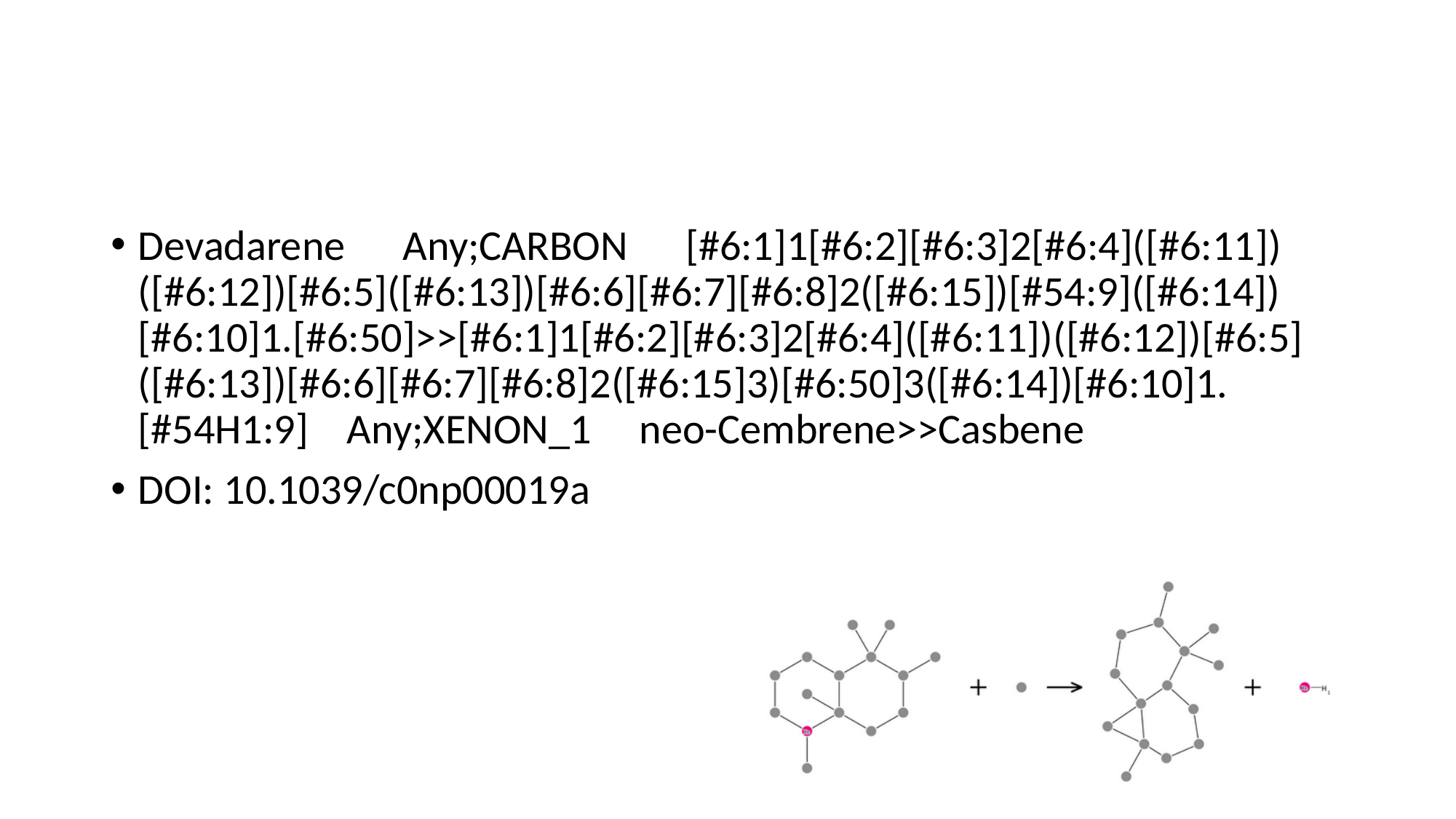

#
Devadarene Any;CARBON [#6:1]1[#6:2][#6:3]2[#6:4]([#6:11])([#6:12])[#6:5]([#6:13])[#6:6][#6:7][#6:8]2([#6:15])[#54:9]([#6:14])[#6:10]1.[#6:50]>>[#6:1]1[#6:2][#6:3]2[#6:4]([#6:11])([#6:12])[#6:5]([#6:13])[#6:6][#6:7][#6:8]2([#6:15]3)[#6:50]3([#6:14])[#6:10]1.[#54H1:9] Any;XENON_1 neo-Cembrene>>Casbene
DOI: 10.1039/c0np00019a

### Slide 59
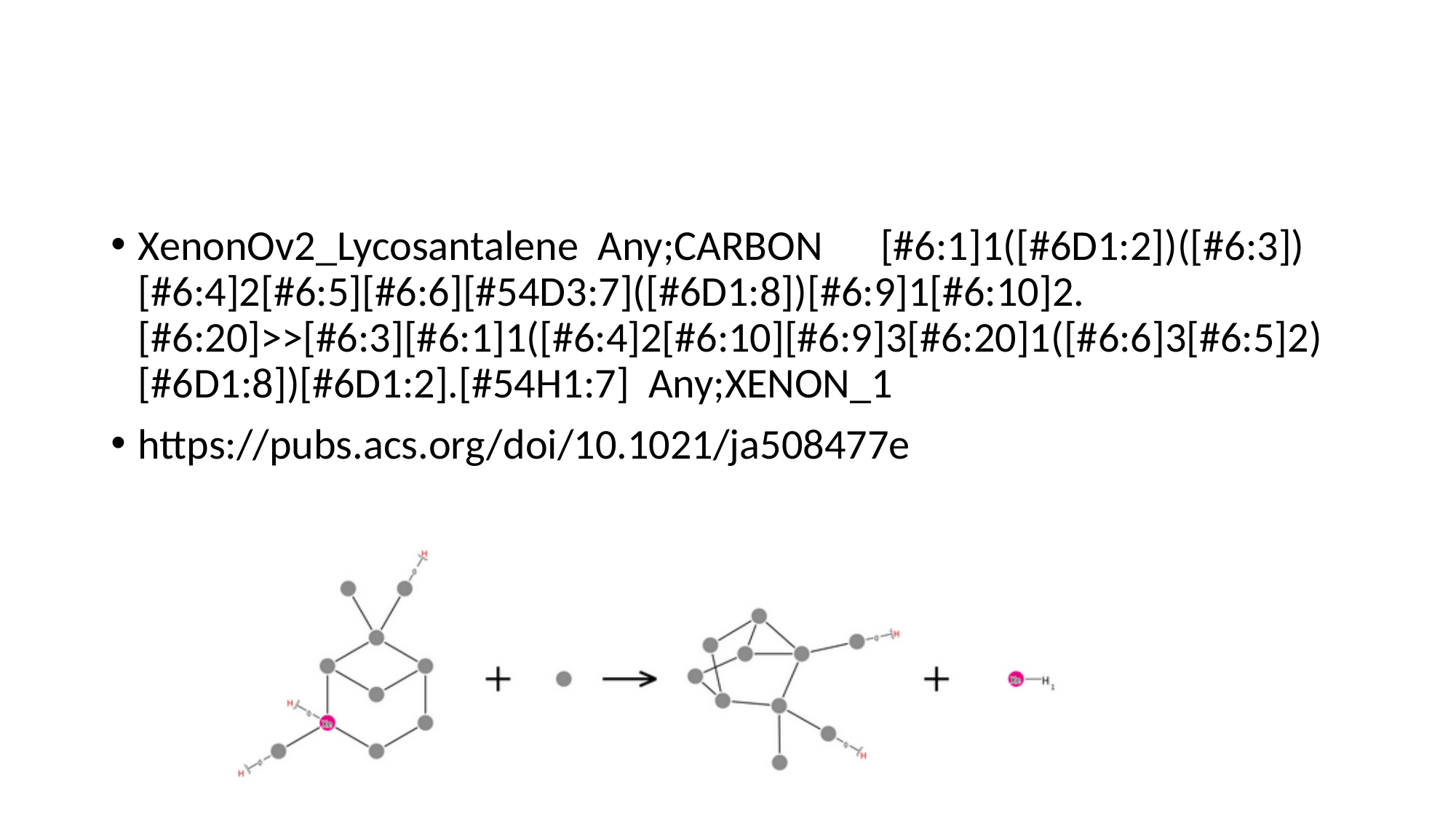

#
XenonOv2_Lycosantalene Any;CARBON [#6:1]1([#6D1:2])([#6:3])[#6:4]2[#6:5][#6:6][#54D3:7]([#6D1:8])[#6:9]1[#6:10]2.[#6:20]>>[#6:3][#6:1]1([#6:4]2[#6:10][#6:9]3[#6:20]1([#6:6]3[#6:5]2)[#6D1:8])[#6D1:2].[#54H1:7] Any;XENON_1
https://pubs.acs.org/doi/10.1021/ja508477e

### Slide 60
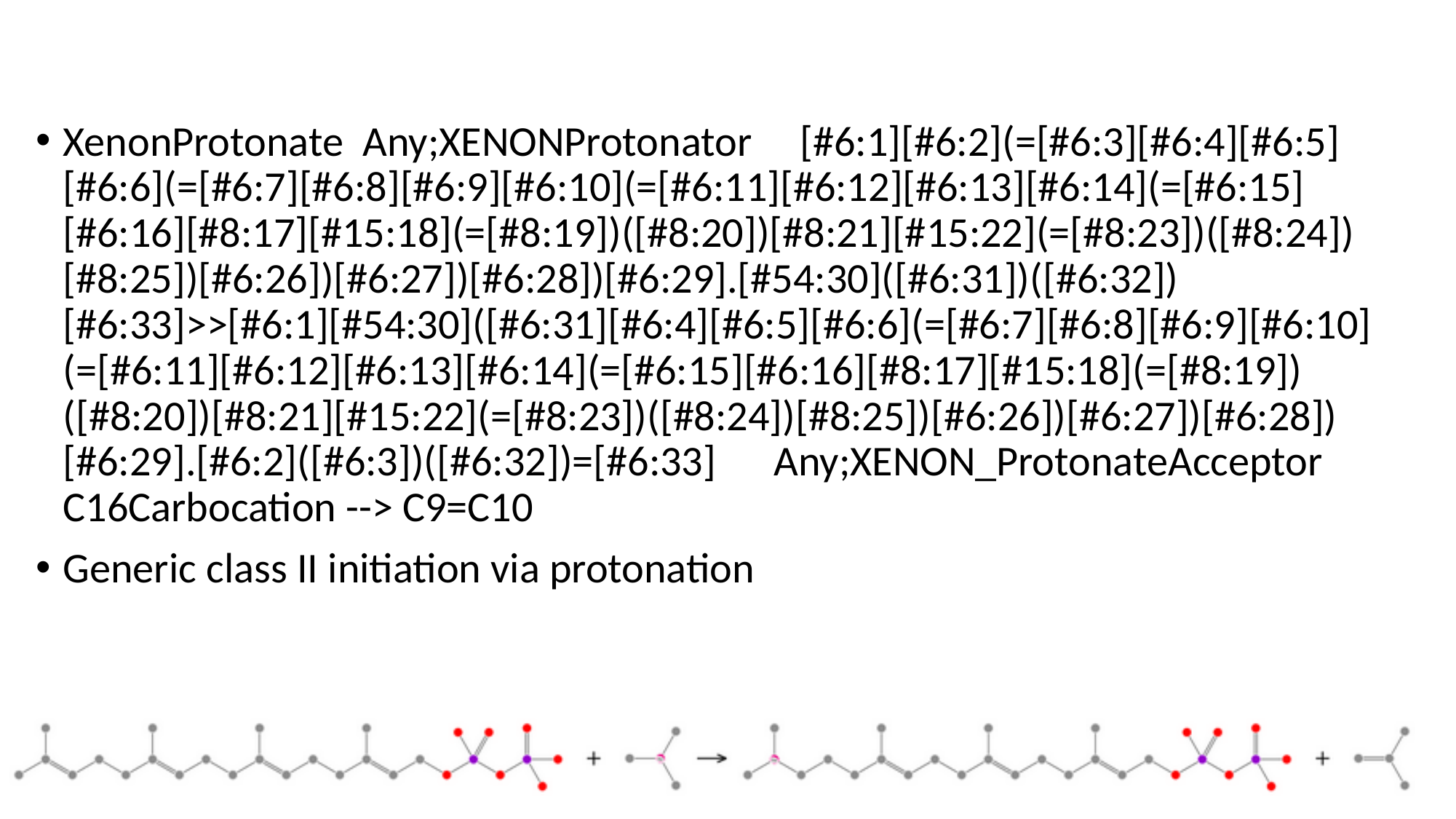

XenonProtonate Any;XENONProtonator [#6:1][#6:2](=[#6:3][#6:4][#6:5][#6:6](=[#6:7][#6:8][#6:9][#6:10](=[#6:11][#6:12][#6:13][#6:14](=[#6:15][#6:16][#8:17][#15:18](=[#8:19])([#8:20])[#8:21][#15:22](=[#8:23])([#8:24])[#8:25])[#6:26])[#6:27])[#6:28])[#6:29].[#54:30]([#6:31])([#6:32])[#6:33]>>[#6:1][#54:30]([#6:31][#6:4][#6:5][#6:6](=[#6:7][#6:8][#6:9][#6:10](=[#6:11][#6:12][#6:13][#6:14](=[#6:15][#6:16][#8:17][#15:18](=[#8:19])([#8:20])[#8:21][#15:22](=[#8:23])([#8:24])[#8:25])[#6:26])[#6:27])[#6:28])[#6:29].[#6:2]([#6:3])([#6:32])=[#6:33] Any;XENON_ProtonateAcceptor C16Carbocation --> C9=C10
Generic class II initiation via protonation

### Slide 61
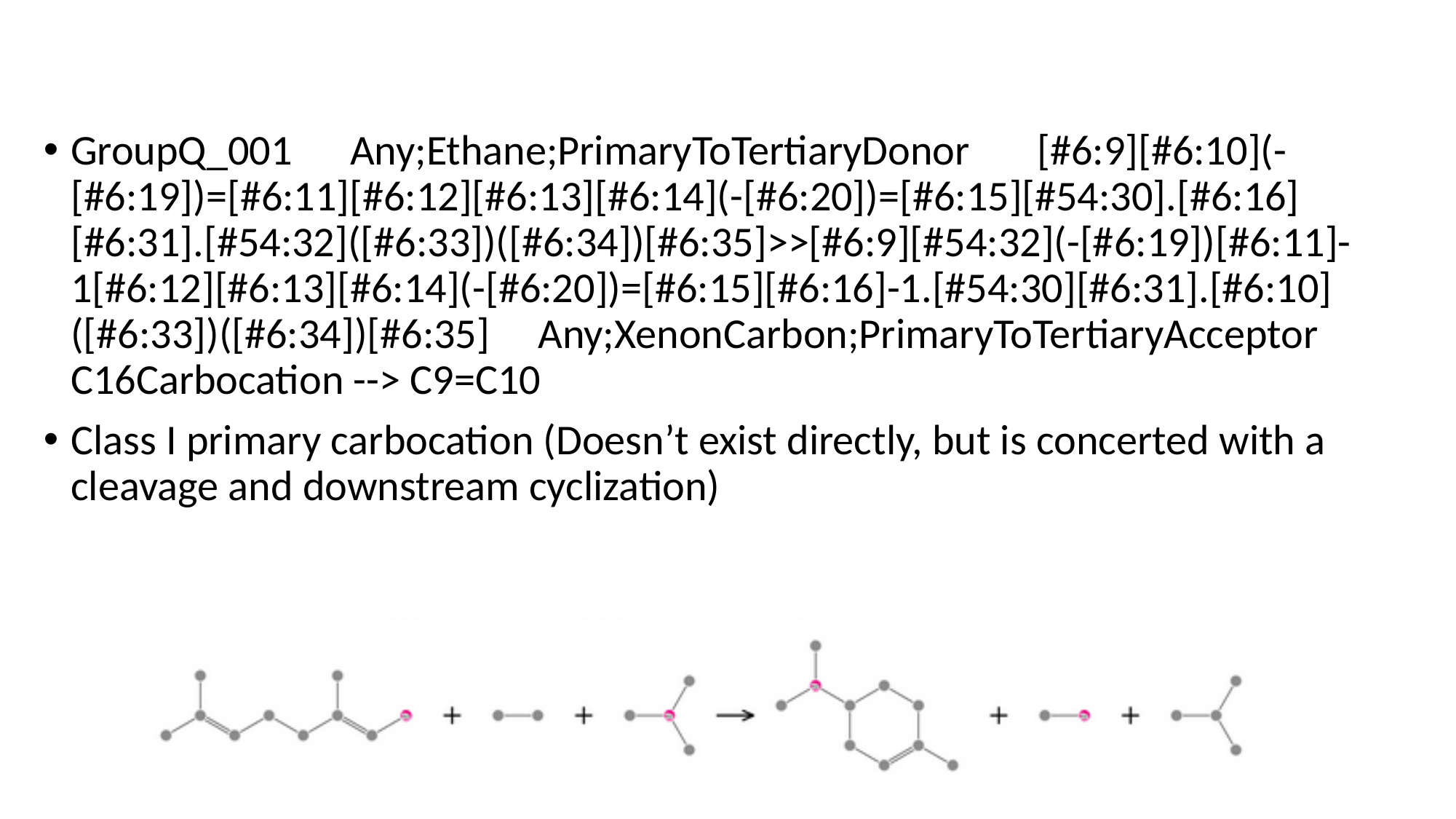

GroupQ_001 Any;Ethane;PrimaryToTertiaryDonor [#6:9][#6:10](-[#6:19])=[#6:11][#6:12][#6:13][#6:14](-[#6:20])=[#6:15][#54:30].[#6:16][#6:31].[#54:32]([#6:33])([#6:34])[#6:35]>>[#6:9][#54:32](-[#6:19])[#6:11]-1[#6:12][#6:13][#6:14](-[#6:20])=[#6:15][#6:16]-1.[#54:30][#6:31].[#6:10]([#6:33])([#6:34])[#6:35] Any;XenonCarbon;PrimaryToTertiaryAcceptor C16Carbocation --> C9=C10
Class I primary carbocation (Doesn’t exist directly, but is concerted with a cleavage and downstream cyclization)

### Slide 62
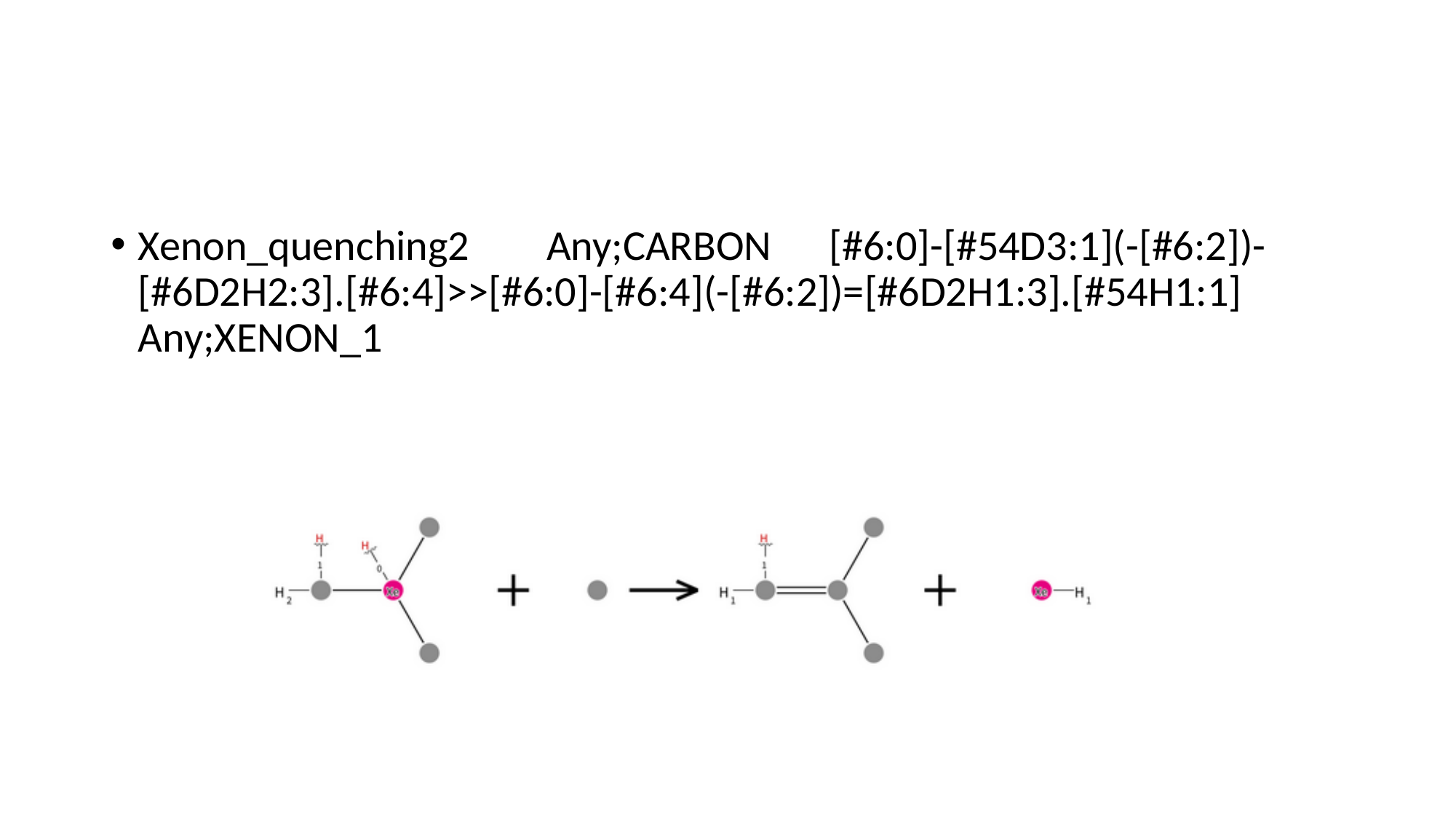

#
Xenon_quenching2 Any;CARBON [#6:0]-[#54D3:1](-[#6:2])-[#6D2H2:3].[#6:4]>>[#6:0]-[#6:4](-[#6:2])=[#6D2H1:3].[#54H1:1] Any;XENON_1

### Slide 63
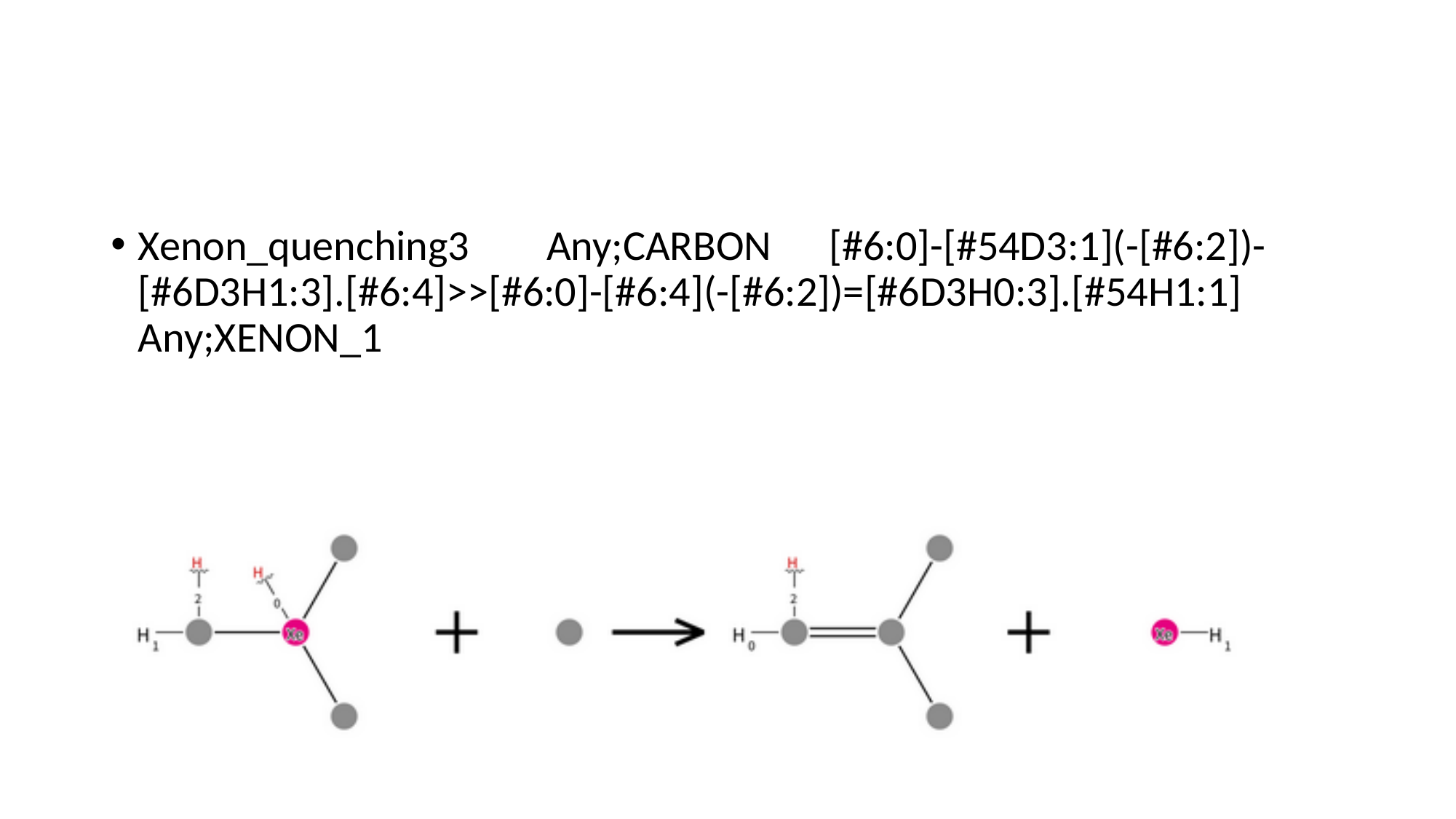

#
Xenon_quenching3 Any;CARBON [#6:0]-[#54D3:1](-[#6:2])-[#6D3H1:3].[#6:4]>>[#6:0]-[#6:4](-[#6:2])=[#6D3H0:3].[#54H1:1] Any;XENON_1

### Slide 64
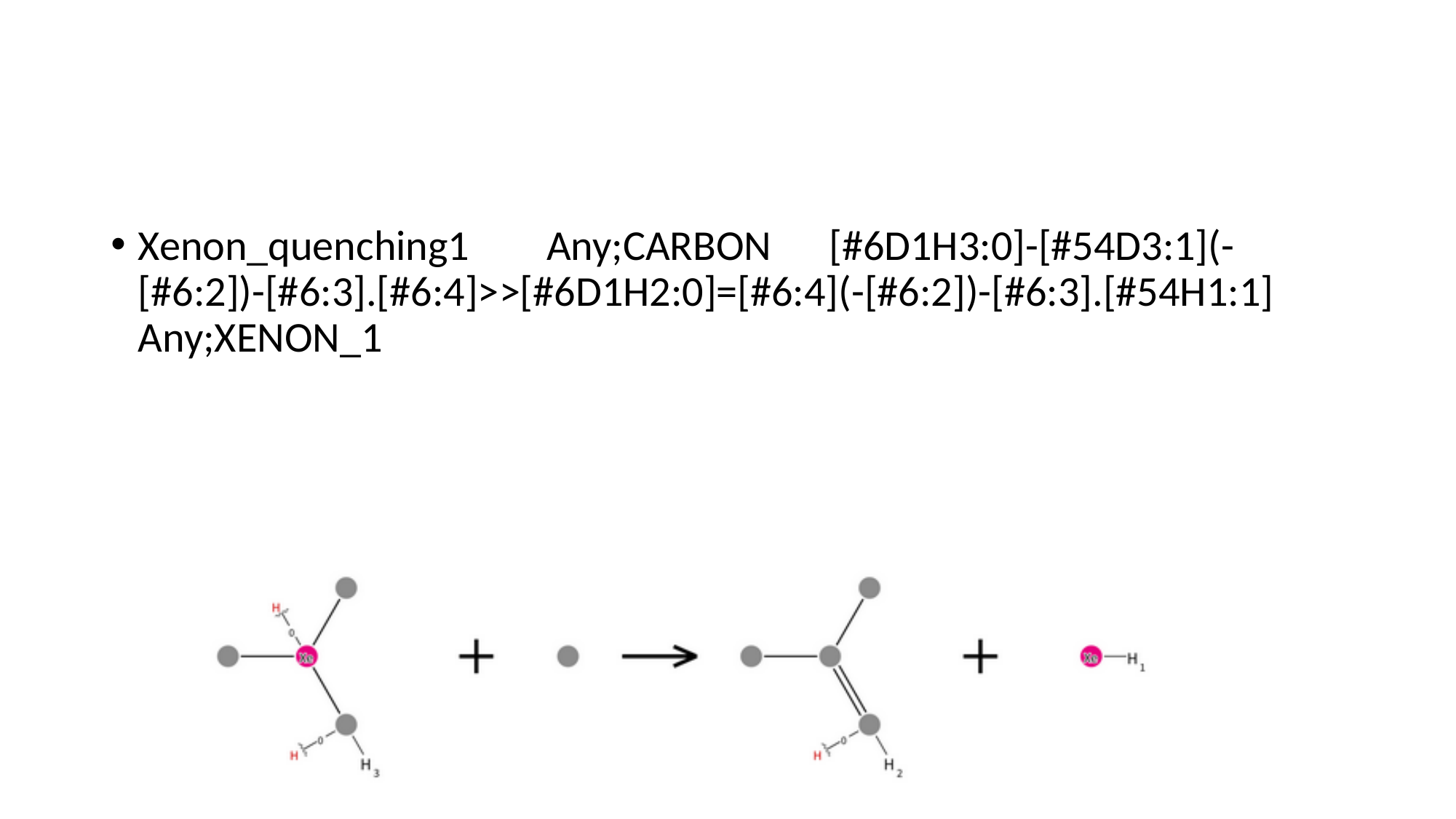

#
Xenon_quenching1 Any;CARBON [#6D1H3:0]-[#54D3:1](-[#6:2])-[#6:3].[#6:4]>>[#6D1H2:0]=[#6:4](-[#6:2])-[#6:3].[#54H1:1] Any;XENON_1

### Slide 65
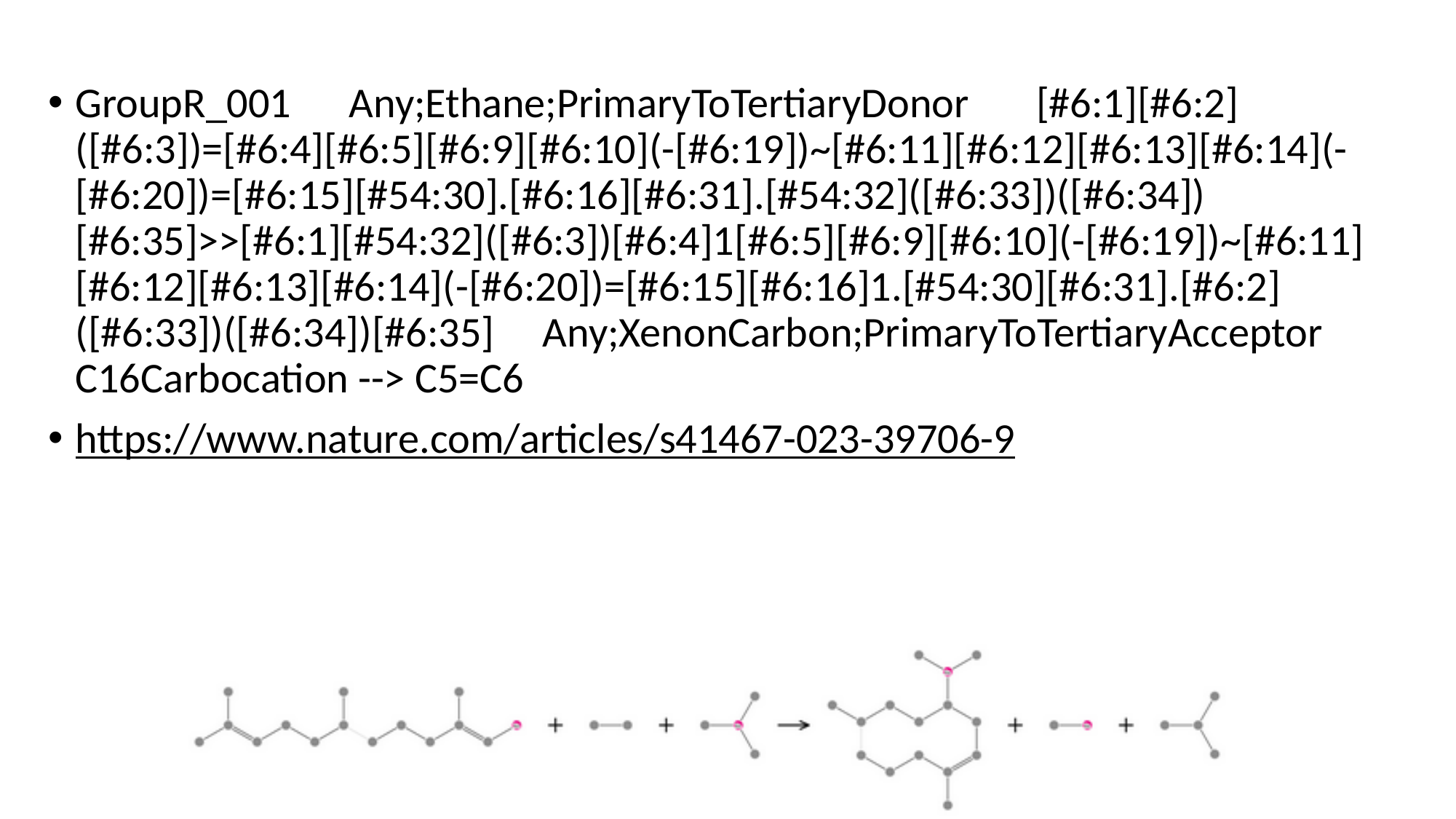

GroupR_001 Any;Ethane;PrimaryToTertiaryDonor [#6:1][#6:2]([#6:3])=[#6:4][#6:5][#6:9][#6:10](-[#6:19])~[#6:11][#6:12][#6:13][#6:14](-[#6:20])=[#6:15][#54:30].[#6:16][#6:31].[#54:32]([#6:33])([#6:34])[#6:35]>>[#6:1][#54:32]([#6:3])[#6:4]1[#6:5][#6:9][#6:10](-[#6:19])~[#6:11][#6:12][#6:13][#6:14](-[#6:20])=[#6:15][#6:16]1.[#54:30][#6:31].[#6:2]([#6:33])([#6:34])[#6:35] Any;XenonCarbon;PrimaryToTertiaryAcceptor C16Carbocation --> C5=C6
https://www.nature.com/articles/s41467-023-39706-9

### Slide 66
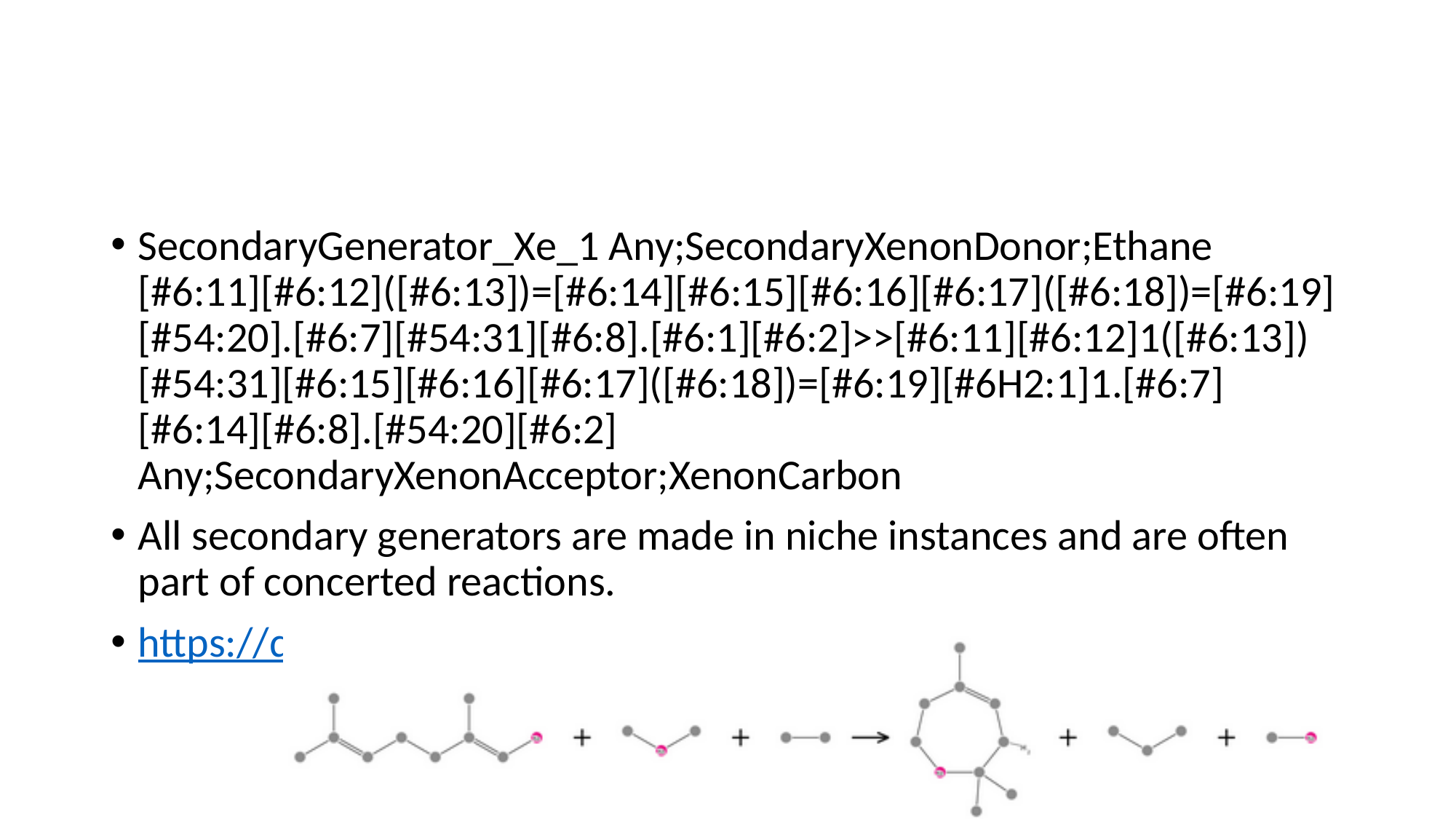

#
SecondaryGenerator_Xe_1 Any;SecondaryXenonDonor;Ethane [#6:11][#6:12]([#6:13])=[#6:14][#6:15][#6:16][#6:17]([#6:18])=[#6:19][#54:20].[#6:7][#54:31][#6:8].[#6:1][#6:2]>>[#6:11][#6:12]1([#6:13])[#54:31][#6:15][#6:16][#6:17]([#6:18])=[#6:19][#6H2:1]1.[#6:7][#6:14][#6:8].[#54:20][#6:2] Any;SecondaryXenonAcceptor;XenonCarbon
All secondary generators are made in niche instances and are often part of concerted reactions.
https://doi.org/10.1016/B978-0-444-63602-7.00005-9

### Slide 67
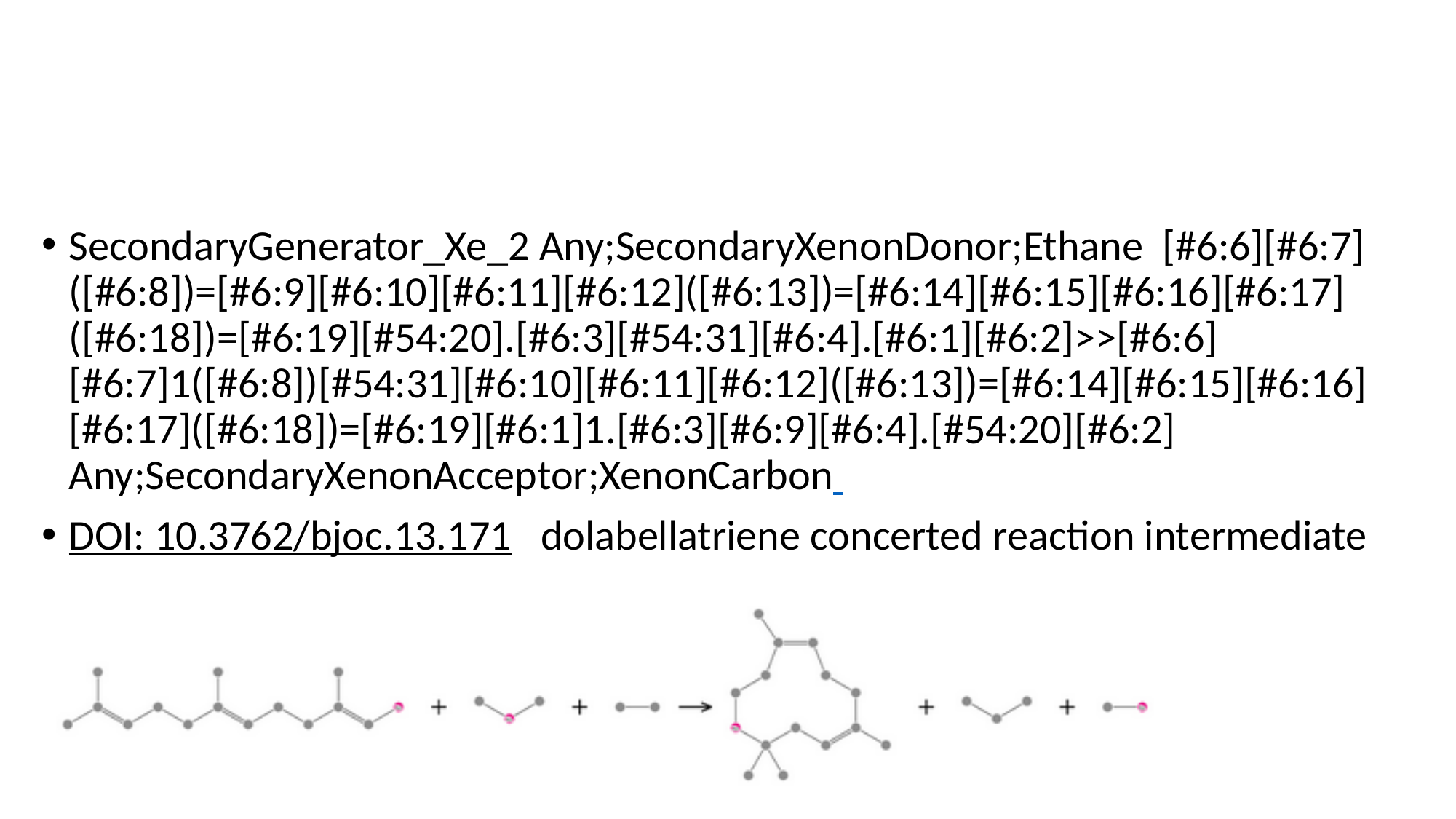

SecondaryGenerator_Xe_2 Any;SecondaryXenonDonor;Ethane [#6:6][#6:7]([#6:8])=[#6:9][#6:10][#6:11][#6:12]([#6:13])=[#6:14][#6:15][#6:16][#6:17]([#6:18])=[#6:19][#54:20].[#6:3][#54:31][#6:4].[#6:1][#6:2]>>[#6:6][#6:7]1([#6:8])[#54:31][#6:10][#6:11][#6:12]([#6:13])=[#6:14][#6:15][#6:16][#6:17]([#6:18])=[#6:19][#6:1]1.[#6:3][#6:9][#6:4].[#54:20][#6:2] Any;SecondaryXenonAcceptor;XenonCarbon
DOI: 10.3762/bjoc.13.171 dolabellatriene concerted reaction intermediate

### Slide 68
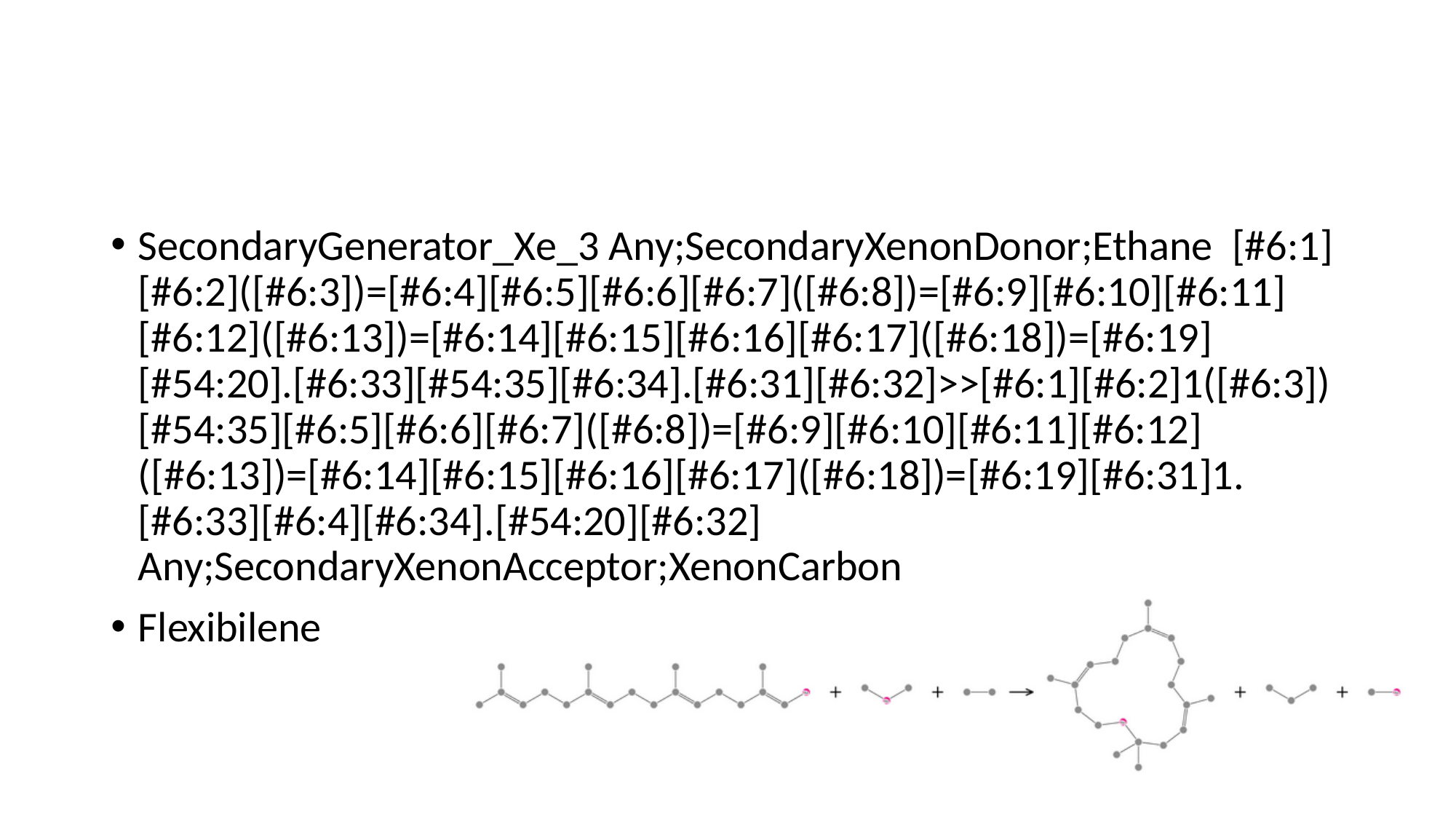

#
SecondaryGenerator_Xe_3 Any;SecondaryXenonDonor;Ethane [#6:1][#6:2]([#6:3])=[#6:4][#6:5][#6:6][#6:7]([#6:8])=[#6:9][#6:10][#6:11][#6:12]([#6:13])=[#6:14][#6:15][#6:16][#6:17]([#6:18])=[#6:19][#54:20].[#6:33][#54:35][#6:34].[#6:31][#6:32]>>[#6:1][#6:2]1([#6:3])[#54:35][#6:5][#6:6][#6:7]([#6:8])=[#6:9][#6:10][#6:11][#6:12]([#6:13])=[#6:14][#6:15][#6:16][#6:17]([#6:18])=[#6:19][#6:31]1.[#6:33][#6:4][#6:34].[#54:20][#6:32] Any;SecondaryXenonAcceptor;XenonCarbon
Flexibilene

### Slide 69
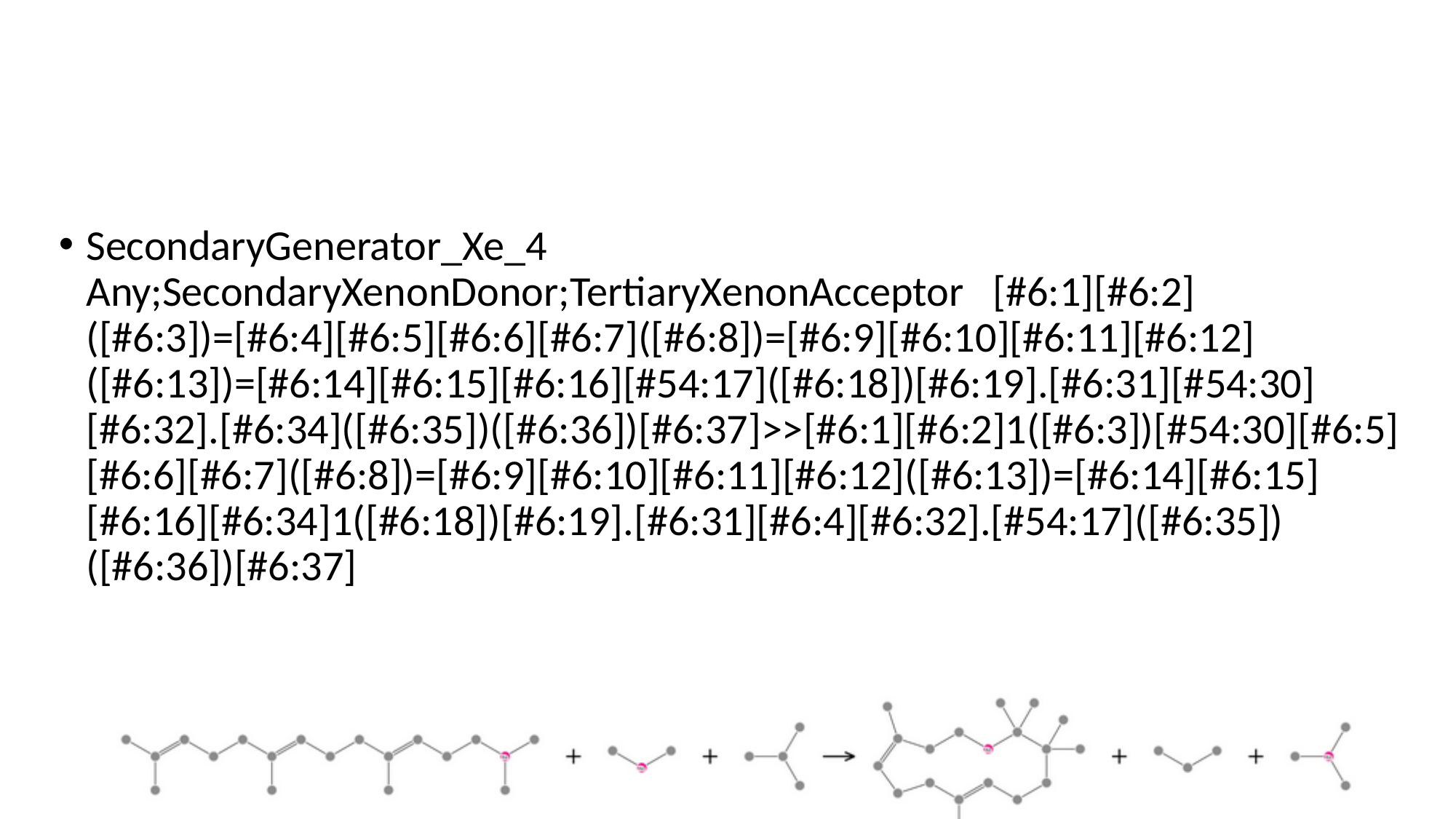

#
SecondaryGenerator_Xe_4 Any;SecondaryXenonDonor;TertiaryXenonAcceptor [#6:1][#6:2]([#6:3])=[#6:4][#6:5][#6:6][#6:7]([#6:8])=[#6:9][#6:10][#6:11][#6:12]([#6:13])=[#6:14][#6:15][#6:16][#54:17]([#6:18])[#6:19].[#6:31][#54:30][#6:32].[#6:34]([#6:35])([#6:36])[#6:37]>>[#6:1][#6:2]1([#6:3])[#54:30][#6:5][#6:6][#6:7]([#6:8])=[#6:9][#6:10][#6:11][#6:12]([#6:13])=[#6:14][#6:15][#6:16][#6:34]1([#6:18])[#6:19].[#6:31][#6:4][#6:32].[#54:17]([#6:35])([#6:36])[#6:37]

### Slide 70
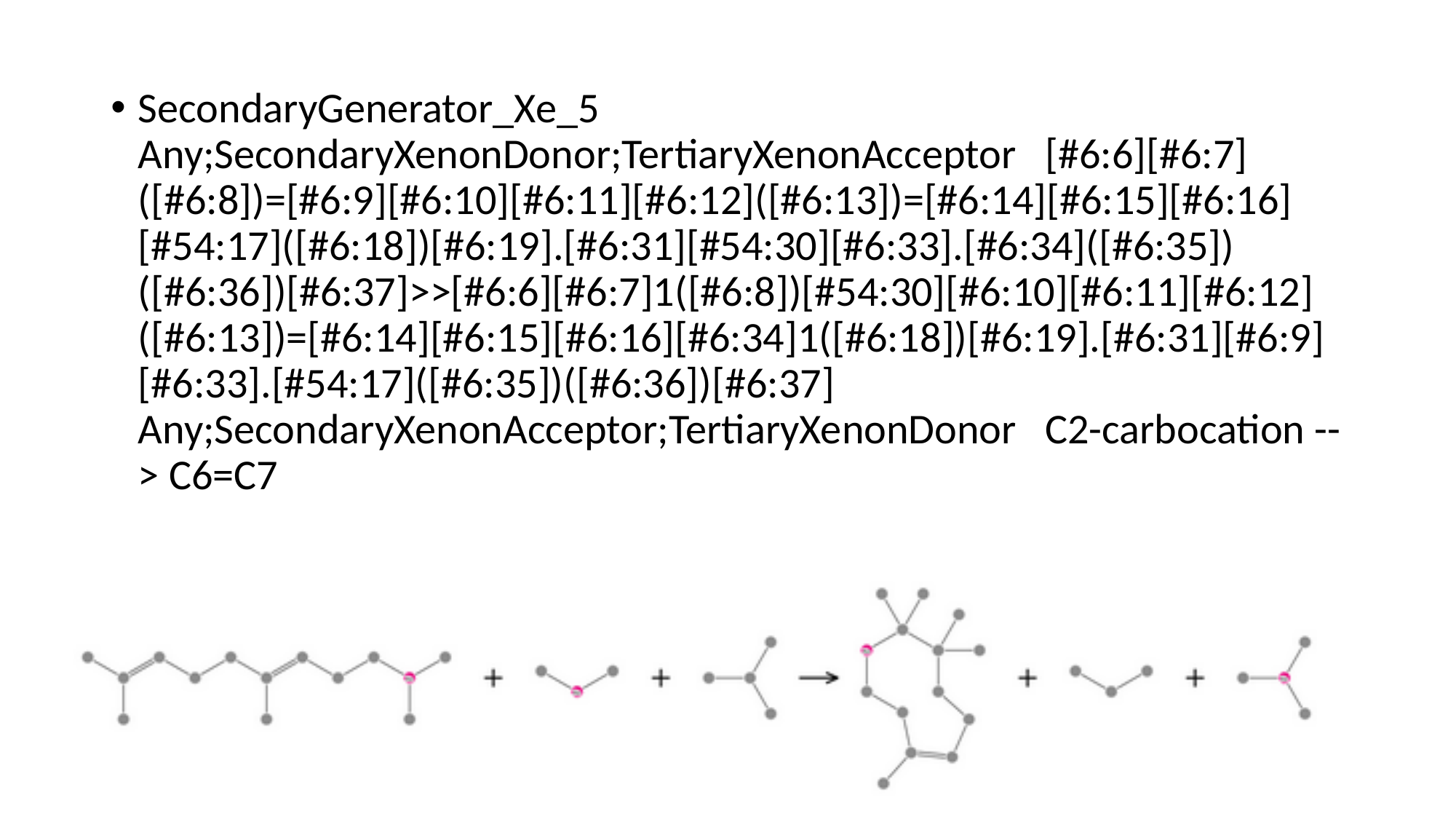

SecondaryGenerator_Xe_5 Any;SecondaryXenonDonor;TertiaryXenonAcceptor [#6:6][#6:7]([#6:8])=[#6:9][#6:10][#6:11][#6:12]([#6:13])=[#6:14][#6:15][#6:16][#54:17]([#6:18])[#6:19].[#6:31][#54:30][#6:33].[#6:34]([#6:35])([#6:36])[#6:37]>>[#6:6][#6:7]1([#6:8])[#54:30][#6:10][#6:11][#6:12]([#6:13])=[#6:14][#6:15][#6:16][#6:34]1([#6:18])[#6:19].[#6:31][#6:9][#6:33].[#54:17]([#6:35])([#6:36])[#6:37] Any;SecondaryXenonAcceptor;TertiaryXenonDonor C2-carbocation --> C6=C7

### Slide 71
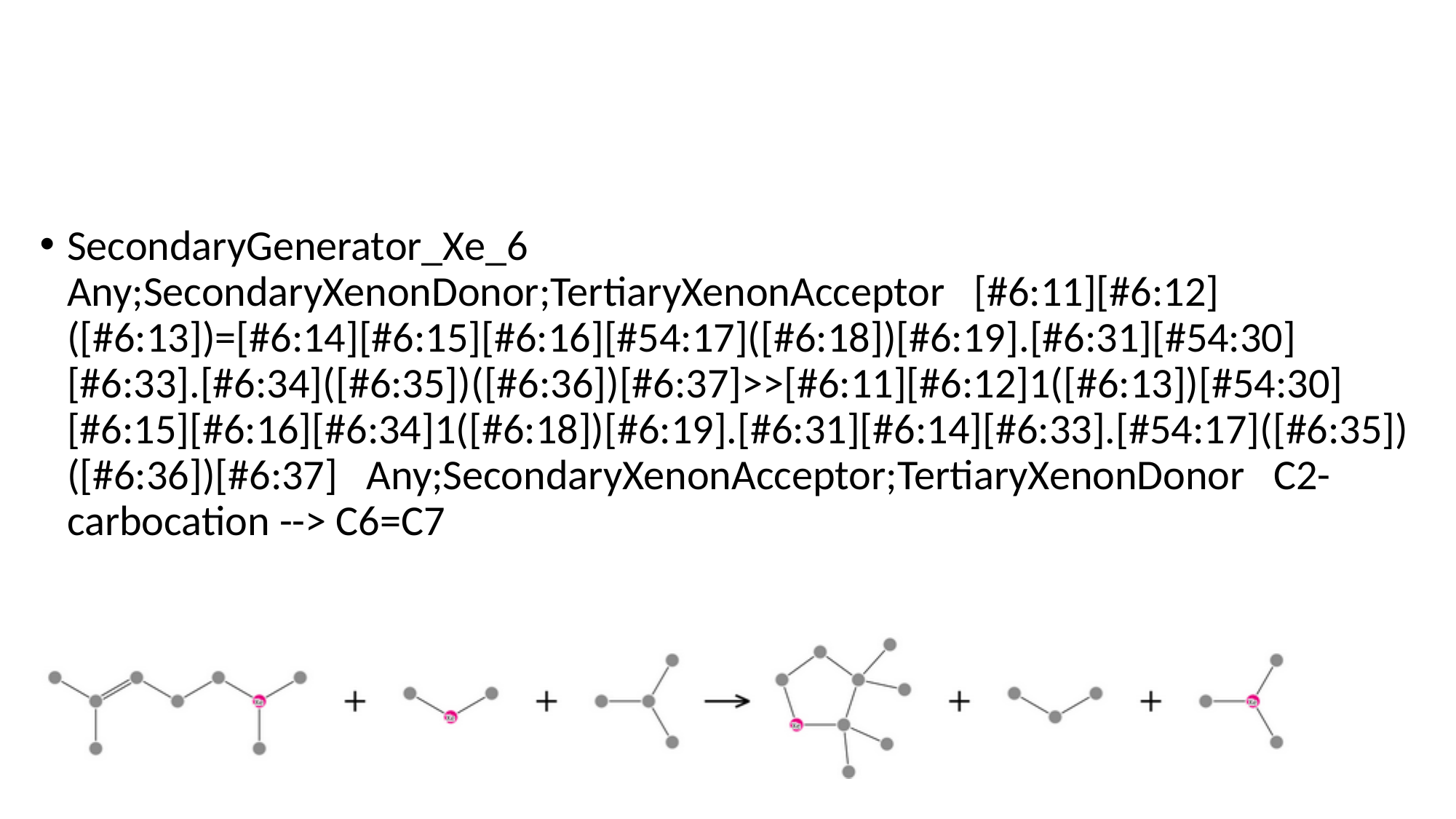

#
SecondaryGenerator_Xe_6 Any;SecondaryXenonDonor;TertiaryXenonAcceptor [#6:11][#6:12]([#6:13])=[#6:14][#6:15][#6:16][#54:17]([#6:18])[#6:19].[#6:31][#54:30][#6:33].[#6:34]([#6:35])([#6:36])[#6:37]>>[#6:11][#6:12]1([#6:13])[#54:30][#6:15][#6:16][#6:34]1([#6:18])[#6:19].[#6:31][#6:14][#6:33].[#54:17]([#6:35])([#6:36])[#6:37] Any;SecondaryXenonAcceptor;TertiaryXenonDonor C2-carbocation --> C6=C7

### Slide 72
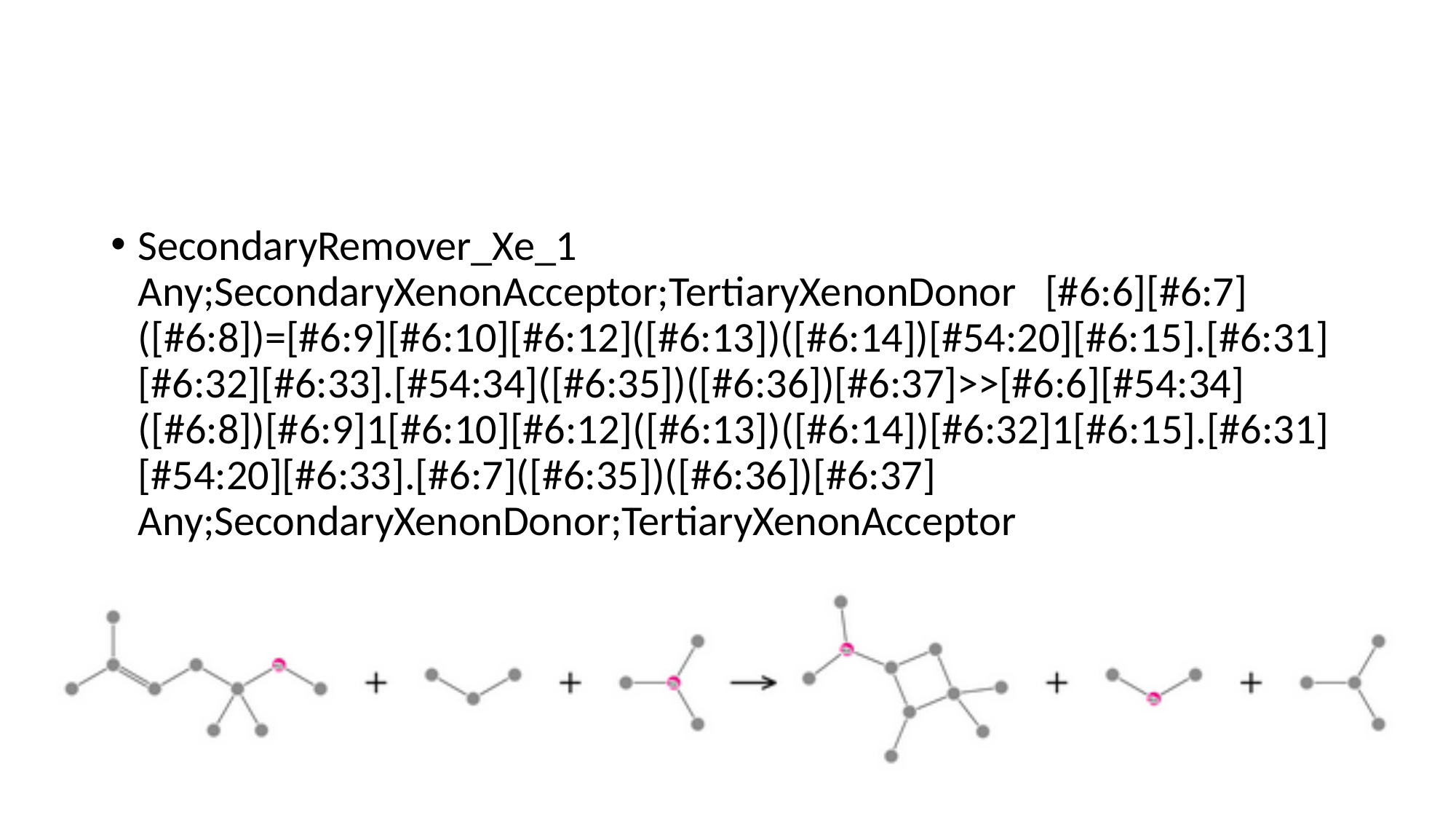

#
SecondaryRemover_Xe_1 Any;SecondaryXenonAcceptor;TertiaryXenonDonor [#6:6][#6:7]([#6:8])=[#6:9][#6:10][#6:12]([#6:13])([#6:14])[#54:20][#6:15].[#6:31][#6:32][#6:33].[#54:34]([#6:35])([#6:36])[#6:37]>>[#6:6][#54:34]([#6:8])[#6:9]1[#6:10][#6:12]([#6:13])([#6:14])[#6:32]1[#6:15].[#6:31][#54:20][#6:33].[#6:7]([#6:35])([#6:36])[#6:37] Any;SecondaryXenonDonor;TertiaryXenonAcceptor

### Slide 73
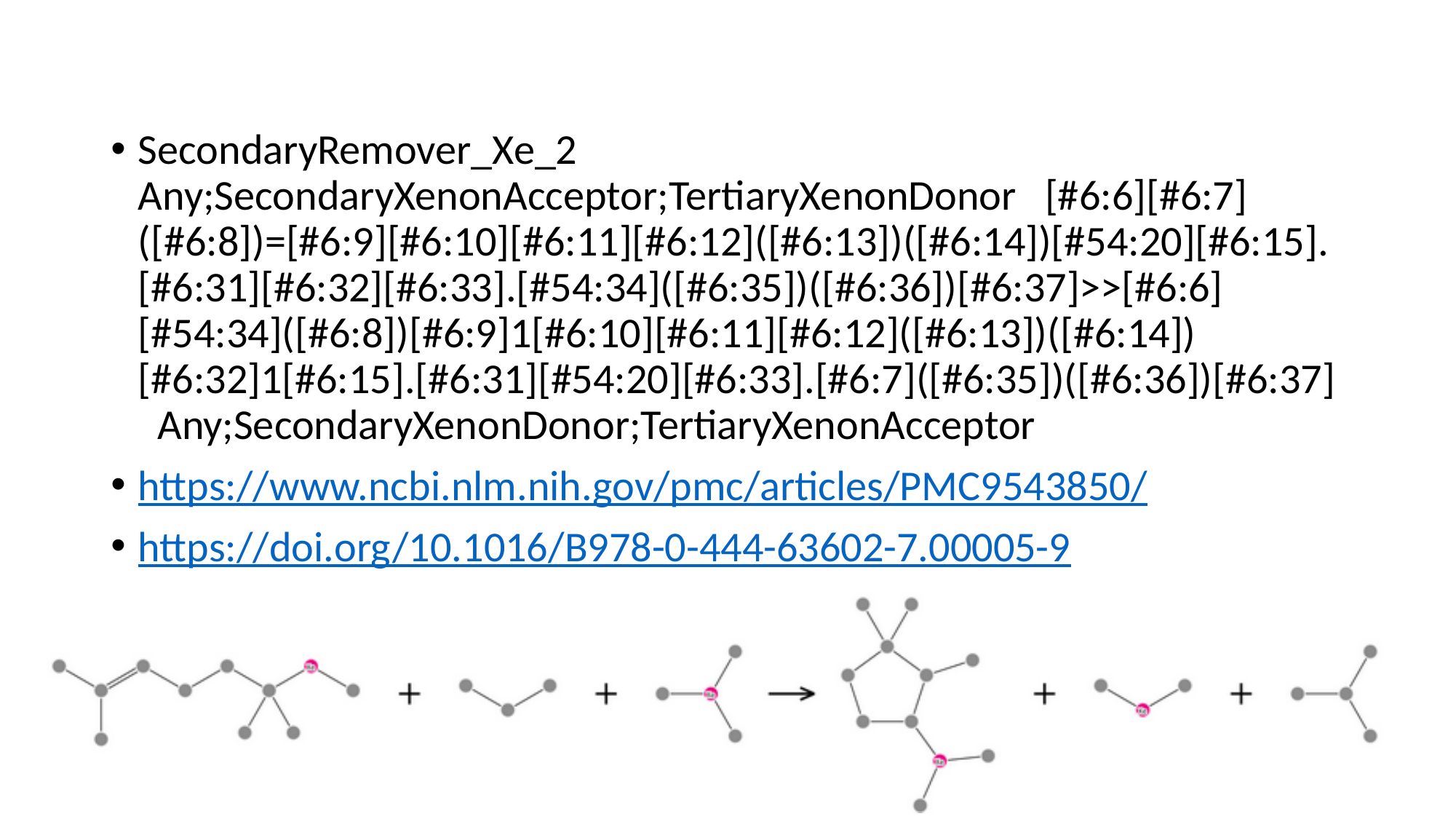

SecondaryRemover_Xe_2 Any;SecondaryXenonAcceptor;TertiaryXenonDonor [#6:6][#6:7]([#6:8])=[#6:9][#6:10][#6:11][#6:12]([#6:13])([#6:14])[#54:20][#6:15].[#6:31][#6:32][#6:33].[#54:34]([#6:35])([#6:36])[#6:37]>>[#6:6][#54:34]([#6:8])[#6:9]1[#6:10][#6:11][#6:12]([#6:13])([#6:14])[#6:32]1[#6:15].[#6:31][#54:20][#6:33].[#6:7]([#6:35])([#6:36])[#6:37] Any;SecondaryXenonDonor;TertiaryXenonAcceptor
https://www.ncbi.nlm.nih.gov/pmc/articles/PMC9543850/
https://doi.org/10.1016/B978-0-444-63602-7.00005-9

### Slide 74
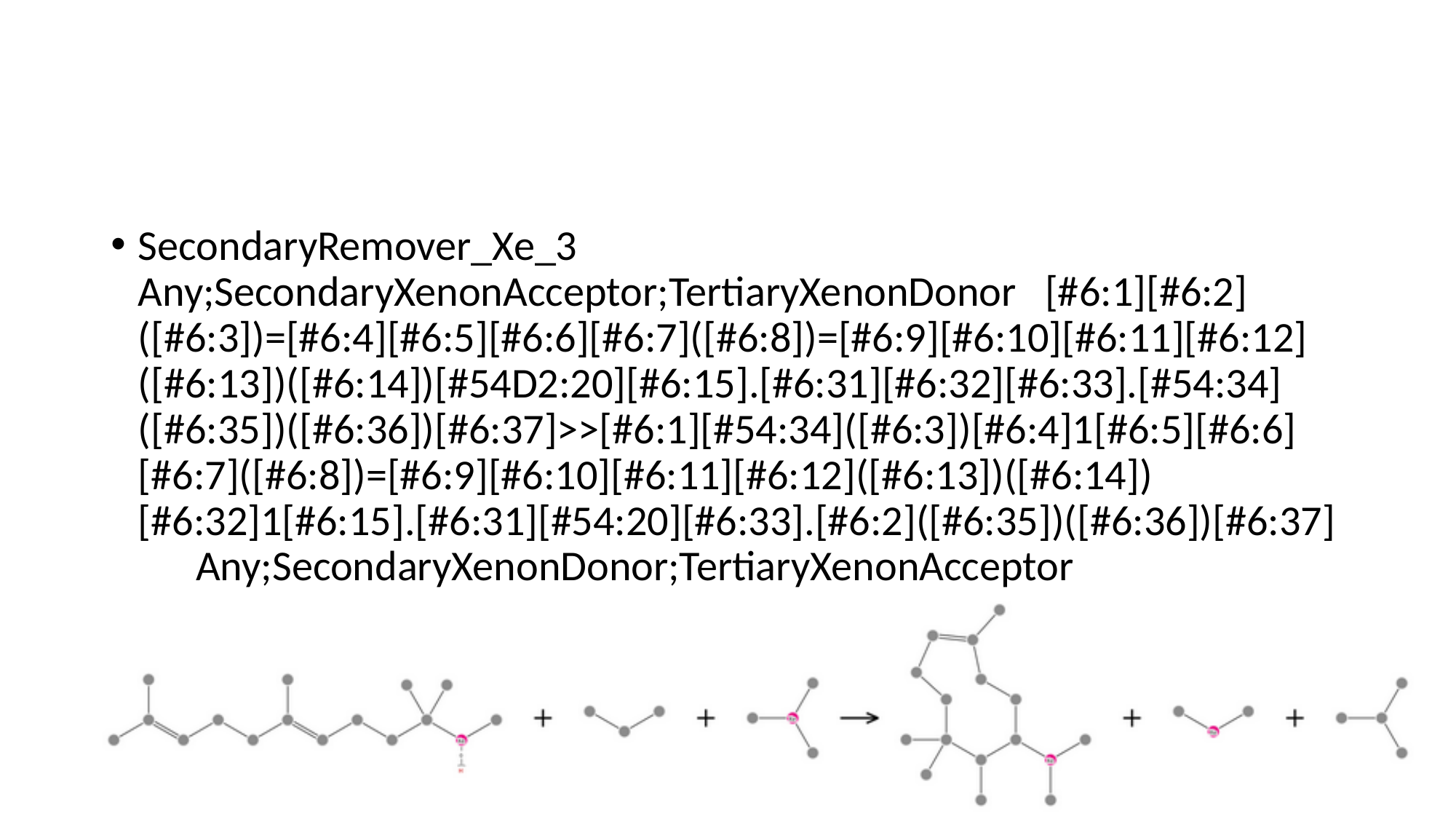

#
SecondaryRemover_Xe_3 Any;SecondaryXenonAcceptor;TertiaryXenonDonor [#6:1][#6:2]([#6:3])=[#6:4][#6:5][#6:6][#6:7]([#6:8])=[#6:9][#6:10][#6:11][#6:12]([#6:13])([#6:14])[#54D2:20][#6:15].[#6:31][#6:32][#6:33].[#54:34]([#6:35])([#6:36])[#6:37]>>[#6:1][#54:34]([#6:3])[#6:4]1[#6:5][#6:6][#6:7]([#6:8])=[#6:9][#6:10][#6:11][#6:12]([#6:13])([#6:14])[#6:32]1[#6:15].[#6:31][#54:20][#6:33].[#6:2]([#6:35])([#6:36])[#6:37] Any;SecondaryXenonDonor;TertiaryXenonAcceptor

### Slide 75
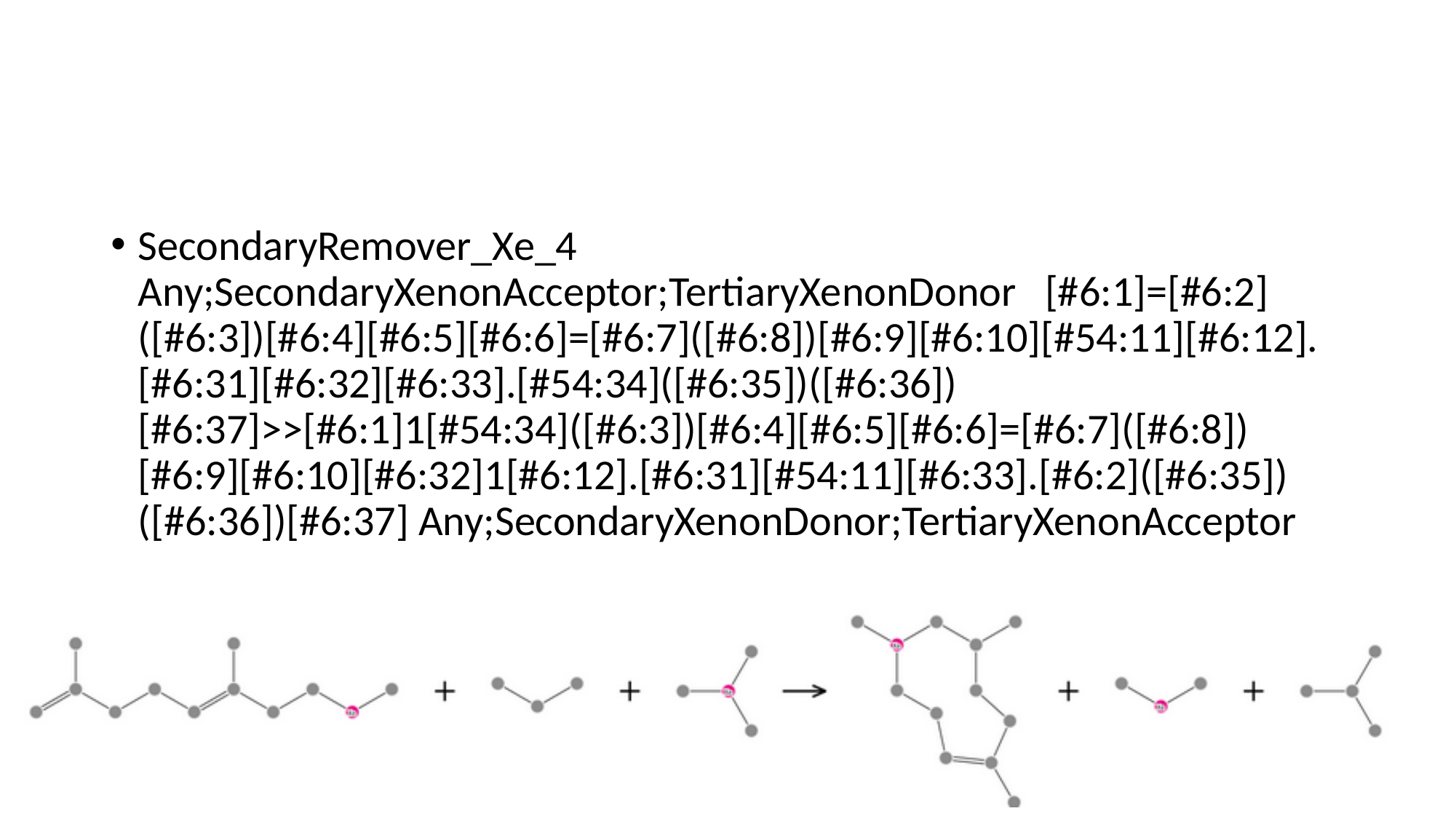

#
SecondaryRemover_Xe_4 Any;SecondaryXenonAcceptor;TertiaryXenonDonor [#6:1]=[#6:2]([#6:3])[#6:4][#6:5][#6:6]=[#6:7]([#6:8])[#6:9][#6:10][#54:11][#6:12].[#6:31][#6:32][#6:33].[#54:34]([#6:35])([#6:36])[#6:37]>>[#6:1]1[#54:34]([#6:3])[#6:4][#6:5][#6:6]=[#6:7]([#6:8])[#6:9][#6:10][#6:32]1[#6:12].[#6:31][#54:11][#6:33].[#6:2]([#6:35])([#6:36])[#6:37] Any;SecondaryXenonDonor;TertiaryXenonAcceptor

### Slide 76
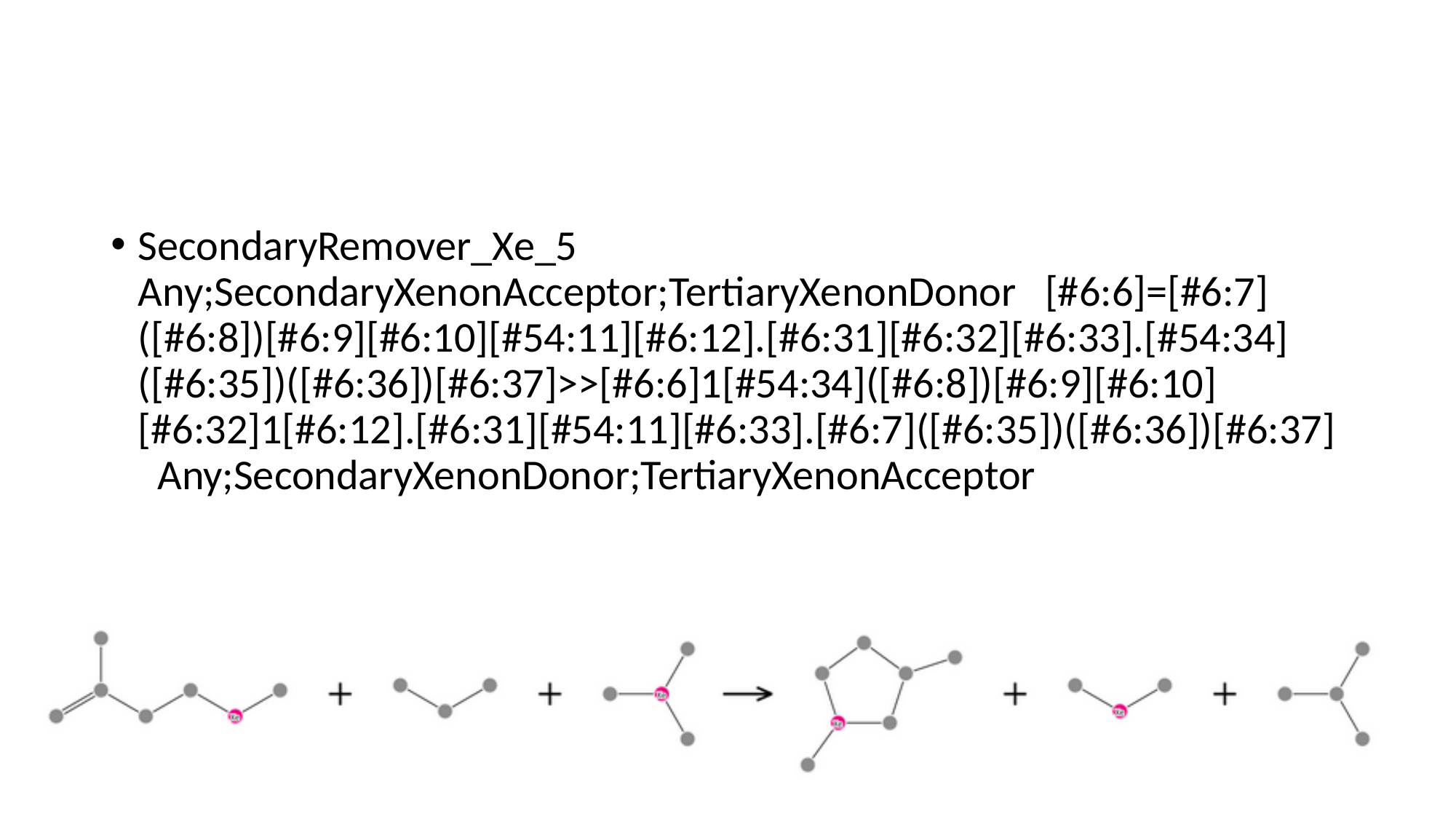

#
SecondaryRemover_Xe_5 Any;SecondaryXenonAcceptor;TertiaryXenonDonor [#6:6]=[#6:7]([#6:8])[#6:9][#6:10][#54:11][#6:12].[#6:31][#6:32][#6:33].[#54:34]([#6:35])([#6:36])[#6:37]>>[#6:6]1[#54:34]([#6:8])[#6:9][#6:10][#6:32]1[#6:12].[#6:31][#54:11][#6:33].[#6:7]([#6:35])([#6:36])[#6:37] Any;SecondaryXenonDonor;TertiaryXenonAcceptor

### Slide 77
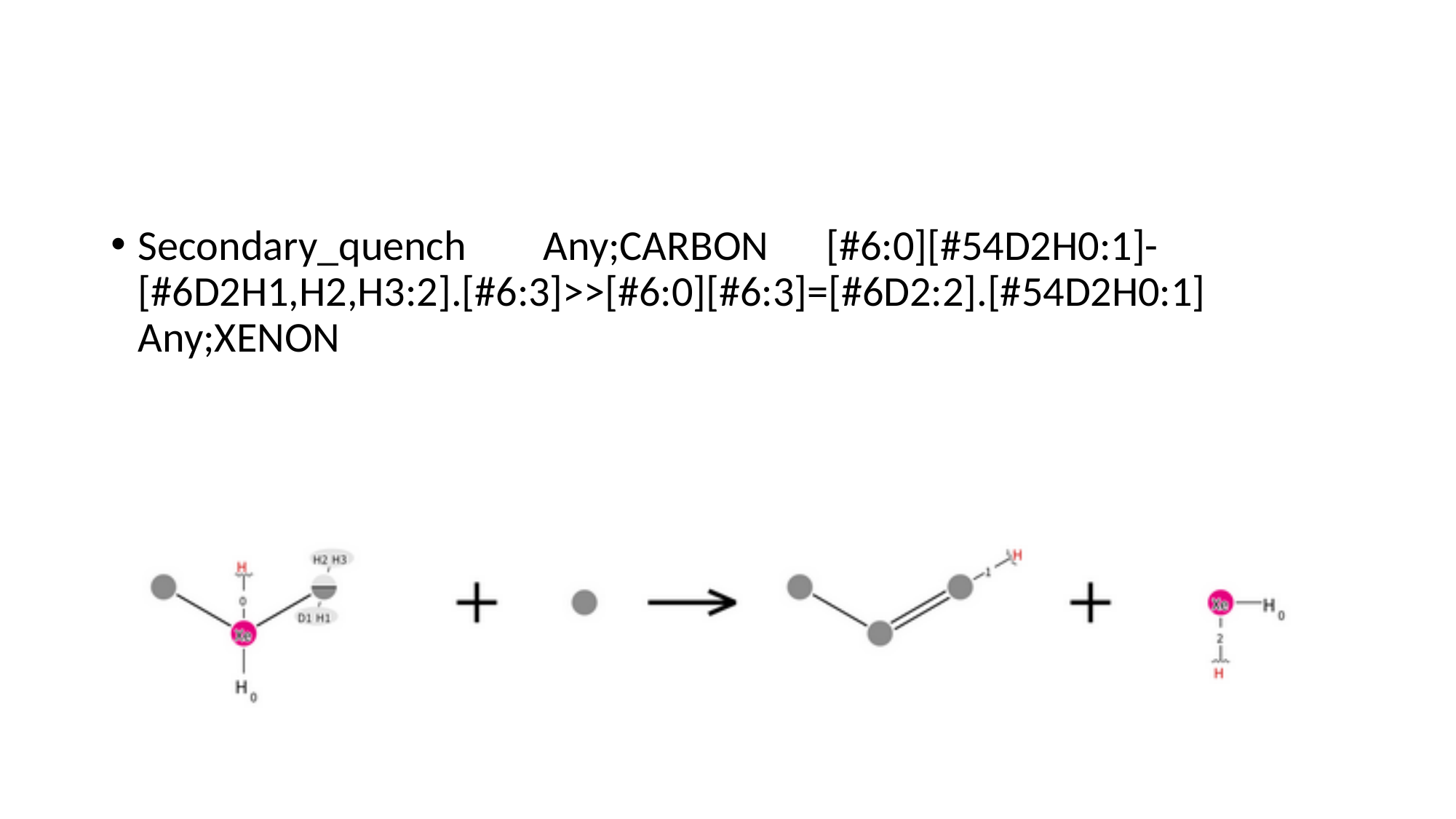

#
Secondary_quench Any;CARBON [#6:0][#54D2H0:1]-[#6D2H1,H2,H3:2].[#6:3]>>[#6:0][#6:3]=[#6D2:2].[#54D2H0:1] Any;XENON

### Slide 78
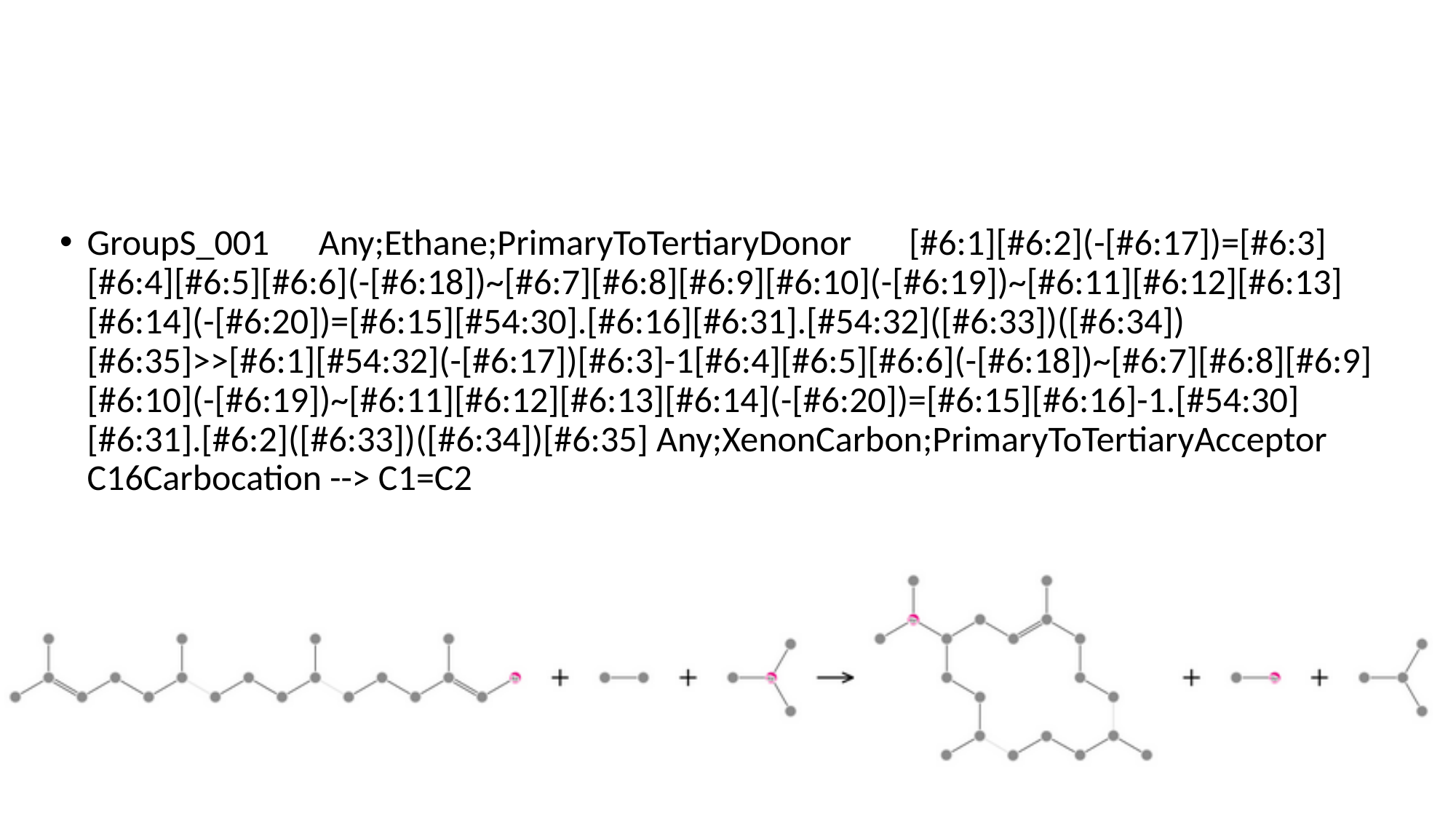

#
GroupS_001 Any;Ethane;PrimaryToTertiaryDonor [#6:1][#6:2](-[#6:17])=[#6:3][#6:4][#6:5][#6:6](-[#6:18])~[#6:7][#6:8][#6:9][#6:10](-[#6:19])~[#6:11][#6:12][#6:13][#6:14](-[#6:20])=[#6:15][#54:30].[#6:16][#6:31].[#54:32]([#6:33])([#6:34])[#6:35]>>[#6:1][#54:32](-[#6:17])[#6:3]-1[#6:4][#6:5][#6:6](-[#6:18])~[#6:7][#6:8][#6:9][#6:10](-[#6:19])~[#6:11][#6:12][#6:13][#6:14](-[#6:20])=[#6:15][#6:16]-1.[#54:30][#6:31].[#6:2]([#6:33])([#6:34])[#6:35] Any;XenonCarbon;PrimaryToTertiaryAcceptor C16Carbocation --> C1=C2

### Slide 79
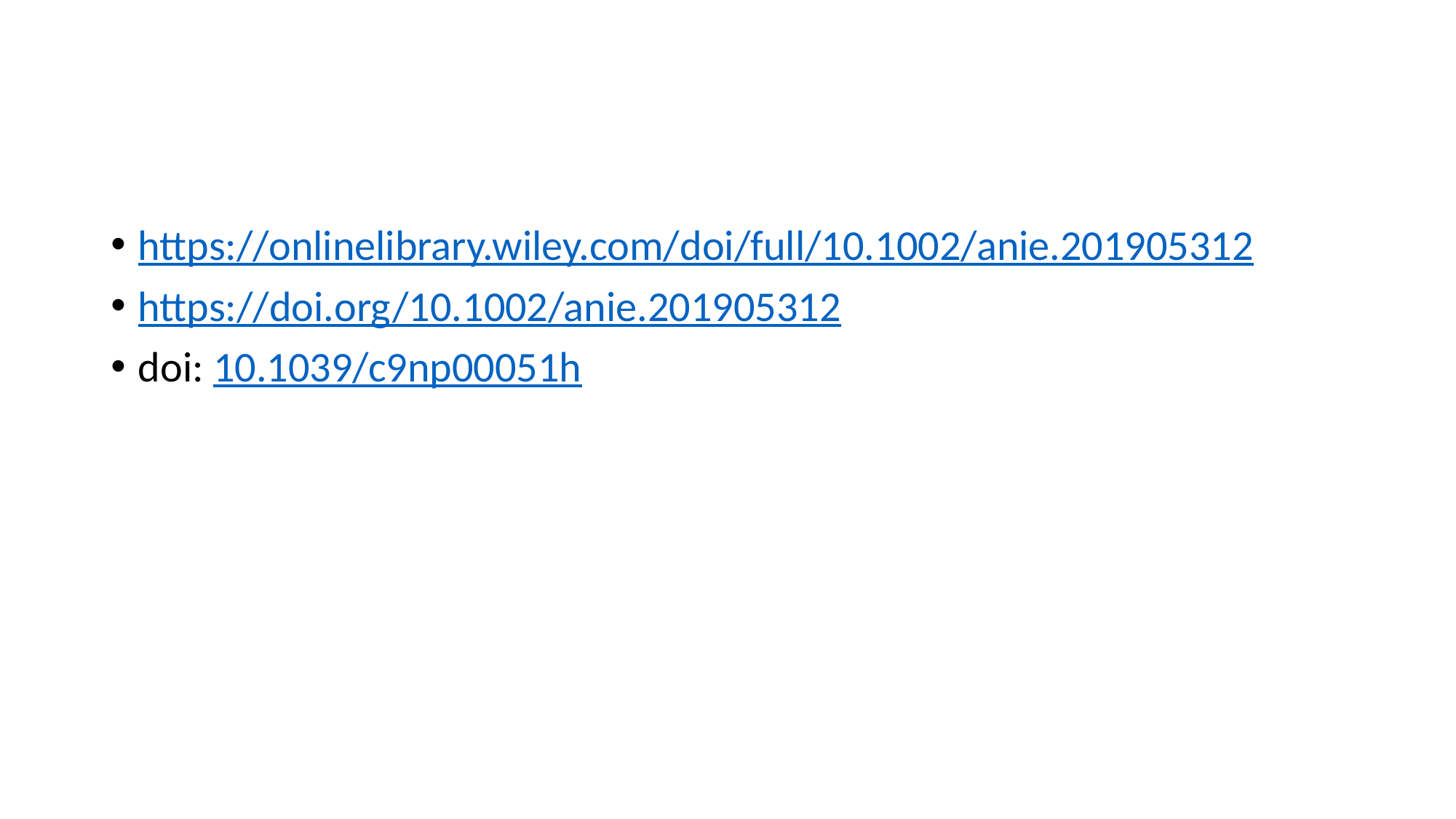

#
https://onlinelibrary.wiley.com/doi/full/10.1002/anie.201905312
https://doi.org/10.1002/anie.201905312
doi: 10.1039/c9np00051h
