## Supplementary figures and images for "Rule-Based Deconstruction and Reconstruction of Diterpene Libraries: Categorizing Patterns & Unravelling the Structural Landscape"

### Supp.2g.Terokit_Diterpene_Skeleton_Abundance_Distribution.png

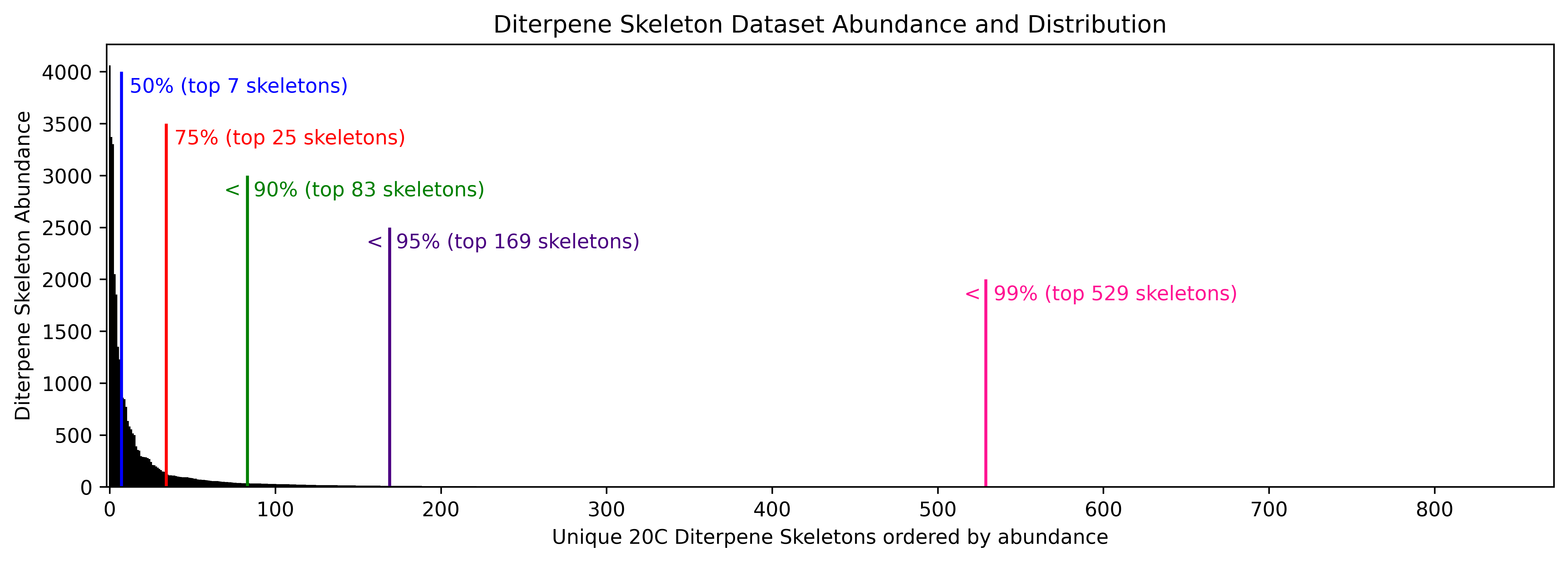

### Supp.2h.Terokit_Diterpene_Deconstructed_Skeleton_Carbon_Distribution.png

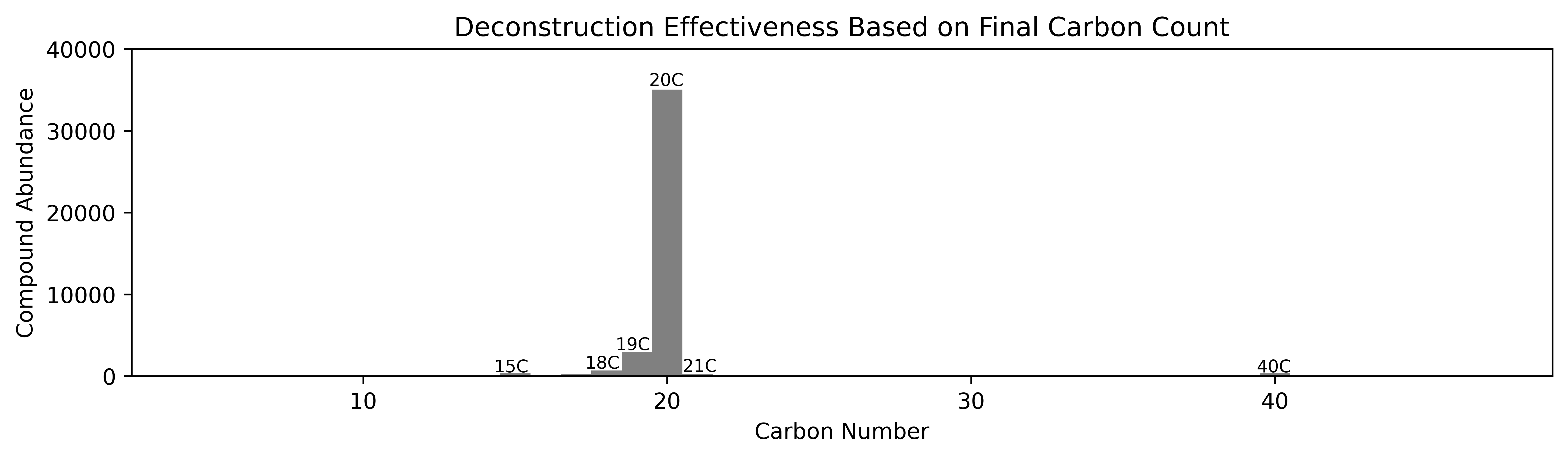

### Supp.2i.10X_Zoomed_Terokit_Diterpene_Deconstructed_Skeleton_Carbon_Distribution.png

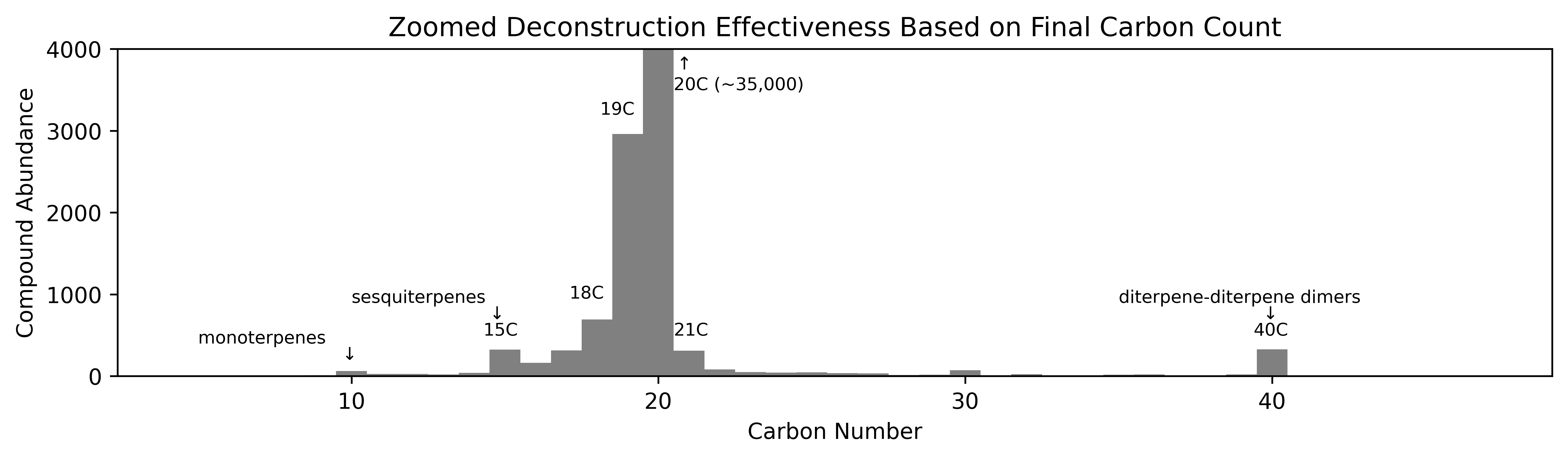

### Supp.8.SpeciesPerFamily_count_V_Compound_count.png

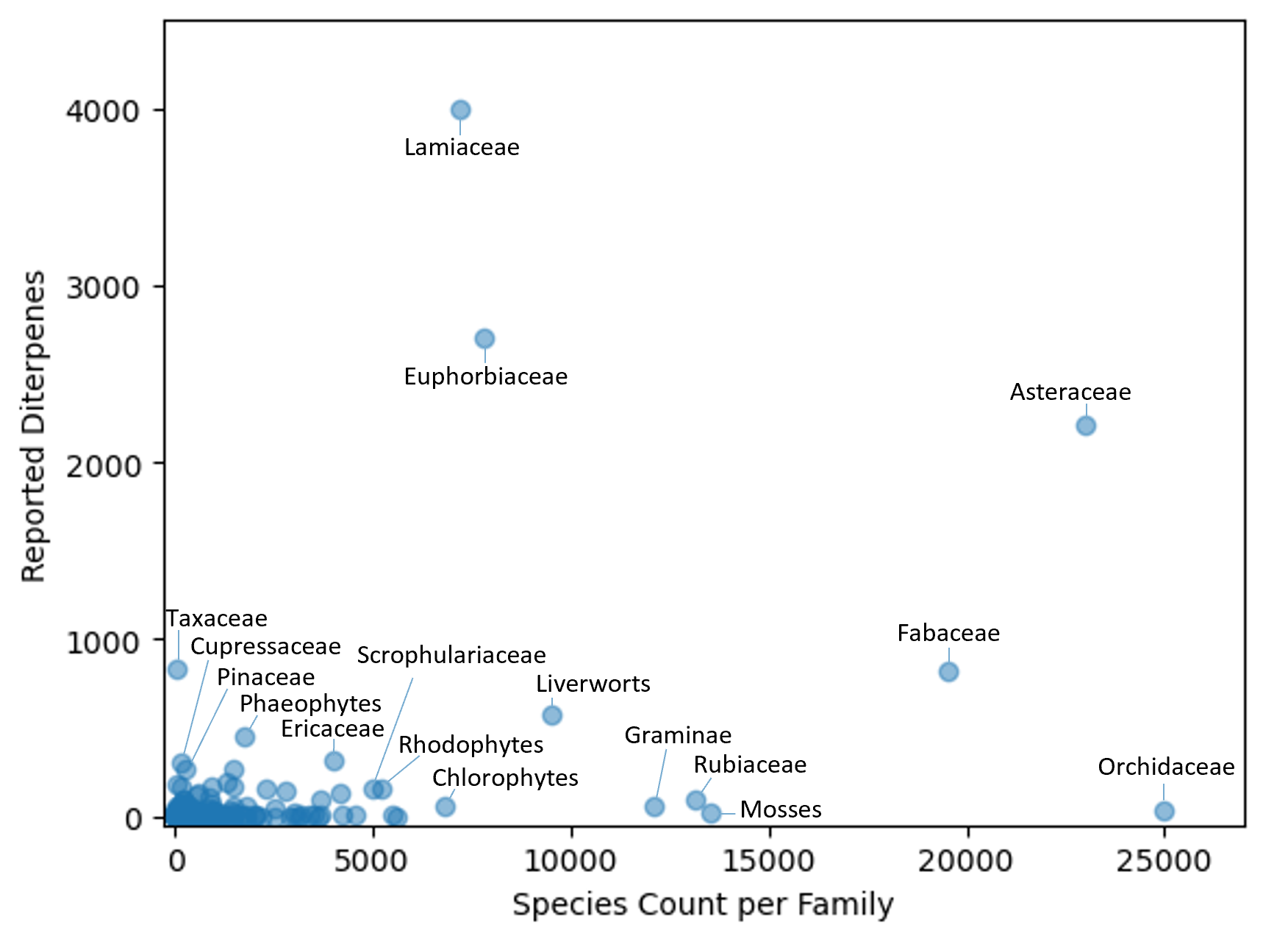

### Supp.9.Predicted_TaxaneRelated_Mechanisms.png

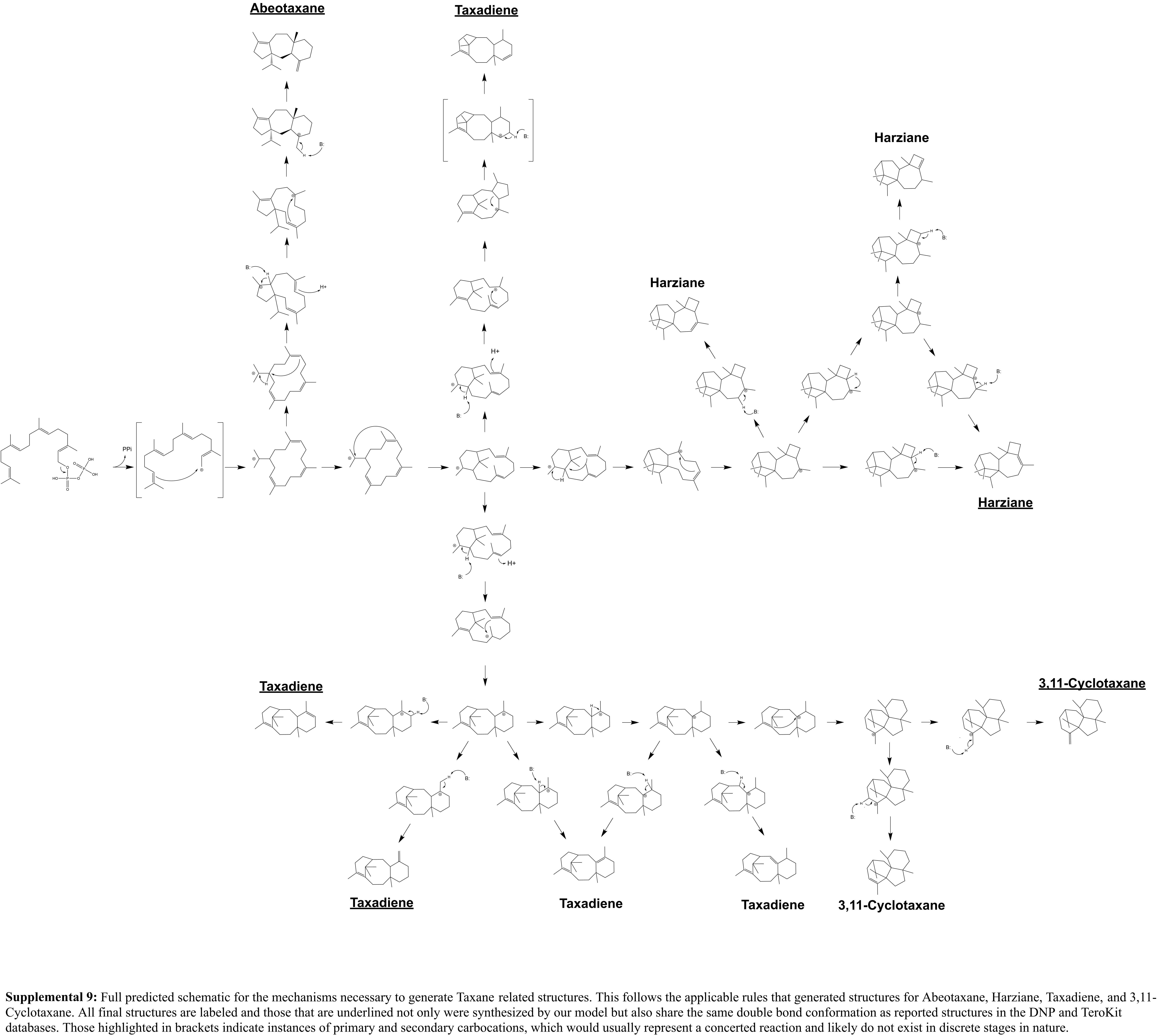
